# ER-to-Golgi Trafficking is a Nutrient-Sensitive Checkpoint Linking Glucose Starvation to Cell Surface Remodeling

**DOI:** 10.1101/2025.10.31.685804

**Authors:** Joung Hyuck Joo, William Kasberg, Spencer Douglas, Uwemedimo Udoh, Alexandre Carisey, James Messing, Yong-Dong Wang, Shilpa Narina, Shondra M. Pruett-Miller, Myriam Labelle, Jennifer Lippincott-Schwartz, Chi-Lun Chang, Mondira Kundu

## Abstract

Cancer cells adapt to nutrient stress by remodeling the repertoire of proteins on their surface, enabling survival and progression under starvation conditions. However, the molecular mechanisms by which nutrient cues reshape the cell surface proteome to influence cell behavior remain largely unresolved. Here, we show that acute glucose starvation, but not amino acid deprivation or mTOR inhibition, selectively impairs ER-to-Golgi export of specific cargoes, such as E-cadherin, in a SEC24C-dependent manner. Quantitative cell surface proteomics reveal that glucose deprivation remodels the cell surface proteome, notably reducing surface expression of key adhesion molecules. This nutrient-sensitive reprogramming enhances cell migration *in vitro* and promotes metastasis *in vivo*. Mechanistically, we show that AMPK and ULK1 signaling orchestrate this process independent of autophagy, with ULK1-mediated phosphorylation of SEC31A driving SEC24C-dependent COPII reorganization. These findings establish ER-to-Golgi trafficking as a nutrient-sensitive regulatory node that links metabolic stress to cell surface remodeling and metastatic potential.

## Introduction

Eukaryotic cells possess a remarkable capacity to adapt to dynamic metabolic and environmental conditions, an ability that is pushed to extremes in cancer as transformed cells must survive and proliferate within nutrient-poor, hypoxic microenvironments^1, 2^. Adaptation to metabolic stress involves not only rewiring of intracellular metabolism and signaling^3, 4^, but also dynamic remodeling of the cell surface proteome^5–7^. Nutrient and metabolic stresses, including glucose starvation and hypoxia, are well known to induce broad remodeling of the cell surface proteome, affecting transporters, adhesion molecules, and signaling receptors in ways that can influence cancer progression^5, 8–10^. However, the precise molecular mechanisms by which nutrient cues engage the secretory machinery and reshape the cell surface proteome remain poorly understood. Addressing this gap in our knowledge is critical not only for understanding cancer progression, but also for answering fundamental questions in cell biology, such as how metabolic cues are transmitted through the secretory pathway to shape the cell surface proteome, and how these changes in surface composition, in turn, influence core cellular behaviors.

A key step in remodeling the cell surface proteome is the export of newly synthesized proteins from the endoplasmic reticulum (ER) to the Golgi apparatus, a process mediated by the conserved COPII (coat protein complex II) machinery. COPII components assemble at specialized regions of the ER membrane called ER exit sites (ERES), where they select cargo and shape the membrane to form transport carriers^11, 12^. Vesicle formation begins when the small GTPase Sar1 is activated by its guanine nucleotide exchange factor, Sec12, which is embedded in the ER membrane^13, 14^. Once activated, GTP-bound Sar1 inserts into the membrane and recruits the Sec23–Sec24 inner coat complex^15–19^, followed by the Sec13–Sec31 outer coat complex^20–22^. Sec31 enhances the activity of Sec23, which acts as a GTPase-activating protein for Sar1, promoting GTP hydrolysis^23–25^. This step is crucial for regulating vesicle size and ensuring timely release of the COPII coat, a prerequisite for subsequent tethering and membrane fusion events during cargo delivery^26–28^. COPII’s assembly and function are further coordinated by Sec16, a large scaffold protein that marks and organizes ERES, facilitates COPII coat recruitment, and modulates Sar1 GTPase activity by counteracting Sec31-driven GTP hydrolysis^29–32^.

Recent work has demonstrated that COPII function can by modulated by a variety of signaling inputs, with multiple kinases and post-translational modifications tuning ER export efficiency^33–36^. Kinases such as ERK7 (extracellular signal-regulated kinase 7), CK2 (casein kinase 2), and MAPKs (mitogen-activated protein kinases), LTK (leukocyte receptor tyrosine kinase) have been shown to phosphorylate COPII components and influence secretory flux^35, 37–39^, while additional post-translational modifications, including O-GlcNAcylation and ubiquitination, of COPII subunits further remodel COPII machinery, thereby impacting vesicle formation and cargo selection^34^. Although nutrient deprivation in mammalian cells has been shown to slow COPII subunit recruitment and turnover at ER subdomains^33^, the molecular mechanisms by which nutrient availability is communicated to the COPII machinery to regulate ER export, and thereby shape the cell surface proteome and cellular behavior, remain poorly understood.

Here, we systematically investigate the effects of acute nutrient starvation on ER-to-Golgi trafficking and the cell surface proteome. We find that glucose starvation (but not amino acid starvation) disrupts ER export and cell surface display of cell-cell adhesion molecules such as E-cadherin, thereby promoting cell migration and metastatic potential. This effect depends on SEC24C, one of four mammalian SEC24 isoforms responsible for cargo selection during COPII-mediated transport^12^. We further demonstrate that this nutrient-sensitive, SEC24C-dependent process is regulated by the energy-sensing kinase AMPK (adenosine monophosphate-activated kinase) and the autophagy-initiating kinase ULK1 (Unc-51-like kinase 1), independent of conventional autophagy pathways. Finally, we identify SEC31 as a target of ULK1 and show that its phosphorylation is required for the SEC24C-mediated inhibition of ER-to-Golgi trafficking during starvation. Together, our findings identify ER-to-Golgi trafficking as a key regulatory hub linking metabolic stress to cell surface dynamics and reveal how nutrient availability can directly impact cancer cell invasiveness by regulating this pathway.

## Results

### Glucose, but Not Amino Acid, Starvation Selectively Inhibits ER-to-Golgi Trafficking of Specific Cargoes

To investigate how nutrient availability influences protein trafficking from the ER to the Golgi apparatus, we utilized the Retention Using Selective Hooks (RUSH) system, a well-established approach to synchronizing and visualizing the transport of specific cargo proteins^40^. In this system, cargo proteins are initially retained in the ER via a streptavidin “hook” and are rapidly released upon addition of biotin, allowing real-time monitoring of their trafficking.

We examined the ER-to-Golgi transport of three distinct cargoes in U2OS cells: E-cadherin, GPI (GPI-anchored proteins), and VSVG (vesicular stomatitis virus G protein). As expected, the addition of biotin to the culture media triggered robust trafficking of all three cargoes to the Golgi (Fig. 1a–i). However, when cells were subjected to glucose starvation for 1 h prior to biotin addition, we observed a striking inhibition of E-cadherin (reduced by ∼66%) and GPI trafficking (reduced by ∼75%) to the Golgi; VSVG transport was unaffected (Fig. 1a, b, d, e, g, h). In contrast, amino acid starvation had no effect on the ER-to-Golgi trafficking of any of the three cargoes (Fig. 1c, f, i). Using E-cadherin trafficking as a model, we further tested the effect of directly inhibiting mTOR signaling with the mTOR inhibitor Torin2. Consistent with the results from amino acid starvation, Torin2 did not impair ER-to-Golgi trafficking of E-cadherin (Extended data Fig. 1a, b). To confirm that the changes in nutrient availability were functionally effective, we performed immunoblot analyses in U2OS cells. Glucose starvation resulted in the expected activation of AMPK (Extended data Fig. 1c), indicated by increased phosphorylation of acetyl-CoA carboxylase^41^. Conversely, both amino acid starvation and Torin2 treatment led to decreased phosphorylation of the ribosomal protein S6 (Extended data Fig. 1c), confirming effective mTOR inhibition^42^.

**Fig. 1:**
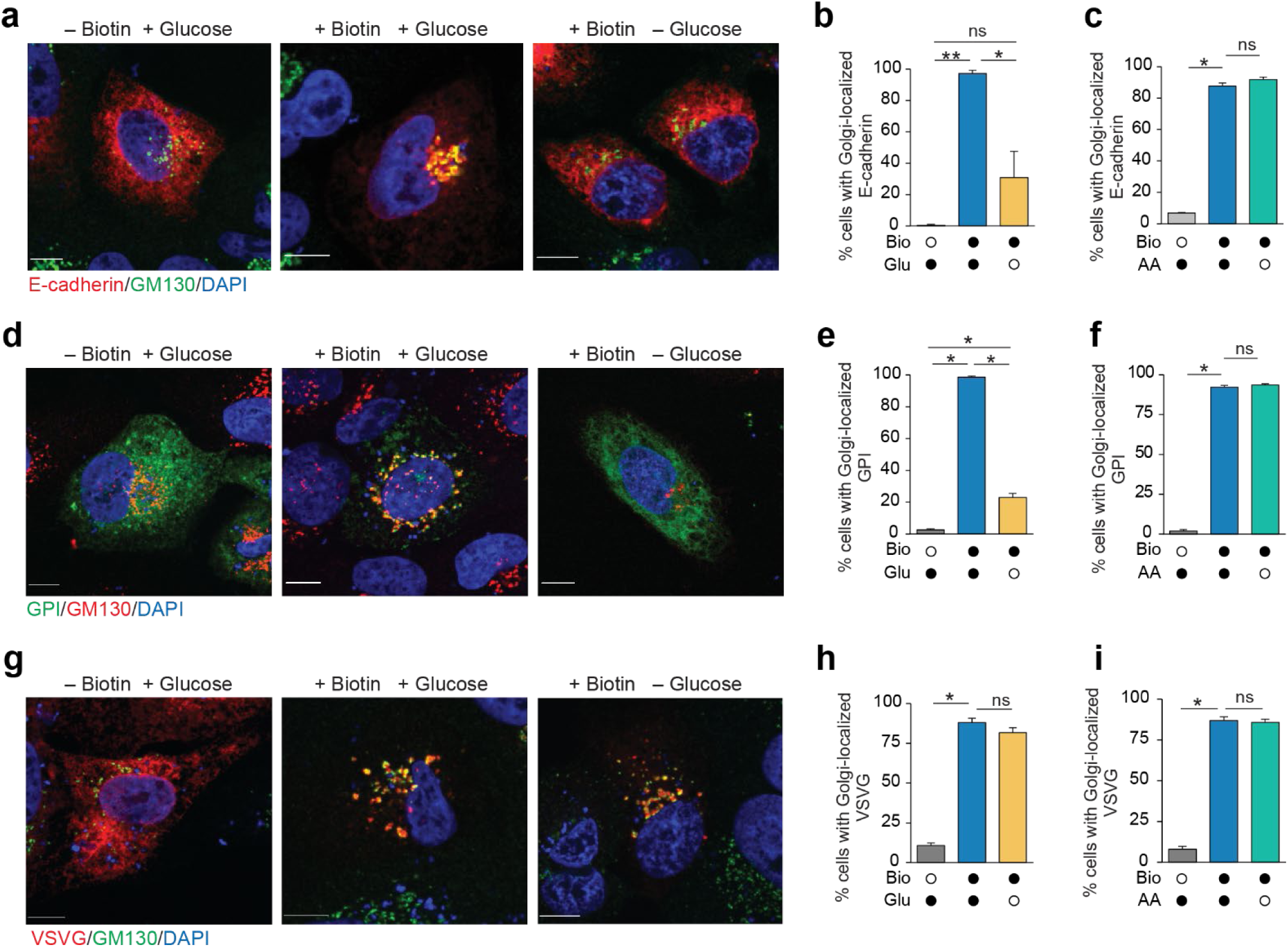
Acute nutrient deprivation differentially regulates ER-to-Golgi trafficking. (**a–i**) U2OS cells were transiently transfected with the indicated RUSH constructs (E-cadherin, GPI, or vesicular stomatitis virus G protein [VSVG]). After 48 h, cells were subjected to the following treatments: (1) incubation in Dulbecco’s Modified Eagle Medium (DMEM) with or without glucose (Glu) for 1 h, followed by biotin (Bio) treatment for 30 min to release reporters from ER hooks (**a, b, d, e, g, h**); (2) incubation in DMEM or Earle’s Balanced Salt Solution (amino acid [AA] starvation media) for 2 h, followed by biotin addition for 30 min (**c, f, i**). All conditions included fetal bovine serum (FBS) or dialyzed FBS. Representative immunofluorescence images (**a, d, g**) show reporter distribution following staining with anti-GM130 and DAPI. Quantification (mean ± S.E.M., n = 3 biological replicates, >45 cells per replicate (**b, c, e, f, h, i**) shows the percentage of cells with Golgi-localized reporters following biotin addition (closed circles) and/or nutrient starvation (open circles). Statistical significance was analyzed by one-way ANOVA followed by Holm-Sidak multiple comparisons. **P < 0.001, *P < 0.01; ns, not significant. Scale bar, 10 μm.

**Extended Data Fig. 1.**
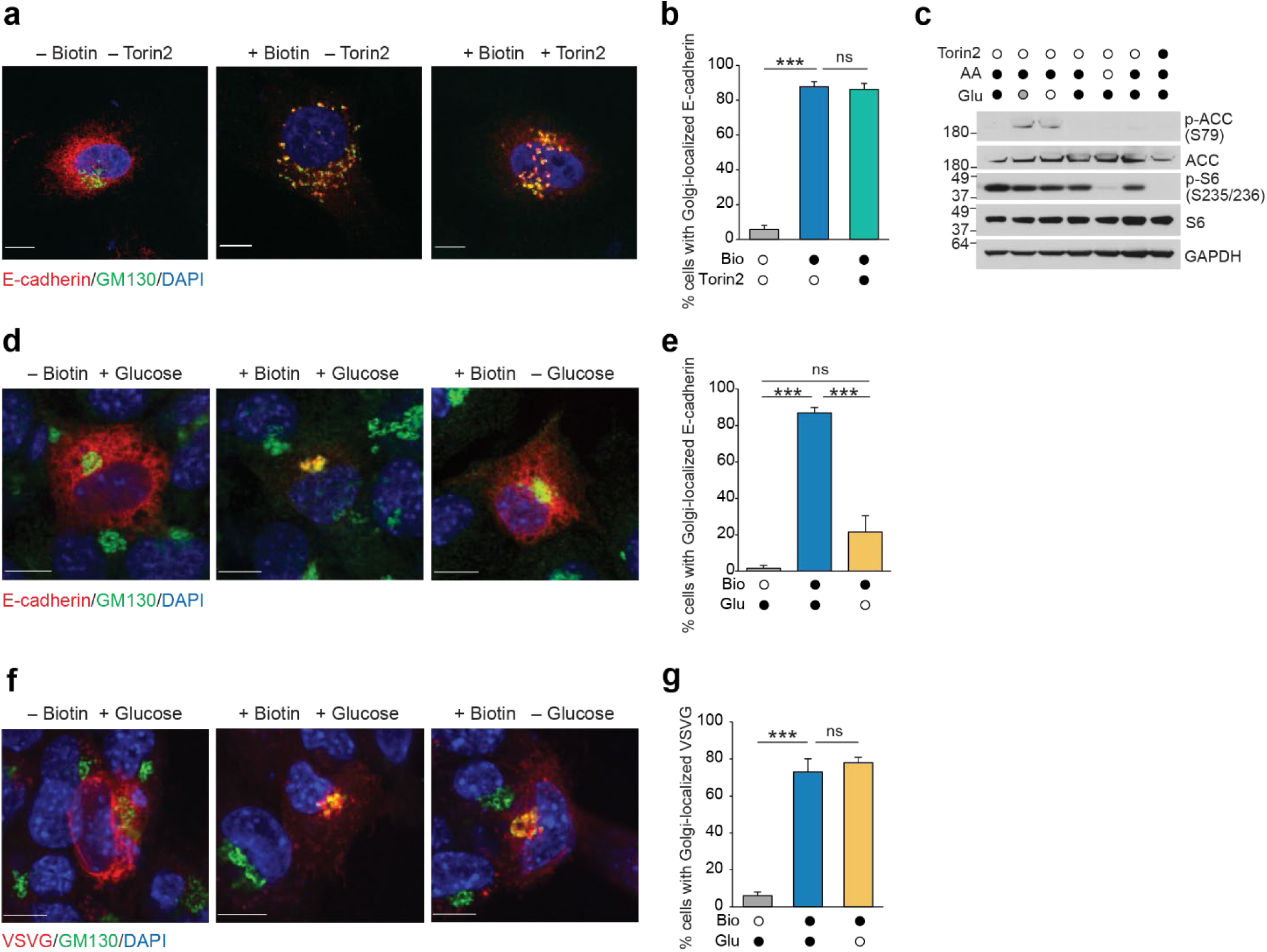
Supplementary data for Fig. 1. (**a, b**) U2OS cells transiently transfected with E-cadherin RUSH construct were treated with Torin2 (final conc. 250 nM) for 6 h, followed by biotin treatment for 30 min. (**a**) Representative immunofluorescence images following anti-GM130 and DAPI staining. (**b**) Quantification (mean ± S.E.M., n = 3 biological replicates, >63 cells per replicate of cells with Golgi-localized reporters following Torin2 treatment. (**c**) Representative immunoblots for p-ACC (S79), ACC, p-S6 (S235/236), S6, and GAPDH following glucose starvation (0.5 h, gray circle; 1 h, open circle), amino acid starvation (2 h, open circle), or Torin2 (6 h) treatment. (**d–g**) Mouse embryonic fibroblasts transiently transfected with the indicated RUSH constructs (E-cadherin or vesicular stomatitis virus G protein [VSVG]) were incubated in Dulbecco’s Modified Eagle Medium with or without glucose for 1 h, followed by biotin addition for 30 min. (**d, f)** Representative immunofluorescence images following anti-GM130 and DAPI staining. (**e, g**) Quantification (mean ± S.E.M., n = 3 biological replicates, >41 cells per replicate) of cells with Golgi-localized reporters following biotin addition and/or glucose starvation. Statistical significance was analyzed with one-way ANOVA followed by Tukey’s multiple comparisons. ***P < 0.001; ns, not significant. Scale bar, 10 μm.

This cargo-selective blockade of ER-to-Golgi transport under glucose starvation was not unique to U2OS cells; similar effects were observed in mouse embryonic fibroblasts (MEFs): E-cadherin trafficking was inhibited and VSVG trafficking was unaffected (Extended data Fig. 1d–g). Collectively, these findings demonstrate that glucose starvation, but not amino acid starvation or mTOR inhibition, selectively impaired ER-to-Golgi trafficking of a subset of cargoes, suggesting a cargo- and nutrient-specific regulatory mechanism.

### SEC24C Regulates E-cadherin Trafficking Under Glucose Starvation

The four mammalian SEC24 isoforms (A–D) confer cargo specificity^12^. SEC24A and SEC24B form one subfamily while SEC24C and SEC24D form the other, each with distinct but partially redundant cargo specificities^43, 44^. Given our observation that glucose starvation selectively impairs the trafficking of certain cargoes, we hypothesized that individual SEC24 isoforms might respond differently to nutrient stress, thereby contributing to this cargo selectivity.

To address this, we first sought to determine whether glucose starvation elicits isoform-specific changes in the subcellular organization of SEC24 isoforms. We first examined the distribution of overexpressed hemagglutinin (HA)-tagged SEC24A–D via quantitative fluorescence microscopy. Analysis revealed isoform-specific changes in puncta number and sum intensity (reflecting both area and average pixel intensity per punctum) under glucose starvation (Extended data Fig. 2a, b). The most striking changes were observed with SEC24C: glucose starvation led to a pronounced decrease in puncta number and a concomitant increase in sum intensity, indicating a redistribution and clustering of SEC24C at fewer, but more intensely labeled, sites. Starvation-induced changes in the distribution of other SEC24 isoforms were less pronounced.

Building on these findings, we next investigated the functional role of each SEC24 isoform in E-cadherin trafficking. Consistent with the established role of SEC24A in E-cadherin trafficking, and the functional overlap between SEC24A and SEC24B,^44^ RNA interference (RNAi)-mediated knockdown of SEC24A or SEC24B in U2OS cells significantly reduced E-cadherin trafficking under nutrient-replete conditions (P<0.001; Fig. 2a, b). Upon glucose starvation, E-cadherin trafficking was further diminished in these knockdown cells. In contrast, SEC24D knockdown had no effect under either nutrient-replete or glucose-starved conditions, mirroring the behavior of the control RNAi-treated cells (Fig. 2a, b). Notably, knockdown of SEC24C had no effect under nutrient-replete conditions but nullified the inhibitory effect of glucose starvation on E-cadherin trafficking (Fig. 2a, b). Immunoblot analysis was used to confirm efficient knockdown of the SEC24 isoforms (Extended data Fig. 2c). To assess the specificity of these findings, we transiently expressed an RNAi-resistant SEC24C in *SEC24C* knockdown cells, which restored the glucose starvation–induced inhibition of E-cadherin trafficking (Fig. 2c–e and Extended data Fig. 2d).

**Fig. 2:**
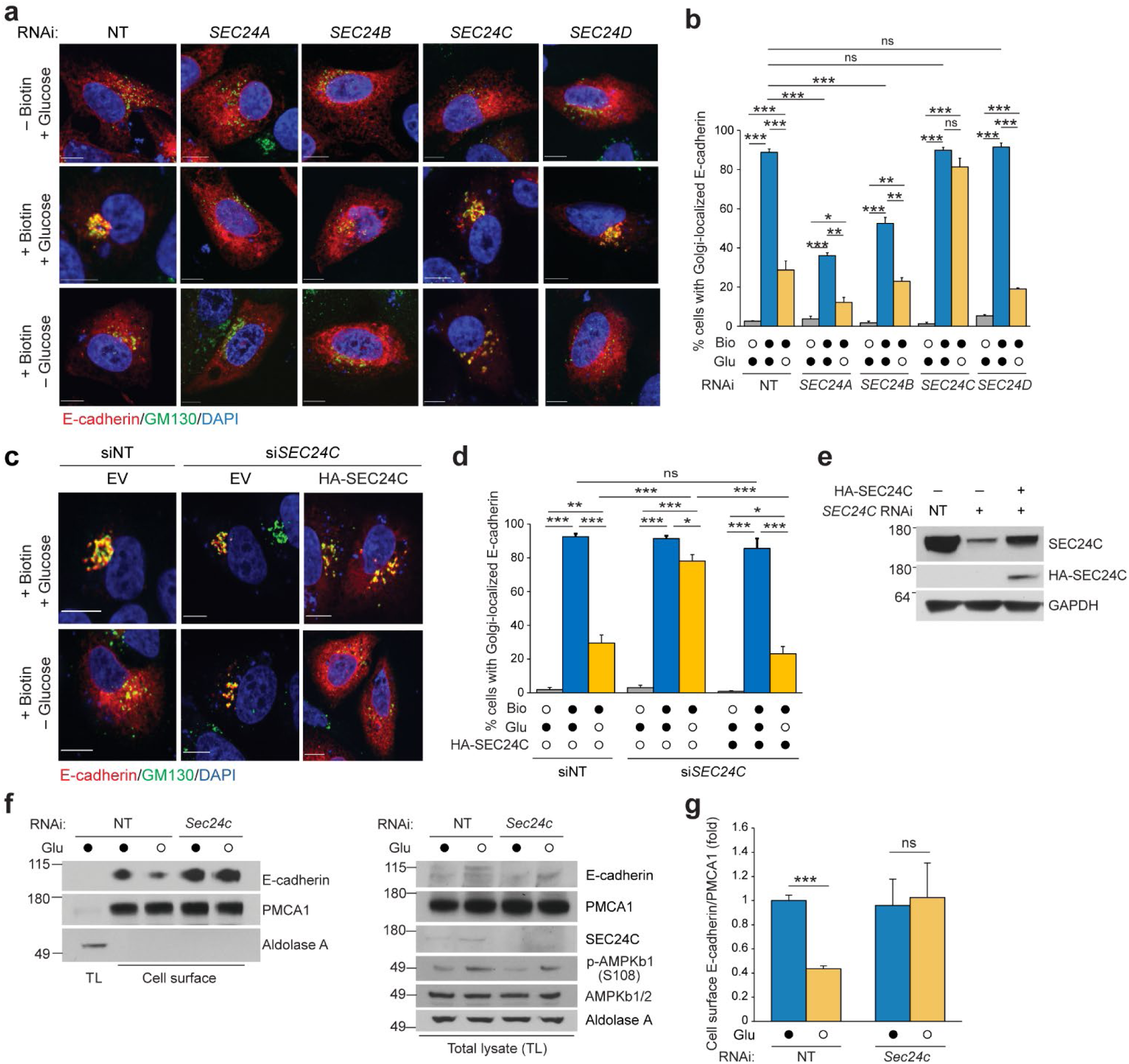
Glucose starvation impairs trafficking of E-cadherin in a SEC24C-dependent manner. (**a, b**) U2OS cells were transiently transfected with the E-cadherin RUSH construct and siRNAs targeting *SEC24A*, *SEC24B*, *SEC24C*, or *SEC24D*, as indicated. After 48 h, cells were incubated in Dulbecco’s Modified Eagle Medium (DMEM) with or without glucose for 1 h, followed by biotin treatment for 30 min. (**a**) Representative immunofluorescence images following anti-GM130 and DAPI staining show changes in reporter distribution after biotin addition and/or glucose starvation. (**b**) Percentage of cells (mean ± S.E.M., n = 3 biological replicates, >54 cells per replicate) with Golgi-localized E-cadherin following biotin treatment and/or glucose starvation. Scale bar, 10 μm. (**c–e)** U2OS cells were transiently transfected with the E-cadherin RUSH construct and either non-targeting control RNAi (siNT) or si*SEC24C*, with or without RNAi-resistant HA-tagged SEC24C. After 48 h, cells were incubated in DMEM with or without glucose for 1 h, followed by biotin addition for 30 min. (**c**) Representative immunofluorescence images following anti-GM130 and DAPI staining show changes in reporter distribution. EV, empty vector. (**d**) Percentage of cells (mean ± S.E.M., n = 3 biological replicates, >31 cells per replicate) with Golgi-localized reporters under the indicated conditions. (**e**) *SEC24C* knockdown efficiency and HA-SEC24C expression were confirmed by immunoblot analysis using the indicated antibodies. (**f, g**) Mouse embryonic fibroblasts transfected with non-targeting control RNAi (NT) or si*Sec24c* were incubated in DMEM with or without glucose for 2 h prior to treatment with membrane-impermeable NHS-biotin to biotinylate surface proteins. Biotinylated proteins were collected and analyzed by immunoblot using the indicated antibodies. (**f**) Representative immunoblots show enrichment of PCMA1 on the cell surface compared to total lysates. Cell surface levels of PCMA1 were unchanged after glucose starvation. (**g**) Levels (mean ± S.E.M., n = 3 biological replicates) of cell surface E-cadherin (relative to PCMA1) under the indicated conditions. Statistical significance was analyzed by one-way ANOVA followed by Tukey’s multiple comparisons. ***P < 0.001, **P < 0.01, *P < 0.05; ns, not significant. Scale bar, 10 μm.

**Extended Data Fig. 2:**
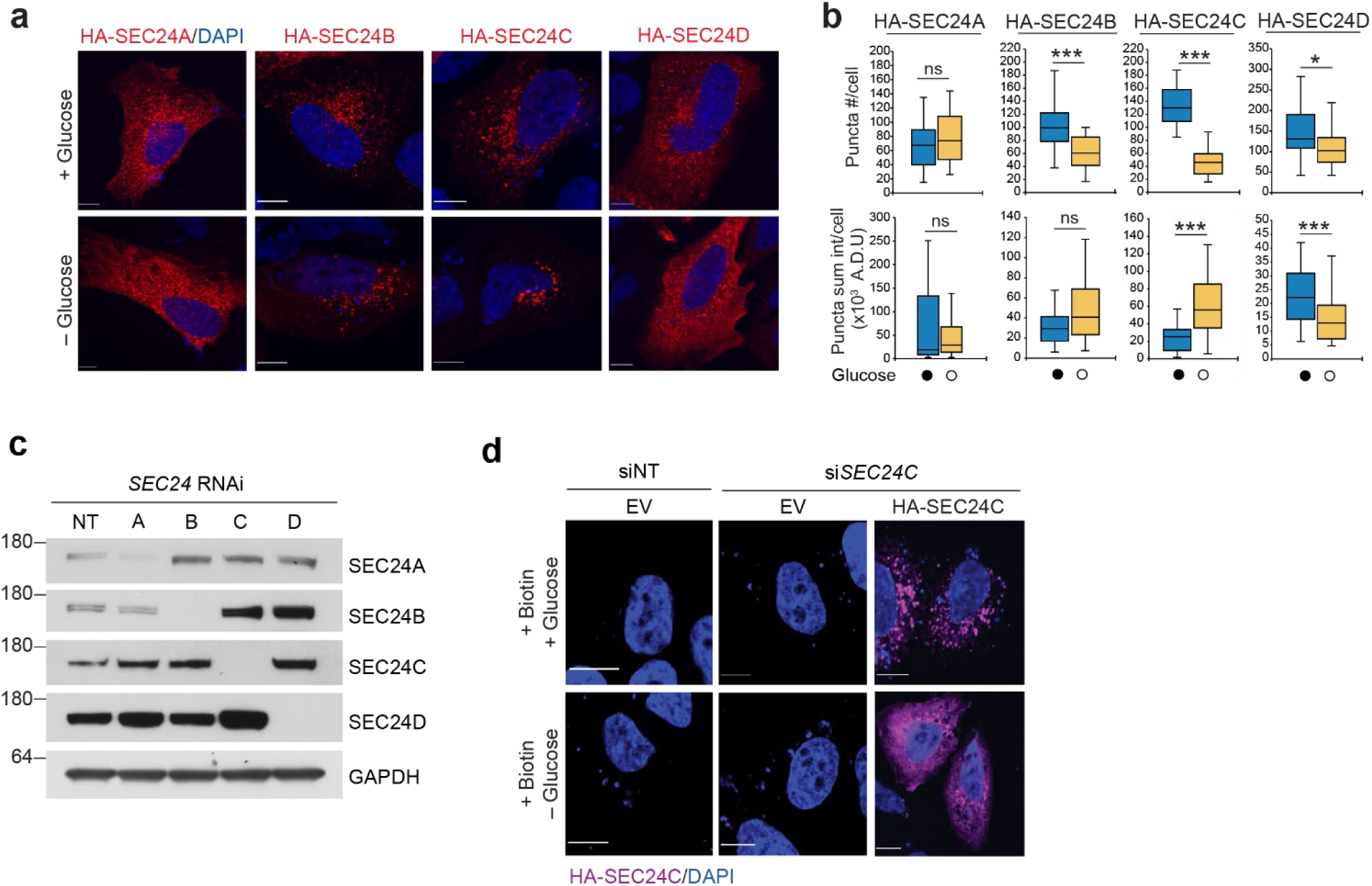
Supplementary data for Fig. 2. (**a, b**) U2OS cells were transiently transfected with HA-SEC24A, HA-SEC24B, HA-SEC24C, or HA-SEC24D expression constructs as indicated. After 48 h, cells were incubated in Dulbecco’s Modified Eagle Medium (DMEM) with or without glucose for 1 h, followed by anti-hemagglutinin (HA) antibody staining. (**a**) Representative images showing HA-SEC24 redistribution. (**b**) SEC24 puncta number and puncta intensity (mean ± S.E.M., n = 3 biological replicates, >30 cells per replicate). Statistical significance was assessed using a two-tailed paired Student’s *t*-test. ***P < 0.001, *P < 0.05. Scale bar, 10 μm. A.D.U., analog-to-digital unit. (**c**) U2OS cells were transiently transfected with the E-cadherin RUSH construct and siRNAs targeting *SEC24A*, *SEC24B*, *SEC24C*, or *SEC24D*, as indicated. After 48 h, representative immunoblot analyses demonstrated the level of knockdown using the respective SEC24 antibodies. (**d**) U2OS cells were transiently transfected with the E-cadherin RUSH construct and either non-targeting control RNAi (siNT) or si*SEC24C*, with or without RNAi-resistant HA-tagged SEC24C. After 48 h, cells were incubated in DMEM with or without glucose for 1 h, followed by biotin treatment for 30 min. Representative immunofluorescence images following anti-HA and DAPI staining show expression of HA-SEC24C in the cells. EV, empty vector. Scale bar, 10 μm.

To determine if the trafficking effects observed using the E-cadherin RUSH vector reflected changes in endogenous protein trafficking, we next assessed steady-state cell surface levels of E-cadherin by NHS-biotinylation, streptavidin-agarose pulldown, and immunoblotting in MEFs (chosen due to higher surface E-cadherin expression than found in U2OS cells). Consistent with the RUSH system results, glucose starvation led to a marked decrease in surface E-cadherin, which was prevented by *Sec24c* knockdown (Fig. 2f, g). These findings highlight the importance of SEC24C in mediating the glucose starvation–induced inhibition of E-cadherin trafficking.

### SEC24C Remodels the Cell Surface Proteome in Response to Glucose Starvation, Promoting Cellular Migration and Metastasis

To elucidate the broader impact of glucose starvation and SEC24C on the plasma membrane proteome, we performed quantitative cell surface proteomics in MEFs using NHS-biotin labeling, streptavidin-agarose pulldown, and tandem mass tag–based mass spectrometry (Fig. 3a). This analysis revealed widespread remodeling of the cell surface proteome upon glucose starvation, with both increases and decreases in surface protein abundance (Fig. 3b, Table S1a-c). Notably, a subset of proteins whose surface levels were reduced by glucose starvation, including E-cadherin, showed minimal changes under starvation conditions when *Sec24c* was knocked down (Fig. 3b, Table S1a). Further pathway enrichment analysis revealed that cell-cell adhesion molecules were enriched among these Sec24c-dependent, glucose-sensitive proteins (Fig. 3c and Table S1d), suggesting that SEC24C regulates the surface availability of certain adhesion proteins in response to metabolic stress.

**Fig. 3:**
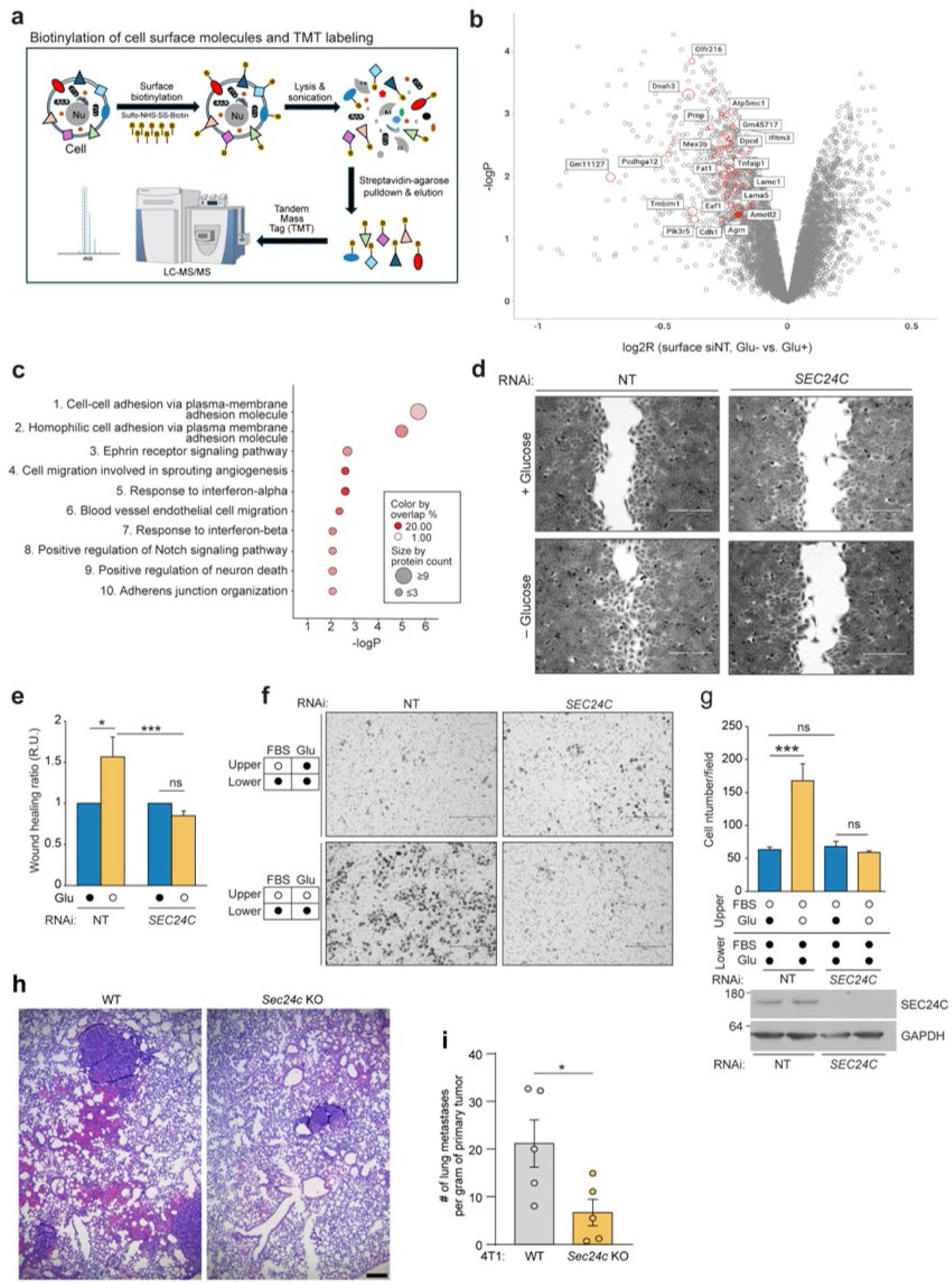
SEC24C regulates the cell surface proteome and cell migration in response to glucose starvation and promotes metastasis *in vivo*. (**a**) Schematic of cell surface biotinylation and tandem mass tag (TMT)–based proteomics analysis. MEFs transfected with non-targeting control RNAi (NT) or si*Sec24c* were incubated in Dulbecco’s Modified Eagle Medium (DMEM) with or without glucose for 2 h prior to biotinylation of surface proteins with membrane-impermeable NHS-biotin. Biotinylated proteins were collected and analyzed via TMT proteomics. (**b**) Proteins whose cell surface levels were altered by glucose starvation. Open circles outlined in red highlight the 20 proteins whose surface levels were most decreased in NT RNAi-treated MEFs (fold-change>10%, P<0.05) but not in si*Sec24c*-treated MEFs; the degree of reversal is indicated by the circle size. The solid red circle denotes E-cadherin (Cdh1). (**c**) Pathway enrichment analysis of proteins whose cell surface levels decreased upon glucose starvation in NT RNAi-treated MEFs but not in si*Sec24c*-treated MEFs. (**d, e**) U2OS cells grown in wound healing assay chambers were transfected with NT or si*SEC24C*. Cells were incubated in DMEM with or without glucose for 2 h prior to removal of the insert, then incubated in DMEM with glucose for 24 h and imaged. (**d**) Representative brightfield images showing wound closure status. (**e**) Wound closure ratio (mean ± S.E.M., n = 3 biological replicates, 5 fields per replicate) under the indicated conditions. Statistical significance was analyzed with one-way ANOVA followed by Holm-Sidak multiple comparisons. ***P < 0.001, *P < 0.05; ns, not significant. Scale bar, 300 μm. R.U., relative unit. (**f, g**) U2OS cells were transfected with NT or si*SEC24C*. After 48 h, cells were seeded into the upper chambers of a Transwell plate in media with or without FBS and/or glucose. After 18 h, migrated cells were stained with crystal violet. (**f**) Representative images of crystal violet-stained Transwell membranes illustrating the extent of cell migration, as indicated by the number of cells that have crossed to the underside of the membrane. (**g**) Top: Number of migrated U2OS cells (mean ± S.E.M., n = 3 biological replicates) under the indicated conditions. Bottom: Immunoblot analysis of *SEC24C* knockdown efficiency using anti-SEC24C and anti-GAPDH antibodies. Statistical significance was analyzed with one-way ANOVA followed by Holm-Sidak multiple comparisons. ***P < 0.001, **P < 0.01; ns, not significant. Scale bar, 300 μm. (**h, i**) Sec24c-dependent primary tumor growth and spontaneous metastasis were assessed in a mouse cancer metastasis model. Wild-type (WT) and *Sec24c* knockout 4T1 breast cancer cells were injected into the mammary fat pad of WT female mice. After 28 days, primary tumors were resected and weighed, and lungs were fixed. (**h**) Paraffin-embedded lung sections were stained with hematoxylin and eosin (H&E) and imaged. Scale bar, 200 μm. (**i**) The number of metastatic foci per gram of primary tumor was determined after counting metastatic sites in H&E-stained lung sections (mean ± S.E.M., n = 5 mice). Statistical significance was assessed using a two-tailed unpaired Student’s *t*-test. *P < 0.05; ns, not significant.

**Extended Data Fig. 3:**
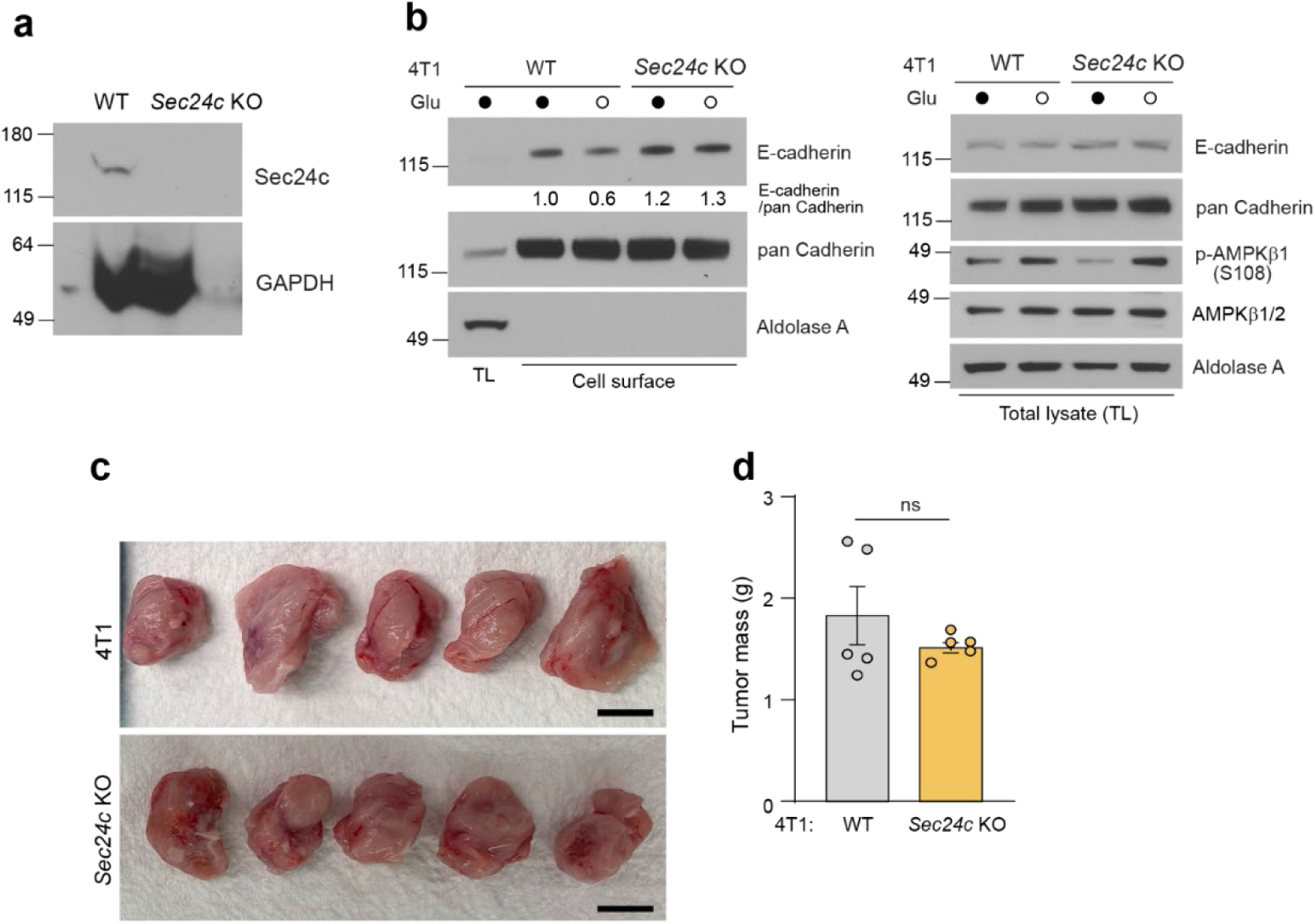
Supplementary data for Fig. 3. (**a**) CRISPR-mediated *Sec24c* knockout KO 4T1 cells were analyzed by immunoblot using anti-Sec24C and anti-GAPDH antibodies. (**b)** Wild-type (WT) and *Sec24c* KO 4T1 cells were incubated in Dulbecco’s Modified Eagle Medium with or without glucose for 4 h prior to biotinylation of surface proteins with membrane-impermeable NHS-biotin. Biotinylated proteins were collected and analyzed by immunoblot using the indicated antibodies. Representative immunoblots show enrichment of cadherins on the cell surface compared to total lysates; cell surface levels of pan Cadherin were unchanged with glucose starvation. (**c, d)** Sec24c-dependent primary tumor growth and spontaneous metastasis were assessed in a mouse cancer metastasis model. WT and *Sec24c* KO 4T1 cells were injected into the mammary fat pad of WT female mice. After 28 days, primary tumors for each condition were resected, weighed, and imaged (n = 5 mice per group). (**c**) Representative image of primary tumors resected from the mammary fat pads of female recipient mice. Scale bar, 1 cm. (**d**) Mass of primary tumors (in grams; mean ± S.E.M., n = 5 mice) derived from WT and *Sec24c* KO 4T1 cells. Statistical significance was assessed using a two-tailed unpaired Student’s *t*-test. ns, not significant.

**Table S1.** Altered Levels of Cell Surface Proteins Following Glucose Starvation and Associated Pathway Enrichment Analyses in Murine Embryonic Fibroblasts. This table presents lists of cell surface proteins with significantly altered levels in murine embryonic fibroblasts (MEFs) following glucose starvation, together with results from pathway enrichment analyses. Protein quantification was performed using TMT-labeled samples subjected to NHS-biotinylation and streptavidin pull-down to enrich for cell surface proteins, comparing wild-type (WT), non-targeting siRNA-treated control (CT), and siSec24c-treated (KD) MEFs. (**a**) Quantitative and statistical analysis of cell surface protein levels across conditions. (**b**) Pathway enrichment analysis (Enrichr) of proteins significantly down-regulated by glucose starvation in WT and CT MEFs. (**c**) Pathway enrichment analysis (Enrichr) of proteins significantly up-regulated under the same conditions. (**d**) Pathway enrichment analysis (DAVID) of proteins whose down-regulation in response to glucose starvation is dependent on Sec24c. See accompanying Excel file for complete data.

Given the critical role of cellular adhesion in cell migration and metastasis^45, 46^, we investigated how glucose starvation and SEC24C influence cell motility. In wound-healing assays, glucose-starved U2OS cells exhibited accelerated wound closure compared to controls, an effect abolished by *SEC24C* knockdown (Fig. 3d, e). In Transwell migration assays, knocking down *SEC24C* did not affect migration toward a fetal bovine serum (FBS) gradient under nutrient-replete conditions (Fig. 3f, g). However, culturing cells in media without FBS and glucose significantly enhanced migration toward the lower chamber containing both FBS and glucose; this accelerated migration was also blocked upon *SEC24C* knockdown (P<0.001; Fig. 3f, g).

To assess the *in vivo* relevance of these findings, we used the 4T1 murine mammary carcinoma model, which forms primary tumors and spontaneously metastasizes to the lung following mammary fat pad injection^47^. *Sec24c* knockout (KO) 4T1 cells were generated via CRISPR-mediated gene editing, and loss of Sec24c was confirmed by immunoblot analysis (Extended data Fig. 3a). Glucose starvation resulted in a modest decrease in surface E-cadherin levels in wild-type (WT) but not in *Sec24c* KO 4T1 cells *in vitro* (Extended data Fig. 3b), consistent with our observations in MEFs (Fig. 2f, g). *In vivo, Sec24c* KO did not alter primary tumor size (Extended data Fig. 3c, d), but resulted in significantly fewer lung metastases than those found in WT controls (P<0.05; Fig. 3h, i). Collectively, these results identify SEC24C as a key mediator of glucose starvation– induced remodeling of the cell surface proteome with functional consequences for cell migration and metastatic potential in a murine cancer model.

### SEC24C-Dependent, Starvation-induced Inhibition of ER-to-Golgi Trafficking Requires AMPK and ULK1, but Not Autophagy

Given the importance of SEC24C in mediating glucose starvation–induced remodeling of the cell surface proteome and its effect on cell migration and metastasis, we next sought to elucidate the mechanism underlying the inhibition of ER-to-Golgi trafficking observed under glucose starvation. SEC24C has been previously implicated in the selective autophagy of ERES (ERES-phagy), particularly in response to mTOR inhibition, when it mediates the relocalization of ERES components to lysosomes for protein quality control^48^. Given this established role and the fact that glucose starvation is a potent activator of autophagy through AMPK and ULK1/2 signaling^49, 50^, we hypothesized that the decrease in E-cadherin trafficking observed during glucose starvation might involve an autophagy-dependent mechanism targeting ERES.

To test this, we first examined the requirement for ULK1 and ULK2, related serine-threonine kinases that initiate autophagy and are directly activated by AMPK^49–52^. Using CRISPR-mediated gene editing, we generated *ULK1/2* double KO (DKO) U2OS cells^53^. These cells were resistant to the trafficking inhibition imposed by glucose starvation, indicating that ULK1/2 are essential for this process (Fig. 4a–d).

**Fig. 4:**
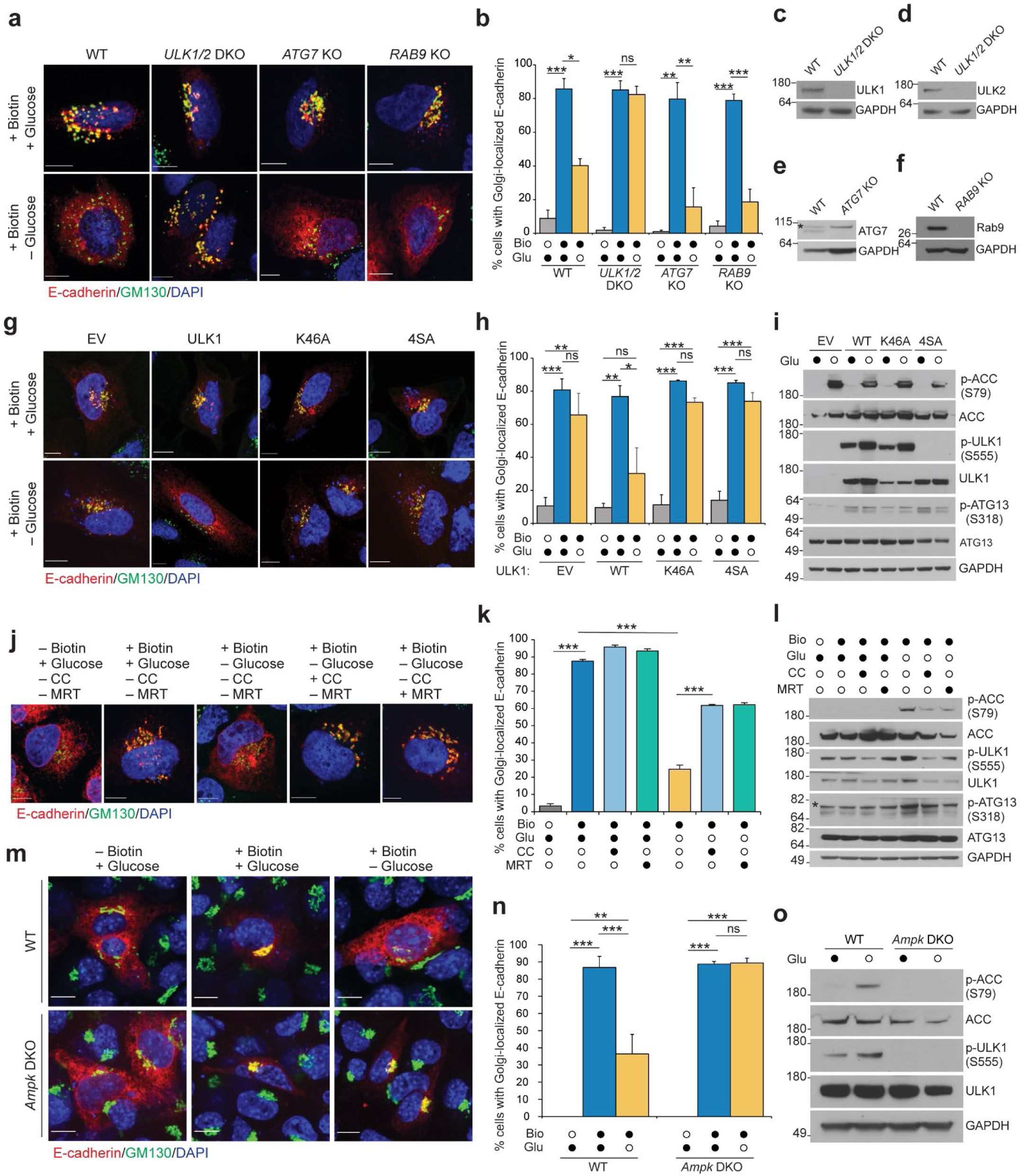
AMPK and ULK1/2, but not other autophagy-related proteins, are required for starvation-induced inhibition of ER-to-Golgi trafficking. **(a–f)** U2OS cells (wild-type [WT], *ULK1/2* double knockout [DKO], *ATG7* knockout [KO], or *RAB9* KO were transfected with the E-cadherin RUSH construct. After 48 h, cells were incubated in Dulbecco’s Modified Eagle Medium (DMEM) with or without glucose for 1 h, followed by biotin treatment for 30 min. (**a**) Representative immunofluorescence images captured following anti-GM130 and DAPI staining show changes in reporter distribution after biotin addition and/or glucose starvation. (**b**) Percentage of cells (mean ± S.E.M., n = 3 biological replicates, >37 cells per replicate) with Golgi-localized E-cadherin. (**c–f**) Representative immunoblot analyses show protein levels of ULK1, ULK2, ATG7, or RAB9 in WT and indicated KO cell lines; * indicates nonspecific band in (**e**). (**g–i**) Rescue of glucose starvation–induced ER-to-Golgi trafficking inhibition was evaluated in U2OS *ULK1/2* DKO cells stably expressing empty vector (EV), WT ULK1, kinase-dead ULK1 (K46A), or AMPK-resistant ULK1 (4SA). Cells were transfected with the E-cadherin RUSH construct, incubated in DMEM with or without glucose for 1 h, and treated with biotin for 30 min. (**g**) Representative immunofluorescence images captured following anti-GM130 and DAPI staining show changes in reporter distribution after biotin addition and/or glucose starvation. (**h**) Golgi-localized E-cadherin in U2OS *ULK1/2* DKO cells expressing WT or mutant forms of ULK (mean ± S.E.M., n = 3 biological replicates, >30 cells per replicate). (**i**) Representative immunoblot analysis shows AMPK activity as indicated by total and phospho-ACC, and ULK1 expression, and AMPK-mediated phosphorylation of ULK1 at S555 in the indicated cell lines. (**j–l**) WT U2OS cells were transfected with the E-cadherin RUSH construct. After 48 h, cells were pre-treated with compound C (5 μM, 4 h) or MRT68921 (1 μM, 24 h) and incubated in DMEM with or without glucose for 1 h, followed by biotin treatment for 30 min. (**j**) Representative immunofluorescence images following anti-GM130 and DAPI staining show changes in reporter distribution after biotin addition and/or glucose starvation. (**k**) Golgi-localized E-cadherin in U2OS cells before and after glucose starvation (mean ± S.E.M., n = 3 biological replicates, >103 cells per replicate). (**l**) Representative immunoblot analysis demonstrates activity of each inhibitor post biotin treatment. (**m–o**) WT or *Prkaa1/2* (AMPKα1/2) DKO murine embryonic fibroblasts (MEFs) were transfected with the E-cadherin RUSH construct. After 48 h, cells were incubated in DMEM with or without glucose for 1 h, followed by biotin treatment for 30 min. (**m**) Representative immunofluorescence images captured following anti-GM130 and DAPI staining show changes in reporter distribution after biotin addition and/or glucose starvation. (**n**) Golgi-localized E-cadherin in in WT and AMPK α1/2 DKO MEFs before and after glucose starvation (mean ± S.E.M., n = 3 biological replicates, >33 cells per replicate). (**o**) Levels of phosphorylated ACC and ULK1 in WT and AMPK α1/2 DKO MEFs after 1-h glucose starvation. Statistical significance was analyzed with one-way ANOVA followed by Tukey’s multiple comparisons. ***P < 0.001, **P < 0.01, *P < 0.05; ns, not significant. Scale bar, 10 μm.

**Extended Data Fig. 4:**
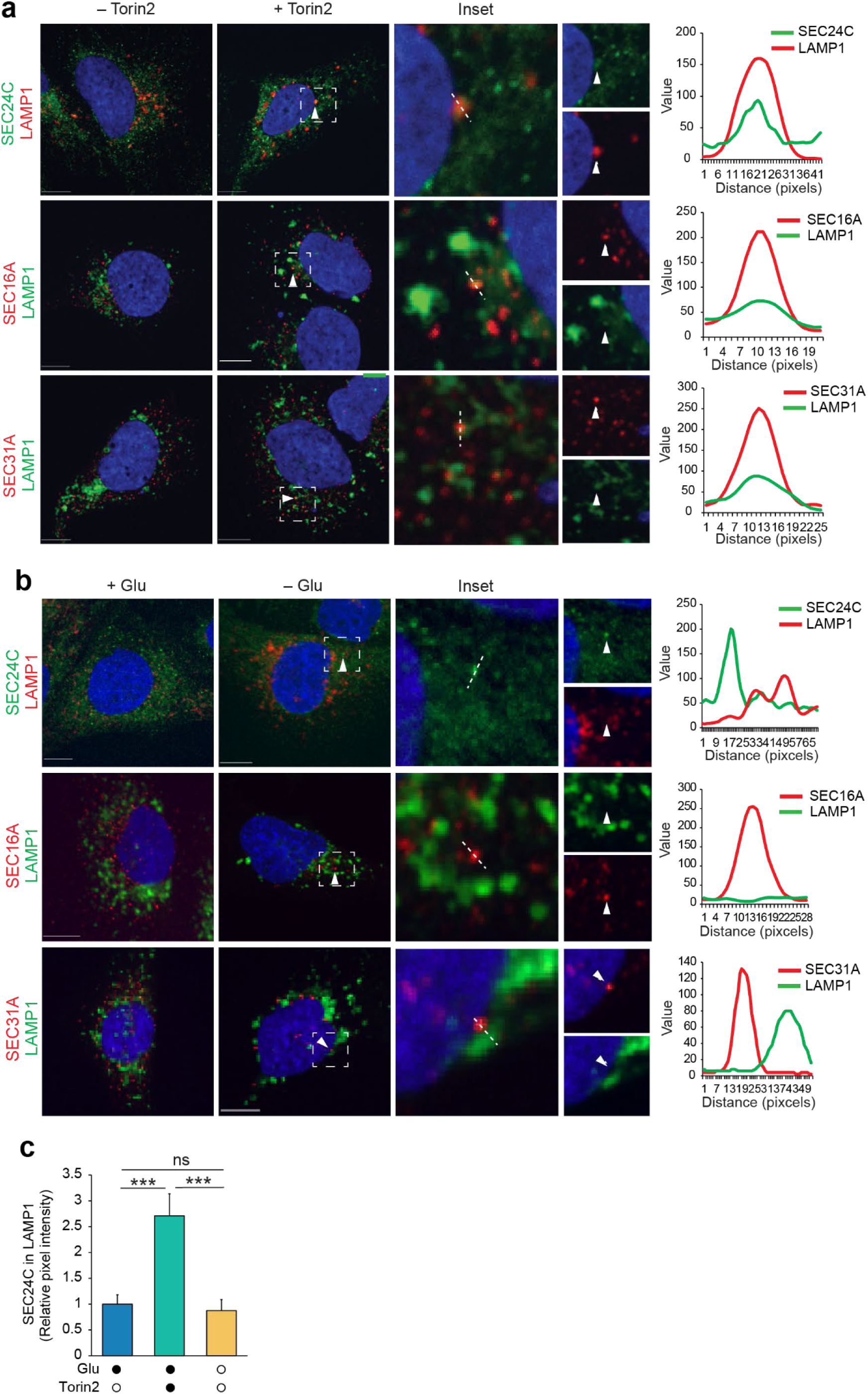
Supplemental data for Fig. 4. **(a–c)** U2OS FLAG-SEC24C, FLAG-SEC16A, and FLAG-SEC31A knock-in cells were treated with Torin2 (250 nM) (A) or starved of glucose for 6 h (B). Representative immunofluorescence images taken after staining cells using an anti-FLAG antibody and LAMP1 as indicated. Line scans indicate the degree of colocalization between LAMP1 and COPII component and correlate to the lines drawn in the magnified images (inset) and color separation. Intensity profiles (value) were obtained by using Fiji software. White solid arrow indicates targeted foci to line scans. (C) SEC24C signal in LAMP1 structures was quantified after segmentation of each color (mean ± S.E.M., n=31 cells). Statistical significance was analyzed using one-way ANOVA followed by Tukey’s multiple comparisons. ***P < 0.001; ns, not significant. Scale bar represents 10 µM.

As ULK1/2 have been implicated in both canonical (ATG7-dependent) and non-canonical (RAB9-dependent) forms of autophagy^54^, we next tested whether either of these pathways were involved. To examine canonical autophagy, we generated U2OS cells lacking ATG7, an E1-like enzyme required for autophagosome formation^55^. To examine non-canonical autophagy, we generated U2OS cells lacking RAB9, a small GTPase that mediates an alternative, ULK1/2-dependent, ATG7-independent autophagy pathway involving autophagosome formation from the trans-Golgi and late endosomes^56^. Notably, both *ATG7* and *RAB9* KO cells retained the glucose starvation–induced inhibition of E-cadherin trafficking (Fig. 4a, b, e, f), indicating that neither canonical ATG7-dependent autophagy nor the ATG7-independent, RAB9-dependent pathway are required for this trafficking block.

To confirm that the starvation-induced trafficking block was not due to ERES-phagy, we examined the subcellular localization of COPII components (i.e., SEC24C, SEC16A, and SEC31A) and the lysosomal marker LAMP1 (Extended data Fig. 4a, b). Treatment with Torin2 was used as a positive control. Consistent with previous reports of ERES-phagy following mTOR inhibition^48, 57^, treatment with Torin2 induced the co-localization of SEC24C and LAMP1 (Extended data Fig. 4a, c). In contrast, glucose starvation did not increase SEC24C-LAMP1 co-localization, supporting our genetic data that trafficking inhibition occurs independently of ERES-phagy or lysosomal targeting of ERES (Extended data Fig. 4b, c).

To further investigate the role(s) of ULK1 kinase activity and its regulation by AMPK, we performed rescue experiments in *ULK1/2* DKO cells. Re-expression of WT ULK1 restored the glucose starvation–induced inhibition of E-cadherin trafficking, whereas neither a kinase-dead (KD) ULK1 mutant (K46A) nor the ULK1-4SA mutant^49^, which harbors alanine substitutions at four AMPK phosphorylation sites (S467A, S555A, T574A, S637A), could rescue this phenotype (Fig. 4g–i). This demonstrates that both ULK1 kinase activity and its phosphorylation by AMPK are essential for the starvation-induced trafficking block.

Consistent with these genetic data, pharmacological inhibition of AMPK by Compound C^58^ or inhibition of ULK1/2 by MRT68921^59^ reduced the glucose starvation–induced inhibition of ER-to-Golgi trafficking (Fig. 4j–l). Moreover, glucose starvation failed to inhibit E-cadherin trafficking in MEFs lacking both AMPKα1 and AMPKα2 isoforms^60^, further confirming this inhibitory effect depends on AMPK signaling (Fig. 4m–o).

Collectively, these results demonstrate that the inhibition of ER-to-Golgi trafficking of E-cadherin in response to glucose starvation does not depend on either canonical (ATG7-dependent) or non-canonical (RAB9-dependent) autophagy. Instead, this process requires AMPK activation, ULK1 phosphorylation, and ULK1 kinase activity.

### SEC24C Drives Reorganization of COPII Components and Alters ERES Dynamics in Response to Glucose Starvation

Having established that AMPK and ULK1 are both essential for the inhibition of ER-to-Golgi trafficking during glucose starvation, we sought to define the downstream cellular events through which SEC24C mediates this process. Building on our preliminary findings that glucose starvation induces a unique redistribution of overexpressed HA-tagged SEC24C into fewer, more intensely labeled puncta (Extended data Fig. 2a, b), we evaluated the subcellular distribution and interactions of endogenous SEC24C and other COPII components to determine whether these changes were associated with a broader reorganization of COPII complexes.

First, using antibodies against endogenous SEC24C and the outer coat component SEC13, we found that glucose starvation led to a marked decrease in the number of SEC24C and SEC13 puncta per cell, accompanied by an increase in summed fluorescence intensity (reflecting both the area of the puncta and the average pixel intensity) for each component (Fig. 5a–e). Co-localization analysis revealed a significant increase in overlap between SEC24C and SEC13 staining (P<0.001; Fig. 5f). Together, these results suggest increased clustering and stabilization of COPII complexes under nutrient stress.

**Figure 5.**
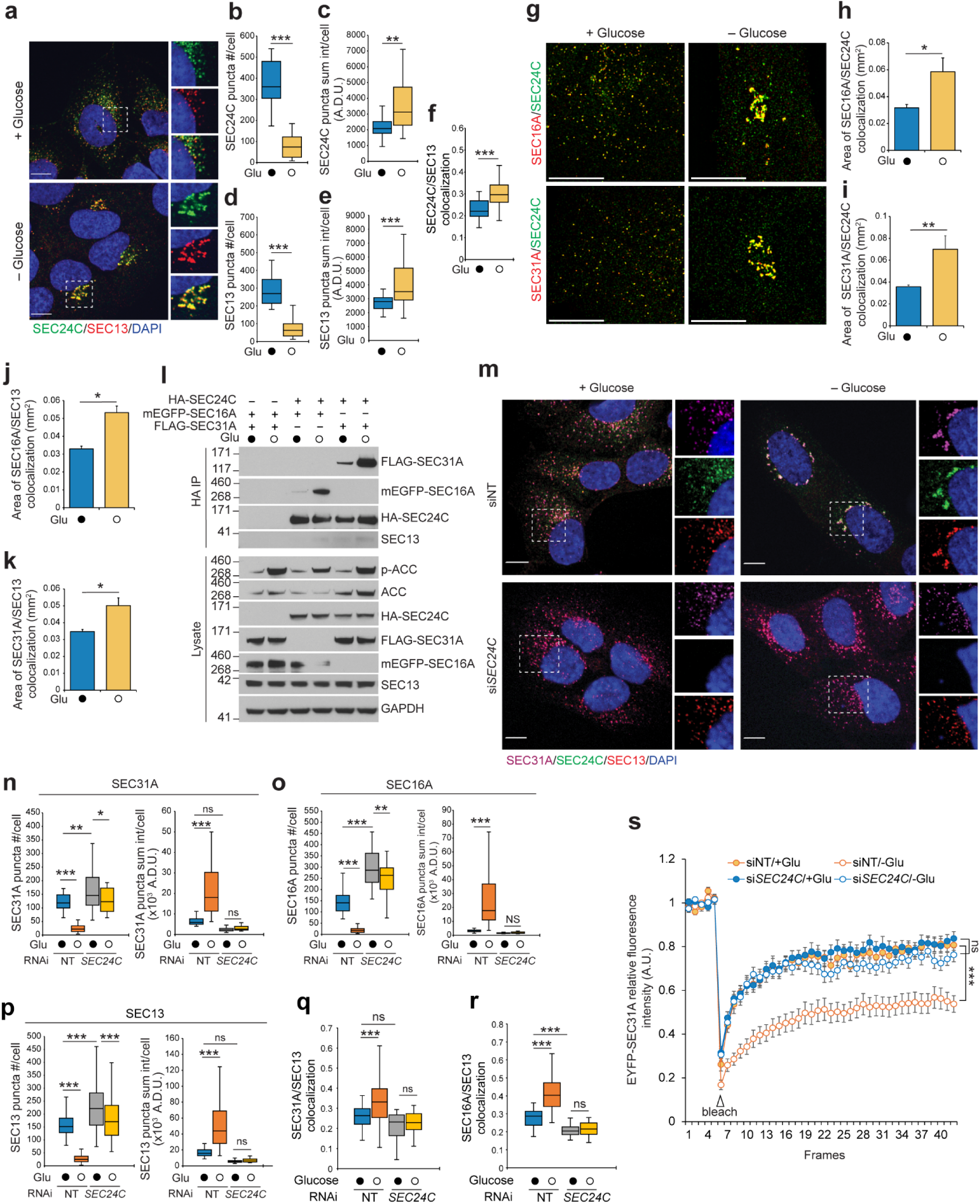
SEC24C drives reorganization of COPII components in response to glucose starvation. (**a–f**) U2OS cells were incubated in Dulbecco’s Modified Eagle Medium (DMEM) with or without glucose for 1 h, followed by antibody staining for SEC24C and SEC13. (**a**) Representative immunofluorescence images of SEC24C and SEC13 staining show changes in the distribution of COPII components after glucose depletion. Dotted squares indicate magnified areas; individual colors represent different components. (**b, d**) Number of SEC24C and SEC13 puncta; (**c, e**) puncta sum intensity; and (**f**) degree of colocalization between SEC24C and SEC13, as assessed by the Pearson’s correlation coefficient. Data are shown as mean ± S.E.M., n > 30 cells per replicate. (**g–k**) U2OS FLAG-SEC16A knock-in or FLAG-SEC31A knock-in cells were incubated in DMEM with or without glucose for 1 h and stained with anti-FLAG and SEC24C antibodies. (**g**) Super-resolution confocal images show redistribution of SEC24C, SEC31A, and SEC16A. Quantification of colocalization area: SEC16A/SEC24C (**h**), SEC31A/SEC24C (**i**), SEC16A/SEC13 (**j**), and SEC31A/SEC13 (**k**). (**l**) U2OS cells were transfected with expression plasmids encoding HA-SEC24C, mEGFP-SEC16A, and FLAG-SEC31A as indicated. After 48 h, cells were incubated in DMEM with or without glucose for 1 h, followed by HA-SEC24C immunoprecipitation. Representative immunoblot analyses demonstrate molecular interaction between HA-SEC24C and mEGFP-SEC16A or FLAG-SEC31A. (**m–r**) U2OS FLAG-SEC16A and FLAG-SEC31A knock-in cells were transfected with either non-targeting siRNA (siNT) or si*SEC24C*. After 48 h, cells were incubated in DMEM with or without glucose for 1 h, followed by antibody staining for FLAG, SEC24C, and SEC13. (**m**) Representative immunofluorescence images show changes in the distribution of COPII components after glucose starvation in U2OS FLAG-SEC31A knock-in cells (see Extended data Fig. 5b for representative images from FLAG-SEC16A knock-in cells). Dotted squares indicate magnified areas; individual colors represent different components. Quantification of SEC31A (**n**), SEC16A (**o**), and SEC13 (**p**) puncta number and intensity data are shown as mean ± S.E.M., n > 30 cells per replicate. Pearson’s correlation coefficients between the indicated protein pairs reflect the degree of colocalization: SEC31A/SEC13 (**q**), and SEC16A/SEC13 (**r**). A.D.U., analog-to-digital unit. (**s**) U2OS cells were transfected with EYFP-SEC31A and either non-targeting siRNA (siNT) or si*SEC24C*. After 48 h, cells were incubated in DMEM with or without glucose for 1 h, and EYFP-SEC31A mobility was measured by fluorescence recovery after photobleaching (FRAP) assay (mean ± S.E.M., n > 11 puncta per condition). Images were acquired every 5 seconds (5 sec per frame). Mobile fractions: siNT/+Glu, 75.6%; siNT/−Glu, 45.7%; si*SEC24C*/+Glu, 75.7%; si*SEC24C*/−Glu, 68.4%. Statistical significance was analyzed using one-way ANOVA followed by Holm-Sidak multiple comparisons. ***P < 0.001, **P < 0.01, *P < 0.05; ns, not significant. Scale bar, 10 μm. A.U., arbitrary unit.

To further characterize these changes and visualize the organization of additional COPII components at endogenous expression levels, we used CRISPR/Cas9 genome editing to generate U2OS cell lines in which endogenous SEC31A and SEC16A were tagged with FLAG. Super-resolution microscopy of these tagged lines revealed that glucose starvation led to larger areas of co-localization of SEC24C or SEC13 and the endogenously tagged SEC16A or SEC31A (Fig. 5g–k). To determine whether these changes reflected altered physical interactions among COPII components, we performed co-immunoprecipitation experiments using overexpressed, epitope-tagged proteins. Immunoprecipitation of HA-tagged SEC24C revealed increased association with modified EGFP (mEGFP)-tagged SEC16A and FLAG-tagged SEC31A in glucose-starved cells (Fig. 5l). Together, these findings suggest that glucose starvation induces a reorganization of the COPII machinery at ERES that involves the stabilization of certain COPII complexes.

These observations raised the possibility that SEC24C actively drives the reorganization of COPII components during glucose starvation. To test this, we performed RNAi-mediated knockdown of *SEC24C* in the endogenous FLAG-tagged SEC16A and SEC31A lines (Extended data Fig. 5a). Depletion of SEC24C abolished the starvation-induced decrease in puncta number and increase in summed fluorescence intensity for SEC13, SEC16A, and SEC31A, as well as the increase in co-localization among these components (Fig. 5m–r and Extended data Fig. 5b–f).

Given prior evidence that SEC31A plays a key role in COPII vesicle dynamics and is regulated by cellular conditions such as combined amino acid and serum starvation^33^, we next assessed how glucose starvation and *SEC24C* knockdown influence SEC31A behavior. Using fluorescence recovery after photobleaching, we found that the mobile fraction of SEC31A was significantly reduced in non-targeting siRNA (siNT)-treated cells following glucose starvation (P<0.001; Fig. 5s), suggesting that SEC31A becomes more stably associated with COPII structures upon glucose starvation. This starvation-induced reduction in SEC31A mobility was not observed in *SEC24C*-knockdown cells (Fig. 5s), indicating that SEC24C is necessary to stabilize COPII complexes during glucose starvation.

Collectively, these results demonstrate that SEC24C orchestrates a reorganization of COPII components in response to nutrient availability, leading to altered ERES dynamics and inhibition of ER-to-Golgi trafficking during glucose starvation.

### ULK1 Is Recruited to ERES and Regulates SEC24C-Dependent COPII Reorganization During Glucose Starvation

To understand how upstream signaling pathways regulate this SEC24C-dependent process, we investigated ULK1’s coordinating role in the recruitment and organization of COPII components during glucose starvation. Previously, we established that ULK1/2 interact with and phosphorylate SEC16A to promote ER-to-Golgi trafficking of specific cargoes under nutrient-replete conditions^61^. To build on this, we first examined the interaction between ULK1 and SEC16A during glucose starvation and found that this interaction was further stabilized under nutrient stress (Fig. 6a, b).

**Figure 6.**
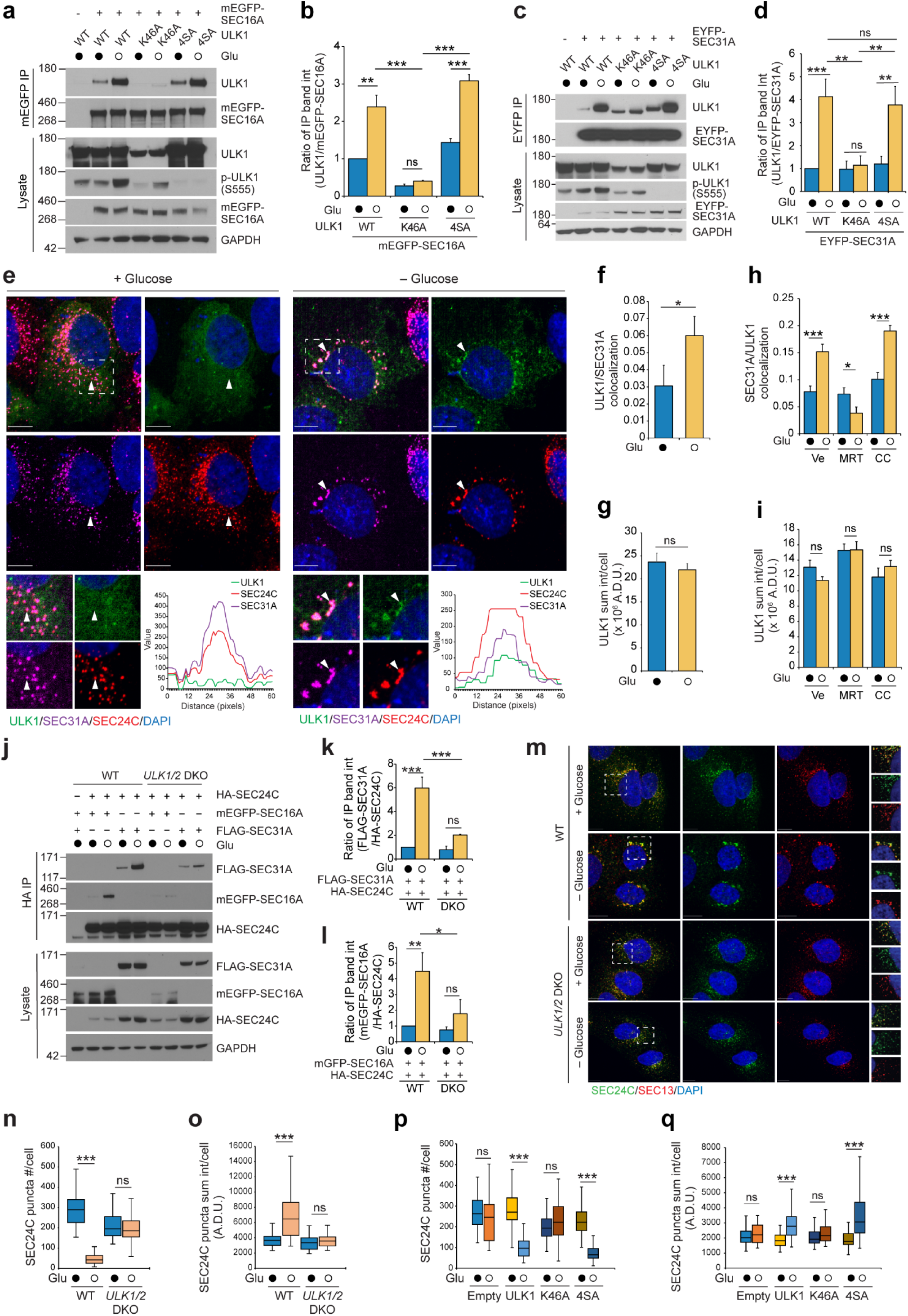
ULK1 kinase activity is required for the starvation-induced redistribution of the COPII machinery. **(a–d)** HEK293T cells were transfected with expression plasmids of mEGFP-SEC16A (**a, b**) or EYFP-SEC31A (**c, d**) with different versions of ULK1: wild-type (WT), K46A (kinase defective), 4SA (AMPK phosphorylation resistant) as indicated. After 48 h, cells were incubated in Dulbecco’s Modified Eagle Medium (DMEM) with or without glucose for 1 h followed by mEGFP or Enhanced Yellow Fluorescent Protein (EYFP) immunoprecipitation. Representative immunoblot analyses demonstrate the level of molecular interaction between mEGFP-SEC16A or EYFP-SEC3A and different ULK1 versions. (**b, d**) The ratio of co-immunoprecipitated band intensity was calculated from three independent experiments (mean ± S.E.M). **(e–g)** U2OS FLAG-SEC31A knock-in cells were transfected with the expression plasmids of EGFP-ULK1. After 48 h, cells were incubated in DMEM with or without glucose for 1 h and stained with FLAG and SEC24C antibody. (**e**) Representative immunofluorescence images taken after staining cells show changes in the distribution of EGFP-ULK1 and COPII components after glucose starvation. The degree of co-localization between ULK1 and SEC31, as determined by Pearson’s correlation coefficient **(f)**, and the total EGFP-ULK1 intensity **(g)** are shown as bar graphs (mean ± S.E.M., n > 30). Dotted squares indicate magnified areas; individual colors represent different components. Solid arrowheads highlight regions covered by line scans (bottom panels), which show the relative fluorescence intensities of ULK1 (green), SEC24C (red), and SEC31A (purple). **(h, i)** U2OS FLAG-SEC31A knock-in cells were transfected with EGFP-ULK1 expression plasmids. After 48 h, cells were pretreated with MRT 68921 (1 µM final conc.) or compound C (5 µM final conc.) for 4 h, then incubated in DMEM with or without glucose for 1 h and stained with antibodies against FLAG and SEC24C. The degree of co-localization between ULK1 and SEC31, as determined by Pearson’s correlation coefficient **(h)**, and the total GFP-ULK1 intensity **(h)** are shown as bar graphs (mean ± S.E.M., n > 30 cells). **(j–l)** U2OS WT and *ULK1/2* DKO cells were transfected with HA-SEC24C, mEGFP-SEC16A, and FLAG-SEC31A expression plasmids. After 48 h, cells were incubated in DMEM with or without glucose for 1 h followed by HA immunoprecipitation. (**j**) Representative immunoblot analyses demonstrate the level of molecular interaction between HA-SEC24C and mEGFP-SEC16A or FLAG-SEC31A. The ratio of co-immunoprecipitated FLAG-SEC31A (**k**) and mEGFP-SEC16A (**l**) band intensities, relative to SEC24C, were calculated from 3 independent experiments (mean ± S.E.M). **(m–o)** U2OS WT and *ULK1/2* DKO cells were incubated in DMEM with or without glucose for 1 h followed by antibody staining with SEC24C and SEC13. **(m)** Representative immunofluorescence images taken after staining cells show changes in the distribution of the COPII component after glucose starvation. Dotted squares indicate the areas that are magnified and shown in the rightmost panels. Differences in SEC24C and SEC13 puncta number (**n**) and intensity (**o**) between WT and *ULK1/2* DKO cells (mean ± S.E.M., n>30 cells per group). **(p, q)** *ULK1/2* DKO U2OS cells stably expressing UK1, K46A (kinase defective), or 4SA (AMPK phosphorylation-resistant) were incubated in DMEM with or without glucose for 1 h followed by staining with antibodies against SEC24C. SEC24C puncta number (**p**) and intensity (**q**) were measured and compared across WT and mutants (mean ± S.E.M., n>30 cells per group). Statistical significance was analyzed using one-way ANOVA followed by Holm-Sidak multiple comparisons. ***P < 0.001, **P < 0.01, *P < 0.05; ns, not significant. Scale bar, 10 μm. A.D.U., analog-to-digital unit.

**Extended Data Fig. 5:**
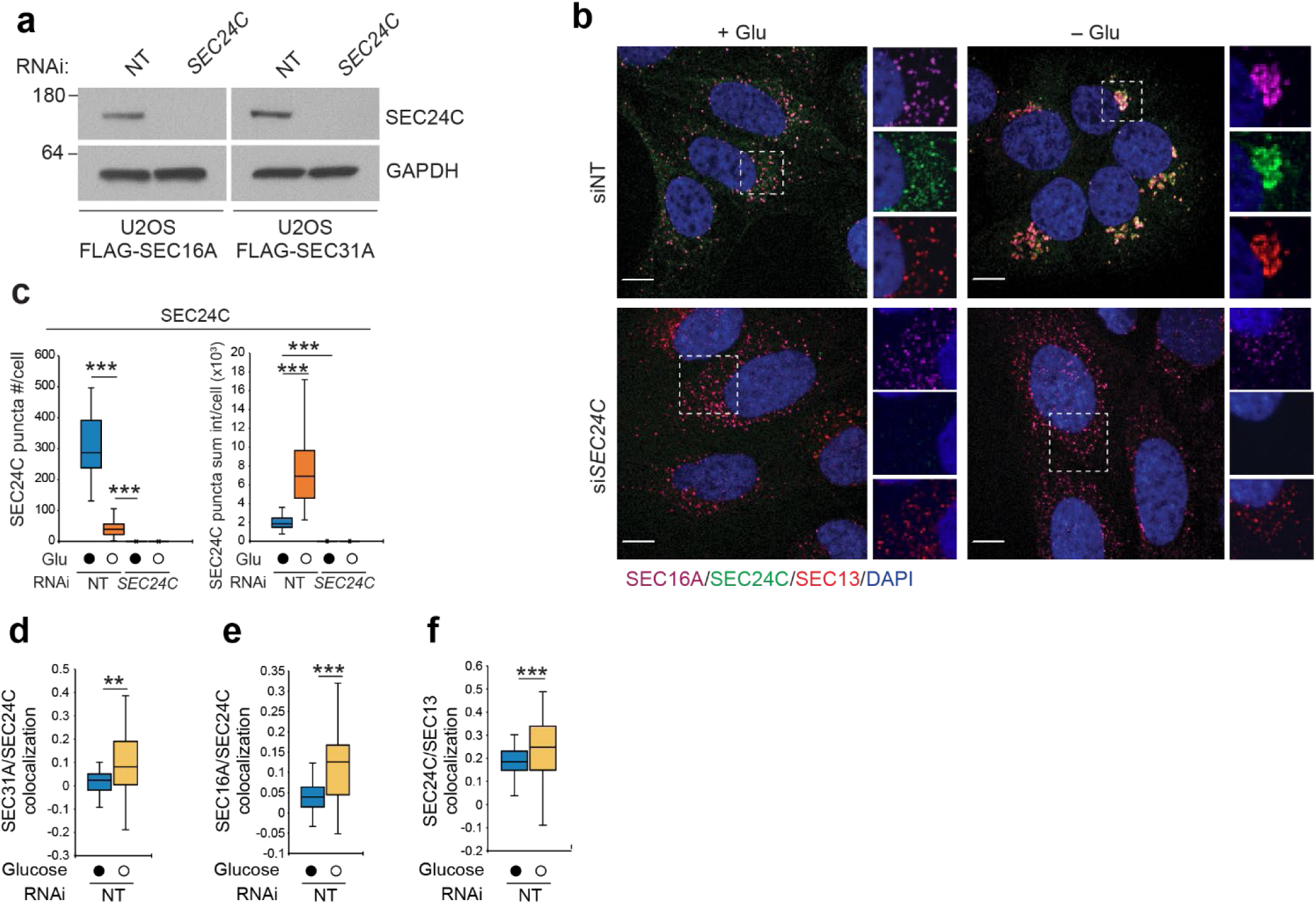
Supplemental data for Fig. 5. (**a**) Immunoblot showing siRNA knockdown efficiency of *SEC24C* in U2OS FLAG-SEC16A and FLAG-SEC31A knock-in cells. NT, non-targeting siRNA. **(b–f)** U2OS FLAG-SEC16A knock-in cells were transfected with either siNT or si*SEC24C*. After 48 h, cells were incubated in Dulbecco’s Modified Eagle Medium with or without glucose for 1 h followed by antibody staining with FLAG, SEC24C, and SEC13. Representative immunofluorescence images (**b**) taken after staining cells show changes in the distribution of the COPII components after glucose starvation. (**c**) Quantification of SEC24C puncta number and intensity data are shown as mean ± S.E.M., n >30 cells. Pearson’s correlation coefficients between the indicated protein pairs reflect the degree of colocalization: SEC31A/SEC24C (**d**), SEC16A/SEC24C (**e**), and SEC24C/SEC13 (**f**). Statistical significance was assessed using a two-tailed paired Student’s *t*-test. ***P < 0.001, **P < 0.01. Scale bar, 10 μm.

**Extended Data Fig. 6:**
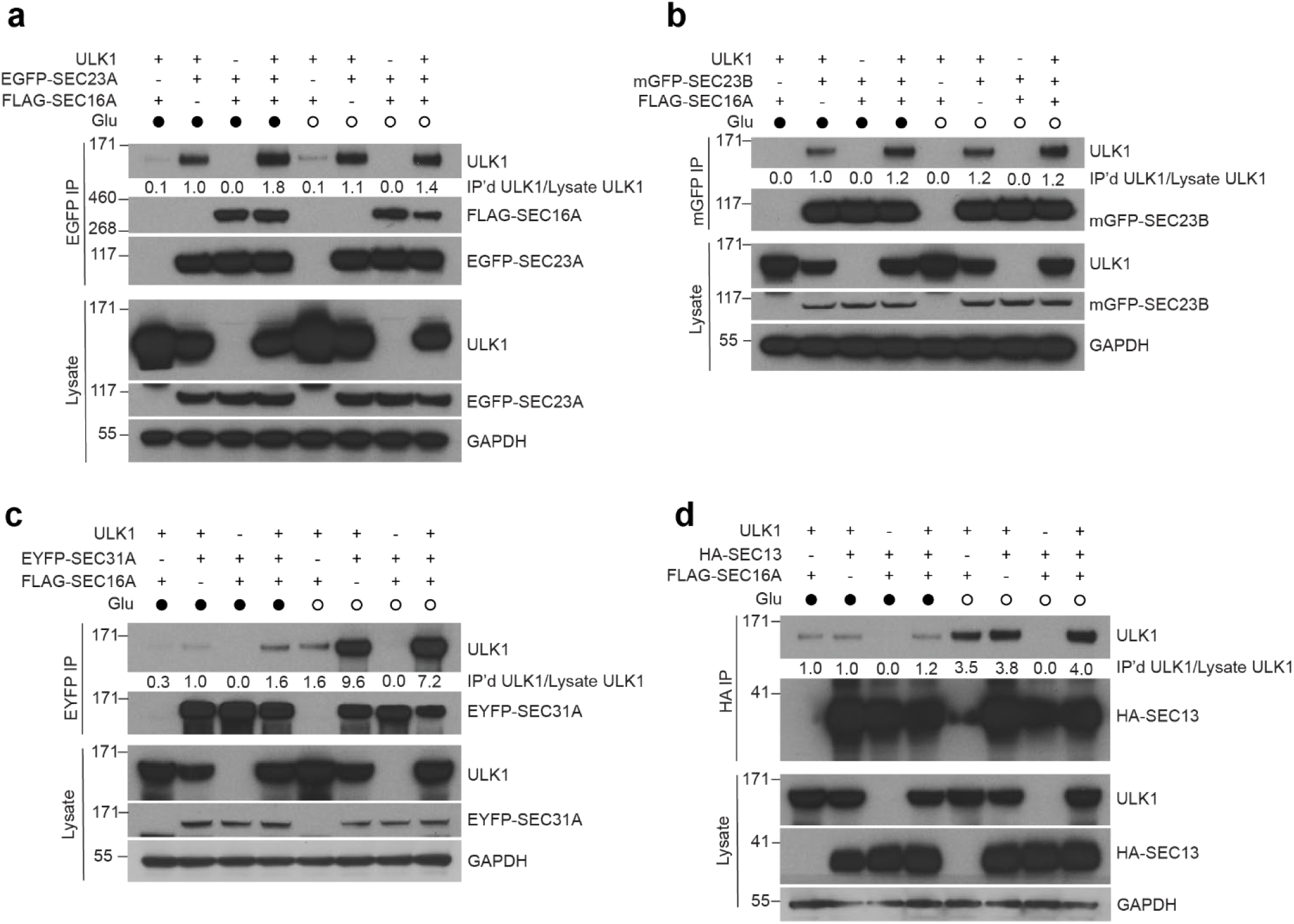
Supplemental data for Fig. 6. **(a–d)** HEK293T cells were transfected with the indicated expression vectors. After 48 h, cells were incubated in Dulbecco’s Modified Eagle Medium (DMEM) with or without glucose for 1 h followed by GFP or hemagglutinin (HA) immunoprecipitation. Representative immunoblot analyses demonstrate the level of molecular interaction between the indicated COPII components and ULK1.

**Extended Data Fig. 7:**
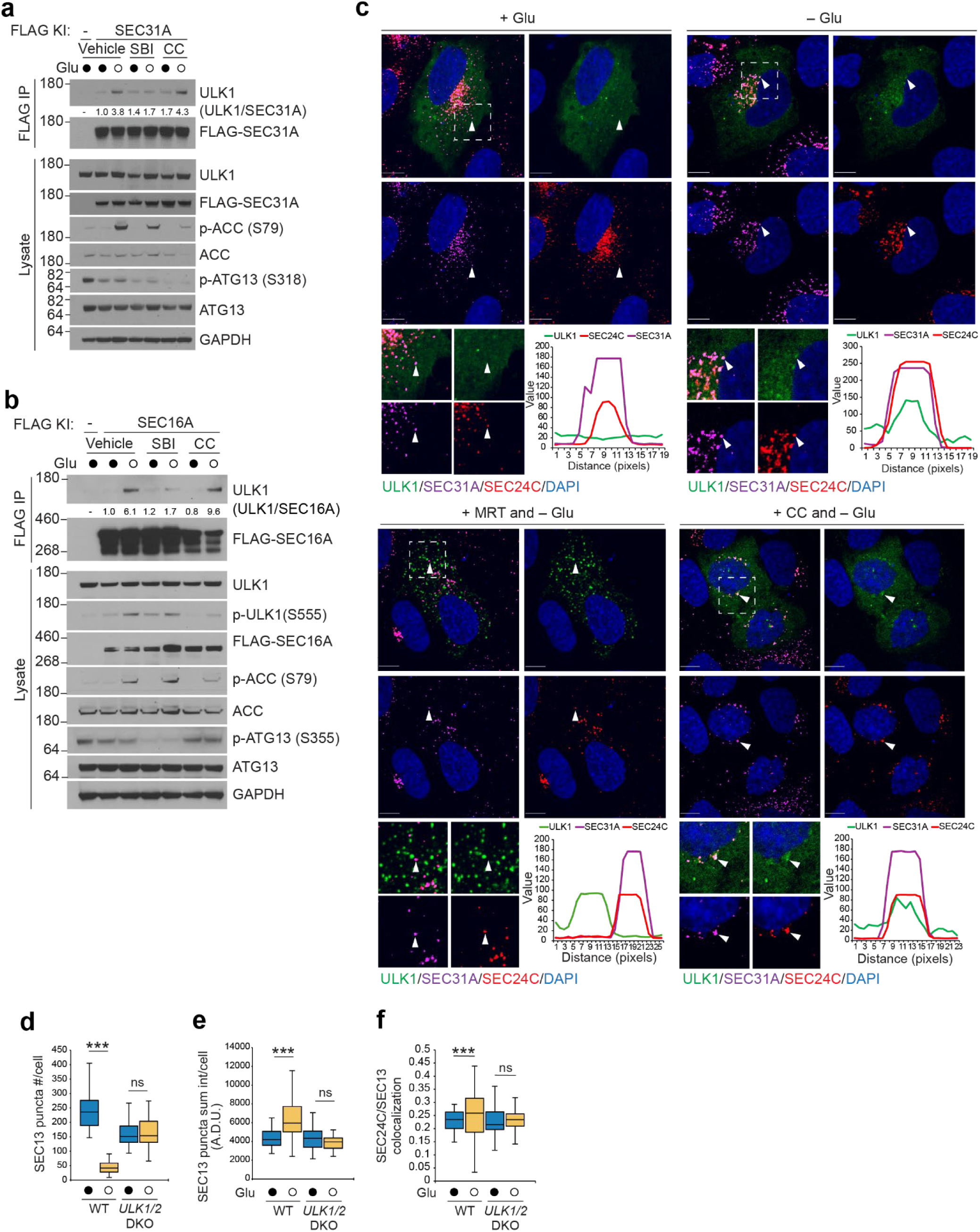
Additional supplemental data for Fig. 6. **(a, b)** U2OS FLAG-SEC31A knock-in (**a**) or FLAG-SEC16A knock-in cells (**b**) were pretreated with SBI 0206965 (10 µM) or compound C (20 µM) for 1 h, then incubated in Dulbecco’s Modified Eagle Medium (DMEM) with or without glucose for 1 h followed by FLAG immunoprecipitation. Representative immunoblot analyses demonstrate the level of molecular interaction between FLAG-SEC31A or FLAG-SEC16A and endogenous ULK1. **(c)** U2OS FLAG-SEC31A knock-in cells were transfected with EGFP-ULK1expression plasmids. After 48 h, cells were pretreated with MRT 68921 (1 µM) or compound C (5 µM) for 4 h, then incubated in DMEM with or without glucose for 1 h and stained with FLAG and SEC24C antibody. Representative immunofluorescence images taken after staining cells show changes in the distribution of the EGFP-ULK1 and COPII components after glucose starvation. Dotted squares indicate magnified area and color separate as individual color (bottom panel). The solid arrowheads indicate regions covered by line scans (bottom panels), which show the relative fluorescence intensities of ULK1 (green), SEC24C (red), and SEC31A (purple). **(d–f)** U2OS WT and *ULK1/2* DKO cells were incubated in DMEM with or without glucose for 1 h followed by antibody staining with SEC24C and SEC13. Representative immunofluorescence images (**Fig. 6, m**) taken after staining cells show changes in the distribution of the COPII components after glucose starvation. SEC13 puncta number (**d**), intensity (**e**), and the degree of co-localization between ULK1 and SEC31 as measured by Pearson’s correlation coefficient (**f**) were quantified and are presented as mean ± S.E.M. (n > 30 cells per condition). Statistical significance was analyzed using one-way ANOVA followed by Holm-Sidak multiple comparisons. ***P < 0.001; ns, not significant. Scale bar, 10 μm. A.D.U., analog-to-digital unit.

We then extended our analysis to examine how glucose starvation affects interactions between overexpressed ULK1 and additional COPII components, specifically SEC23A, SEC23B, SEC31A, and SEC13, in the presence or absence of SEC16A (Extended data Fig. 6a–d). Consistent with previous studies^62, 63^, we detected interactions between ULK1 and SEC23A or SEC23B under nutrient-replete conditions. Our analysis further showed that these interactions were independent of SEC16A expression and remained largely unchanged during glucose starvation. ULK1 did not show any specific interaction with SEC13 under any condition. In contrast, we observed a weak interaction between ULK1 and SEC31A under nutrient-replete conditions that became notably stronger during glucose starvation, independent of SEC16A expression. Together, these findings indicate that ULK1 can be recruited by several COPII components, including SEC23A, SEC23B, SEC16A, and SEC31A, but its interactions with SEC16A and SEC31A are particularly responsive to changes in glucose availability.

We next evaluated whether ULK1 kinase activity and AMPK-mediated phosphorylation were required for the glucose starvation–induced stabilization of the interactions between ULK1 and SEC16A or SEC31A. Using the KD (K46A) and AMPK phosphorylation-resistant (4SA) ULK1 mutants, we observed that the starvation-induced increases in ULK1–SEC16A and ULK1–SEC31A interactions persisted with the AMPK phosphorylation-resistant mutant but were abolished with the KD mutant, indicating that these interactions depend on ULK1 kinase activity but not direct AMPK phosphorylation (Fig. 6a–d). These findings were recapitulated using the CRISPR-engineered SEC16A-FLAG and SEC31A-FLAG knock-in lines, in which endogenous ULK1 interactions with SEC16A and SEC31A were similarly stabilized by glucose starvation (Extended Data Fig.7 a, b). Importantly, this stabilization was sensitive to the ULK inhibitor SBI 0206965 but not to the AMPK inhibitor Compound C, further suggesting a direct role for ULK1 (but not AMPK) activity. Consistent with these biochemical interactions, we observed enhanced recruitment of ULK1 to ERES components in glucose-starved U2OS cells, as shown by increased colocalization of overexpressed ULK1 with endogenous SEC31A by confocal microscopy (Fig. 6e–g). This recruitment was also blocked by ULK inhibitor MRT68921, but not by Compound C (Fig. 6h, i, and Extended data Fig. 7c).

Given the central role of SEC24C in mediating COPII reorganization, we next asked whether ULK1/2 activity influences the interaction of SEC24C with other COPII components under glucose starvation. Remarkably, in ULK1/2 DKO U2OS cells, the starvation–induced stabilization of SEC24C–SEC16A and SEC24C–SEC31A interactions observed in WT cells was lost (Fig. 6j–l). This underscores ULK1/2’s role in orchestrating the molecular interactions necessary for COPII complex remodeling during nutrient stress.

To directly test the consequences of ULK1/2 loss on COPII organization, we examined the subcellular distribution of COPII components in *ULK1/2* DKO U2OS cells. The characteristic decrease in SEC24C puncta number, increase in sum intensity, and enhanced co-localization with SEC13 observed during glucose starvation were all abolished in *ULK1/2* DKO cells (Fig. 6m–o and Extended data Fig. 7d–f). These defects were rescued by re-expressing WT ULK1 or the AMPK phosphorylation-resistant mutant, but not by re-expressing the KD mutant (Fig. 6p, q), confirming that ULK1 kinase activity (but not its phosphorylation by AMPK) is essential for mediating COPII reorganization in response to nutrient stress.

Together, these findings reveal that ULK1’s kinase activity regulates both its recruitment to ERES and the SEC24C-dependent remodeling of COPII machinery, thereby enabling adaptive reorganization of the secretory pathway during glucose starvation. The observation that ULK1-driven COPII stabilization at ERES is insensitive to AMPK inhibition suggests that this remodeling represents an initial, AMPK-independent phase of the cellular response to glucose deprivation, potentially setting the stage for later AMPK-dependent changes in trafficking dynamics.

### ULK1-Mediated Phosphorylation of SEC31A Regulates COPII Dynamics and Trafficking Inhibition During Glucose Starvation

To characterize the mechanism underlying how ULK1 signaling coordinates COPII complex assembly and trafficking inhibition in response to glucose starvation, we investigated whether SEC31A, given its enhanced interaction with ULK1 under glucose starvation, might serve as one of ULK1’s direct substrates. First, we transiently overexpressed FLAG-tagged SEC31A together with either WT or KD ULK1 in HEK293T cells. Following glucose starvation and/or treatment of cells with the phosphatase inhibitor calyculin A^64^, we immunoprecipitated FLAG-tagged SEC31A from cell lysates and analyzed the samples via Phos-tag gel electrophoresis. We found an ULK1-dependent mobility shift of SEC31A in the presence of calyculin A, indicating phosphorylation (Fig. 7a). Unexpectedly, this shift was observed regardless of glucose availability (Fig. 7a).

**Figure 7.**
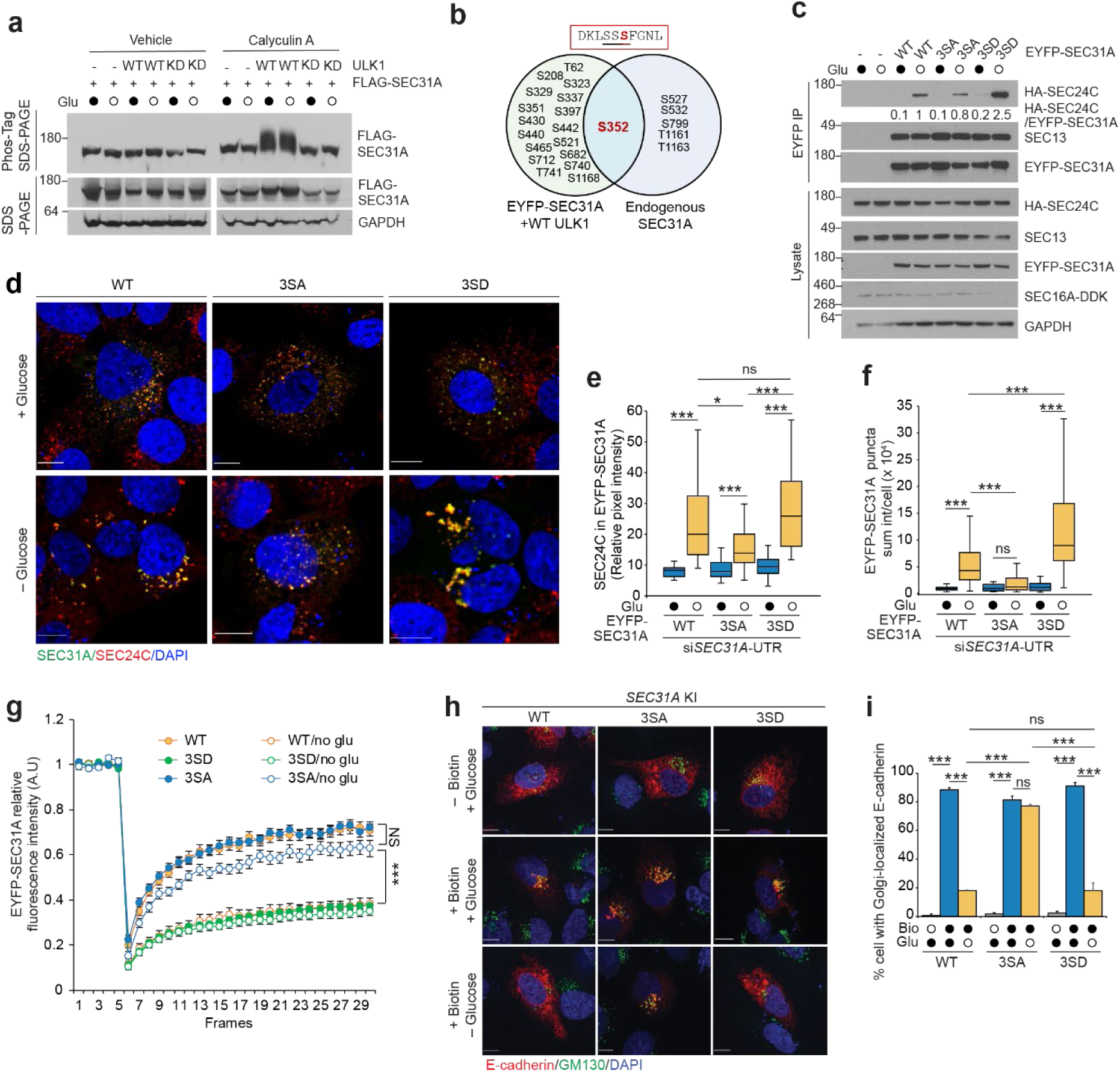
ULK1-mediated phosphorylation of SEC31A is required for COPII dynamics and trafficking inhibition during glucose starvation. **(a)** HEK293T cells were co-transfected with FLAG-SEC31A and either empty vector (EV), wild-type (WT) ULK1 or kinase-dead (KD) K46A ULK1. After 48 h, cells were incubated in Dulbecco’s Modified Eagle Medium (DMEM) with or without glucose and/or calyculin A (80 nM) for 1 h. FLAG immunoprecipitated protein samples were separated in SuperSep Phos-tag (50 µmole/L) 7.5% gel and 4%–12% SDS-PAGE gel. Representative immunoblot analyses demonstrate the level of SEC31A phosphorylation. **(b)** Two complementary phosphoproteomics approaches were used to identify ULK1- and starvation-dependent phosphorylation sites on SEC31A. The Venn diagram illustrates phosphorylated residues detected on EYFP-SEC31A immunoprecipitated from HEK293T cells co-transfected with WT ULK1 (but not with empty vector or kinase-dead ULK1) (left circle), and phosphorylated residues detected on endogenous SEC31A immunoprecipitated from FLAG-SEC31A knock-in U2OS cells cultured with or without glucose (right circle). S352 was the only phosphorylation site found in both analyses, indicating it is a common site under both ULK1 expression and physiologic conditions. **(c)** ULK1/2 double knockout U2OS cells were co-transfected with plasmids encoding HA-SEC24C, SEC16A-DDK, and either wild-type (WT), phospho-deficient (3SA), or phospho-mimetic (3SD) EYFP-SEC31A, as indicated. Representative immunoblots show the interaction levels between each form of EYFP-SEC31A and HA-SEC24C. **(d–f)** FLAG-SEC16A knock-in U2OS cells were transfected with *siSEC31A* and with plasmids encoding RNAi resistant EYFP-tagged WT, 3SA, or 3SD SEC31A. After 48 h, cells were incubated in DMEM with or without glucose for 1 h and stained with SEC24C antibody. Representative immunofluorescence images (d) taken after staining cells show changes in the distribution of the COPII component after glucose starvation. SEC24C intensity in SEC31A puncta (e) and SEC31A puncta intensity (f) were measured and plotted (mean ± S.E.M., n > 32). **(g)** WT U2OS cells were transfected with plasmids encoding EYFP-tagged WT, 3SA, or 3SD SEC31A. After 48 h, cells were incubated in DMEM with or without glucose for 1 h followed by measuring EYFP-SEC31A mobility with fluorescence recovery after photobleaching assay (mean ± S.E.M., n > 22 puncta per condition). Images were acquired every 5 seconds (5 sec per frame). The mobile fractions of EYFP-SEC31A under various conditions were as follows: WT with glucose, 63.4%; WT without glucose, 30.2%; 3SA mutant with glucose, 63.4%; 3SA mutant without glucose, 56.8%; 3SD mutant with glucose, 29.6%; and 3SD mutant without glucose, 27.6%. **(h, i)** FLAG-SEC31A knock-in, FLAG-SEC31A 3SA knock-in, and FLAG-SEC31A 3SD knock-in U2OS cells were transfected with E-cadherin RUSH construct. After 48 h, cells were incubated in DMEM with or without glucose for 1 h, followed by biotin treatment for 30 min. Representative immunofluorescence images (h) taken after staining cells using an anti-GM130 antibody and DAPI show changes in the subcellular distribution of the reporters after biotin treatment and/or glucose starvation. (i) Percentage of cells with Golgi-localized E-cadherin (mean ± S.E.M., n=3 biological replicates, >38 cells per experiment) in the knock-in U2OS cell lines cultured with or without glucose. Statistical significance was analyzed using one-way ANOVA followed by Tukey’s multiple comparisons. ***P < 0.001, *P < 0.05; ns, not significant. Scale bar, 10 μm.

**Extended Data Fig. 8:**
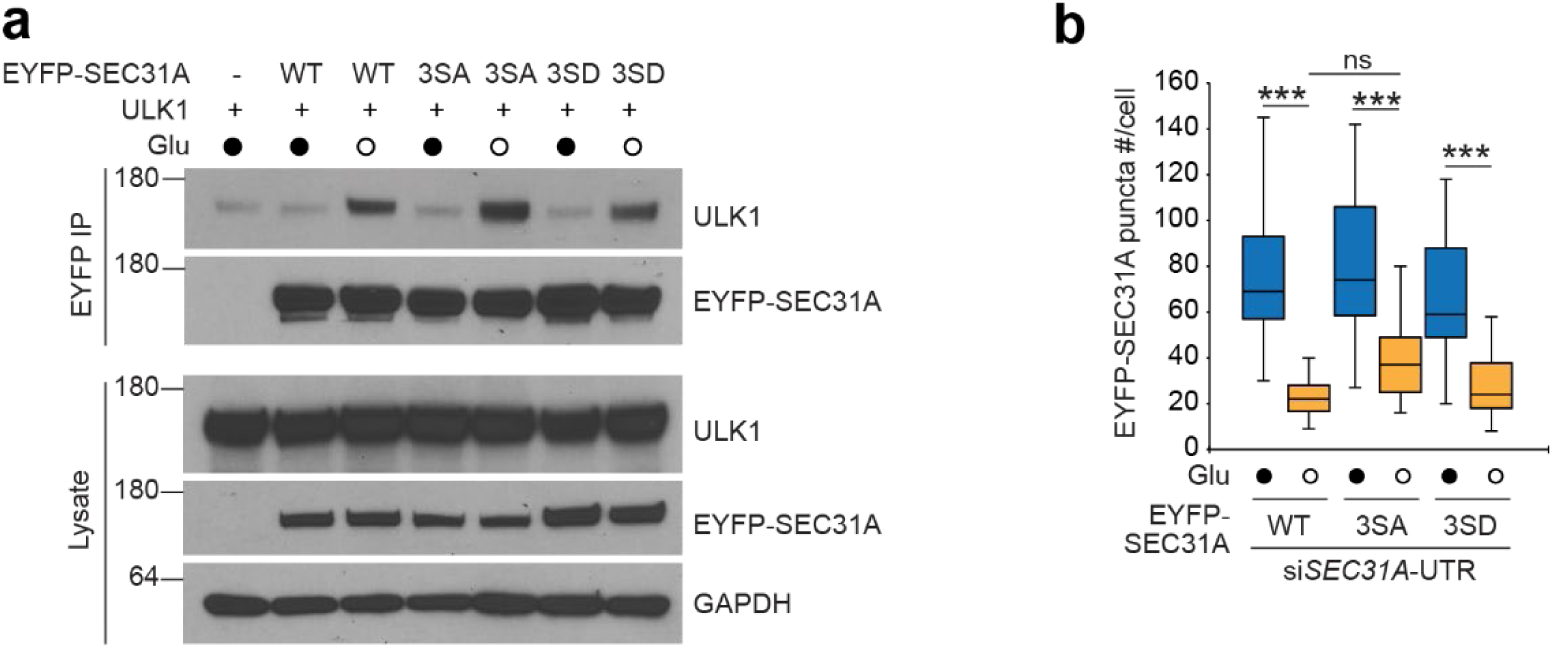
Supplemental data for Fig. 7. **(a)** HEK293T cells were transfected with ULK1 and EYFP-SEC31 wild-type (WT) or mutants (3SA or 3SD) as indicated. After 48 h, cells were incubated in Dulbecco’s Modified Eagle Medium (DMEM) with or without glucose for 1 h followed by EYFP immunoprecipitation. Representative immunoblot analyses demonstrate the level of molecular interaction between EYFP-SEC31A and ULK1. **(b)** FLAG-SEC16A knock-in U2OS cells were transfected with *siSEC31A* and RNAi-resistant EYFP-tagged WT, 3SA, or 3SD SEC31A. After 48 h, cells were incubated in DMEM with or without glucose for 1 h and stained with SEC24C antibody. Representative immunofluorescence images taken after staining cells show changes in the distribution of EYFP-SEC31A after glucose starvation. EYFP-SEC31A puncta were counted and plotted (mean ± S.E.M., n>32). Statistical significance was analyzed using one-way ANOVA followed by Tukey’s multiple comparisons. ***P < 0.001; ns, not significant.

**Table S2.** ULK1-regulated phosphorylation sites on SEC31: motif scores, conservation, and mass spectrometry results. This supplemental table summarizes the analysis of phosphorylation sites on SEC31A, with an emphasis on ULK1-dependent regulation. ULK1 motif-matching scores (PSSM) and evolutionary conservation across vertebrate SEC31A were determined using a motif prediction program^84, 85^. Phosphorylation status was evaluated by mass spectrometry in two experimental contexts: (1) overexpressed EYFP-tagged SEC31A isolated from cells co-expressing empty vector (EV), wild-type (WT) ULK1, or kinase-dead (KD) ULK1; and (2) endogenous FLAG-tagged SEC31A from cells cultured in Dulbecco’s Modified Eagle Medium with or without glucose for 1 h to model nutrient stress. By integrating motif prediction, conservation, and empirical phosphorylation data, this data highlights candidate ULK1-regulated sites on SEC31A and their sensitivity to cellular nutrient stress. Although several ULK1-dependent phosphorylation sites (identified in the overexpression context) exhibit high PSSM scores, S352 is the only site for which mass spectrometry detected phosphorylation on endogenous SEC31A. Abbreviations: D, detected; ND, not detected. See accompanying Excel file for complete data.

To identify specific ULK1-dependent phosphorylation sites, we performed phosphoproteomic analysis on EYFP-SEC31A immunoprecipitated from HEK293T cells transiently overexpressing EYFP-SEC31A along with either WT or KD ULK1. This analysis uncovered numerous ULK1-dependent phosphorylation sites on SEC31A (Fig. 7b and Table S2). To determine which sites are phosphorylated under physiological conditions, we conducted phosphoproteomic profiling of endogenous FLAG-tagged SEC31A immunoprecipitated from CRISPR knock-in cells cultured with or without glucose. While multiple phosphorylation sites were identified on endogenous SEC31A, no glucose-dependent changes were observed, and only one site, S352, was consistently detected in both experimental approaches (Fig. 7b and Table S2). This highlights S352 as a physiologically relevant site that can be phosphorylated in a ULK1-dependent manner. S352 is part of a cluster of three serine residues (Fig. 7b), which led us to generate expression constructs in which these three serines were mutated to alanine (3SA, phospho-deficient) or aspartate (3SD, phospho-mimetic). These constructs were then used to assess the functional consequences of SEC31A phosphorylation.

Since our data indicated that ULK1 activity was required to stabilize both the ULK1–SEC31A (Fig. 6c, d) and SEC31–SEC24C (Fig. 6j, k) interactions during glucose starvation, we first examined whether phospho-deficient or phospho-mimetic SEC31A mutants affected these protein interactions. The interaction between SEC31A and ULK1 was unchanged for both mutant forms compared to WT SEC31A (Extended data Fig. 8a). However, the SEC31A–SEC24C interaction was sensitive to the phosphorylation state of SEC31A during glucose starvation: the phospho-deficient mutant showed slightly reduced complex formation with SEC24C, while the phospho-mimetic mutant displayed increased association (Fig. 7c). These results suggest that ULK1-mediated phosphorylation of SEC31A promotes stabilization of the SEC31A–SEC24C complex during glucose starvation.

We found that the recruitment of endogenous SEC24C to puncta formed by overexpressed SEC31A was also dependent on SEC31A phosphorylation status, with markedly reduced colocalization observed for the phospho-deficient mutant compared to WT or the phospho-mimetic mutant (Fig. 7d, e). Notably, although the glucose starvation-induced decrease in the number of SEC31A+ puncta was not affected by SEC31A phosphorylation status (Extended data Fig. 8b), the size of puncta formed by the phospho-deficient SEC31A mutant did not increase in glucose-starved cells, whereas those formed by the phospho-mimetic mutant were larger than those observed with WT SEC31A upon starvation (Fig 7f). Thus, ULK1-dependent phosphorylation of SEC31A not only stabilizes its interaction with SEC24C but also facilitates their spatial colocalization and the formation of larger ERES puncta under nutrient stress.

To further probe the function of SEC31A phosphorylation, we performed fluorescence recovery after photobleaching on puncta generated by each of these constructs. The phospho-mimetic mutant displayed delayed recovery and a lower mobile fraction under both nutrient-replete and glucose-starved conditions, mirroring the behavior of WT SEC31A in glucose-starved cells (Fig. 7g). Importantly, the delay in recovery and reduction in mobile fraction seen with WT SEC31A during glucose starvation were not observed in the phospho-deficient mutant (Fig. 7g), highlighting the functional significance of SEC31A phosphorylation for ERES dynamics.

Finally, to determine whether ULK1-mediated phosphorylation of SEC31A contributes to the trafficking inhibition observed during glucose starvation, we used CRISPR/Cas9 editing to generate SEC31A phospho-deficient (3SA) and phospho-mimetic (3SD) knock-in U2OS cell lines. When these cells were transfected with the E-cadherin RUSH construct, both phospho-deficient and phospho-mimetic SEC31A supported normal trafficking of E-cadherin under nutrient-replete conditions (Fig. 7h, i). However, upon glucose starvation, the phospho-mimetic mutant exhibited robust inhibition of E-cadherin trafficking, similar to WT SEC31A; this starvation-induced inhibition was not observed in cells expressing the phospho-deficient mutant (Fig. 7h, i).

Collectively, these data reveal that ULK1-mediated phosphorylation of SEC31A regulates its interaction and colocalization with SEC24C, alters COPII complex dynamics, and is required for the full inhibition of ER-to-Golgi trafficking in response to glucose starvation. This phosphorylation-dependent modulation of COPII assembly represents a key mechanistic step linking nutrient sensing to adaptive secretory pathway remodeling.

## Discussion

Cellular adaptation to metabolic stress is fundamental for survival and plays a crucial role in the progression of diseases such as cancer. Diverse metabolic challenges, such as glucose deprivation and hypoxia, drive extensive changes in the composition of proteins at the cell surface, with downstream effects on cell signaling, nutrient uptake, and interactions with the microenvironment^5, 8–10^. Although nutrient-sensing kinases such as mTOR and AMPK are known to regulate autophagy and membrane trafficking^65–68^, their direct impact on the selective export of specific cargoes to the plasma membrane has remained unclear. Prior studies have implicated mTOR signaling at the Golgi in unconventional protein secretion^69, 70^ and AMPK activation in the reduction of surface levels of adhesion proteins^71^, but the molecular details linking these pathways to cargo-specific secretory trafficking have not been fully resolved.

In this study, we identify a previously unrecognized mechanism by which glucose starvation regulates ER-to-Golgi trafficking, directly linking metabolic signals to dynamic remodeling of the cell surface proteome. We show that glucose starvation, but not amino acid starvation, selectively impairs the ER export of key adhesion molecules such as E-cadherin, while other cargoes remain unaffected. This process is mediated by a non-canonical AMPK-ULK1/2-SEC24C signaling axis, independent of autophagy-related pathways. The associated remodeling of the cell surface proteome is accompanied by increased cell motility and metastatic potential, consistent with established roles for E-cadherin loss in epithelial–mesenchymal transition and metastatic dissemination^45, 72^. Importantly, our findings support emerging models in which dynamic regulation of E-cadherin trafficking, rather than its complete loss, underlies cancer cell plasticity and metastatic competence^73, 74^. Our findings reveal how metabolic cues orchestrate targeted changes in secretory trafficking, establishing a new framework for understanding nutrient-driven cell surface reprogramming and its potential impact on cancer progression.

A key discovery of our study is the requirement for both AMPK and ULK1/2 in this selective secretory blockade, which occurs independently of both canonical and non-canonical autophagy. While AMPK–ULK1 signaling is classically linked to autophagy initiation^49, 50^, and though ULK1 has been implicated in the diversion of COPII components for autophagosome biogenesis^62, 75–77^, our work demonstrates a distinct, autophagy-independent role for ULK1/2 in modulating ER export in response to starvation. Notably, SEC24C emerges as a central mediator, required for the glucose-dependent inhibition of E-cadherin trafficking. While SEC24C has established roles in cargo export and ER homeostasis^44, 48, 78, 79^, we have revealed a previously unreported function for SEC24C in glucose starvation–induced, cargo-selective secretory control.

Mechanistically, glucose starvation triggers SEC24C-dependent clustering and stabilization of COPII components; ULK1 is robustly recruited to these sites. This reorganization is contingent on ULK1 kinase activity and the phosphorylation of SEC31A, but it does not depend on AMPK-mediated phosphorylation of ULK1. Consistent with this, expression of the AMPK-resistant ULK1-4SA mutant supports ULK1 recruitment and stabilization at ERES but fails to support glucose starvation–induced trafficking inhibition. In contrast, ULK1 kinase activity and SEC31A phosphorylation are essential for both COPII stabilization and trafficking inhibition. Although SEC31A phosphorylation occurs under both basal and stress conditions, its role in trafficking inhibition is unmasked specifically during glucose starvation, suggesting a “priming” mechanism that enables rapid adaptation to metabolic cues. This two-tiered regulatory system allows cells to fine-tune secretory trafficking and surface composition in response to fluctuating nutrient availability.

Importantly, these findings reveal that COPII stabilization at ERES is likely an early and AMPK-independent event that may prime cells for a subsequent, AMPK-dependent trafficking blockade. This suggests that distinct upstream inputs converge on ULK1, enabling independent regulation of ERES remodeling and secretory inhibition as temporally and mechanistically separate steps during glucose starvation. However, the mechanism by which glucose starvation stabilizes the interaction between ULK1 and SEC31A remains unresolved, indicating that additional factors or post-translational modifications may contribute to this process. Future studies will be needed to elucidate the upstream signals governing ULK1 recruitment to ERES, define the molecular determinants of cargo selectivity in this pathway, and explore its broader relevance in physiological and pathological contexts such as development, tissue homeostasis, and cancer progression.

Our findings establish a new paradigm in which glucose availability exerts selective, cargo-specific control over ER-to-Golgi trafficking via an autophagy-independent AMPK-ULK1/2-SEC24C axis. This mechanism enables rapid remodeling of the cell surface proteome, with direct implications for cell adhesion, migration, and thus, metastatic potential. Given the central role of glucose limitation in the tumor microenvironment^80–83^, our results suggest that targeting nutrient-sensitive secretory pathways could represent a novel strategy to limit cancer metastasis and modulate cellular adaptation to metabolic stress going forward.

## MATERIALS AND METHODS

### Cell Lines and Cell Culture

U2OS (Cat. No. HTB-96), HEK293T (Cat. No. CRL-3216), and 4T1 (Cat. No. CRL-2539) cells were obtained from American Type Culture Collection. Cells were mycoplasma free and verified by PCR analysis (VenorGeM Mycoplasma Detection Kit, PCR-based, Cat. No. MP0025, Sigma-Aldrich). U2OS, MEF, and HEK293T cells were cultured in Dulbecco’s Modified Eagle Medium (DMEM; Cat. No. 11965-092, Gibco) containing 10% fetal bovine serum (FBS; Cat. No. SH30396.03, Cytiva), 1XPenStrep (Cat. No. 15140-122, Gibco), and 1XGlutaMax-1 (Cat. No. 35050-061, Gibco). 4T1 cells were cultured in RPMI (Cat. No. 11875119, Gibco) containing 10% FBS and 1XPenStrep. For glucose starvation, 10% dialyzed FBS (Cat. No. 26400044, Gibco), 1XPenStrep, and 1XGlutaMax-1 was added to DMEM without glucose (Cat. No. 11966-025, Gibco). Pre-plated cells were washed three times with glucose-free complete medium and then cultured in complete medium containing glucose or no glucose for indicated time periods. For amino acid starvation, U2OS cells were cultured in Earle’s Balanced Salts (EBSS; Cat. No. E3024, Millipore Sigma) containing 10% dialyzed FBS, and 1XPenStrep for 2 h. All cells were incubated in a humidified environment containing 5% CO2 at 37°C. Cells were treated with 250 nM Torin 2 (Cat. No. 4248, R&D systems) for 6 h, 5 µM Compound C (Cat. No. B3252, Apexbio) for 4 h, 20 µM compound C for 1 h, 10 µM SBI 0206965 (Cat. No. HY-16966, MCE) for 1 h, 1 µM MRT 68921 (Cat. No. S7949, Selleckchem) for 24 h or 4 h, or 80 nM Calyculin A (Cat. No. 1336, TOCRIS) for 1 h as indicated.

### Transient Transfection and Generation of Genetically Modified Cells

Plasmid DNA transfection was performed using FuGENE 6 (Cat. No. E2691, Promega) for HEK293T cells or FuGENE HD (Cat. No. E2311, Promega) for U2OS cells and MEFs per the manufacturer’s protocol. Genetically modified cells were generated using CRISPR technology in the Center for Advanced Genome Engineering (St. Jude Children’s Research Hospital [St. Jude]). Briefly, 400,000 U2OS cells were transiently co-transfected with precomplexed ribonuclear proteins (RNPs) consisting of 50–150 pmol of each chemically modified sgRNA (IDT), 50 pmol of 3X NLS *Sp*Cas9 protein (St. Jude Protein Production Core), 400 ng of pMaxGFP (Lonza), and, if required, 3 μg of ssODN or 1 μg of dsDNA plasmid donor. The transfections were performed via nucleofection (Lonza, 4D-Nucleofector X-unit) using solution P3 and program CM-104 in a 20-μl cuvette according to the manufacturer’s recommended protocol. Single cells were sorted based on transfected cells (GFP+) 5 days post-nucleofection into 96-well plates containing prewarmed media and were clonally expanded. Clones were screened and verified for the desired modification using targeted deep sequencing analyzed with CRIS.py as previously described^86, 87^. Final clones tested negative for mycoplasma by the MycoAlert Plus Mycoplasma Detection Kit (Lonza) and were authenticated using the PowerPlex Fusion System (Promega) performed at the Hartwell Center (St. Jude).

Editing construct sequences and screening primers are listed in Table 1.

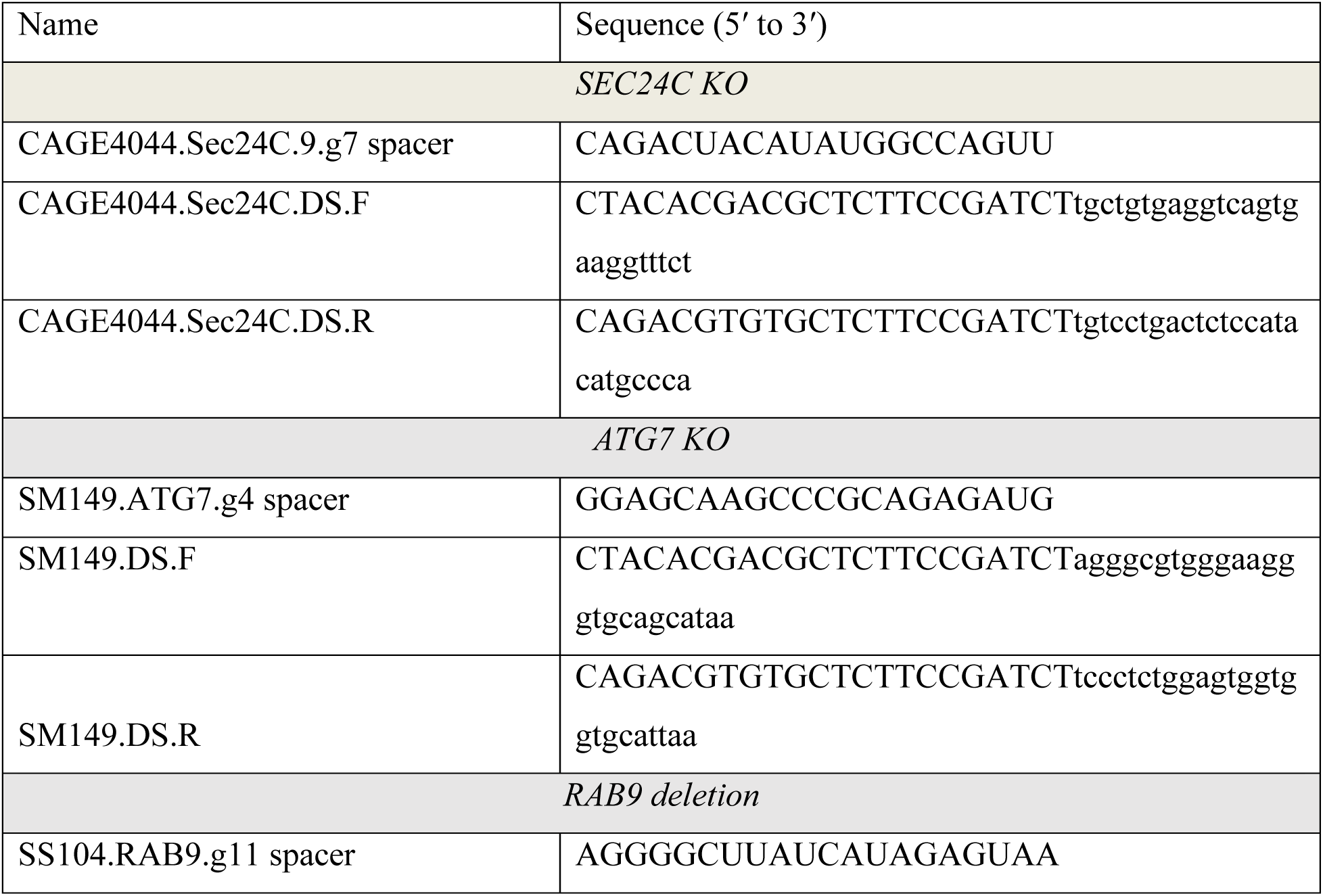
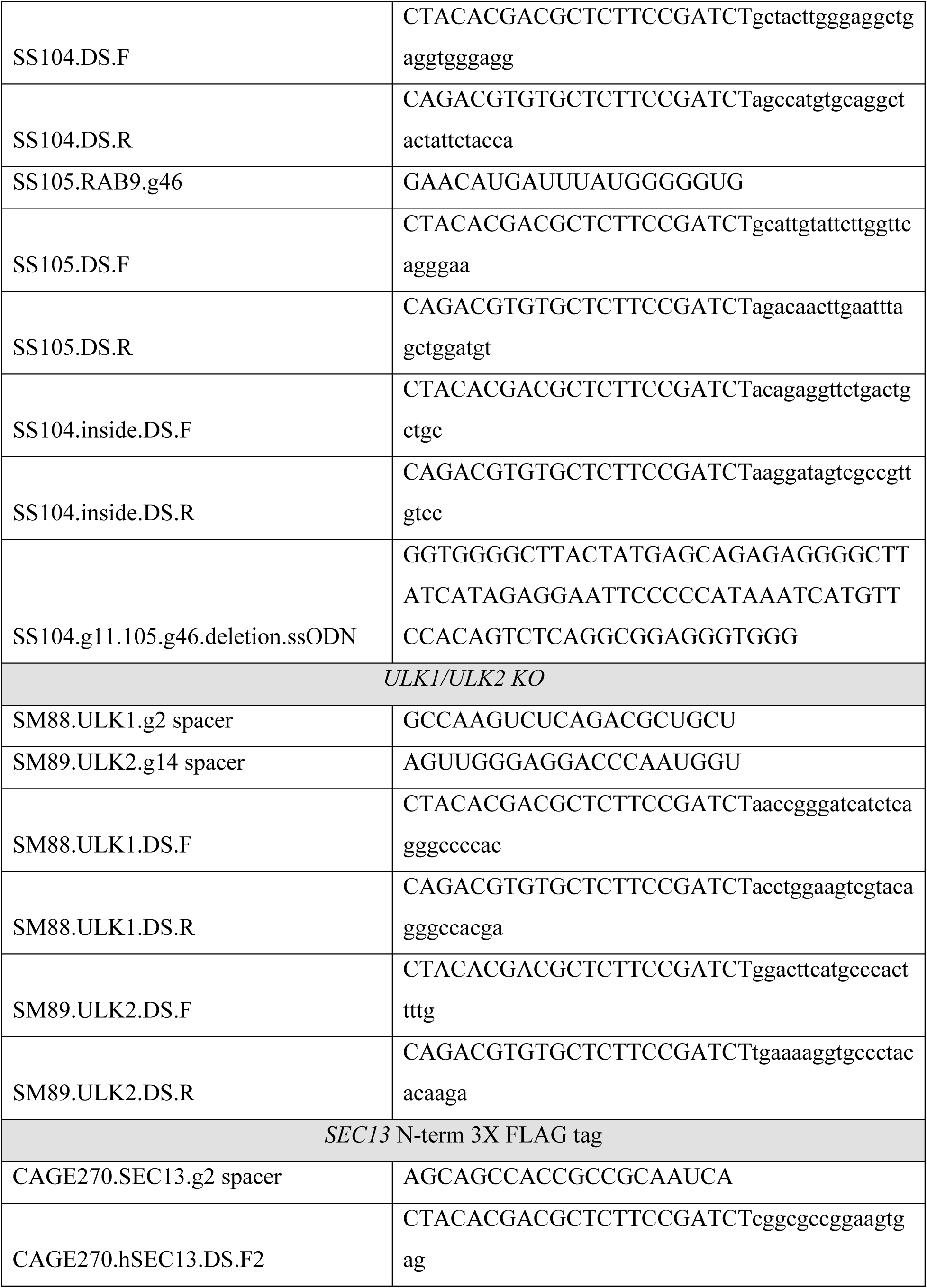
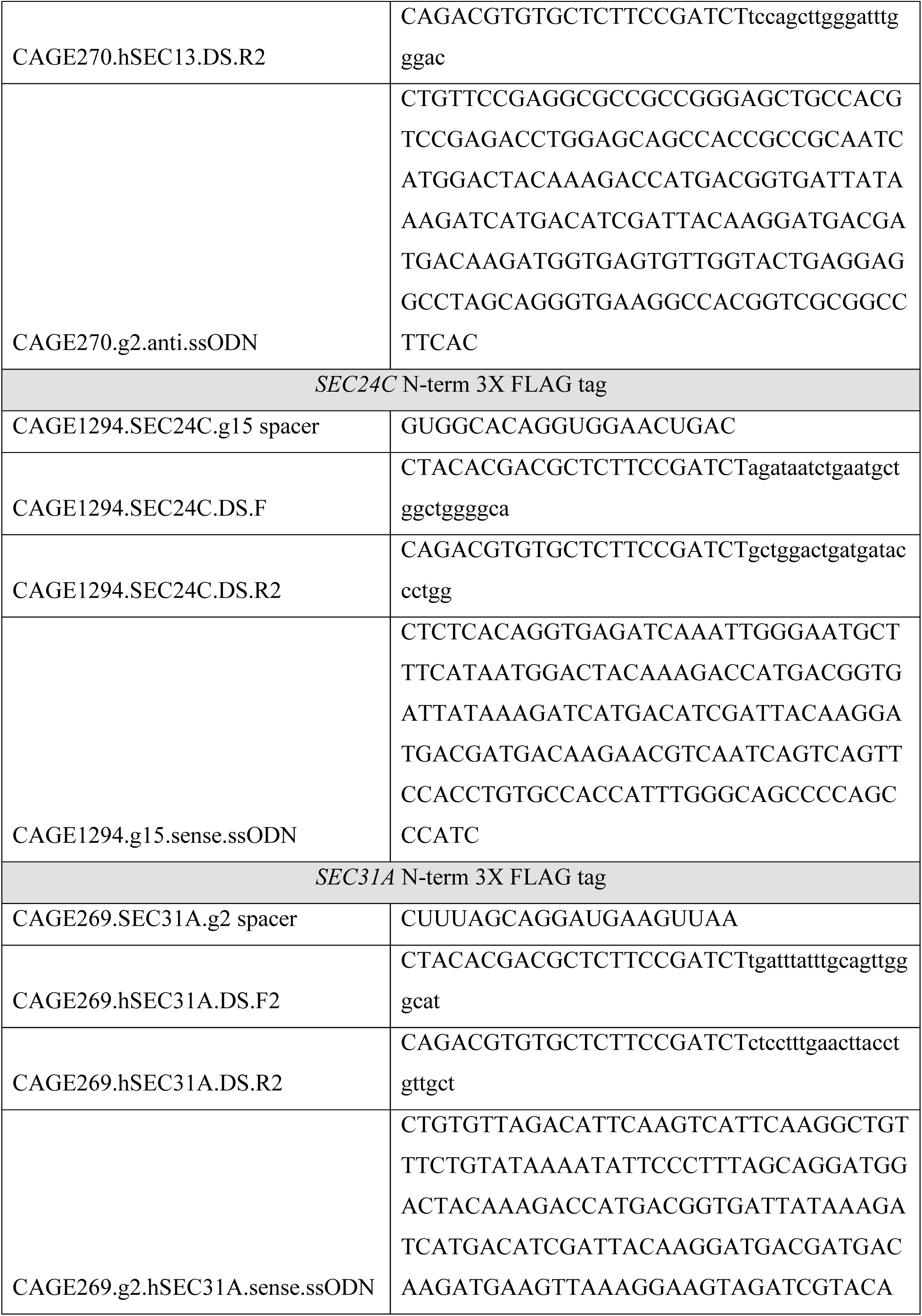
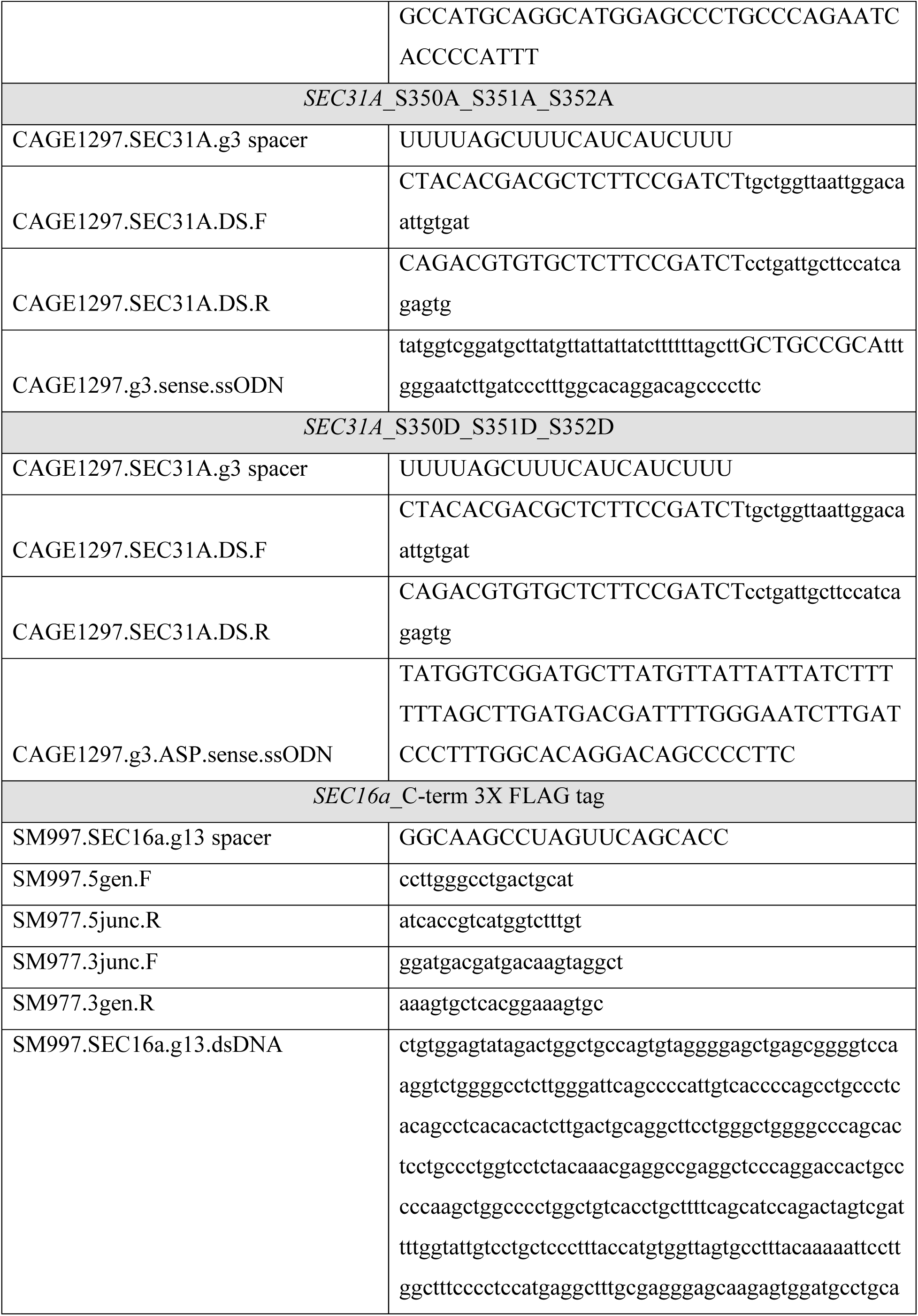
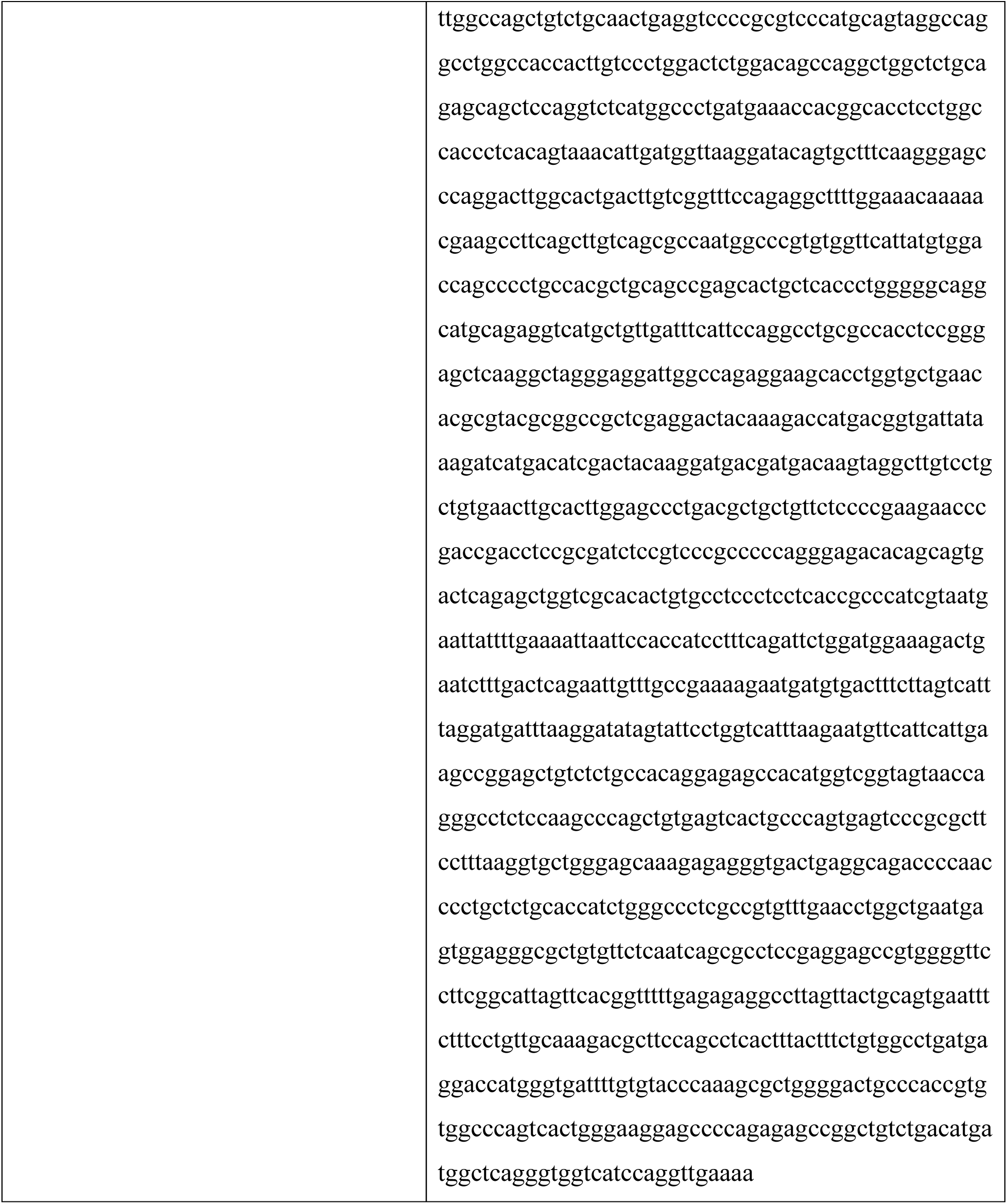

### Plasmids constructs

The pmGFP-SEC16A(L) (Cat. No. 15776), pFLAG-SEC31A (Cat. No. 42110), pEYFP-Sec31A (Cat. No. 66613), pEGFP-SEC23A (Cat. No. 66609), pmGFP-SEC23A (Cat. No. 66598), pHA-SEC13 pRK5 (Cat. No. 46332), and pMXs-IP-EGFP-mULK1 (Cat. No. 38193) were purchased from Addgene. The pCMV6-Entry-Sec16A-myc-DDK (Cat. No. RC223625) was purchased from OriGene Technologies. HA-tagged SEC24 constructs, HA-SEC24A, HA-SEC24B, HA-SEC24C, and HA-SEC24D were generously gifted by Dr. Randy Scheckman (Department of Molecular & Cell Biology, UC Berkeley, CA). The pcDNA3.1 backbone-based and retroviral pMSCV-IGFP-MII backbone-based WT ULK1, KD (K46A) mutant, and 4SA mutant (AMPK phosphorylation–resistant mutant) were generated for transient transfection and retrovirus generation, respectively, as described previously^61^. SEC31A phosphorylation-site mutants (3SA and 3SD) were introduced into the pEYFP–SEC31A using mutagenic oligonucleotides. SEC31A 3SA; forward 5’-CAAGTTGACAAGCTTGCAGCAGCTTTTGGGAATCTTG-3’, reverse 5’-CAAGATTCCCAAAAGCTGCTGCAAGCTTGTCAACTTG-3’, SEC31A 3SD; forward 5’-CAAGTTGACAAGCTTGATGATGATTTTGGGAATCTTG-3’, reverse 5’-CAAGATTCCCAAAATCATCATCAAGCTTGTCAACTTG-3’

### RNAi Knockdown

Knockdown experiments in U2OS cells and MEFs were performed using Lipofectamine RNAi Max (Cat. No. 13778-150, Thermo Fisher Scientific) per the manufacturer’s protocol and with the following siRNAs obtained from Horizon Discovery: ON-TARGETplus Non-targeting pool (D-001810-10-05), siGENOME Non-Targeting siRNA pool (D001206-13-05), siGENOME Human *Sec24C* siRNA-individual (D-008467-18-0005), ON-TARGETplus Mouse *Sec24C* siRNA-SMARTpool (J-05952-12-0005), ON-TARGETplus Human *Sec24A* siRNA-SMARTpool (L-024405-01-0005), ON-TARGETplus Human *Sec24B* siRNA-SMARTpool (L-008299-02-0005), ON-TARGETplus Human *Sec24C* siRNA-SMARTpool (L-008467-02-0010), and ON-TARGETplus Human *Sec24D* siRNA-SMARTpool (L-008493-01-0005). The knockdown efficiency was confirmed by immunoblot analysis.

### Immunoprecipitation and Immunoblotting

Cell lysates obtained with Triton-based cell lysis buffer (40 mM HEPES, 120 mM NaCl, 1 mM EDTA, 1.5 mM Na3VO4, 50 mM NaF, 10 mM β-glycerophosphate, 20 mM MoO4, 0.5% Triton X-100, protease inhibitor [Cat. No. 11836170001, Sigma Aldrich]) and phosphatase inhibitor (Cat. No. 4906837001, SigmaAldrich). The lysates were incubated with anti-HA (Cat. No. 3724, Cell Signaling) or anti-GFP antibody (Cat. No. 6556, Abcam) overnight at 4°C and precipitated with protein G agarose beads (Cat. No. 20399, Thermo Fisher Scientific). Anti-FLAG M2-agarose beads (Cat. No. A2220, Sigma-Aldrich) were used for immunoprecipitation of FLAG-tagged proteins. The beads were washed three times with cold Triton-based buffer and incubated at 95°C for 5 min in sodium dodecyl sulfate (SDS) sample buffer (Cat. No. S3401, Sigma-Aldrich).

Proteins were electrophoretically separated on 4%–12% bis-Tris gels (Cat. No. NP0335BOX, Thermo Fisher Scientific). Proteins were then transferred to a PVDF membrane. After blocking the membrane with 5% skim milk, blots were probed with antibodies directed against the following targets: ULK1 (Cat. No. 8058, Cell Signaling), p-ULK1 (S555) (Cat. No. 5869, Cell Signaling), ATG7 (Cat. No. 2631, Cell Signaling), ULK2 (Cat. No. HPA009027, Sigma-Aldrich), ATG13 (Cat. No. SAB4200100, Millipore Sigma), p-ATG13 (S318) (Cat. No. 600-401-C49 Rockland), GFP (Cat. No. ab6556, Abcam), FLAG (Cat. No. F1804, Millipore Sigma), HA (Cat. No. 3724, Cell Signaling), GAPDH (Cat. No. G9545, Sigma Aldrich), p-ACC (S79) (Cat. No. 11818, Cell Signaling), ACC (Cat. No. 3676, Cell Signaling), p-S6 (S235/236) (Cat. No. 4858, Cell Signaling), S6 (Cat. No. 2317, Cell Signaling), SEC24C (Cat. No. 8531, Cell Signaling), SEC24A (Cat. No. 15958-1-AP, Proteintech), SEC24B (Cat. No. A304-876A, Bethyl), SEC24D (Cat. No. 14687, Cell Signaling), and RAB9 (Cat. No. 5133, Cell Signaling). Membranes were then incubated with horseradish peroxidase–conjugated secondary antibodies, and bands were detected using chemiluminescence-detection kits (Cat. No. RPN2106, Cytiva).

### Retention Using Selective Hooks (RUSH) Assay

RUSH constructs, Str-KDEL-SBP-mCherry-Ecadherin (Cat. No. 65287), Str-KDEL-SBP-EGFP-GPI (Cat. No. 65294), and Str-KDEL_IRES_SBP-mCherry-VSVG (Cat. No. 203647) were purchased from Addgene. U2OS cells and MEFs were transiently transfected with the indicated RUSH construct using FuGENE 6 (Cat. No. E2691, Promega) per the manufacturer’s protocol. After 48 h, cells were incubated in DMEM with or without glucose for 1 h, followed by the addition of biotin (final conc. 40 µM) for 30 min to release the reporters from their ER hooks; cells were then stained using an anti-GM130 antibody (Cat. No. 610823, BD Trabsduction Laboratories) and DAPI to show changes in the subcellular distribution of the reporters after biotin addition.

### Immunofluorescence for Protein

To assess COPII puncta quantity, U2OS were incubated in complete growth media with or without glucose for 1 h, fixed in 4% formaldehyde, permeabilized with 0.5 % Triton X-100, blocked in 5% goat serum, and labeled with one of the following antibodies: anti-SEC24C (Cat. No. 14676, Cell Signaling), anti-SEC13 (Cat. No. MAB9055, R&D Systems), anti-FLAG (Cat. No. F1804, Millipore Sigma), or HA (Cat. No. 3724, Cell Signaling). Cells were then treated with secondary antibodies conjugated to Alexa-555 (Cat. No. A21428, Invitrogen, Thermo Fisher Scientific), Alexa-488 (Cat. No. A11008, Invitrogen, Thermo Fisher Scientific), or Alexa-647 (Cat. No. A11001, A21236, Invitrogen, Thermo Fisher Scientific). Alternatively, antibodies were labeled with Mix-n-Stain Antibody Labeling Kit (Biotium), CF^®^488A (Cat. No. 92273), CF^®^555 (Cat. No. 92274), or CF^®^647 (Cat. No. 92279) per manufacturers’ instructions. Images were acquired on a Marianas system (Intelligent Imaging Innovations) configured with a CSU-W1 (Yokogawa) spinning disk on a Zeiss Axio Observer (Zeiss) microscope. Images were collected with a Plan Apo 100X/1.40 NA objective (Zeiss) on a Prime 95B camera (Teledyne Photometrics), resulting in 110-nm pixels. Illumination was provided by either a 405-, 488-, 561-, or 638-nm laser (200/150/150/200 mW, Coherent) passing through the CSU-W1 with a Uniformizer for image acquisition. Channels were defined by a combination of a ZT405/488/561/640tpc-uf1 dichroic mirror (Chroma) and either a ET445/58m, ET525/50m, ET600/50m, or ET706/95m emission filter (Chroma), as appropriate. Image acquisition was controlled using Slidebook 2024 (Intelligent Imaging Innovations). The number or intensity of puncta was analyzed using Fiji^88^ or SlideBook 2024 software (Intelligent Imaging Innovations, Denver, CO, USA).

### Fluorescence Recovery After Photobleaching

U2OS cells were transfected with siRNA by TransIT-TKO (Cat. No. MIR 2154, Mirus Bio) for 24–30 h followed by transient transfection with 100 ng EYFP-SEC31A expression plasmid by Lipofectamine 3000 (Cat. No. L3000015, Thermo Fisher Scientific) in Lab-Tek II chambered #1.5 coverglass (Cat. No. 155382PK, Thermo Fisher Scientific) for 20 h. Prior to imaging, media were replaced with Fluoro-Brite DMEM (Thermo Fisher Scientific) supplemented with 5% FBS, 1XPenicillin/streptomycin (Cat. No. 15140-122, Gibco), and 1XGlutaMax^TM^-1 (Cat. No. 35050-061, Gibco) and imaged at 37°C. Exposure time was 200 ms and frame rate was 5 s per frame for 3–4 min. Confocal microscopy was conducted on a custom-built Nikon microscope equipped with a Yokogawa CSU-W1 Spinning Disk unit, XY galvo scanning module, and Tokai Hit STXG CO2 incubation system using a 60× (CFI APO TIRF, NA 1.49 oil) objective. Photobleaching experiments were performed on a Nikon microscope using XY galvo scanning module. All image analyses were performed using Fiji^88^. All intensity analyses were subjected to background subtraction. To obtain relative intensity profiles, background-subtracted intensity values from different conditions were normalized to that at the first time point or in control groups.

### Super Resolution Microscopy

FLAG-SEC31A knock-in or FLAG-SEC16A knock-in U2OS cells were seeded onto coverglass. The next day, cells were incubated in complete growth media with or without glucose for 1 h, fixed in 4% formaldehyde, permeabilized with 0.5 % Triton X-100, blocked in 5% goat serum, and then labeled with anti-FLAG (Cat. No. F1804, Millipore Sigma) and anti-SEC24C (Cat. No. 14676, Cell Signaling) antibody, or anti-FLAG (Cat. No. F1804, Millipore Sigma) and anti-SEC13 (Cat. No. MAB9055, R&D systems) antibody. Cells were then treated with secondary antibodies conjugated to Alexa-594 (Cat. No. A21203, Invitrogen, Thermo Fisher Scientific) and Alexa-488 (Cat. No. A11008, Invitrogen, Thermo Fisher Scientific). The stained coverglass was mounted with ProLong Glass Antifade mountant (Cat. No. P36984, Invitrogen, Thermo Fisher Scientific). Image acquisition was performed on an Elyra 7 Lattice SIM system (Zeiss) fitted with two pco.edge CLHS sCMOS cameras, running Zen software (Black Edition, Zeiss). All acquisition was done on a Plan-Apochromat 63×/1.4NA oil immersion objective (Zeiss). Alignment between the cameras and for chromatic aberration was done with an in-built patterned grating and 200-nm 4-color beads respectively immediately prior to acquiring the sample. All acquisition was done with a lattice grating period of 819 nm, 13 phases, a z-step size of 90 nm, quad 405/488/561/642 beam splitter filter between the objective and grating, and a 560-nm LP filter before the cameras splitting the signal between them. Acquisition was done in stack mode, with two tracks, one for 488 nm and one for 561 nm, and a camera exposure time set at 50 ms for all channels. The first track was using 4% power 561-nm laser, with a BP570-620+LP655 emission filter, and detected on camera 1. The second track was using 2%–4% power 488-nm laser (depending on the primary antibody used), with a BP420-480+BP495-550 emission filter and detected on camera 2.

After acquisition, images were processed in Zeiss’s proprietary SIM processing module, using the SIM2 feature, with adjusted parameters set to the “Weak Fixed” preset (Low input SNR, 15 iterations, 0.065 regularization weight) with 2× Proc Sampling and 4× Out Sampling. After SIM processing, channel alignment was performed using the shift matrix obtained from the SIM2 processed 4-color bead images acquired immediately prior to sample acquisition.

Colocalization between the different fluorescent channels was measured using FiJi v1.53c^88^ and the JACoP plugin^89^ to extract the size of the colocalized area between the two fluorescent signals after thresholding. All thresholding parameters were kept identical throughout the analysis that was batch processed and plotted in Excel (Microsoft) or Prizm 10 (GraphPad) for data analysis.

### Wound-Healing Assay

U2OS cells grown in wound healing assay chambers, Culture-Insert 2 Well (Cat. No. 81176, ibidi), with fixed width inserts around 500 µm of cell free gap. U2OS cells were transfected with NT control RNAi or si*SEC24C* and seeded at 0.25×10^5^ cells in each well. After 48 h, cells were washed and incubated in DMEM with or without glucose for 2 h prior to removal of the insert. Cells were further incubated in DMEM with glucose for 16 h and stained with 0.1% crystal violet staining solution for 30 min. The stained cells were imaged with EVOS M5000 (objective 10X, Evos_AMEP4981, Invitrogen Thermo Fisher) to measure remaining width of cell free gap.

### Transwell Cell Migration Assay

U2OS cells were transfected with NT control RNAi or si*SEC24C*. After 48 h, cells were trypsinized and seeded at 0.5×10^5^ cells onto upper chamber of Transwell plate (Cat. No. 3464, Corning Costar) in media with or without 20% FBS and/or glucose. After an 18-h incubation, the Transwell insert was washed twice with PBS and the cells on the upper chamber were gently removed with PBS-moistened cotton swabs. The cells that had migrated to the lower chamber of the membrane were stained with 0.1% crystal violet staining solution for 30 min before being washed and dried for imaging and counting.

### Mouse Model of Breast Cancer Metastasis

Primary tumor growth and spontaneous metastasis were assessed essentially as described previously^90^. Eight-week-old BALB/c (Cat. No. BALB-F, Taconic) female mice were randomized and anesthetized with isoflurane, and 10^6^ 4T1 breast cancer cells resuspended in 200 μL HBSS were injected into the fourth mammary fat pad. Mice were euthanized with CO_2_ 28 days after inoculation, and the primary tumors were resected and weighed. Lungs were fixed by tracheal perfusion with 4% formaldehyde for 15 min, removed en bloc, and then fixed in 4% formaldehyde for an additional 24 h. Paraffin embedded lung sections were prepared, stained with haematoxylin and eosin (H&E) and imaged. The number of metastases per animal was determined by counting metastatic foci in H&E-stained lung sections. For each mouse, the metastasis count was then divided by the mass of the primary tumor to yield the number of metastases per gram of primary tumor. All mice were housed and handled in accordance with approved St. Jude’s Institutional Animal Care and Use Committee protocols (IACUC# 560-100583).

### Cell Surface Protein Purification

The levels of cell surface protein expression in MEFs were compared using the membrane-impermeant biotinylation reagent, EZ-Link™ Sulfo-NHS-SS-Biotin (Cat. No. 21331, Thermo Fisher Scientific), as described previously^61^. Briefly, MEFs cultured in dishes were washed twice with ice-cold PBSCM solution (PBS containing 0.1 mmol/L CaCl2 and 1 mmol/L MgCl2), incubated twice with Sulfo-NHS-SS-biotin (1.5 mg/mL) on ice with very gentle shaking for 20 min, rinsed briefly, incubated with PBSCM containing 100 mmol/L glycine on ice for 20 min, and lysed with 1% SDS–1%/Triton X-100 lysis buffer. SDS concentration was diluted with SDS-free lysis buffer as 0.1% SDS. Biotinylated plasma membrane proteins were recovered from the cell lysates by using streptavidin-agarose beads (Cat. No. 20349, Thermo Fisher Scientific). The beads were washed with lysis buffer three times, high-salt buffer twice, and low salt buffer once. Then, the biotinylated proteins were eluted from the beads with Laemmli Lysis-buffer containing 400 mM DTT (Cat. No. 38733, Millipore Sigma) and processed for tandem mass tag assay or separated by SDS-PAGE for western blotting. Western bot membranes were probed with antibodies directed against the following targets: E-Cadherin (Cat. No. GTX100443, GeneTex), pan-Cadherin (Cat. No. ab51034, abcam), PMCA1 (Cat. No. ab190355, abcam), Aldolase A (Cat. No. 8060, Cell Signaling), SEC24C (Cat. No. 8531, Cell Signaling), p-AMPKβ1/2 (Cat. No. 4181, Cell Signaling), and AMPKβ1/2 (Cat. No. 4150, Cell Signaling).

### Tandem Mass Tag Assay and Data Analysis

From cell surface protein purification procedure, eluted cell surface proteins were digested into peptides for tandem mass tag analysis. After desalting, the peptides are labeled with tandem mass tag reagents (Cat. No. A52045, Thermo Fisher Scientific). The labeled samples were equally mixed and further fractionated by neutral pH reverse phase liquid chromatography. Forty fractions were collected and further analyzed by low pH reverse phase liquid chromatography-tandem mass spectrometry (LC-MS/MS). The collected data was searched against a database to identify peptides. While the peptides were identified by MS/MS, the quantification was achieved by the fragmented reporter ions in the same MS/MS scans. The peptide quantification data were then corrected for mixing errors and summarized to derive protein quantification results.

The summarized protein quantification was scaled by sample averages across the dataset and followed by log2 transformation. The differential expression analysis was performed using log2 folder change between average of three samples from each of the comparison groups, and p-value was calculated correspondingly using a Student’s t-Test. The volcano plot (Fig. 3B) was plotted using Spotfire v7.5.0 software (TIBCO, Palo Alto, CA). Those proteins of significantly differentially down regulated (p-value < 0.05 and folder change > 10%) responding to lack of glucose, in wild type and control samples but not in Sec24C knockdown samples, were marked in red. The larger size of the dot reflects higher level of rescuing in corresponding comparison of Sec24C knockdown samples, while the top twenty proteins were labeled along with positive control protein Cdh1. The pathway analysis for these down regulated proteins (in red) in response to lack of glucose dependent on the presence of Sec24C was performed using Enrichr^91^ (https://maayanlab.cloud/Enrichr) or DAVID (v2023q4, https://davidbioinformatics.nih.gov/). The result of enriched pathway was listed in Table S1, and the top 10 pathways from Biological Process (GO Term) were plotted in Fig 3C.

### Phosphoproteomics

SEC31A proteins samples were prepared from two different set-ups. In the first set up, HEK293T cells were co-transfected with EYFP-SEC31A and empty vector, WT ULK1, or KD ULK1. After 48 h, cells were lysed and YFP immunoprecipitated. In the second set up, FLAG-SEC31A knock-in U2OS cells were incubated in DMEM with or without glucose for 1 h, followed by FLAG immunoprecipitation. Each protein sample was resolved on 4%–12% bis-Tris gels (Cat. No. NP0335BOX, Thermo Fisher Scientific) and excised from a Coomassie-blue stained SDS gel followed by dithiothreitol reduction, iodoacetamide alkylation, and in-gel digestion (12.5 ng/µl trypsin overnight). The resulting tryptic peptides were concentrated and analyzed by C_18_ capillary reverse phase liquid chromatography coupled with tandem mass spectrometry (LC-MS/MS) using a LTQ Orbitrap FUSION (Thermo Fisher Scientific) mass spectrometer operating under optimized high-resolution mode^92, 93^. Acquired MS/MS data were searched against the Uniprot Human database^94^. Proteins and sites modified by phosphorylation were identified by a target/decoy search strategy^95^ in PEAKS Studio 8.5 software (Build 20180507, Bioinformatics Solutions Inc.)^96^ by dynamically assigning a mass addition of +79.9663 Da for modified Serine/Threonine/Tyrosine residues. All matched MS/MS spectra were filtered by mass accuracy and matching scores to reduce protein false discovery rate to less than 1%. Peptide and modified residue assignments were further validated by manually examining raw MS/MS spectra and confirmed based on unambiguous assignment of characteristic site-specific fragment ions.

### Statistical Analyses

Statistical analysis was performed using SigmaPlot 13 (Systat Software) or GraphPad Prizm 10 (GraphPad); significance was assessed using a two-tailed paired Student’s *t*-test or a one- or two-factor ANOVA followed by Holm-Sidak or Tukey’s *post hoc* multiple comparisons test.

## Data Availability

All data supporting the findings of this study are included within the manuscript and its supplementary information files. No additional datasets were generated or analyzed.

## Code Availability

No custom code was used in this study.

## Acknowledgements

We thank all the Kundu Lab. Members for assistance and feedback. We thank A. Mishra, A. High, K. Yu (Center for Proteomics and Metabolomics, St. Jude Children’s Research Hospital) for processing and acquiring data of TMT and phosphoproteomics. We thank D. D’Amore, N. Nedelsky, and C. Guess for editing the manuscript. This work was supported by the National Institutes of Health grant R01MH115058 (to M.K.) and by ALSAC (American Lebanese Syrian Associated Charities).

## Author Information

### Contributions

**Conceptualization – M.K., J.H.J, W.K., M.L.**

Ideas; formulation or evolution of overarching research goals and aims.

**Methodology – J.H.J., W.K., A.C., S.N., S.M.P., M.L., C.-L.C.**

Development or design of methodology; creation of models.

**Formal Analysis – J.H.J., W.K., A.C., J.M., Y.-D.W., M.L., C.-L.C.**

Application of statistical, mathematical, computational, or other formal techniques to analyze or synthesize study data.

**Funding Acquisition – M.K.**

Acquisition of the financial support for the project leading to this publication.

**Investigation – J.H.J., W.K., S.D., U.U., A.C., S.N., S.M.P., M.L., C.-L.C.**

Conducting a research and investigation process, specifically performing the experiments, or data/evidence collection.

**Project Administration – M.K.**

Management and coordination responsibility for the research activity planning and execution.

**Resources – W.K., J.L.S.**

Provision of study materials, reagents, materials, patients, laboratory samples, animals, instrumentation, computing resources, or other analysis tools.

**Supervision – M.K., A.C., M.L., C.-L.C., J.L.S.**

Oversight and leadership responsibility for the research activity planning and execution, including mentorship external to the core team.

**Validation – W.K., U.U.**

Verification, whether as a part of the activity or separate, of the overall replication/reproducibility of results/experiments and other research outputs.

**Visualization – J.H.J, A.C., J.M., D.S., M.L.**

Preparation, creation and/or presentation of the published work, specifically visualization/data presentation.

**Writing – Original Draft – M.K., J.H.J, S.M.P., M.L.**

Preparation, creation and/or presentation of the published work, specifically writing the initial draft (including substantive translation).

**Writing – Review & Editing – All authors**

Preparation, creation and/or presentation of the published work by those from the original research group, specifically critical review, commentary, or revision—including pre- or post-publication stages. All authors revised and reviewed the manuscript and approved the final version.

## Ethics Declaration

The authors declare no competing interests.

