## Supplemental Table S1 for "ER-to-Golgi Trafficking is a Nutrient-Sensitive Checkpoint Linking Glucose Starvation to Cell Surface Remodeling"

### Table S1. Altered Levels of Cell Surface Proteins Following Glucose Starvation and Associated Pathway Enrichment Analyses in Murine Embryonic Fibroblasts

This supplemental table presents lists of cell surface proteins with significantly altered levels in murine embryonic fibroblasts (MEFs) following glucose starvation, as well as the results of pathway enrichment analyses for these proteins. All protein quantification and statistical analyses are based on TMT-labeled samples subjected to NHS-biotinylation and streptavidin pull-down to specifically enrich for cell surface proteins. The analysis compares the response to glucose starvation in wild-type (WT), non-targeting siRNA-treated control (CT), and siSec24c-treated (KD) MEFs. Cell surface proteins showing significant changes in response to glucose starvation (p-value < 0.05 and fold change > 10%) were included in downstream pathway analyses.

#### Tab Descriptions

**Tab a_Surface protein changes:** Contains quantitative and statistical analysis of cell surface protein levels across conditions.

Column Descriptions:

*All values reflect changes in cell surface protein abundance as measured by TMT-based proteomics following NHS-biotinylation and streptavidin pull-* [illegible]

| Column Heading | Description |
|---|---|
| Symbol | Official gene symbol or protein identifier for each cell surface protein. |
| p-value_WT_Glu-/+ | p-value for the change in cell surface protein abundance in WT MEFs (glucose starvation vs. control). |
| log2R_WT_Glu-/+ | Log2 fold change in cell surface protein levels in WT MEFs (glucose starvation vs. control). |
| -logP(WT_Glu-/+) | -log10(p-value) for the change in WT MEFs (higher values = greater significance). |
| p-value_CT_Glu-/+ | p-value for the change in cell surface protein abundance in CT MEFs (glucose starvation vs. control). |
| log2R_CT_Glu-/+ | Log2 fold change in CT MEFs (glucose starvation vs. control). |
| -logP(CT_Glu-/+) | -log10(p-value) for the change in CT MEFs. |
| p-value_KD_Glu-/+ | p-value for the change in cell surface protein abundance in KD MEFs (glucose starvation vs. control). |
| log2R_KD_Glu-/+ | Log2 fold change in KD MEFs (glucose starvation vs. control). |
| -logP(KD_Glu-/+) | -log10(p-value) for the change in KD MEFs. |
| sig_WT | Direction and significance of change in WT MEFs: -1 = significantly down-regulated; 1 = significantly up-regulated; 0 = not significant (cutoffs: p-value < 0.05 and fold change > 10%). |
| sig_CT | Direction and significance of change in CT MEFs: -1 = significantly down-regulated; 1 = significantly up-regulated; 0 = not significant (same cutoffs). |
| sig_KD | Direction and significance of change in KD MEFs: -1 = significantly down-regulated; 1 = significantly up-regulated; 0 = not significant (same cutoffs). |
| Flag_sig | Protein classification: 3 = positive control (Cdh1/E-Cadherin); 2 = significantly down-regulated in WT and CT, but not KD; 1 = significantly up-regulated in WT and CT, but not KD; 0 = all others. |
| log2R_diff | Difference in log2 fold change of cell surface protein levels between KD and the average of WT and CT, reflecting SEC24C dependence or rescue. For visualization in the volcano plot, a minimum value of 0.05 was applied to determine point size. |
| Volcano_label | Top 20 cell surface proteins most significantly down-regulated in WT and CT but not in KD, ranked by “log2R_diff [KD vs. average(WT/CT)]”; used for volcano plot annotation. |

**Tab b_Down-regulated:** Contains pathway enrichment analysis (Enrichr) of 244 cell surface proteins significantly down-regulated by glucose starvation in both WT and CT MEFs.

**Tab c_Up-regulated:** Contains pathway enrichment analysis (Enrichr) of 95 cell surface proteins significantly up-regulated by glucose starvation in both WT and CT MEFs.

**Tab d_Sec24c-sependent down:** Contains pathway enrichment analysis (DAVID, v2023q4) of cell surface proteins whose down-regulation in response to glucose starvation is dependent on Sec24c.

#### Pathway Analysis Tools:

Enrichr (https://maayanlab.cloud/Enrichr/) for tabs b and c

DAVID (https://davidbioinformatics.nih.gov/) for tab d

**Tab a_Surface protein changes:** Contains quantitative and statistical analysis of cell surface protein levels across conditions.

| | WT_Glu-/+ | | | CT_Glu-/+ | | | KD_Glu-/+ | | | | | | | | |
|---|---|---|---|---|---|---|---|---|---|---|---|---|---|---|---|
| **Symbol** | **p-value** | **log2R** | **-logP** | **p-value** | **log2R** | **-logP** | **p-value** | **log2R** | **-logP** | **sig_WT** | **sig_CT** | **sig_KD** | **Flag_sig** | **log2R_diff** | **Volcano_label** |
| Cdh1 | 0.0127 | -0.156 | 1.897 | 0.041 | -0.198 | 1.392 | 0.514 | -0.050 | 0.289 | -1 | -1 | 0 | 3 | 0.127 | Cdh1 |
| Dnah3 | 0.0092 | -0.237 | 2.037 | 0.000 | -0.398 | 3.310 | 0.825 | -0.038 | 0.084 | -1 | -1 | 0 | 2 | 0.280 | Dnah3 |
| Tmbim1 | 0.0003 | -0.384 | 3.501 | 0.036 | -0.384 | 1.447 | 0.481 | -0.177 | 0.318 | -1 | -1 | 0 | 2 | 0.207 | Tmbim1 |
| Amotl2 | 0.0150 | -0.195 | 1.825 | 0.032 | -0.196 | 1.489 | 0.827 | -0.011 | 0.083 | -1 | -1 | 0 | 2 | 0.184 | Amotl2 |
| Gm11127 | 0.0161 | -0.799 | 1.794 | 0.010 | -0.712 | 1.988 | 0.085 | -0.572 | 1.071 | -1 | -1 | 0 | 2 | 0.184 | Gm11127 |
| Mex3b | 0.0084 | -0.268 | 2.075 | 0.004 | -0.274 | 2.430 | 0.141 | -0.099 | 0.850 | -1 | -1 | 0 | 2 | 0.173 | Mex3b |
| Pik3r5 | 0.0309 | -0.338 | 1.510 | 0.048 | -0.375 | 1.321 | 0.139 | -0.205 | 0.857 | -1 | -1 | 0 | 2 | 0.152 | Pik3r5 |
| Fat1 | 0.0053 | -0.264 | 2.276 | 0.009 | -0.258 | 2.040 | 0.190 | -0.110 | 0.721 | -1 | -1 | 0 | 2 | 0.151 | Fat1 |
| Eaf1 | 0.0164 | -0.221 | 1.785 | 0.019 | -0.242 | 1.716 | 0.093 | -0.105 | 1.032 | -1 | -1 | 0 | 2 | 0.127 | Eaf1 |
| Dpcd | 0.0174 | -0.223 | 1.759 | 0.004 | -0.235 | 2.443 | 0.123 | -0.104 | 0.909 | -1 | -1 | 0 | 2 | 0.125 | Dpcd |
| Lama5 | 0.0096 | -0.281 | 2.016 | 0.013 | -0.195 | 1.886 | 0.011 | -0.116 | 1.954 | -1 | -1 | 0 | 2 | 0.122 | Lama5 |
| Tnfaip1 | 0.0025 | -0.222 | 2.608 | 0.009 | -0.226 | 2.029 | 0.006 | -0.111 | 2.216 | -1 | -1 | 0 | 2 | 0.113 | Tnfaip1 |
| Olfr216 | 0.0100 | -0.230 | 2.001 | 0.000 | -0.386 | 3.845 | 0.269 | -0.196 | 0.570 | -1 | -1 | 0 | 2 | 0.112 | Olfr216 |
| Lamc1 | 0.0031 | -0.180 | 2.507 | 0.016 | -0.213 | 1.793 | 0.014 | -0.087 | 1.869 | -1 | -1 | 0 | 2 | 0.109 | Lamc1 |
| Pcdhga12 | 0.0003 | -0.362 | 3.463 | 0.004 | -0.480 | 2.356 | 0.072 | -0.317 | 1.144 | -1 | -1 | 0 | 2 | 0.104 | Pcdhga12 |
| Atp5mc1 | 0.0230 | -0.374 | 1.639 | 0.001 | -0.245 | 2.985 | 0.051 | -0.208 | 1.291 | -1 | -1 | 0 | 2 | 0.102 | Atp5mc1 |
| Prnp | 0.0345 | -0.250 | 1.462 | 0.002 | -0.310 | 2.794 | 0.140 | -0.180 | 0.854 | -1 | -1 | 0 | 2 | 0.100 | Prnp |
| Agrn | 0.0238 | -0.218 | 1.624 | 0.048 | -0.262 | 1.321 | 0.206 | -0.143 | 0.687 | -1 | -1 | 0 | 2 | 0.097 | Agrn |
| Gm45717 | 0.0150 | -0.254 | 1.825 | 0.002 | -0.230 | 2.615 | 0.176 | -0.146 | 0.754 | -1 | -1 | 0 | 2 | 0.096 | Gm45717 |
| Ifitm3 | 0.0168 | -0.255 | 1.774 | 0.002 | -0.221 | 2.786 | 0.173 | -0.142 | 0.761 | -1 | -1 | 0 | 2 | 0.096 | Ifitm3 |
| Scd4 | 0.0045 | -0.267 | 2.342 | 0.028 | -0.229 | 1.555 | 0.210 | -0.156 | 0.677 | -1 | -1 | 0 | 2 | 0.091 | |
| Rims1 | 0.0286 | -0.163 | 1.544 | 0.004 | -0.187 | 2.392 | 0.022 | -0.096 | 1.651 | -1 | -1 | 0 | 2 | 0.079 | |
| Klhl11 | 0.0199 | -0.194 | 1.702 | 0.006 | -0.143 | 2.252 | 0.280 | -0.090 | 0.553 | -1 | -1 | 0 | 2 | 0.079 | |
| Adgre5 | 0.0183 | -0.169 | 1.737 | 0.002 | -0.246 | 2.640 | 0.017 | -0.131 | 1.776 | -1 | -1 | 0 | 2 | 0.076 | |
| Zeb2 | 0.0109 | -0.161 | 1.962 | 0.003 | -0.145 | 2.486 | 0.390 | -0.078 | 0.409 | -1 | -1 | 0 | 2 | 0.074 | |
| Bud23 | 0.0076 | -0.202 | 2.119 | 0.009 | -0.209 | 2.070 | 0.039 | -0.133 | 1.410 | -1 | -1 | 0 | 2 | 0.072 | |
| Rrn3 | 0.0003 | -0.229 | 3.569 | 0.015 | -0.167 | 1.820 | 0.104 | -0.132 | 0.982 | -1 | -1 | 0 | 2 | 0.066 | |
| Arhgap11a | 0.0174 | -0.181 | 1.759 | 0.042 | -0.151 | 1.380 | 0.124 | -0.100 | 0.905 | -1 | -1 | 0 | 2 | 0.066 | |
| Mtbp | 0.0095 | -0.189 | 2.020 | 0.029 | -0.141 | 1.533 | 0.024 | -0.101 | 1.620 | -1 | -1 | 0 | 2 | 0.064 | |
| Shisa4 | 0.0284 | -0.200 | 1.546 | 0.002 | -0.186 | 2.822 | 0.514 | -0.133 | 0.289 | -1 | -1 | 0 | 2 | 0.060 | |
| Ifitm2 | 0.0138 | -0.238 | 1.861 | 0.010 | -0.242 | 2.005 | 0.085 | -0.182 | 1.070 | -1 | -1 | 0 | 2 | 0.058 | |
| Angptl1 | 0.0383 | -0.202 | 1.417 | 0.013 | -0.140 | 1.893 | 0.088 | -0.114 | 1.056 | -1 | -1 | 0 | 2 | 0.057 | |
| Jag1 | 0.0306 | -0.145 | 1.514 | 0.002 | -0.456 | 2.620 | 0.053 | -0.390 | 1.276 | -1 | -1 | 0 | 2 | 0.050 | |
| Vasn | 0.0217 | -0.306 | 1.664 | 0.012 | -0.282 | 1.921 | 0.101 | -0.270 | 0.995 | -1 | -1 | 0 | 2 | 0.050 | |
| Efnb1 | 0.0286 | -0.180 | 1.544 | 0.006 | -0.277 | 2.223 | 0.061 | -0.263 | 1.213 | -1 | -1 | 0 | 2 | 0.050 | |
| Ptk7 | 0.0202 | -0.178 | 1.696 | 0.000 | -0.298 | 3.374 | 0.066 | -0.257 | 1.179 | -1 | -1 | 0 | 2 | 0.050 | |
| Gpc6 | 0.0359 | -0.171 | 1.444 | 0.003 | -0.276 | 2.560 | 0.066 | -0.250 | 1.181 | -1 | -1 | 0 | 2 | 0.050 | |
| Adgrg6 | 0.0081 | -0.207 | 2.092 | 0.004 | -0.275 | 2.436 | 0.068 | -0.249 | 1.170 | -1 | -1 | 0 | 2 | 0.050 | |
| Klf13 | 0.0090 | -0.230 | 2.048 | 0.012 | -0.250 | 1.904 | 0.065 | -0.242 | 1.188 | -1 | -1 | 0 | 2 | 0.050 | |
| Efnb2 | 0.0282 | -0.170 | 1.549 | 0.003 | -0.247 | 2.472 | 0.160 | -0.233 | 0.796 | -1 | -1 | 0 | 2 | 0.050 | |
| Ptprm | 0.0090 | -0.144 | 2.047 | 0.001 | -0.266 | 3.006 | 0.063 | -0.231 | 1.204 | -1 | -1 | 0 | 2 | 0.050 | |
| Ptprf | 0.0232 | -0.159 | 1.634 | 0.000 | -0.294 | 3.546 | 0.080 | -0.222 | 1.096 | -1 | -1 | 0 | 2 | 0.050 | |
| Robo1 | 0.0061 | -0.161 | 2.213 | 0.001 | -0.222 | 2.824 | 0.085 | -0.208 | 1.069 | -1 | -1 | 0 | 2 | 0.050 | |
| Nectin3 | 0.0199 | -0.159 | 1.702 | 0.001 | -0.208 | 3.030 | 0.114 | -0.208 | 0.942 | -1 | -1 | 0 | 2 | 0.050 | |
| Bcam | 0.0078 | -0.153 | 2.106 | 0.005 | -0.294 | 2.325 | 0.085 | -0.207 | 1.071 | -1 | -1 | 0 | 2 | 0.050 | |
| Cdh3 | 0.0040 | -0.183 | 2.401 | 0.002 | -0.259 | 2.710 | 0.114 | -0.204 | 0.941 | -1 | -1 | 0 | 2 | 0.050 | |
| Slc39a10 | 0.0332 | -0.138 | 1.479 | 0.014 | -0.251 | 1.860 | 0.098 | -0.199 | 1.008 | -1 | -1 | 0 | 2 | 0.050 | |
| Cxadr | 0.0456 | -0.222 | 1.341 | 0.007 | -0.239 | 2.168 | 0.103 | -0.197 | 0.987 | -1 | -1 | 0 | 2 | 0.050 | |
| Tspan14 | 0.0325 | -0.240 | 1.488 | 0.009 | -0.198 | 2.047 | 0.084 | -0.196 | 1.077 | -1 | -1 | 0 | 2 | 0.050 | |
| Ephb4 | 0.0413 | -0.152 | 1.384 | 0.005 | -0.211 | 2.284 | 0.081 | -0.194 | 1.091 | -1 | -1 | 0 | 2 | 0.050 | |
| Anapc10 | 0.0081 | -0.208 | 2.089 | 0.043 | -0.216 | 1.362 | 0.064 | -0.185 | 1.194 | -1 | -1 | 0 | 2 | 0.050 | |
| Ephb3 | 0.0225 | -0.148 | 1.648 | 0.003 | -0.226 | 2.550 | 0.058 | -0.183 | 1.235 | -1 | -1 | 0 | 2 | 0.050 | |
| Lrrc42 | 0.0401 | -0.150 | 1.397 | 0.005 | -0.175 | 2.307 | 0.052 | -0.172 | 1.280 | -1 | -1 | 0 | 2 | 0.050 | |
| Ifnar2 | 0.0142 | -0.182 | 1.848 | 0.034 | -0.192 | 1.466 | 0.060 | -0.170 | 1.222 | -1 | -1 | 0 | 2 | 0.050 | |
| Ctnnb1 | 0.0192 | -0.144 | 1.716 | 0.002 | -0.202 | 2.636 | 0.180 | -0.165 | 0.744 | -1 | -1 | 0 | 2 | 0.050 | |
| Cbarp | 0.0165 | -0.155 | 1.782 | 0.019 | -0.148 | 1.727 | 0.161 | -0.163 | 0.794 | -1 | -1 | 0 | 2 | 0.050 | |
| Ap5m1 | 0.0236 | -0.149 | 1.627 | 0.009 | -0.152 | 2.070 | 0.175 | -0.160 | 0.757 | -1 | -1 | 0 | 2 | 0.050 | |
| F11r | 0.0151 | -0.155 | 1.820 | 0.013 | -0.212 | 1.882 | 0.074 | -0.155 | 1.132 | -1 | -1 | 0 | 2 | 0.050 | |
| Gm21970 | 0.0348 | -0.148 | 1.458 | 0.007 | -0.220 | 2.177 | 0.201 | -0.151 | 0.697 | -1 | -1 | 0 | 2 | 0.050 | |
| Nectin2 | 0.0470 | -0.188 | 1.328 | 0.047 | -0.152 | 1.324 | 0.060 | -0.149 | 1.223 | -1 | -1 | 0 | 2 | 0.050 | |
| Akr7a2 | 0.0232 | -0.177 | 1.634 | 0.045 | -0.169 | 1.350 | 0.105 | -0.134 | 0.981 | -1 | -1 | 0 | 2 | 0.050 | |
| Smtn | 0.0139 | -0.150 | 1.857 | 0.028 | -0.150 | 1.547 | 0.023 | -0.125 | 1.639 | -1 | -1 | 0 | 2 | 0.050 | |
| Ube2j1 | 0.0298 | -0.145 | 1.526 | 0.047 | -0.141 | 1.326 | 0.025 | -0.102 | 1.603 | -1 | -1 | 0 | 2 | 0.050 | |
| Zfyve21 | 0.0338 | 0.176 | 1.472 | 0.011 | 0.228 | 1.958 | 0.345 | 0.040 | 0.463 | 1 | 1 | 0 | 1 | 0.050 | |
| Rpf2 | 0.0071 | 0.203 | 2.147 | 0.038 | 0.199 | 1.425 | 0.144 | 0.046 | 0.842 | 1 | 1 | 0 | 1 | 0.050 | |
| Ctcf | 0.0132 | 0.154 | 1.879 | 0.019 | 0.186 | 1.727 | 0.189 | 0.058 | 0.723 | 1 | 1 | 0 | 1 | 0.050 | |

| | | | | | | | | | | | | | | |
|---|---|---|---|---|---|---|---|---|---|---|---|---|---|---|
| Sptbn5 | 0.0003 | 0.210 | 3.558 | 0.031 | 0.159 | 1.510 | 0.175 | 0.081 | 0.756 | 1 | 1 | 0 | 1 | 0.050 |
| Plce1 | 0.0312 | 0.144 | 1.505 | 0.046 | 0.211 | 1.333 | 0.310 | 0.083 | 0.509 | 1 | 1 | 0 | 1 | 0.050 |
| Rab10 | 0.0007 | 0.595 | 3.164 | 0.022 | 0.162 | 1.667 | 0.195 | 0.086 | 0.710 | 1 | 1 | 0 | 1 | 0.050 |
| Mrps34 | 0.0013 | 0.141 | 2.900 | 0.043 | 0.219 | 1.371 | 0.123 | 0.090 | 0.910 | 1 | 1 | 0 | 1 | 0.050 |
| Plekho2 | 0.0166 | 0.173 | 1.780 | 0.013 | 0.174 | 1.874 | 0.358 | 0.093 | 0.446 | 1 | 1 | 0 | 1 | 0.050 |
| Pgk2 | 0.0090 | 0.239 | 2.047 | 0.049 | 0.194 | 1.311 | 0.261 | 0.098 | 0.583 | 1 | 1 | 0 | 1 | 0.050 |
| Rab35 | 0.0043 | 0.397 | 2.371 | 0.008 | 0.157 | 2.077 | 0.309 | 0.100 | 0.510 | 1 | 1 | 0 | 1 | 0.050 |
| Rab15 | 0.0015 | 0.154 | 2.833 | 0.049 | 0.203 | 1.308 | 0.155 | 0.101 | 0.808 | 1 | 1 | 0 | 1 | 0.050 |
| Lias | 0.0085 | 0.317 | 2.072 | 0.001 | 0.249 | 2.972 | 0.392 | 0.108 | 0.407 | 1 | 1 | 0 | 1 | 0.050 |
| Rhoa | 0.0009 | 0.253 | 3.045 | 0.033 | 0.255 | 1.479 | 0.239 | 0.113 | 0.621 | 1 | 1 | 0 | 1 | 0.050 |
| Pip4k2b | 0.0030 | 0.199 | 2.528 | 0.003 | 0.232 | 2.567 | 0.017 | 0.118 | 1.774 | 1 | 1 | 0 | 1 | 0.050 |
| Pgk1 | 0.0064 | 0.273 | 2.194 | 0.024 | 0.222 | 1.626 | 0.197 | 0.122 | 0.706 | 1 | 1 | 0 | 1 | 0.050 |
| Cbx5 | 0.0205 | 0.198 | 1.687 | 0.044 | 0.210 | 1.352 | 0.006 | 0.125 | 2.259 | 1 | 1 | 0 | 1 | 0.050 |
| Rap2b | 0.0145 | 0.170 | 1.840 | 0.001 | 0.255 | 2.916 | 0.002 | 0.128 | 2.658 | 1 | 1 | 0 | 1 | 0.050 |
| Wdr77 | 0.0108 | 0.188 | 1.967 | 0.008 | 0.152 | 2.078 | 0.122 | 0.130 | 0.912 | 1 | 1 | 0 | 1 | 0.050 |
| Nifk | 0.0303 | 0.172 | 1.518 | 0.006 | 0.170 | 2.197 | 0.003 | 0.131 | 2.505 | 1 | 1 | 0 | 1 | 0.050 |
| Rab2b | 0.0070 | 0.160 | 2.155 | 0.032 | 0.158 | 1.501 | 0.065 | 0.133 | 1.185 | 1 | 1 | 0 | 1 | 0.050 |
| Rpap3 | 0.0042 | 0.151 | 2.375 | 0.007 | 0.146 | 2.184 | 0.004 | 0.137 | 2.371 | 1 | 1 | 0 | 1 | 0.050 |
| Prmt5 | 0.0004 | 0.191 | 3.375 | 0.003 | 0.157 | 2.535 | 0.106 | 0.146 | 0.974 | 1 | 1 | 0 | 1 | 0.050 |
| Rab2a | 0.0047 | 0.169 | 2.330 | 0.037 | 0.197 | 1.428 | 0.078 | 0.150 | 1.106 | 1 | 1 | 0 | 1 | 0.050 |
| Scrib | 0.0004 | 0.202 | 3.372 | 0.041 | 0.147 | 1.384 | 0.094 | 0.170 | 1.028 | 1 | 1 | 0 | 1 | 0.050 |
| Gldc | 0.0415 | 0.216 | 1.382 | 0.004 | 0.193 | 2.426 | 0.095 | 0.187 | 1.021 | 1 | 1 | 0 | 1 | 0.050 |
| Mrps28 | 0.0079 | 0.195 | 2.101 | 0.003 | 0.182 | 2.565 | 0.097 | 0.231 | 1.013 | 1 | 1 | 0 | 1 | 0.050 |
| Rab11a | 0.0088 | 0.285 | 2.056 | 0.035 | 0.213 | 1.452 | 0.060 | 0.261 | 1.224 | 1 | 1 | 0 | 1 | 0.050 |
| Ncf2 | 0.0056 | 0.244 | 2.249 | 0.000 | 0.293 | 3.642 | 0.299 | 0.465 | 0.524 | 1 | 1 | 0 | 1 | 0.050 |
| Gys1 | 0.2395 | 0.088 | 0.621 | 0.021 | 0.689 | 1.679 | 0.017 | 0.745 | 1.767 | 0 | 1 | 1 | 0 | 0.050 |
| Gys2 | 0.3312 | 0.117 | 0.480 | 0.008 | 0.894 | 2.086 | 0.023 | 0.858 | 1.641 | 0 | 1 | 1 | 0 | 0.050 |
| Agl | 0.0368 | 0.129 | 1.434 | 0.000 | 0.875 | 3.401 | 0.001 | 0.887 | 2.915 | 0 | 1 | 1 | 0 | 0.050 |
| Sumo2 | 0.0001 | 1.265 | 4.226 | 0.127 | 0.724 | 0.897 | 0.020 | 0.852 | 1.701 | 1 | 0 | 1 | 0 | 0.050 |
| Hk1 | 0.0023 | 0.186 | 2.635 | 0.004 | 0.720 | 2.405 | 0.005 | 0.677 | 2.314 | 1 | 1 | 1 | 0 | 0.050 |
| Dnah7b | 0.0002 | 0.532 | 3.707 | 0.001 | 0.648 | 3.160 | 0.003 | 0.677 | 2.562 | 1 | 1 | 1 | 0 | 0.050 |
| Hkdc1 | 0.0013 | 0.211 | 2.871 | 0.006 | 0.709 | 2.240 | 0.005 | 0.713 | 2.290 | 1 | 1 | 1 | 0 | 0.050 |
| Dnah7a | 0.0001 | 0.632 | 4.187 | 0.001 | 0.791 | 3.158 | 0.002 | 0.878 | 2.780 | 1 | 1 | 1 | 0 | 0.050 |
| Gale | 0.0000 | 0.447 | 4.552 | 0.005 | 1.222 | 2.260 | 0.008 | 1.033 | 2.073 | 1 | 1 | 1 | 0 | 0.050 |
| Pklr | 0.0000 | 1.757 | 5.469 | 0.001 | 1.664 | 2.846 | 0.001 | 1.740 | 3.220 | 1 | 1 | 1 | 0 | 0.050 |
| Pkm | 0.0000 | 1.735 | 5.273 | 0.001 | 1.674 | 2.868 | 0.000 | 1.747 | 3.771 | 1 | 1 | 1 | 0 | 0.050 |
| Dhfr | 0.0005 | -0.463 | 3.318 | 0.014 | -0.916 | 1.861 | 0.000 | -1.336 | 5.872 | -1 | -1 | -1 | 0 | 0.050 |
| Ifnar1 | 0.0014 | -0.582 | 2.860 | 0.001 | -1.113 | 3.280 | 0.001 | -0.998 | 3.241 | -1 | -1 | -1 | 0 | 0.050 |
| Bgn | 0.0012 | -0.344 | 2.921 | 0.001 | -0.907 | 2.890 | 0.001 | -0.785 | 2.918 | -1 | -1 | -1 | 0 | 0.050 |
| Nmral1 | 0.0001 | -0.829 | 3.836 | 0.008 | -0.890 | 2.079 | 0.004 | -0.779 | 2.359 | -1 | -1 | -1 | 0 | 0.050 |
| Ghr | 0.0011 | -0.570 | 2.976 | 0.001 | -0.865 | 3.133 | 0.003 | -0.742 | 2.487 | -1 | -1 | -1 | 0 | 0.050 |
| Hmgcs1 | 0.0002 | -0.463 | 3.782 | 0.004 | -0.732 | 2.422 | 0.005 | -0.715 | 2.302 | -1 | -1 | -1 | 0 | 0.050 |
| Tnfrsf1a | 0.0001 | -0.386 | 4.010 | 0.000 | -0.843 | 4.081 | 0.001 | -0.687 | 3.233 | -1 | -1 | -1 | 0 | 0.050 |
| Il6st | 0.0022 | -0.345 | 2.653 | 0.001 | -0.803 | 3.029 | 0.002 | -0.668 | 2.670 | -1 | -1 | -1 | 0 | 0.050 |
| Sdc4 | 0.0013 | -0.534 | 2.873 | 0.000 | -0.650 | 3.460 | 0.000 | -0.667 | 3.528 | -1 | -1 | -1 | 0 | 0.050 |
| Snai1 | 0.0381 | -0.376 | 1.419 | 0.000 | -0.721 | 3.465 | 0.010 | -0.619 | 1.993 | -1 | -1 | -1 | 0 | 0.050 |
| Il4r | 0.0001 | -0.583 | 4.034 | 0.002 | -0.623 | 2.738 | 0.000 | -0.573 | 4.312 | -1 | -1 | -1 | 0 | 0.050 |
| Aprt | 0.0020 | -0.423 | 2.706 | 0.004 | -0.475 | 2.423 | 0.000 | -0.569 | 3.788 | -1 | -1 | -1 | 0 | 0.050 |
| Il1rl1 | 0.0011 | -0.282 | 2.942 | 0.001 | -0.568 | 3.120 | 0.003 | -0.539 | 2.534 | -1 | -1 | -1 | 0 | 0.050 |
| Sppl2a | 0.0013 | -0.296 | 2.893 | 0.000 | -0.673 | 3.707 | 0.003 | -0.537 | 2.571 | -1 | -1 | -1 | 0 | 0.050 |
| Klf16 | 0.0021 | -0.461 | 2.680 | 0.000 | -0.398 | 3.531 | 0.000 | -0.529 | 3.802 | -1 | -1 | -1 | 0 | 0.050 |
| Lgals1 | 0.0025 | -0.232 | 2.602 | 0.001 | -0.460 | 2.879 | 0.006 | -0.529 | 2.237 | -1 | -1 | -1 | 0 | 0.050 |
| Rnf149 | 0.0031 | -0.403 | 2.507 | 0.001 | -0.516 | 3.205 | 0.000 | -0.525 | 4.327 | -1 | -1 | -1 | 0 | 0.050 |
| LOC633332 | 0.0038 | -0.362 | 2.423 | 0.003 | -0.562 | 2.564 | 0.008 | -0.508 | 2.107 | -1 | -1 | -1 | 0 | 0.050 |
| Itm2b | 0.0002 | -0.237 | 3.630 | 0.006 | -0.625 | 2.256 | 0.009 | -0.506 | 2.045 | -1 | -1 | -1 | 0 | 0.050 |
| Crim1 | 0.0021 | -0.385 | 2.684 | 0.000 | -0.531 | 3.327 | 0.000 | -0.502 | 5.107 | -1 | -1 | -1 | 0 | 0.050 |
| D17H6S56E-[illegible] | 0.0087 | -0.195 | 2.060 | 0.012 | -0.676 | 1.915 | 0.049 | -0.501 | 1.312 | -1 | -1 | -1 | 0 | 0.050 |
| Sema3c | 0.0094 | -0.212 | 2.029 | 0.001 | -0.561 | 2.836 | 0.017 | -0.497 | 1.776 | -1 | -1 | -1 | 0 | 0.050 |
| Sema4b | 0.0024 | -0.354 | 2.617 | 0.000 | -0.585 | 3.804 | 0.002 | -0.492 | 2.702 | -1 | -1 | -1 | 0 | 0.050 |
| Chmp1a | 0.0065 | -0.235 | 2.185 | 0.001 | -0.347 | 2.903 | 0.007 | -0.484 | 2.176 | -1 | -1 | -1 | 0 | 0.050 |
| Pcdhb14 | 0.0016 | -0.296 | 2.805 | 0.003 | -0.507 | 2.531 | 0.008 | -0.474 | 2.120 | -1 | -1 | -1 | 0 | 0.050 |
| Pcdhb22 | 0.0050 | -0.258 | 2.303 | 0.012 | -0.498 | 1.938 | 0.014 | -0.472 | 1.851 | -1 | -1 | -1 | 0 | 0.050 |
| Akr1b1 | 0.0003 | -0.599 | 3.583 | 0.046 | -0.254 | 1.335 | 0.016 | -0.468 | 1.807 | -1 | -1 | -1 | 0 | 0.050 |
| Ttyh2 | 0.0014 | -0.388 | 2.860 | 0.000 | -0.459 | 4.278 | 0.000 | -0.467 | 4.225 | -1 | -1 | -1 | 0 | 0.050 |
| Klf10 | 0.0354 | -0.241 | 1.451 | 0.006 | -0.487 | 2.252 | 0.000 | -0.464 | 4.887 | -1 | -1 | -1 | 0 | 0.050 |
| Aplp2 | 0.0449 | -0.138 | 1.348 | 0.000 | -0.575 | 4.000 | 0.014 | -0.456 | 1.860 | -1 | -1 | -1 | 0 | 0.050 |
| Lipg | 0.0030 | -0.312 | 2.530 | 0.002 | -0.470 | 2.818 | 0.028 | -0.454 | 1.552 | -1 | -1 | -1 | 0 | 0.050 |
| Il17ra | 0.0015 | -0.500 | 2.811 | 0.002 | -0.435 | 2.670 | 0.000 | -0.447 | 4.420 | -1 | -1 | -1 | 0 | 0.050 |
| Efna5 | 0.0046 | -0.234 | 2.334 | 0.005 | -0.595 | 2.324 | 0.012 | -0.444 | 1.905 | -1 | -1 | -1 | 0 | 0.050 |
| Pclaf | 0.0001 | -0.303 | 4.001 | 0.006 | -0.397 | 2.187 | 0.000 | -0.441 | 3.456 | -1 | -1 | -1 | 0 | 0.050 |
| Apex2 | 0.0011 | -0.278 | 2.973 | 0.014 | -0.280 | 1.858 | 0.004 | -0.434 | 2.431 | -1 | -1 | -1 | 0 | 0.050 |

| | | | | | | | | | | | | | |
|---|---|---|---|---|---|---|---|---|---|---|---|---|---|
| Ppat | 0.0209 | -0.291 | 1.679 | 0.001 | -0.450 | 2.960 | 0.002 | -0.431 | 2.610 | -1 | -1 | -1 | 0 | 0.050 |
| Ca12 | 0.0231 | -0.315 | 1.636 | 0.011 | -0.457 | 1.946 | 0.000 | -0.423 | 4.175 | -1 | -1 | -1 | 0 | 0.050 |
| Il1rn | 0.0004 | -0.336 | 3.415 | 0.001 | -0.453 | 2.964 | 0.002 | -0.416 | 2.613 | -1 | -1 | -1 | 0 | 0.050 |
| Rdh11 | 0.0011 | -0.204 | 2.966 | 0.002 | -0.321 | 2.772 | 0.000 | -0.407 | 3.913 | -1 | -1 | -1 | 0 | 0.050 |
| Ddr1 | 0.0029 | -0.286 | 2.536 | 0.002 | -0.421 | 2.662 | 0.003 | -0.397 | 2.537 | -1 | -1 | -1 | 0 | 0.050 |
| Hs6st2 | 0.0292 | -0.206 | 1.535 | 0.021 | -0.438 | 1.672 | 0.020 | -0.393 | 1.695 | -1 | -1 | -1 | 0 | 0.050 |
| Pcdhgb6 | 0.0002 | -0.308 | 3.806 | 0.004 | -0.454 | 2.379 | 0.042 | -0.391 | 1.374 | -1 | -1 | -1 | 0 | 0.050 |
| Crlf2 | 0.0271 | -0.295 | 1.566 | 0.001 | -0.365 | 2.862 | 0.009 | -0.391 | 2.050 | -1 | -1 | -1 | 0 | 0.050 |
| Ttyh3 | 0.0050 | -0.216 | 2.297 | 0.001 | -0.425 | 2.849 | 0.022 | -0.390 | 1.667 | -1 | -1 | -1 | 0 | 0.050 |
| Emilin1 | 0.0008 | -0.341 | 3.076 | 0.004 | -0.345 | 2.397 | 0.005 | -0.381 | 2.289 | -1 | -1 | -1 | 0 | 0.050 |
| Slit1 | 0.0259 | -0.230 | 1.586 | 0.003 | -0.460 | 2.547 | 0.005 | -0.381 | 2.325 | -1 | -1 | -1 | 0 | 0.050 |
| Tapbp | 0.0377 | -0.140 | 1.423 | 0.002 | -0.306 | 2.629 | 0.003 | -0.380 | 2.573 | -1 | -1 | -1 | 0 | 0.050 |
| St3gal5 | 0.0046 | -0.157 | 2.342 | 0.001 | -0.432 | 3.208 | 0.012 | -0.379 | 1.917 | -1 | -1 | -1 | 0 | 0.050 |
| Pcdhga1 | 0.0012 | -0.226 | 2.909 | 0.003 | -0.372 | 2.480 | 0.004 | -0.378 | 2.381 | -1 | -1 | -1 | 0 | 0.050 |
| Hbegf | 0.0001 | -0.468 | 3.951 | 0.035 | -0.489 | 1.459 | 0.001 | -0.373 | 3.264 | -1 | -1 | -1 | 0 | 0.050 |
| Ptprs | 0.0069 | -0.179 | 2.161 | 0.000 | -0.426 | 4.030 | 0.013 | -0.371 | 1.874 | -1 | -1 | -1 | 0 | 0.050 |
| Tmem132a | 0.0018 | -0.198 | 2.753 | 0.001 | -0.338 | 3.018 | 0.003 | -0.365 | 2.499 | -1 | -1 | -1 | 0 | 0.050 |
| Pcdhgb7 | 0.0033 | -0.328 | 2.482 | 0.003 | -0.368 | 2.593 | 0.012 | -0.363 | 1.916 | -1 | -1 | -1 | 0 | 0.050 |
| Elovl5 | 0.0003 | -0.315 | 3.464 | 0.004 | -0.265 | 2.441 | 0.029 | -0.360 | 1.543 | -1 | -1 | -1 | 0 | 0.050 |
| Pcdhga5 | 0.0020 | -0.245 | 2.706 | 0.008 | -0.310 | 2.073 | 0.007 | -0.359 | 2.145 | -1 | -1 | -1 | 0 | 0.050 |
| Egfr | 0.0151 | -0.258 | 1.821 | 0.001 | -0.477 | 2.988 | 0.016 | -0.356 | 1.799 | -1 | -1 | -1 | 0 | 0.050 |
| Mphosph9 | 0.0152 | -0.317 | 1.818 | 0.018 | -0.300 | 1.746 | 0.009 | -0.352 | 2.036 | -1 | -1 | -1 | 0 | 0.050 |
| Arl4c | 0.0126 | -0.204 | 1.900 | 0.003 | -0.460 | 2.519 | 0.024 | -0.352 | 1.626 | -1 | -1 | -1 | 0 | 0.050 |
| Lrrcc1 | 0.0195 | -0.437 | 1.710 | 0.000 | -0.355 | 3.942 | 0.000 | -0.350 | 3.662 | -1 | -1 | -1 | 0 | 0.050 |
| Pcdhga6 | 0.0062 | -0.226 | 2.205 | 0.008 | -0.300 | 2.116 | 0.008 | -0.350 | 2.093 | -1 | -1 | -1 | 0 | 0.050 |
| Unc5c | 0.0009 | -0.227 | 3.068 | 0.000 | -0.348 | 3.933 | 0.007 | -0.347 | 2.141 | -1 | -1 | -1 | 0 | 0.050 |
| Dcbld2 | 0.0008 | -0.189 | 3.088 | 0.002 | -0.405 | 2.687 | 0.006 | -0.346 | 2.253 | -1 | -1 | -1 | 0 | 0.050 |
| Unc5b | 0.0178 | -0.185 | 1.749 | 0.012 | -0.328 | 1.919 | 0.009 | -0.343 | 2.054 | -1 | -1 | -1 | 0 | 0.050 |
| Cd63 | 0.0083 | -0.287 | 2.079 | 0.007 | -0.385 | 2.126 | 0.008 | -0.343 | 2.076 | -1 | -1 | -1 | 0 | 0.050 |
| Tgfbr3 | 0.0019 | -0.234 | 2.718 | 0.004 | -0.412 | 2.374 | 0.000 | -0.343 | 3.762 | -1 | -1 | -1 | 0 | 0.050 |
| Senp2 | 0.0006 | -0.213 | 3.187 | 0.024 | -0.322 | 1.621 | 0.006 | -0.340 | 2.254 | -1 | -1 | -1 | 0 | 0.050 |
| Tll1 | 0.0114 | -0.169 | 1.943 | 0.003 | -0.360 | 2.565 | 0.008 | -0.338 | 2.088 | -1 | -1 | -1 | 0 | 0.050 |
| Pcdhgc5 | 0.0020 | -0.346 | 2.691 | 0.003 | -0.319 | 2.594 | 0.008 | -0.336 | 2.117 | -1 | -1 | -1 | 0 | 0.050 |
| Pcdhga10 | 0.0014 | -0.216 | 2.839 | 0.020 | -0.260 | 1.694 | 0.005 | -0.335 | 2.309 | -1 | -1 | -1 | 0 | 0.050 |
| Thsd7a | 0.0066 | -0.143 | 2.182 | 0.000 | -0.410 | 3.708 | 0.027 | -0.334 | 1.574 | -1 | -1 | -1 | 0 | 0.050 |
| Sema4c | 0.0293 | -0.187 | 1.533 | 0.004 | -0.425 | 2.436 | 0.013 | -0.333 | 1.871 | -1 | -1 | -1 | 0 | 0.050 |
| Pcdhgb5 | 0.0027 | -0.298 | 2.563 | 0.003 | -0.328 | 2.487 | 0.010 | -0.333 | 2.001 | -1 | -1 | -1 | 0 | 0.050 |
| Pcdhgb4 | 0.0044 | -0.309 | 2.359 | 0.003 | -0.310 | 2.498 | 0.007 | -0.332 | 2.163 | -1 | -1 | -1 | 0 | 0.050 |
| Pcdhgb8 | 0.0028 | -0.329 | 2.548 | 0.003 | -0.318 | 2.566 | 0.008 | -0.332 | 2.110 | -1 | -1 | -1 | 0 | 0.050 |
| MACIR | 0.0171 | -0.229 | 1.767 | 0.013 | -0.229 | 1.894 | 0.005 | -0.332 | 2.324 | -1 | -1 | -1 | 0 | 0.050 |
| Emilin2 | 0.0090 | -0.215 | 2.045 | 0.000 | -0.338 | 3.355 | 0.001 | -0.331 | 2.935 | -1 | -1 | -1 | 0 | 0.050 |
| Dctn5 | 0.0010 | -0.286 | 2.986 | 0.016 | -0.231 | 1.800 | 0.004 | -0.330 | 2.351 | -1 | -1 | -1 | 0 | 0.050 |
| Mrps12 | 0.0041 | -0.276 | 2.391 | 0.009 | -0.458 | 2.064 | 0.000 | -0.330 | 3.856 | -1 | -1 | -1 | 0 | 0.050 |
| Pcdh18 | 0.0009 | -0.176 | 3.060 | 0.000 | -0.340 | 3.767 | 0.002 | -0.329 | 2.739 | -1 | -1 | -1 | 0 | 0.050 |
| Fgfrl1 | 0.0002 | -0.239 | 3.816 | 0.003 | -0.436 | 2.564 | 0.008 | -0.329 | 2.085 | -1 | -1 | -1 | 0 | 0.050 |
| Zfp523 | 0.0063 | -0.200 | 2.203 | 0.002 | -0.230 | 2.777 | 0.000 | -0.327 | 3.502 | -1 | -1 | -1 | 0 | 0.050 |
| Ephb6 | 0.0151 | -0.181 | 1.821 | 0.000 | -0.307 | 3.315 | 0.022 | -0.326 | 1.654 | -1 | -1 | -1 | 0 | 0.050 |
| Fhl1 | 0.0176 | -0.165 | 1.754 | 0.010 | -0.163 | 1.990 | 0.006 | -0.321 | 2.256 | -1 | -1 | -1 | 0 | 0.050 |
| H60a | 0.0007 | -0.231 | 3.180 | 0.014 | -0.384 | 1.867 | 0.046 | -0.317 | 1.334 | -1 | -1 | -1 | 0 | 0.050 |
| Lrp10 | 0.0239 | -0.217 | 1.621 | 0.035 | -0.302 | 1.461 | 0.037 | -0.316 | 1.429 | -1 | -1 | -1 | 0 | 0.050 |
| Pcdha4 | 0.0022 | -0.329 | 2.661 | 0.003 | -0.303 | 2.478 | 0.009 | -0.316 | 2.023 | -1 | -1 | -1 | 0 | 0.050 |
| Kirrel3 | 0.0262 | -0.169 | 1.582 | 0.001 | -0.275 | 3.065 | 0.010 | -0.316 | 2.010 | -1 | -1 | -1 | 0 | 0.050 |
| Clcf1 | 0.0251 | -0.326 | 1.601 | 0.002 | -0.215 | 2.701 | 0.021 | -0.314 | 1.670 | -1 | -1 | -1 | 0 | 0.050 |
| Pcdhgc4 | 0.0021 | -0.325 | 2.680 | 0.004 | -0.297 | 2.374 | 0.009 | -0.313 | 2.040 | -1 | -1 | -1 | 0 | 0.050 |
| Pcdhga2 | 0.0039 | -0.298 | 2.412 | 0.002 | -0.292 | 2.642 | 0.012 | -0.312 | 1.921 | -1 | -1 | -1 | 0 | 0.050 |
| Ddr2 | 0.0038 | -0.230 | 2.421 | 0.005 | -0.311 | 2.311 | 0.002 | -0.312 | 2.671 | -1 | -1 | -1 | 0 | 0.050 |
| Cdk4 | 0.0168 | -0.147 | 1.774 | 0.006 | -0.263 | 2.192 | 0.005 | -0.310 | 2.269 | -1 | -1 | -1 | 0 | 0.050 |
| Pcdhga4 | 0.0041 | -0.293 | 2.391 | 0.004 | -0.277 | 2.414 | 0.008 | -0.302 | 2.074 | -1 | -1 | -1 | 0 | 0.050 |
| Fzd8 | 0.0262 | -0.215 | 1.582 | 0.018 | -0.229 | 1.734 | 0.002 | -0.299 | 2.811 | -1 | -1 | -1 | 0 | 0.050 |
| H2-T23 | 0.0291 | -0.160 | 1.535 | 0.001 | -0.332 | 2.958 | 0.029 | -0.297 | 1.530 | -1 | -1 | -1 | 0 | 0.050 |
| Pdlim2 | 0.0036 | -0.164 | 2.446 | 0.008 | -0.211 | 2.085 | 0.002 | -0.296 | 2.707 | -1 | -1 | -1 | 0 | 0.050 |
| Tspan4 | 0.0096 | -0.525 | 2.020 | 0.006 | -0.158 | 2.257 | 0.014 | -0.295 | 1.846 | -1 | -1 | -1 | 0 | 0.050 |
| Pcdhgb1 | 0.0045 | -0.177 | 2.344 | 0.016 | -0.360 | 1.805 | 0.027 | -0.294 | 1.562 | -1 | -1 | -1 | 0 | 0.050 |
| Il1rap | 0.0339 | -0.213 | 1.470 | 0.001 | -0.338 | 2.861 | 0.009 | -0.293 | 2.040 | -1 | -1 | -1 | 0 | 0.050 |
| Cachd1 | 0.0109 | -0.145 | 1.963 | 0.003 | -0.353 | 2.599 | 0.028 | -0.293 | 1.553 | -1 | -1 | -1 | 0 | 0.050 |
| Antxr1 | 0.0199 | -0.178 | 1.700 | 0.017 | -0.289 | 1.757 | 0.005 | -0.292 | 2.267 | -1 | -1 | -1 | 0 | 0.050 |
| Pcdhga11 | 0.0111 | -0.195 | 1.956 | 0.024 | -0.281 | 1.625 | 0.013 | -0.290 | 1.902 | -1 | -1 | -1 | 0 | 0.050 |
| Fcgrt | 0.0018 | -0.219 | 2.743 | 0.038 | -0.473 | 1.420 | 0.002 | -0.290 | 2.656 | -1 | -1 | -1 | 0 | 0.050 |
| Fem1aa | 0.0002 | -0.249 | 3.753 | 0.001 | -0.212 | 3.016 | 0.003 | -0.289 | 2.578 | -1 | -1 | -1 | 0 | 0.050 |
| Aifm1 | 0.0372 | -0.191 | 1.430 | 0.021 | -0.241 | 1.671 | 0.003 | -0.289 | 2.514 | -1 | -1 | -1 | 0 | 0.050 |
| Tll2 | 0.0175 | -0.156 | 1.757 | 0.002 | -0.351 | 2.683 | 0.007 | -0.285 | 2.134 | -1 | -1 | -1 | 0 | 0.050 |

| | | | | | | | | | | | | | |
|---|---|---|---|---|---|---|---|---|---|---|---|---|---|
| Pcdhga3 | 0.0119 | -0.168 | 1.925 | 0.001 | -0.248 | 2.919 | 0.025 | -0.285 | 1.599 | -1 | -1 | -1 | 0 | 0.050 |
| Fem1c | 0.0010 | -0.307 | 3.017 | 0.017 | -0.313 | 1.758 | 0.004 | -0.283 | 2.443 | -1 | -1 | -1 | 0 | 0.050 |
| Mpzl1 | 0.0008 | -0.267 | 3.115 | 0.002 | -0.265 | 2.678 | 0.034 | -0.281 | 1.472 | -1 | -1 | -1 | 0 | 0.050 |
| Pdgfra | 0.0182 | -0.209 | 1.741 | 0.007 | -0.314 | 2.166 | 0.040 | -0.279 | 1.399 | -1 | -1 | -1 | 0 | 0.050 |
| Lrig3 | 0.0391 | -0.183 | 1.408 | 0.009 | -0.301 | 2.062 | 0.000 | -0.279 | 4.332 | -1 | -1 | -1 | 0 | 0.050 |
| Ptprd | 0.0280 | -0.148 | 1.553 | 0.000 | -0.321 | 3.470 | 0.034 | -0.278 | 1.465 | -1 | -1 | -1 | 0 | 0.050 |
| Ptprk | 0.0098 | -0.222 | 2.007 | 0.001 | -0.278 | 2.864 | 0.018 | -0.277 | 1.737 | -1 | -1 | -1 | 0 | 0.050 |
| Klhl13 | 0.0049 | -0.193 | 2.314 | 0.005 | -0.300 | 2.301 | 0.011 | -0.276 | 1.960 | -1 | -1 | -1 | 0 | 0.050 |
| L | 0.0041 | -0.440 | 2.385 | 0.000 | -0.332 | 3.708 | 0.024 | -0.275 | 1.614 | -1 | -1 | -1 | 0 | 0.050 |
| Vcam1 | 0.0110 | -0.154 | 1.959 | 0.011 | -0.335 | 1.962 | 0.038 | -0.274 | 1.417 | -1 | -1 | -1 | 0 | 0.050 |
| Nrg1 | 0.0418 | -0.233 | 1.379 | 0.002 | -0.236 | 2.603 | 0.015 | -0.273 | 1.817 | -1 | -1 | -1 | 0 | 0.050 |
| Scd1 | 0.0036 | -0.290 | 2.442 | 0.002 | -0.249 | 2.797 | 0.011 | -0.272 | 1.960 | -1 | -1 | -1 | 0 | 0.050 |
| Klhl9 | 0.0021 | -0.201 | 2.681 | 0.004 | -0.314 | 2.407 | 0.008 | -0.271 | 2.094 | -1 | -1 | -1 | 0 | 0.050 |
| Rrm2 | 0.0043 | -0.232 | 2.370 | 0.021 | -0.175 | 1.668 | 0.001 | -0.270 | 3.127 | -1 | -1 | -1 | 0 | 0.050 |
| Igsf8 | 0.0198 | -0.188 | 1.703 | 0.001 | -0.286 | 3.263 | 0.039 | -0.269 | 1.413 | -1 | -1 | -1 | 0 | 0.050 |
| Fgfr1 | 0.0343 | -0.145 | 1.465 | 0.000 | -0.268 | 3.473 | 0.009 | -0.266 | 2.029 | -1 | -1 | -1 | 0 | 0.050 |
| Spn | 0.0014 | -0.161 | 2.868 | 0.009 | -0.176 | 2.042 | 0.037 | -0.264 | 1.427 | -1 | -1 | -1 | 0 | 0.050 |
| Znf143 | 0.0012 | -0.160 | 2.925 | 0.008 | -0.181 | 2.116 | 0.004 | -0.264 | 2.399 | -1 | -1 | -1 | 0 | 0.050 |
| Pom121 | 0.0057 | -0.251 | 2.246 | 0.014 | -0.232 | 1.861 | 0.020 | -0.263 | 1.695 | -1 | -1 | -1 | 0 | 0.050 |
| Enkd1 | 0.0229 | -0.233 | 1.639 | 0.035 | -0.321 | 1.457 | 0.036 | -0.260 | 1.442 | -1 | -1 | -1 | 0 | 0.050 |
| Lgals8 | 0.0265 | -0.265 | 1.577 | 0.001 | -0.269 | 2.965 | 0.001 | -0.259 | 2.979 | -1 | -1 | -1 | 0 | 0.050 |
| Pcdhb19 | 0.0046 | -0.326 | 2.338 | 0.002 | -0.200 | 2.720 | 0.032 | -0.257 | 1.491 | -1 | -1 | -1 | 0 | 0.050 |
| Lrig1 | 0.0259 | -0.190 | 1.587 | 0.020 | -0.259 | 1.698 | 0.001 | -0.254 | 3.026 | -1 | -1 | -1 | 0 | 0.050 |
| Fgfr3 | 0.0191 | -0.145 | 1.720 | 0.000 | -0.261 | 3.391 | 0.013 | -0.253 | 1.899 | -1 | -1 | -1 | 0 | 0.050 |
| Lrp4 | 0.0414 | -0.157 | 1.383 | 0.001 | -0.188 | 2.871 | 0.023 | -0.253 | 1.647 | -1 | -1 | -1 | 0 | 0.050 |
| Cdon | 0.0074 | -0.150 | 2.129 | 0.001 | -0.331 | 3.199 | 0.015 | -0.251 | 1.837 | -1 | -1 | -1 | 0 | 0.050 |
| Scd2 | 0.0150 | -0.273 | 1.825 | 0.007 | -0.198 | 2.156 | 0.027 | -0.251 | 1.568 | -1 | -1 | -1 | 0 | 0.050 |
| Epha6 | 0.0390 | -0.198 | 1.409 | 0.024 | -0.249 | 1.616 | 0.006 | -0.250 | 2.193 | -1 | -1 | -1 | 0 | 0.050 |
| Lrp6 | 0.0102 | -0.184 | 1.991 | 0.020 | -0.258 | 1.691 | 0.019 | -0.249 | 1.710 | -1 | -1 | -1 | 0 | 0.050 |
| Pacc1 | 0.0363 | -0.182 | 1.440 | 0.009 | -0.303 | 2.037 | 0.026 | -0.247 | 1.583 | -1 | -1 | -1 | 0 | 0.050 |
| Tmsb10 | 0.0341 | -0.199 | 1.467 | 0.038 | -0.252 | 1.418 | 0.003 | -0.247 | 2.568 | -1 | -1 | -1 | 0 | 0.050 |
| Slit2 | 0.0304 | -0.175 | 1.518 | 0.004 | -0.296 | 2.383 | 0.023 | -0.246 | 1.641 | -1 | -1 | -1 | 0 | 0.050 |
| Nid1 | 0.0054 | -0.147 | 2.267 | 0.007 | -0.373 | 2.169 | 0.029 | -0.245 | 1.542 | -1 | -1 | -1 | 0 | 0.050 |
| Polh | 0.0180 | -0.156 | 1.744 | 0.022 | -0.212 | 1.664 | 0.038 | -0.242 | 1.419 | -1 | -1 | -1 | 0 | 0.050 |
| Heg1 | 0.0075 | -0.224 | 2.127 | 0.000 | -0.287 | 3.404 | 0.011 | -0.242 | 1.950 | -1 | -1 | -1 | 0 | 0.050 |
| Polr1d | 0.0235 | -0.163 | 1.628 | 0.020 | -0.202 | 1.690 | 0.002 | -0.241 | 2.736 | -1 | -1 | -1 | 0 | 0.050 |
| Bmpr2 | 0.0404 | -0.176 | 1.393 | 0.010 | -0.270 | 2.022 | 0.028 | -0.240 | 1.556 | -1 | -1 | -1 | 0 | 0.050 |
| Ahr | 0.0013 | -0.197 | 2.887 | 0.032 | -0.182 | 1.493 | 0.001 | -0.238 | 2.886 | -1 | -1 | -1 | 0 | 0.050 |
| Btk | 0.0253 | -0.215 | 1.596 | 0.045 | -0.256 | 1.347 | 0.015 | -0.236 | 1.828 | -1 | -1 | -1 | 0 | 0.050 |
| Acaca | 0.0305 | -0.206 | 1.515 | 0.006 | -0.326 | 2.204 | 0.038 | -0.233 | 1.421 | -1 | -1 | -1 | 0 | 0.050 |
| Ryk | 0.0032 | -0.159 | 2.494 | 0.002 | -0.273 | 2.642 | 0.031 | -0.231 | 1.506 | -1 | -1 | -1 | 0 | 0.050 |
| Mkln1 | 0.0152 | -0.161 | 1.819 | 0.007 | -0.246 | 2.159 | 0.009 | -0.230 | 2.045 | -1 | -1 | -1 | 0 | 0.050 |
| Acacb | 0.0453 | -0.196 | 1.343 | 0.005 | -0.321 | 2.282 | 0.038 | -0.229 | 1.423 | -1 | -1 | -1 | 0 | 0.050 |
| Layn | 0.0080 | -0.249 | 2.097 | 0.016 | -0.229 | 1.785 | 0.004 | -0.224 | 2.376 | -1 | -1 | -1 | 0 | 0.050 |
| Epha3 | 0.0386 | -0.157 | 1.414 | 0.011 | -0.219 | 1.960 | 0.025 | -0.221 | 1.603 | -1 | -1 | -1 | 0 | 0.050 |
| Dcaf7 | 0.0027 | -0.167 | 2.577 | 0.033 | -0.182 | 1.475 | 0.006 | -0.218 | 2.215 | -1 | -1 | -1 | 0 | 0.050 |
| Epha7 | 0.0282 | -0.189 | 1.549 | 0.047 | -0.234 | 1.324 | 0.012 | -0.218 | 1.937 | -1 | -1 | -1 | 0 | 0.050 |
| Nsmce2 | 0.0010 | -0.186 | 2.988 | 0.018 | -0.186 | 1.738 | 0.010 | -0.217 | 1.985 | -1 | -1 | -1 | 0 | 0.050 |
| Cercam | 0.0375 | -0.178 | 1.426 | 0.010 | -0.217 | 2.016 | 0.024 | -0.217 | 1.620 | -1 | -1 | -1 | 0 | 0.050 |
| Tk1 | 0.0065 | -0.197 | 2.184 | 0.005 | -0.196 | 2.270 | 0.004 | -0.216 | 2.449 | -1 | -1 | -1 | 0 | 0.050 |
| Gm38396 | 0.0009 | -0.288 | 3.061 | 0.002 | -0.161 | 2.626 | 0.037 | -0.213 | 1.426 | -1 | -1 | -1 | 0 | 0.050 |
| Pcdh19 | 0.0019 | -0.305 | 2.720 | 0.047 | -0.220 | 1.327 | 0.002 | -0.212 | 2.729 | -1 | -1 | -1 | 0 | 0.050 |
| Notch1 | 0.0068 | -0.267 | 2.169 | 0.006 | -0.258 | 2.230 | 0.020 | -0.208 | 1.707 | -1 | -1 | -1 | 0 | 0.050 |
| Wdr90 | 0.0097 | -0.363 | 2.012 | 0.002 | -0.196 | 2.686 | 0.028 | -0.207 | 1.553 | -1 | -1 | -1 | 0 | 0.050 |
| Angptl2 | 0.0038 | -0.206 | 2.416 | 0.004 | -0.305 | 2.439 | 0.012 | -0.206 | 1.939 | -1 | -1 | -1 | 0 | 0.050 |
| Plxna2 | 0.0468 | -0.160 | 1.330 | 0.005 | -0.210 | 2.265 | 0.047 | -0.205 | 1.328 | -1 | -1 | -1 | 0 | 0.050 |
| Pcdhgc3 | 0.0098 | -0.304 | 2.007 | 0.029 | -0.230 | 1.538 | 0.009 | -0.204 | 2.046 | -1 | -1 | -1 | 0 | 0.050 |
| Adgra2 | 0.0058 | -0.176 | 2.240 | 0.001 | -0.244 | 3.244 | 0.047 | -0.204 | 1.328 | -1 | -1 | -1 | 0 | 0.050 |
| Morf4l2 | 0.0284 | -0.196 | 1.547 | 0.018 | -0.240 | 1.734 | 0.024 | -0.200 | 1.619 | -1 | -1 | -1 | 0 | 0.050 |
| S1pr2 | 0.0021 | -0.155 | 2.688 | 0.010 | -0.204 | 1.982 | 0.043 | -0.198 | 1.371 | -1 | -1 | -1 | 0 | 0.050 |
| Mxra8 | 0.0183 | -0.222 | 1.737 | 0.030 | -0.208 | 1.517 | 0.003 | -0.198 | 2.500 | -1 | -1 | -1 | 0 | 0.050 |
| Kirrel1 | 0.0131 | -0.160 | 1.883 | 0.037 | -0.232 | 1.428 | 0.042 | -0.197 | 1.380 | -1 | -1 | -1 | 0 | 0.050 |
| Tnfrsf1b | 0.0145 | -0.147 | 1.839 | 0.024 | -0.260 | 1.619 | 0.038 | -0.195 | 1.418 | -1 | -1 | -1 | 0 | 0.050 |
| Ephb1 | 0.0325 | -0.142 | 1.489 | 0.010 | -0.208 | 2.007 | 0.041 | -0.190 | 1.382 | -1 | -1 | -1 | 0 | 0.050 |
| Mcl1 | 0.0387 | -0.164 | 1.412 | 0.031 | -0.311 | 1.507 | 0.038 | -0.190 | 1.423 | -1 | -1 | -1 | 0 | 0.050 |
| Gjc1 | 0.0273 | -0.174 | 1.564 | 0.004 | -0.149 | 2.430 | 0.044 | -0.184 | 1.357 | -1 | -1 | -1 | 0 | 0.050 |
| Fzd6 | 0.0127 | -0.264 | 1.896 | 0.003 | -0.223 | 2.499 | 0.041 | -0.181 | 1.391 | -1 | -1 | -1 | 0 | 0.050 |
| Flt4 | 0.0330 | -0.196 | 1.482 | 0.022 | -0.204 | 1.666 | 0.004 | -0.174 | 2.436 | -1 | -1 | -1 | 0 | 0.050 |
| Kctd10 | 0.0020 | -0.297 | 2.694 | 0.001 | -0.254 | 2.958 | 0.002 | -0.171 | 2.679 | -1 | -1 | -1 | 0 | 0.050 |
| Pvr | 0.0235 | -0.175 | 1.630 | 0.042 | -0.160 | 1.375 | 0.043 | -0.167 | 1.370 | -1 | -1 | -1 | 0 | 0.050 |
| Dnajb14 | 0.0028 | -0.183 | 2.552 | 0.009 | -0.165 | 2.043 | 0.001 | -0.159 | 2.869 | -1 | -1 | -1 | 0 | 0.050 |

| | | | | | | | | | | | | | |
|---|---|---|---|---|---|---|---|---|---|---|---|---|---|
| Tab3 | 0.0035 | -0.240 | 2.453 | 0.008 | -0.162 | 2.106 | 0.018 | -0.157 | 1.736 | -1 | -1 | -1 | 0 | 0.050 |
| Tsen34 | 0.0187 | -0.159 | 1.729 | 0.007 | -0.149 | 2.164 | 0.003 | -0.150 | 2.508 | -1 | -1 | -1 | 0 | 0.050 |
| Aen | 0.0321 | -0.312 | 1.493 | 0.006 | -0.227 | 2.195 | 0.019 | -0.144 | 1.729 | -1 | -1 | -1 | 0 | 0.050 |
| Impa2 | 0.0162 | -0.184 | 1.791 | 0.030 | -0.140 | 1.528 | 0.001 | -0.144 | 2.849 | -1 | -1 | -1 | 0 | 0.050 |
| Itih2 | 0.0414 | -0.252 | 1.383 | 0.136 | -0.492 | 0.865 | 0.007 | -0.667 | 2.149 | -1 | 0 | -1 | 0 | 0.050 |
| Ereg | 0.0056 | -0.335 | 2.253 | 0.054 | -0.520 | 1.265 | 0.000 | -0.561 | 3.306 | -1 | 0 | -1 | 0 | 0.050 |
| Has2 | 0.0040 | -0.390 | 2.398 | 0.063 | -0.369 | 1.202 | 0.025 | -0.454 | 1.610 | -1 | 0 | -1 | 0 | 0.050 |
| Arpin | 0.0058 | -0.176 | 2.233 | 0.225 | -0.188 | 0.647 | 0.002 | -0.449 | 2.808 | -1 | 0 | -1 | 0 | 0.050 |
| Pcolce | 0.0016 | -0.157 | 2.791 | 0.058 | -0.395 | 1.234 | 0.002 | -0.429 | 2.787 | -1 | 0 | -1 | 0 | 0.050 |
| Itih4 | 0.0212 | -0.350 | 1.674 | 0.895 | -0.050 | 0.048 | 0.046 | -0.423 | 1.337 | -1 | 0 | -1 | 0 | 0.050 |
| Sft2d2 | 0.0080 | -0.398 | 2.098 | 0.074 | -0.261 | 1.132 | 0.003 | -0.342 | 2.494 | -1 | 0 | -1 | 0 | 0.050 |
| Cntnap2 | 0.0199 | -0.340 | 1.701 | 0.062 | -0.262 | 1.211 | 0.041 | -0.301 | 1.384 | -1 | 0 | -1 | 0 | 0.050 |
| A2m | 0.0003 | -0.296 | 3.585 | 0.549 | -0.132 | 0.260 | 0.028 | -0.300 | 1.556 | -1 | 0 | -1 | 0 | 0.050 |
| Egr1 | 0.0039 | -0.446 | 2.409 | 0.657 | -0.031 | 0.183 | 0.012 | -0.297 | 1.913 | -1 | 0 | -1 | 0 | 0.050 |
| Cep192 | 0.0017 | -0.332 | 2.780 | 0.409 | -0.110 | 0.389 | 0.014 | -0.296 | 1.859 | -1 | 0 | -1 | 0 | 0.050 |
| Cdc20 | 0.0008 | -0.262 | 3.107 | 0.171 | -0.178 | 0.766 | 0.000 | -0.290 | 3.395 | -1 | 0 | -1 | 0 | 0.050 |
| Srp9 | 0.0071 | -0.172 | 2.150 | 0.055 | -0.172 | 1.262 | 0.002 | -0.257 | 2.699 | -1 | 0 | -1 | 0 | 0.050 |
| Sdc1 | 0.0371 | -0.258 | 1.431 | 0.080 | -0.161 | 1.096 | 0.018 | -0.232 | 1.734 | -1 | 0 | -1 | 0 | 0.050 |
| Acvr2a | 0.0455 | -0.295 | 1.342 | 0.307 | -0.105 | 0.513 | 0.000 | -0.231 | 3.607 | -1 | 0 | -1 | 0 | 0.050 |
| Acvr1 | 0.0075 | -0.257 | 2.124 | 0.142 | -0.180 | 0.846 | 0.000 | -0.227 | 3.367 | -1 | 0 | -1 | 0 | 0.050 |
| Nkiras2 | 0.0029 | -0.386 | 2.535 | 0.066 | -0.196 | 1.182 | 0.002 | -0.227 | 2.708 | -1 | 0 | -1 | 0 | 0.050 |
| Ttc12 | 0.0212 | -0.208 | 1.673 | 0.059 | -0.254 | 1.229 | 0.022 | -0.226 | 1.659 | -1 | 0 | -1 | 0 | 0.050 |
| Tmem205 | 0.0270 | -0.216 | 1.569 | 0.357 | -0.112 | 0.448 | 0.020 | -0.226 | 1.689 | -1 | 0 | -1 | 0 | 0.050 |
| Arl14ep | 0.0077 | -0.250 | 2.116 | 0.010 | -0.107 | 2.005 | 0.001 | -0.220 | 3.095 | -1 | 0 | -1 | 0 | 0.050 |
| Rrm2b | 0.0081 | -0.165 | 2.090 | 0.555 | -0.038 | 0.256 | 0.001 | -0.217 | 3.208 | -1 | 0 | -1 | 0 | 0.050 |
| Lrp5 | 0.0068 | -0.199 | 2.168 | 0.086 | -0.205 | 1.066 | 0.011 | -0.211 | 1.950 | -1 | 0 | -1 | 0 | 0.050 |
| Mpc2 | 0.0141 | -0.251 | 1.852 | 0.208 | -0.133 | 0.683 | 0.022 | -0.210 | 1.661 | -1 | 0 | -1 | 0 | 0.050 |
| Znf142 | 0.0019 | -0.202 | 2.710 | 0.149 | -0.127 | 0.826 | 0.011 | -0.209 | 1.963 | -1 | 0 | -1 | 0 | 0.050 |
| Sv2a | 0.0008 | -0.262 | 3.109 | 0.263 | -0.134 | 0.581 | 0.038 | -0.203 | 1.419 | -1 | 0 | -1 | 0 | 0.050 |
| Rad51 | 0.0003 | -0.197 | 3.549 | 0.064 | -0.163 | 1.195 | 0.000 | -0.202 | 3.328 | -1 | 0 | -1 | 0 | 0.050 |
| Alms1 | 0.0015 | -0.370 | 2.812 | 0.662 | -0.034 | 0.179 | 0.012 | -0.197 | 1.925 | -1 | 0 | -1 | 0 | 0.050 |
| Spag9 | 0.0007 | -0.228 | 3.146 | 0.193 | -0.146 | 0.715 | 0.038 | -0.197 | 1.418 | -1 | 0 | -1 | 0 | 0.050 |
| Tyw3 | 0.0053 | -0.142 | 2.274 | 0.340 | -0.053 | 0.468 | 0.016 | -0.194 | 1.783 | -1 | 0 | -1 | 0 | 0.050 |
| Rab40c | 0.0150 | -0.262 | 1.823 | 0.367 | -0.108 | 0.436 | 0.018 | -0.193 | 1.753 | -1 | 0 | -1 | 0 | 0.050 |
| Zkscan1 | 0.0413 | -0.166 | 1.384 | 0.348 | -0.062 | 0.458 | 0.028 | -0.192 | 1.560 | -1 | 0 | -1 | 0 | 0.050 |
| Gpn1 | 0.0458 | -0.147 | 1.339 | 0.189 | -0.070 | 0.724 | 0.000 | -0.192 | 3.722 | -1 | 0 | -1 | 0 | 0.050 |
| Qtrt2 | 0.0016 | -0.184 | 2.801 | 0.078 | -0.200 | 1.106 | 0.033 | -0.191 | 1.476 | -1 | 0 | -1 | 0 | 0.050 |
| Sap30 | 0.0028 | -0.173 | 2.550 | 0.081 | -0.120 | 1.091 | 0.002 | -0.190 | 2.778 | -1 | 0 | -1 | 0 | 0.050 |
| Fgf2 | 0.0400 | -0.191 | 1.398 | 0.126 | -0.119 | 0.900 | 0.009 | -0.190 | 2.031 | -1 | 0 | -1 | 0 | 0.050 |
| Sqle | 0.0045 | -0.169 | 2.351 | 0.093 | -0.145 | 1.030 | 0.022 | -0.189 | 1.663 | -1 | 0 | -1 | 0 | 0.050 |
| Lipa | 0.0032 | -0.246 | 2.498 | 0.564 | -0.035 | 0.249 | 0.005 | -0.187 | 2.296 | -1 | 0 | -1 | 0 | 0.050 |
| Hmces | 0.0396 | -0.178 | 1.402 | 0.181 | -0.091 | 0.743 | 0.001 | -0.185 | 2.846 | -1 | 0 | -1 | 0 | 0.050 |
| Hsdl2 | 0.0327 | -0.140 | 1.485 | 0.271 | -0.104 | 0.567 | 0.050 | -0.182 | 1.303 | -1 | 0 | -1 | 0 | 0.050 |
| Nphs1 | 0.0357 | -0.219 | 1.447 | 0.454 | -0.094 | 0.343 | 0.008 | -0.179 | 2.087 | -1 | 0 | -1 | 0 | 0.050 |
| Hspb1 | 0.0196 | -0.167 | 1.708 | 0.042 | -0.114 | 1.375 | 0.002 | -0.179 | 2.735 | -1 | 0 | -1 | 0 | 0.050 |
| Ddx49 | 0.0050 | -0.151 | 2.301 | 0.055 | -0.144 | 1.259 | 0.013 | -0.177 | 1.881 | -1 | 0 | -1 | 0 | 0.050 |
| Chka | 0.0131 | -0.242 | 1.882 | 0.118 | -0.138 | 0.930 | 0.002 | -0.175 | 2.768 | -1 | 0 | -1 | 0 | 0.050 |
| Kiaa1549 | 0.0200 | -0.270 | 1.699 | 0.115 | -0.169 | 0.940 | 0.007 | -0.174 | 2.132 | -1 | 0 | -1 | 0 | 0.050 |
| Cdh24 | 0.0482 | -0.154 | 1.317 | 0.061 | -0.191 | 1.214 | 0.005 | -0.173 | 2.301 | -1 | 0 | -1 | 0 | 0.050 |
| AW554918 | 0.0207 | -0.252 | 1.684 | 0.044 | -0.119 | 1.360 | 0.001 | -0.172 | 3.246 | -1 | 0 | -1 | 0 | 0.050 |
| Chek2 | 0.0286 | -0.138 | 1.543 | 0.385 | -0.081 | 0.415 | 0.020 | -0.167 | 1.708 | -1 | 0 | -1 | 0 | 0.050 |
| Yy1 | 0.0028 | -0.151 | 2.548 | 0.102 | -0.134 | 0.990 | 0.016 | -0.165 | 1.792 | -1 | 0 | -1 | 0 | 0.050 |
| Prmt9 | 0.0361 | -0.183 | 1.443 | 0.178 | -0.099 | 0.750 | 0.014 | -0.164 | 1.863 | -1 | 0 | -1 | 0 | 0.050 |
| Hjurp | 0.0103 | -0.140 | 1.987 | 0.519 | -0.073 | 0.285 | 0.004 | -0.161 | 2.385 | -1 | 0 | -1 | 0 | 0.050 |
| N4bp2l2 | 0.0061 | -0.190 | 2.216 | 0.083 | -0.244 | 1.083 | 0.001 | -0.161 | 3.103 | -1 | 0 | -1 | 0 | 0.050 |
| Lgals9 | 0.0480 | -0.153 | 1.319 | 0.069 | -0.132 | 1.160 | 0.002 | -0.160 | 2.703 | -1 | 0 | -1 | 0 | 0.050 |
| Plekhf2 | 0.0025 | -0.225 | 2.596 | 0.844 | 0.014 | 0.074 | 0.022 | -0.160 | 1.656 | -1 | 0 | -1 | 0 | 0.050 |
| Aamp | 0.0177 | -0.205 | 1.752 | 0.025 | -0.106 | 1.602 | 0.021 | -0.160 | 1.683 | -1 | 0 | -1 | 0 | 0.050 |
| Fam171a1 | 0.0482 | -0.200 | 1.317 | 0.067 | -0.130 | 1.172 | 0.005 | -0.159 | 2.268 | -1 | 0 | -1 | 0 | 0.050 |
| Nacc1 | 0.0261 | -0.142 | 1.583 | 0.148 | -0.138 | 0.828 | 0.003 | -0.159 | 2.466 | -1 | 0 | -1 | 0 | 0.050 |
| Xk | 0.0142 | -0.243 | 1.849 | 0.347 | -0.078 | 0.460 | 0.050 | -0.159 | 1.305 | -1 | 0 | -1 | 0 | 0.050 |
| Trmt11 | 0.0492 | -0.143 | 1.308 | 0.046 | -0.126 | 1.333 | 0.010 | -0.158 | 2.015 | -1 | 0 | -1 | 0 | 0.050 |
| Rps28 | 0.0041 | -0.193 | 2.389 | 0.476 | -0.069 | 0.323 | 0.040 | -0.158 | 1.399 | -1 | 0 | -1 | 0 | 0.050 |
| Prr14 | 0.0238 | -0.291 | 1.624 | 0.149 | -0.154 | 0.827 | 0.006 | -0.152 | 2.251 | -1 | 0 | -1 | 0 | 0.050 |
| Nop53 | 0.0003 | -0.445 | 3.598 | 0.105 | -0.227 | 0.978 | 0.047 | -0.151 | 1.325 | -1 | 0 | -1 | 0 | 0.050 |
| Prrc2b | 0.0040 | -0.341 | 2.394 | 0.264 | -0.059 | 0.579 | 0.015 | -0.150 | 1.820 | -1 | 0 | -1 | 0 | 0.050 |
| Secisbp2l | 0.0079 | -0.181 | 2.105 | 0.061 | -0.167 | 1.216 | 0.005 | -0.147 | 2.283 | -1 | 0 | -1 | 0 | 0.050 |
| Ldlr | 0.0225 | -0.159 | 1.648 | 0.360 | -0.142 | 0.443 | 0.030 | -0.147 | 1.526 | -1 | 0 | -1 | 0 | 0.050 |
| Tcf25 | 0.0277 | -0.163 | 1.557 | 0.056 | -0.162 | 1.254 | 0.002 | -0.145 | 2.787 | -1 | 0 | -1 | 0 | 0.050 |
| Dcaf15 | 0.0033 | -0.140 | 2.484 | 0.150 | -0.248 | 0.825 | 0.044 | -0.138 | 1.360 | -1 | 0 | -1 | 0 | 0.050 |
| Kif23 | 0.0089 | -0.190 | 2.049 | 0.154 | -0.082 | 0.813 | 0.030 | -0.138 | 1.518 | -1 | 0 | -1 | 0 | 0.050 |

| | | | | | | | | | | | | | | |
|---|---|---|---|---|---|---|---|---|---|---|---|---|---|---|
| Acat1 | 0.0001 | -0.578 | 3.973 | 0.084 | -0.503 | 1.077 | 0.060 | -0.519 | 1.225 | -1 | 0 | 0 | 0 | 0.050 |
| Ptgs2 | 0.0029 | -0.177 | 2.541 | 0.230 | -0.267 | 0.639 | 0.078 | -0.389 | 1.106 | -1 | 0 | 0 | 0 | 0.050 |
| Phlda1 | 0.0004 | -0.418 | 3.398 | 0.209 | -0.188 | 0.681 | 0.064 | -0.319 | 1.192 | -1 | 0 | 0 | 0 | 0.050 |
| Ptgs1 | 0.0309 | -0.201 | 1.510 | 0.444 | -0.176 | 0.352 | 0.150 | -0.305 | 0.824 | -1 | 0 | 0 | 0 | 0.050 |
| Fkbp1a | 0.0071 | -0.154 | 2.151 | 0.144 | 0.077 | 0.842 | 0.132 | -0.303 | 0.881 | -1 | 0 | 0 | 0 | 0.050 |
| Thap4 | 0.0077 | -0.155 | 2.115 | 0.032 | -0.106 | 1.494 | 0.097 | -0.299 | 1.013 | -1 | 0 | 0 | 0 | 0.050 |
| Ccdc162 | 0.0297 | -0.191 | 1.527 | 0.095 | -0.216 | 1.022 | 0.076 | -0.278 | 1.121 | -1 | 0 | 0 | 0 | 0.050 |
| Sc5d | 0.0057 | -0.332 | 2.244 | 0.214 | -0.092 | 0.669 | 0.128 | -0.272 | 0.892 | -1 | 0 | 0 | 0 | 0.050 |
| Epc1 | 0.0072 | -0.290 | 2.145 | 0.208 | -0.528 | 0.682 | 0.507 | -0.247 | 0.295 | -1 | 0 | 0 | 0 | 0.050 |
| Traip | 0.0027 | -0.156 | 2.576 | 0.477 | -0.152 | 0.322 | 0.240 | -0.240 | 0.619 | -1 | 0 | 0 | 0 | 0.050 |
| Dusp5 | 0.0046 | -0.291 | 2.339 | 0.156 | -0.235 | 0.806 | 0.074 | -0.234 | 1.132 | -1 | 0 | 0 | 0 | 0.050 |
| Aldh7a1 | 0.0079 | -0.259 | 2.103 | 0.051 | -0.293 | 1.293 | 0.051 | -0.228 | 1.293 | -1 | 0 | 0 | 0 | 0.050 |
| C2cd3 | 0.0081 | -0.180 | 2.092 | 0.062 | -0.117 | 1.206 | 0.083 | -0.225 | 1.081 | -1 | 0 | 0 | 0 | 0.050 |
| Pcdh1 | 0.0165 | -0.178 | 1.784 | 0.213 | -0.131 | 0.673 | 0.120 | -0.220 | 0.922 | -1 | 0 | 0 | 0 | 0.050 |
| Cfap97 | 0.0013 | -0.382 | 2.896 | 0.281 | -0.151 | 0.551 | 0.160 | -0.207 | 0.795 | -1 | 0 | 0 | 0 | 0.050 |
| Fzr1 | 0.0022 | -0.297 | 2.666 | 0.059 | -0.236 | 1.231 | 0.119 | -0.194 | 0.923 | -1 | 0 | 0 | 0 | 0.050 |
| Sdc3 | 0.0418 | -0.308 | 1.378 | 0.085 | -0.111 | 1.072 | 0.077 | -0.192 | 1.111 | -1 | 0 | 0 | 0 | 0.050 |
| Mtm1 | 0.0011 | -0.242 | 2.960 | 0.559 | 0.076 | 0.252 | 0.121 | -0.186 | 0.917 | -1 | 0 | 0 | 0 | 0.050 |
| Pcyt2 | 0.0213 | -0.171 | 1.671 | 0.940 | -0.012 | 0.027 | 0.145 | -0.185 | 0.837 | -1 | 0 | 0 | 0 | 0.050 |
| Pkhd1l1 | 0.0227 | -0.180 | 1.644 | 0.055 | -0.282 | 1.258 | 0.243 | -0.185 | 0.614 | -1 | 0 | 0 | 0 | 0.050 |
| Stard3nl | 0.0186 | -0.346 | 1.731 | 0.127 | -0.244 | 0.895 | 0.381 | -0.184 | 0.420 | -1 | 0 | 0 | 0 | 0.050 |
| Baiap3 | 0.0236 | -0.229 | 1.627 | 0.804 | -0.044 | 0.095 | 0.187 | -0.181 | 0.729 | -1 | 0 | 0 | 0 | 0.050 |
| Topaz1 | 0.0090 | -0.254 | 2.045 | 0.208 | -0.152 | 0.682 | 0.180 | -0.180 | 0.744 | -1 | 0 | 0 | 0 | 0.050 |
| 3300002I08Ri | 0.0146 | -0.162 | 1.836 | 0.369 | -0.075 | 0.433 | 0.162 | -0.174 | 0.791 | -1 | 0 | 0 | 0 | 0.050 |
| Gm10130 | 0.0146 | -0.162 | 1.836 | 0.369 | -0.075 | 0.433 | 0.162 | -0.174 | 0.791 | -1 | 0 | 0 | 0 | 0.050 |
| Apob | 0.0002 | -1.022 | 3.606 | 0.595 | -0.146 | 0.225 | 0.248 | -0.174 | 0.605 | -1 | 0 | 0 | 0 | 0.050 |
| Gm49337 | 0.0021 | -0.243 | 2.680 | 0.332 | -0.081 | 0.479 | 0.050 | -0.172 | 1.300 | -1 | 0 | 0 | 0 | 0.050 |
| Ankrd13a | 0.0112 | -0.269 | 1.951 | 0.162 | -0.115 | 0.790 | 0.066 | -0.171 | 1.182 | -1 | 0 | 0 | 0 | 0.050 |
| Mis18bp1 | 0.0014 | -0.178 | 2.859 | 0.143 | -0.099 | 0.843 | 0.105 | -0.170 | 0.980 | -1 | 0 | 0 | 0 | 0.050 |
| Selenok | 0.0001 | -0.296 | 4.052 | 0.085 | -0.199 | 1.073 | 0.145 | -0.169 | 0.839 | -1 | 0 | 0 | 0 | 0.050 |
| Dph1 | 0.0421 | -0.143 | 1.376 | 0.888 | -0.030 | 0.051 | 0.150 | -0.168 | 0.823 | -1 | 0 | 0 | 0 | 0.050 |
| Prps2 | 0.0036 | -0.211 | 2.450 | 0.257 | -0.130 | 0.590 | 0.147 | -0.164 | 0.832 | -1 | 0 | 0 | 0 | 0.050 |
| Spon1 | 0.0010 | -0.185 | 3.012 | 0.260 | -0.309 | 0.585 | 0.545 | -0.163 | 0.264 | -1 | 0 | 0 | 0 | 0.050 |
| Stx7 | 0.0128 | -0.205 | 1.894 | 0.056 | -0.152 | 1.251 | 0.114 | -0.160 | 0.943 | -1 | 0 | 0 | 0 | 0.050 |
| Loxl1 | 0.0034 | -0.187 | 2.463 | 0.109 | -0.103 | 0.964 | 0.135 | -0.145 | 0.870 | -1 | 0 | 0 | 0 | 0.050 |
| Slc24a3 | 0.0087 | -0.147 | 2.059 | 0.306 | -0.082 | 0.515 | 0.096 | -0.144 | 1.019 | -1 | 0 | 0 | 0 | 0.050 |
| Kif18a | 0.0030 | -0.178 | 2.528 | 0.714 | -0.034 | 0.146 | 0.064 | -0.140 | 1.194 | -1 | 0 | 0 | 0 | 0.050 |
| Abhd18 | 0.0414 | -0.203 | 1.383 | 0.841 | -0.019 | 0.075 | 0.123 | -0.139 | 0.911 | -1 | 0 | 0 | 0 | 0.050 |
| Dhrsx | 0.0167 | -0.160 | 1.779 | 0.094 | -0.086 | 1.027 | 0.078 | -0.139 | 1.108 | -1 | 0 | 0 | 0 | 0.050 |
| Ip6k1 | 0.0047 | -0.194 | 2.328 | 0.172 | -0.109 | 0.765 | 0.039 | -0.137 | 1.411 | -1 | 0 | 0 | 0 | 0.050 |
| Smad5 | 0.0063 | -0.145 | 2.202 | 0.015 | -0.103 | 1.813 | 0.018 | -0.134 | 1.749 | -1 | 0 | 0 | 0 | 0.050 |
| Slain2 | 0.0171 | -0.147 | 1.767 | 0.324 | -0.128 | 0.490 | 0.046 | -0.131 | 1.339 | -1 | 0 | 0 | 0 | 0.050 |
| Ttc5 | 0.0331 | -0.141 | 1.480 | 0.020 | -0.100 | 1.705 | 0.104 | -0.131 | 0.981 | -1 | 0 | 0 | 0 | 0.050 |
| Sppl2b | 0.0360 | -0.191 | 1.444 | 0.071 | -0.089 | 1.150 | 0.052 | -0.130 | 1.283 | -1 | 0 | 0 | 0 | 0.050 |
| Tnfrsf10b | 0.0323 | -0.200 | 1.490 | 0.055 | -0.146 | 1.259 | 0.007 | -0.130 | 2.167 | -1 | 0 | 0 | 0 | 0.050 |
| Smad1 | 0.0076 | -0.148 | 2.120 | 0.027 | -0.104 | 1.575 | 0.057 | -0.130 | 1.244 | -1 | 0 | 0 | 0 | 0.050 |
| Mgrn1 | 0.0288 | -0.150 | 1.541 | 0.089 | -0.162 | 1.049 | 0.185 | -0.130 | 0.733 | -1 | 0 | 0 | 0 | 0.050 |
| Poldip3 | 0.0212 | -0.215 | 1.674 | 0.058 | -0.133 | 1.238 | 0.026 | -0.130 | 1.590 | -1 | 0 | 0 | 0 | 0.050 |
| Snap47 | 0.0018 | -0.178 | 2.749 | 0.031 | -0.102 | 1.507 | 0.002 | -0.128 | 2.821 | -1 | 0 | 0 | 0 | 0.050 |
| Fbxo42 | 0.0056 | -0.166 | 2.250 | 0.380 | -0.038 | 0.420 | 0.035 | -0.128 | 1.458 | -1 | 0 | 0 | 0 | 0.050 |
| Dtl | 0.0075 | -0.161 | 2.126 | 0.164 | -0.095 | 0.786 | 0.001 | -0.128 | 2.895 | -1 | 0 | 0 | 0 | 0.050 |
| Gnl3l | 0.0068 | -0.221 | 2.167 | 0.094 | -0.162 | 1.025 | 0.075 | -0.127 | 1.125 | -1 | 0 | 0 | 0 | 0.050 |
| Gtse1 | 0.0128 | -0.150 | 1.893 | 0.199 | -0.083 | 0.702 | 0.109 | -0.126 | 0.963 | -1 | 0 | 0 | 0 | 0.050 |
| Rps21 | 0.0104 | -0.183 | 1.985 | 0.180 | -0.092 | 0.744 | 0.069 | -0.124 | 1.164 | -1 | 0 | 0 | 0 | 0.050 |
| Dclk3 | 0.0183 | -0.160 | 1.737 | 0.058 | -0.155 | 1.235 | 0.097 | -0.123 | 1.011 | -1 | 0 | 0 | 0 | 0.050 |
| Adam19 | 0.0292 | -0.178 | 1.534 | 0.309 | -0.068 | 0.510 | 0.024 | -0.122 | 1.623 | -1 | 0 | 0 | 0 | 0.050 |
| Nabp1 | 0.0052 | -0.167 | 2.286 | 0.391 | -0.073 | 0.408 | 0.055 | -0.122 | 1.256 | -1 | 0 | 0 | 0 | 0.050 |
| Six1 | 0.0460 | -0.205 | 1.337 | 0.983 | 0.004 | 0.008 | 0.013 | -0.119 | 1.881 | -1 | 0 | 0 | 0 | 0.050 |
| Dars | 0.0174 | -0.175 | 1.760 | 0.095 | -0.082 | 1.023 | 0.017 | -0.119 | 1.780 | -1 | 0 | 0 | 0 | 0.050 |
| Slc2a3 | 0.0077 | -0.265 | 2.116 | 0.108 | -0.084 | 0.967 | 0.011 | -0.118 | 1.951 | -1 | 0 | 0 | 0 | 0.050 |
| Zfp740 | 0.0221 | -0.148 | 1.656 | 0.059 | -0.176 | 1.228 | 0.087 | -0.117 | 1.061 | -1 | 0 | 0 | 0 | 0.050 |
| Ube3d | 0.0186 | -0.144 | 1.730 | 0.551 | -0.057 | 0.259 | 0.017 | -0.115 | 1.775 | -1 | 0 | 0 | 0 | 0.050 |
| Mvd | 0.0449 | -0.141 | 1.348 | 0.296 | -0.060 | 0.529 | 0.011 | -0.114 | 1.976 | -1 | 0 | 0 | 0 | 0.050 |
| PIK3R2 | 0.0252 | -0.163 | 1.598 | 0.154 | -0.072 | 0.812 | 0.181 | -0.111 | 0.742 | -1 | 0 | 0 | 0 | 0.050 |
| Tent4a | 0.0159 | -0.178 | 1.798 | 0.565 | -0.032 | 0.248 | 0.094 | -0.111 | 1.025 | -1 | 0 | 0 | 0 | 0.050 |
| BC094435 | 0.0174 | -0.227 | 1.761 | 0.484 | -0.121 | 0.315 | 0.176 | -0.110 | 0.754 | -1 | 0 | 0 | 0 | 0.050 |
| AI987944 | 0.0017 | -0.266 | 2.762 | 0.097 | -0.115 | 1.015 | 0.361 | -0.110 | 0.442 | -1 | 0 | 0 | 0 | 0.050 |
| Ccdc113 | 0.0447 | -0.291 | 1.350 | 0.473 | -0.063 | 0.325 | 0.002 | -0.109 | 2.658 | -1 | 0 | 0 | 0 | 0.050 |
| Arhgap32 | 0.0208 | -0.233 | 1.681 | 0.291 | 0.101 | 0.537 | 0.298 | -0.108 | 0.525 | -1 | 0 | 0 | 0 | 0.050 |
| Kcmf1 | 0.0010 | -0.158 | 3.002 | 0.391 | -0.025 | 0.408 | 0.020 | -0.107 | 1.690 | -1 | 0 | 0 | 0 | 0.050 |
| Calcoco1 | 0.0073 | -0.245 | 2.138 | 0.777 | -0.025 | 0.109 | 0.236 | -0.106 | 0.627 | -1 | 0 | 0 | 0 | 0.050 |

| | | | | | | | | | | | | | | |
|---|---|---|---|---|---|---|---|---|---|---|---|---|---|---|
| Ccdc136 | 0.0029 | -0.285 | 2.531 | 0.535 | -0.090 | 0.272 | 0.142 | -0.102 | 0.847 | -1 | 0 | 0 | 0 | 0.050 |
| Rpl12 | 0.0053 | -0.172 | 2.274 | 0.641 | -0.044 | 0.193 | 0.027 | -0.101 | 1.573 | -1 | 0 | 0 | 0 | 0.050 |
| Fam126a | 0.0494 | -0.248 | 1.306 | 0.631 | -0.025 | 0.200 | 0.015 | -0.101 | 1.834 | -1 | 0 | 0 | 0 | 0.050 |
| Tspan9 | 0.0041 | -0.182 | 2.383 | 0.052 | -0.112 | 1.286 | 0.238 | -0.100 | 0.623 | -1 | 0 | 0 | 0 | 0.050 |
| Ccpg1 | 0.0029 | -0.222 | 2.542 | 0.589 | -0.038 | 0.230 | 0.036 | -0.098 | 1.444 | -1 | 0 | 0 | 0 | 0.050 |
| Fam102a | 0.0318 | -0.146 | 1.498 | 0.103 | -0.159 | 0.986 | 0.223 | -0.097 | 0.652 | -1 | 0 | 0 | 0 | 0.050 |
| Prpsap2 | 0.0129 | -0.156 | 1.891 | 0.288 | -0.157 | 0.541 | 0.350 | -0.097 | 0.456 | -1 | 0 | 0 | 0 | 0.050 |
| Smap1 | 0.0077 | -0.139 | 2.113 | 0.787 | -0.012 | 0.104 | 0.085 | -0.097 | 1.069 | -1 | 0 | 0 | 0 | 0.050 |
| Mapk8ip3 | 0.0066 | -0.158 | 2.183 | 0.171 | -0.069 | 0.768 | 0.230 | -0.095 | 0.637 | -1 | 0 | 0 | 0 | 0.050 |
| Mex3a | 0.0339 | -0.224 | 1.470 | 0.219 | 0.032 | 0.659 | 0.158 | -0.095 | 0.801 | -1 | 0 | 0 | 0 | 0.050 |
| Fhl2 | 0.0391 | -0.201 | 1.408 | 0.384 | -0.128 | 0.415 | 0.566 | -0.094 | 0.247 | -1 | 0 | 0 | 0 | 0.050 |
| Srp14 | 0.0182 | -0.145 | 1.739 | 0.705 | 0.029 | 0.152 | 0.070 | -0.091 | 1.155 | -1 | 0 | 0 | 0 | 0.050 |
| Mex3d | 0.0458 | -0.221 | 1.339 | 0.157 | -0.128 | 0.805 | 0.131 | -0.087 | 0.882 | -1 | 0 | 0 | 0 | 0.050 |
| Mfsd2a | 0.0061 | -0.459 | 2.213 | 0.287 | -0.119 | 0.541 | 0.702 | -0.086 | 0.153 | -1 | 0 | 0 | 0 | 0.050 |
| Nav2 | 0.0069 | -0.230 | 2.162 | 0.371 | -0.210 | 0.430 | 0.808 | -0.085 | 0.093 | -1 | 0 | 0 | 0 | 0.050 |
| Prps1 | 0.0140 | -0.187 | 1.854 | 0.517 | -0.075 | 0.287 | 0.365 | -0.083 | 0.438 | -1 | 0 | 0 | 0 | 0.050 |
| Sh3bgrl2 | 0.0018 | -0.162 | 2.755 | 0.071 | -0.171 | 1.150 | 0.259 | -0.083 | 0.587 | -1 | 0 | 0 | 0 | 0.050 |
| Mypop | 0.0177 | -0.504 | 1.752 | 0.740 | 0.097 | 0.131 | 0.482 | -0.083 | 0.317 | -1 | 0 | 0 | 0 | 0.050 |
| Tgm2 | 0.0009 | -0.271 | 3.029 | 0.110 | -0.168 | 0.959 | 0.037 | -0.082 | 1.435 | -1 | 0 | 0 | 0 | 0.050 |
| Cdk5rap2 | 0.0297 | -0.164 | 1.528 | 0.983 | 0.002 | 0.008 | 0.034 | -0.081 | 1.464 | -1 | 0 | 0 | 0 | 0.050 |
| Pear1 | 0.0244 | -0.212 | 1.613 | 0.390 | -0.058 | 0.409 | 0.144 | -0.077 | 0.841 | -1 | 0 | 0 | 0 | 0.050 |
| Pik3r1 | 0.0037 | -0.244 | 2.432 | 0.111 | -0.083 | 0.954 | 0.321 | -0.076 | 0.493 | -1 | 0 | 0 | 0 | 0.050 |
| Tspan11 | 0.0091 | -0.280 | 2.042 | 0.839 | -0.017 | 0.076 | 0.285 | -0.075 | 0.544 | -1 | 0 | 0 | 0 | 0.050 |
| Rtcb | 0.0296 | -0.202 | 1.529 | 0.318 | -0.096 | 0.497 | 0.353 | -0.072 | 0.453 | -1 | 0 | 0 | 0 | 0.050 |
| Troap | 0.0025 | -0.139 | 2.596 | 0.112 | -0.096 | 0.952 | 0.284 | -0.071 | 0.547 | -1 | 0 | 0 | 0 | 0.050 |
| Etnk1 | 0.0019 | -0.236 | 2.720 | 0.331 | 0.038 | 0.480 | 0.030 | -0.070 | 1.516 | -1 | 0 | 0 | 0 | 0.050 |
| Gpat3 | 0.0038 | -0.150 | 2.416 | 0.837 | -0.020 | 0.077 | 0.709 | -0.067 | 0.150 | -1 | 0 | 0 | 0 | 0.050 |
| Sqstm1 | 0.0278 | -0.303 | 1.556 | 0.628 | -0.019 | 0.202 | 0.240 | -0.067 | 0.620 | -1 | 0 | 0 | 0 | 0.050 |
| Nbr1 | 0.0298 | -0.146 | 1.526 | 0.600 | -0.025 | 0.222 | 0.282 | -0.067 | 0.550 | -1 | 0 | 0 | 0 | 0.050 |
| Rnf168 | 0.0006 | -0.235 | 3.241 | 0.053 | -0.187 | 1.274 | 0.408 | -0.065 | 0.389 | -1 | 0 | 0 | 0 | 0.050 |
| Mllt10 | 0.0037 | -0.232 | 2.427 | 0.330 | -0.101 | 0.482 | 0.265 | -0.064 | 0.577 | -1 | 0 | 0 | 0 | 0.050 |
| Slc1a3 | 0.0328 | -0.188 | 1.484 | 0.082 | -0.113 | 1.086 | 0.375 | -0.063 | 0.426 | -1 | 0 | 0 | 0 | 0.050 |
| C6 | 0.0224 | -0.156 | 1.651 | 0.076 | -0.143 | 1.122 | 0.348 | -0.063 | 0.459 | -1 | 0 | 0 | 0 | 0.050 |
| Ube2n | 0.0044 | -0.146 | 2.360 | 0.116 | -0.092 | 0.934 | 0.144 | -0.062 | 0.843 | -1 | 0 | 0 | 0 | 0.050 |
| Ccser2 | 0.0204 | -0.339 | 1.690 | 0.878 | 0.009 | 0.056 | 0.316 | -0.061 | 0.500 | -1 | 0 | 0 | 0 | 0.050 |
| Notch3 | 0.0362 | -0.273 | 1.442 | 0.331 | -0.105 | 0.480 | 0.319 | -0.061 | 0.496 | -1 | 0 | 0 | 0 | 0.050 |
| Pgrmc2 | 0.0089 | -0.149 | 2.052 | 0.511 | -0.045 | 0.292 | 0.370 | -0.058 | 0.431 | -1 | 0 | 0 | 0 | 0.050 |
| Tmem185b | 0.0456 | -0.216 | 1.341 | 0.108 | -0.074 | 0.968 | 0.426 | -0.058 | 0.371 | -1 | 0 | 0 | 0 | 0.050 |
| Tgfbr1 | 0.0110 | -0.217 | 1.959 | 0.256 | -0.065 | 0.591 | 0.155 | -0.054 | 0.810 | -1 | 0 | 0 | 0 | 0.050 |
| Mvb12a | 0.0414 | -0.237 | 1.383 | 0.987 | -0.001 | 0.006 | 0.314 | -0.054 | 0.503 | -1 | 0 | 0 | 0 | 0.050 |
| Tcf12 | 0.0161 | -0.310 | 1.794 | 0.227 | -0.121 | 0.643 | 0.374 | -0.053 | 0.427 | -1 | 0 | 0 | 0 | 0.050 |
| Zc3h12c | 0.0306 | -0.214 | 1.514 | 0.556 | 0.044 | 0.255 | 0.708 | -0.050 | 0.150 | -1 | 0 | 0 | 0 | 0.050 |
| Blm | 0.0097 | -0.311 | 2.012 | 0.953 | 0.002 | 0.021 | 0.115 | -0.050 | 0.940 | -1 | 0 | 0 | 0 | 0.050 |
| Ndufb11 | 0.0346 | -0.304 | 1.461 | 0.053 | -0.182 | 1.279 | 0.470 | -0.049 | 0.328 | -1 | 0 | 0 | 0 | 0.050 |
| Nek6 | 0.0210 | -0.162 | 1.677 | 0.018 | -0.108 | 1.752 | 0.035 | -0.048 | 1.459 | -1 | 0 | 0 | 0 | 0.050 |
| Actr1a | 0.0293 | -0.151 | 1.534 | 0.539 | -0.042 | 0.268 | 0.181 | -0.047 | 0.743 | -1 | 0 | 0 | 0 | 0.050 |
| Mvp | 0.0374 | -0.196 | 1.427 | 0.497 | -0.049 | 0.304 | 0.290 | -0.046 | 0.538 | -1 | 0 | 0 | 0 | 0.050 |
| Slc43a2 | 0.0394 | -0.231 | 1.405 | 0.572 | -0.064 | 0.243 | 0.335 | -0.046 | 0.475 | -1 | 0 | 0 | 0 | 0.050 |
| Cytip | 0.0158 | -0.247 | 1.800 | 0.097 | -0.095 | 1.014 | 0.264 | -0.044 | 0.578 | -1 | 0 | 0 | 0 | 0.050 |
| Dbr1 | 0.0447 | -0.200 | 1.350 | 0.249 | -0.153 | 0.604 | 0.079 | -0.044 | 1.104 | -1 | 0 | 0 | 0 | 0.050 |
| Slc7a7 | 0.0393 | -0.272 | 1.405 | 0.443 | -0.187 | 0.354 | 0.883 | -0.043 | 0.054 | -1 | 0 | 0 | 0 | 0.050 |
| Bhlhb9 | 0.0336 | -0.246 | 1.474 | 0.055 | -0.266 | 1.256 | 0.720 | -0.042 | 0.142 | -1 | 0 | 0 | 0 | 0.050 |
| Pcnt | 0.0002 | -0.321 | 3.719 | 0.464 | -0.057 | 0.333 | 0.304 | -0.038 | 0.517 | -1 | 0 | 0 | 0 | 0.050 |
| Unk | 0.0045 | -0.138 | 2.350 | 0.779 | 0.021 | 0.108 | 0.496 | -0.038 | 0.305 | -1 | 0 | 0 | 0 | 0.050 |
| Ammecr1l | 0.0074 | -0.213 | 2.132 | 0.728 | -0.018 | 0.138 | 0.486 | -0.038 | 0.313 | -1 | 0 | 0 | 0 | 0.050 |
| Ell | 0.0224 | -0.165 | 1.649 | 0.281 | 0.068 | 0.551 | 0.582 | -0.036 | 0.235 | -1 | 0 | 0 | 0 | 0.050 |
| Pphln1 | 0.0293 | -0.145 | 1.533 | 0.676 | 0.021 | 0.170 | 0.433 | -0.036 | 0.364 | -1 | 0 | 0 | 0 | 0.050 |
| Mfsd8 | 0.0106 | -0.164 | 1.973 | 0.465 | 0.079 | 0.332 | 0.747 | -0.035 | 0.127 | -1 | 0 | 0 | 0 | 0.050 |
| Kif20b | 0.0233 | -0.206 | 1.633 | 0.151 | -0.091 | 0.820 | 0.473 | -0.035 | 0.325 | -1 | 0 | 0 | 0 | 0.050 |
| Disp1 | 0.0409 | -0.162 | 1.388 | 0.911 | 0.008 | 0.041 | 0.534 | -0.034 | 0.272 | -1 | 0 | 0 | 0 | 0.050 |
| Setd2 | 0.0282 | -0.154 | 1.550 | 0.703 | 0.012 | 0.153 | 0.197 | -0.033 | 0.705 | -1 | 0 | 0 | 0 | 0.050 |
| Lpar6 | 0.0169 | -0.234 | 1.771 | 0.809 | -0.017 | 0.092 | 0.726 | -0.033 | 0.139 | -1 | 0 | 0 | 0 | 0.050 |
| Foxk1 | 0.0095 | -0.138 | 2.021 | 0.613 | -0.019 | 0.212 | 0.253 | -0.032 | 0.596 | -1 | 0 | 0 | 0 | 0.050 |
| Pygl | 0.0065 | -0.180 | 2.188 | 0.298 | 0.051 | 0.525 | 0.641 | -0.028 | 0.193 | -1 | 0 | 0 | 0 | 0.050 |
| Tmeff1 | 0.0419 | -0.174 | 1.378 | 0.129 | -0.069 | 0.889 | 0.810 | -0.028 | 0.092 | -1 | 0 | 0 | 0 | 0.050 |
| Nxn | 0.0225 | -0.171 | 1.648 | 0.429 | -0.052 | 0.368 | 0.166 | -0.027 | 0.781 | -1 | 0 | 0 | 0 | 0.050 |
| Get1 | 0.0097 | -0.140 | 2.012 | 0.179 | -0.100 | 0.747 | 0.388 | -0.024 | 0.412 | -1 | 0 | 0 | 0 | 0.050 |
| Alg14 | 0.0186 | -0.166 | 1.730 | 0.687 | 0.033 | 0.163 | 0.764 | -0.022 | 0.117 | -1 | 0 | 0 | 0 | 0.050 |
| Fbln1 | 0.0015 | -0.249 | 2.812 | 0.756 | -0.026 | 0.121 | 0.831 | -0.022 | 0.080 | -1 | 0 | 0 | 0 | 0.050 |
| Cbll1 | 0.0205 | -0.151 | 1.687 | 0.806 | 0.006 | 0.094 | 0.286 | -0.021 | 0.544 | -1 | 0 | 0 | 0 | 0.050 |
| pol | 0.0174 | -0.238 | 1.759 | 0.601 | -0.089 | 0.221 | 0.505 | -0.020 | 0.297 | -1 | 0 | 0 | 0 | 0.050 |

| | | | | | | | | | | | | | |
|---|---|---|---|---|---|---|---|---|---|---|---|---|---|
| Neb | 0.0103 | -0.228 | 1.988 | 0.250 | -0.117 | 0.602 | 0.823 | -0.014 | 0.085 | -1 | 0 | 0 | 0 | 0.050 |
| Rc3h2 | 0.0030 | -0.154 | 2.528 | 0.147 | 0.057 | 0.834 | 0.794 | -0.012 | 0.100 | -1 | 0 | 0 | 0 | 0.050 |
| Synj2bp | 0.0055 | -0.210 | 2.263 | 0.418 | -0.071 | 0.378 | 0.915 | -0.008 | 0.039 | -1 | 0 | 0 | 0 | 0.050 |
| Gm20498 | 0.0007 | -0.222 | 3.132 | 0.745 | -0.028 | 0.128 | 0.912 | -0.008 | 0.040 | -1 | 0 | 0 | 0 | 0.050 |
| Pcyt1a | 0.0421 | -0.227 | 1.375 | 0.463 | -0.086 | 0.334 | 0.842 | -0.007 | 0.075 | -1 | 0 | 0 | 0 | 0.050 |
| Tnrc6a | 0.0136 | -0.201 | 1.867 | 0.482 | 0.029 | 0.317 | 0.949 | -0.001 | 0.023 | -1 | 0 | 0 | 0 | 0.050 |
| Tm7sf3 | 0.0032 | -0.150 | 2.494 | 0.021 | -0.118 | 1.669 | 0.992 | -0.001 | 0.004 | -1 | 0 | 0 | 0 | 0.050 |
| Brip1 | 0.0088 | -0.159 | 2.055 | 0.952 | -0.004 | 0.021 | 0.876 | 0.004 | 0.057 | -1 | 0 | 0 | 0 | 0.050 |
| Snx19 | 0.0382 | -0.141 | 1.417 | 0.575 | 0.047 | 0.240 | 0.954 | 0.004 | 0.021 | -1 | 0 | 0 | 0 | 0.050 |
| Gprc5b | 0.0187 | -0.178 | 1.727 | 0.340 | 0.084 | 0.468 | 0.904 | 0.007 | 0.044 | -1 | 0 | 0 | 0 | 0.050 |
| Ccdc50 | 0.0223 | -0.179 | 1.651 | 0.662 | 0.055 | 0.179 | 0.882 | 0.007 | 0.054 | -1 | 0 | 0 | 0 | 0.050 |
| Pus3 | 0.0211 | -0.146 | 1.675 | 0.853 | 0.012 | 0.069 | 0.890 | 0.009 | 0.050 | -1 | 0 | 0 | 0 | 0.050 |
| Cdv3 | 0.0219 | -0.183 | 1.660 | 0.139 | -0.054 | 0.857 | 0.897 | 0.009 | 0.047 | -1 | 0 | 0 | 0 | 0.050 |
| Kdm3a | 0.0012 | -0.187 | 2.916 | 0.789 | -0.011 | 0.103 | 0.668 | 0.018 | 0.175 | -1 | 0 | 0 | 0 | 0.050 |
| Vrk3 | 0.0190 | -0.139 | 1.721 | 0.917 | -0.021 | 0.038 | 0.653 | 0.018 | 0.185 | -1 | 0 | 0 | 0 | 0.050 |
| Cfap157 | 0.0078 | -0.180 | 2.106 | 0.244 | 0.181 | 0.612 | 0.733 | 0.020 | 0.135 | -1 | 0 | 0 | 0 | 0.050 |
| Stx3 | 0.0226 | -0.227 | 1.646 | 0.875 | -0.013 | 0.058 | 0.875 | 0.023 | 0.058 | -1 | 0 | 0 | 0 | 0.050 |
| Sar1a | 0.0155 | -0.166 | 1.810 | 0.894 | 0.018 | 0.048 | 0.640 | 0.025 | 0.194 | -1 | 0 | 0 | 0 | 0.050 |
| C4a | 0.0205 | -0.165 | 1.688 | 0.482 | 0.093 | 0.317 | 0.799 | 0.026 | 0.097 | -1 | 0 | 0 | 0 | 0.050 |
| Tet3 | 0.0374 | -0.152 | 1.427 | 0.763 | 0.008 | 0.117 | 0.311 | 0.029 | 0.508 | -1 | 0 | 0 | 0 | 0.050 |
| Gpbp1 | 0.0122 | -0.271 | 1.912 | 0.426 | -0.054 | 0.371 | 0.124 | 0.047 | 0.905 | -1 | 0 | 0 | 0 | 0.050 |
| Ranbp9 | 0.0044 | -0.143 | 2.360 | 0.644 | 0.038 | 0.191 | 0.454 | 0.052 | 0.343 | -1 | 0 | 0 | 0 | 0.050 |
| Khdrbs1 | 0.0195 | -0.189 | 1.709 | 0.221 | 0.047 | 0.656 | 0.126 | 0.056 | 0.898 | -1 | 0 | 0 | 0 | 0.050 |
| Tanc2 | 0.0126 | -0.143 | 1.899 | 0.853 | 0.007 | 0.069 | 0.250 | 0.065 | 0.601 | -1 | 0 | 0 | 0 | 0.050 |
| Cacna1c | 0.0100 | -0.142 | 1.999 | 0.052 | 0.108 | 1.286 | 0.468 | 0.073 | 0.330 | -1 | 0 | 0 | 0 | 0.050 |
| Cox5b | 0.0098 | -0.232 | 2.009 | 0.844 | 0.020 | 0.073 | 0.480 | 0.092 | 0.318 | -1 | 0 | 0 | 0 | 0.050 |
| Cspp1 | 0.0460 | -0.265 | 1.337 | 0.837 | -0.028 | 0.077 | 0.003 | 0.111 | 2.547 | -1 | 0 | 0 | 0 | 0.050 |
| Notch2 | 0.0209 | -0.227 | 1.680 | 0.275 | 0.087 | 0.561 | 0.009 | 0.120 | 2.025 | -1 | 0 | 0 | 0 | 0.050 |
| Smap | 0.0193 | -0.142 | 1.714 | 0.535 | 0.050 | 0.272 | 0.065 | 0.144 | 1.186 | -1 | 0 | 0 | 0 | 0.050 |
| Ccnd3 | 0.1223 | -0.335 | 0.913 | 0.024 | -0.725 | 1.627 | 0.024 | -0.679 | 1.611 | 0 | -1 | -1 | 0 | 0.050 |
| Fst | 0.1365 | -0.191 | 0.865 | 0.047 | -0.531 | 1.325 | 0.026 | -0.619 | 1.582 | 0 | -1 | -1 | 0 | 0.050 |
| Gmppa | 0.2967 | -0.052 | 0.528 | 0.002 | -0.604 | 2.643 | 0.015 | -0.618 | 1.813 | 0 | -1 | -1 | 0 | 0.050 |
| Tnc | 0.3946 | -0.043 | 0.404 | 0.001 | -0.729 | 3.148 | 0.007 | -0.596 | 2.182 | 0 | -1 | -1 | 0 | 0.050 |
| App | 0.0890 | -0.123 | 1.050 | 0.001 | -0.634 | 2.835 | 0.016 | -0.497 | 1.793 | 0 | -1 | -1 | 0 | 0.050 |
| Tgfbi | 0.8094 | -0.023 | 0.092 | 0.001 | -0.654 | 2.862 | 0.017 | -0.495 | 1.757 | 0 | -1 | -1 | 0 | 0.050 |
| Thbs1 | 0.0138 | -0.131 | 1.859 | 0.002 | -0.476 | 2.712 | 0.004 | -0.494 | 2.362 | 0 | -1 | -1 | 0 | 0.050 |
| Cpe | 0.1001 | 0.151 | 1.000 | 0.003 | -0.470 | 2.493 | 0.010 | -0.460 | 2.014 | 0 | -1 | -1 | 0 | 0.050 |
| Serinc3 | 0.0575 | -0.278 | 1.240 | 0.001 | -0.401 | 3.255 | 0.012 | -0.448 | 1.934 | 0 | -1 | -1 | 0 | 0.050 |
| Lamp1 | 0.9795 | 0.001 | 0.009 | 0.000 | -0.558 | 4.746 | 0.014 | -0.408 | 1.864 | 0 | -1 | -1 | 0 | 0.050 |
| Erlec1 | 0.0451 | -0.097 | 1.346 | 0.001 | -0.397 | 3.289 | 0.009 | -0.405 | 2.034 | 0 | -1 | -1 | 0 | 0.050 |
| Zfand5 | 0.1270 | -0.104 | 0.896 | 0.038 | -0.390 | 1.419 | 0.024 | -0.405 | 1.612 | 0 | -1 | -1 | 0 | 0.050 |
| Cadm2 | 0.1137 | -0.107 | 0.944 | 0.003 | -0.387 | 2.476 | 0.012 | -0.405 | 1.922 | 0 | -1 | -1 | 0 | 0.050 |
| Gclc | 0.0878 | -0.119 | 1.056 | 0.003 | -0.253 | 2.573 | 0.004 | -0.401 | 2.375 | 0 | -1 | -1 | 0 | 0.050 |
| Sema3d | 0.2663 | -0.071 | 0.575 | 0.001 | -0.439 | 3.113 | 0.035 | -0.394 | 1.452 | 0 | -1 | -1 | 0 | 0.050 |
| Timp1 | 0.4715 | 0.042 | 0.327 | 0.030 | -0.326 | 1.523 | 0.000 | -0.380 | 3.569 | 0 | -1 | -1 | 0 | 0.050 |
| Cntrl | 0.0896 | -0.278 | 1.048 | 0.015 | -0.278 | 1.834 | 0.003 | -0.375 | 2.494 | 0 | -1 | -1 | 0 | 0.050 |
| Pdgfc | 0.0995 | -0.270 | 1.002 | 0.005 | -0.371 | 2.268 | 0.017 | -0.368 | 1.769 | 0 | -1 | -1 | 0 | 0.050 |
| Plat | 0.1564 | -0.083 | 0.806 | 0.000 | -0.360 | 3.441 | 0.007 | -0.366 | 2.176 | 0 | -1 | -1 | 0 | 0.050 |
| Dld | 0.7558 | -0.028 | 0.122 | 0.007 | -0.505 | 2.155 | 0.025 | -0.364 | 1.594 | 0 | -1 | -1 | 0 | 0.050 |
| Ltbp2 | 0.3417 | -0.037 | 0.466 | 0.005 | -0.355 | 2.277 | 0.035 | -0.362 | 1.451 | 0 | -1 | -1 | 0 | 0.050 |
| Cd1d1 | 0.0179 | -0.128 | 1.746 | 0.001 | -0.448 | 3.104 | 0.013 | -0.362 | 1.888 | 0 | -1 | -1 | 0 | 0.050 |
| Serpine1 | 0.3286 | -0.118 | 0.483 | 0.027 | -0.377 | 1.567 | 0.000 | -0.361 | 3.480 | 0 | -1 | -1 | 0 | 0.050 |
| Cbr1 | 0.0137 | -0.086 | 1.863 | 0.003 | -0.229 | 2.528 | 0.007 | -0.353 | 2.179 | 0 | -1 | -1 | 0 | 0.050 |
| Os9 | 0.2682 | -0.083 | 0.572 | 0.002 | -0.346 | 2.745 | 0.003 | -0.353 | 2.491 | 0 | -1 | -1 | 0 | 0.050 |
| Neo1 | 0.0215 | -0.136 | 1.667 | 0.000 | -0.408 | 3.777 | 0.017 | -0.353 | 1.760 | 0 | -1 | -1 | 0 | 0.050 |
| Gm38417 | 0.1300 | -0.164 | 0.886 | 0.007 | -0.417 | 2.186 | 0.014 | -0.351 | 1.847 | 0 | -1 | -1 | 0 | 0.050 |
| Ctsb | 0.2439 | -0.040 | 0.613 | 0.035 | -0.465 | 1.454 | 0.037 | -0.351 | 1.436 | 0 | -1 | -1 | 0 | 0.050 |
| Flt1 | 0.0850 | -0.236 | 1.071 | 0.010 | -0.324 | 2.012 | 0.049 | -0.349 | 1.312 | 0 | -1 | -1 | 0 | 0.050 |
| Itgb2 | 0.2990 | -0.116 | 0.524 | 0.021 | -0.430 | 1.684 | 0.036 | -0.348 | 1.447 | 0 | -1 | -1 | 0 | 0.050 |
| Alpk1 | 0.0885 | -0.184 | 1.053 | 0.008 | -0.466 | 2.112 | 0.024 | -0.348 | 1.624 | 0 | -1 | -1 | 0 | 0.050 |
| Fn1 | 0.0989 | -0.219 | 1.005 | 0.035 | -0.518 | 1.452 | 0.049 | -0.343 | 1.305 | 0 | -1 | -1 | 0 | 0.050 |
| Thbs2 | 0.4245 | 0.031 | 0.372 | 0.031 | -0.329 | 1.508 | 0.026 | -0.339 | 1.591 | 0 | -1 | -1 | 0 | 0.050 |
| Dctd | 0.0965 | -0.049 | 1.015 | 0.013 | -0.289 | 1.870 | 0.034 | -0.338 | 1.472 | 0 | -1 | -1 | 0 | 0.050 |
| Znf511 | 0.2644 | -0.104 | 0.578 | 0.019 | -0.246 | 1.726 | 0.037 | -0.338 | 1.434 | 0 | -1 | -1 | 0 | 0.050 |
| Poglut1 | 0.0244 | -0.078 | 1.613 | 0.027 | -0.194 | 1.568 | 0.003 | -0.331 | 2.517 | 0 | -1 | -1 | 0 | 0.050 |
| Limch1 | 0.0797 | -0.179 | 1.099 | 0.003 | -0.299 | 2.461 | 0.001 | -0.331 | 2.873 | 0 | -1 | -1 | 0 | 0.050 |
| Znf768 | 0.2171 | -0.135 | 0.663 | 0.036 | -0.169 | 1.440 | 0.005 | -0.330 | 2.265 | 0 | -1 | -1 | 0 | 0.050 |
| Bmpr1a | 0.0523 | -0.240 | 1.281 | 0.004 | -0.319 | 2.445 | 0.005 | -0.328 | 2.326 | 0 | -1 | -1 | 0 | 0.050 |
| Prdx3 | 0.2909 | 0.082 | 0.536 | 0.012 | -0.445 | 1.936 | 0.004 | -0.323 | 2.390 | 0 | -1 | -1 | 0 | 0.050 |
| Prss23 | 0.1592 | -0.074 | 0.798 | 0.010 | -0.271 | 2.011 | 0.002 | -0.320 | 2.611 | 0 | -1 | -1 | 0 | 0.050 |
| Tnrc18 | 0.3437 | -0.140 | 0.464 | 0.034 | -0.347 | 1.471 | 0.004 | -0.314 | 2.444 | 0 | -1 | -1 | 0 | 0.050 |

| | | | | | | | | | | | | | | |
|---|---|---|---|---|---|---|---|---|---|---|---|---|---|---|
| Znf672 | 0.0947 | -0.152 | 1.023 | 0.009 | -0.145 | 2.024 | 0.007 | -0.304 | 2.152 | 0 | -1 | -1 | 0 | 0.050 |
| Itm2c | 0.3144 | -0.066 | 0.503 | 0.011 | -0.401 | 1.949 | 0.033 | -0.303 | 1.482 | 0 | -1 | -1 | 0 | 0.050 |
| H2-K1 | 0.0622 | -0.109 | 1.206 | 0.001 | -0.321 | 3.084 | 0.008 | -0.303 | 2.115 | 0 | -1 | -1 | 0 | 0.050 |
| Chst14 | 0.1064 | 0.047 | 0.973 | 0.023 | -0.159 | 1.639 | 0.001 | -0.301 | 3.065 | 0 | -1 | -1 | 0 | 0.050 |
| Morf4l1 | 0.2569 | -0.128 | 0.590 | 0.002 | -0.281 | 2.716 | 0.003 | -0.301 | 2.456 | 0 | -1 | -1 | 0 | 0.050 |
| Mfge8 | 0.2783 | -0.062 | 0.555 | 0.008 | -0.305 | 2.091 | 0.015 | -0.299 | 1.835 | 0 | -1 | -1 | 0 | 0.050 |
| Rpl10 | 0.7126 | -0.039 | 0.147 | 0.034 | -0.220 | 1.470 | 0.002 | -0.297 | 2.674 | 0 | -1 | -1 | 0 | 0.050 |
| Msrb3 | 0.0819 | -0.077 | 1.087 | 0.002 | -0.237 | 2.728 | 0.011 | -0.294 | 1.941 | 0 | -1 | -1 | 0 | 0.050 |
| Glud1 | 0.0224 | -0.125 | 1.651 | 0.049 | -0.254 | 1.310 | 0.000 | -0.292 | 3.462 | 0 | -1 | -1 | 0 | 0.050 |
| Rdh10 | 0.1777 | -0.051 | 0.750 | 0.041 | -0.148 | 1.383 | 0.001 | -0.291 | 2.867 | 0 | -1 | -1 | 0 | 0.050 |
| Phc2 | 0.0107 | -0.133 | 1.972 | 0.001 | -0.265 | 3.012 | 0.001 | -0.290 | 2.965 | 0 | -1 | -1 | 0 | 0.050 |
| Psma4 | 0.2272 | -0.062 | 0.644 | 0.002 | -0.286 | 2.737 | 0.007 | -0.289 | 2.137 | 0 | -1 | -1 | 0 | 0.050 |
| Cd109 | 0.0631 | -0.103 | 1.200 | 0.003 | -0.400 | 2.555 | 0.031 | -0.288 | 1.515 | 0 | -1 | -1 | 0 | 0.050 |
| Sdc2 | 0.1477 | -0.192 | 0.831 | 0.019 | -0.175 | 1.729 | 0.002 | -0.283 | 2.650 | 0 | -1 | -1 | 0 | 0.050 |
| Rnf150 | 0.2041 | -0.068 | 0.690 | 0.011 | -0.240 | 1.956 | 0.006 | -0.283 | 2.253 | 0 | -1 | -1 | 0 | 0.050 |
| Zc3h14 | 0.0129 | -0.108 | 1.891 | 0.003 | -0.254 | 2.480 | 0.004 | -0.282 | 2.395 | 0 | -1 | -1 | 0 | 0.050 |
| Map3k11 | 0.1598 | -0.163 | 0.796 | 0.004 | -0.292 | 2.406 | 0.020 | -0.282 | 1.702 | 0 | -1 | -1 | 0 | 0.050 |
| Tbrg1 | 0.2292 | -0.055 | 0.640 | 0.038 | -0.282 | 1.422 | 0.020 | -0.281 | 1.692 | 0 | -1 | -1 | 0 | 0.050 |
| Sema3b | 0.0068 | -0.088 | 2.168 | 0.002 | -0.290 | 2.720 | 0.015 | -0.281 | 1.838 | 0 | -1 | -1 | 0 | 0.050 |
| Tmod2 | 0.8204 | 0.059 | 0.086 | 0.017 | -0.312 | 1.765 | 0.034 | -0.281 | 1.467 | 0 | -1 | -1 | 0 | 0.050 |
| F3 | 0.4739 | -0.037 | 0.324 | 0.027 | -0.323 | 1.563 | 0.019 | -0.280 | 1.727 | 0 | -1 | -1 | 0 | 0.050 |
| Nrp2 | 0.0163 | -0.112 | 1.788 | 0.000 | -0.302 | 3.350 | 0.049 | -0.280 | 1.306 | 0 | -1 | -1 | 0 | 0.050 |
| Crlf1 | 0.8518 | 0.014 | 0.070 | 0.013 | -0.296 | 1.898 | 0.001 | -0.277 | 3.023 | 0 | -1 | -1 | 0 | 0.050 |
| Cpped1 | 0.2598 | -0.054 | 0.585 | 0.010 | -0.235 | 1.981 | 0.019 | -0.276 | 1.719 | 0 | -1 | -1 | 0 | 0.050 |
| Ifngr1 | 0.0115 | -0.133 | 1.938 | 0.003 | -0.267 | 2.460 | 0.010 | -0.275 | 2.012 | 0 | -1 | -1 | 0 | 0.050 |
| Lrig2 | 0.0621 | -0.202 | 1.207 | 0.023 | -0.297 | 1.634 | 0.002 | -0.275 | 2.662 | 0 | -1 | -1 | 0 | 0.050 |
| Dchs1 | 0.1601 | -0.065 | 0.796 | 0.001 | -0.308 | 2.956 | 0.040 | -0.274 | 1.398 | 0 | -1 | -1 | 0 | 0.050 |
| Adgrg1 | 0.1133 | -0.196 | 0.946 | 0.049 | -0.395 | 1.309 | 0.000 | -0.274 | 4.109 | 0 | -1 | -1 | 0 | 0.050 |
| Pcdhb17 | 0.0877 | -0.122 | 1.057 | 0.013 | -0.308 | 1.899 | 0.027 | -0.272 | 1.574 | 0 | -1 | -1 | 0 | 0.050 |
| Tmem176b | 0.1041 | -0.245 | 0.983 | 0.043 | -0.270 | 1.371 | 0.002 | -0.271 | 2.650 | 0 | -1 | -1 | 0 | 0.050 |
| Itprip | 0.8823 | -0.027 | 0.054 | 0.002 | -0.281 | 2.640 | 0.028 | -0.269 | 1.554 | 0 | -1 | -1 | 0 | 0.050 |
| Htra1 | 0.0709 | -0.074 | 1.149 | 0.025 | -0.243 | 1.601 | 0.028 | -0.265 | 1.557 | 0 | -1 | -1 | 0 | 0.050 |
| Axl | 0.0503 | -0.161 | 1.298 | 0.005 | -0.264 | 2.345 | 0.015 | -0.263 | 1.817 | 0 | -1 | -1 | 0 | 0.050 |
| Mtpn | 0.9788 | -0.003 | 0.009 | 0.020 | -0.168 | 1.703 | 0.007 | -0.262 | 2.145 | 0 | -1 | -1 | 0 | 0.050 |
| Bmp1 | 0.0078 | -0.130 | 2.108 | 0.002 | -0.325 | 2.767 | 0.004 | -0.262 | 2.444 | 0 | -1 | -1 | 0 | 0.050 |
| Cd14 | 0.0505 | -0.245 | 1.297 | 0.043 | -0.401 | 1.365 | 0.007 | -0.262 | 2.135 | 0 | -1 | -1 | 0 | 0.050 |
| H2-Q4 | 0.0770 | -0.110 | 1.114 | 0.009 | -0.272 | 2.051 | 0.030 | -0.262 | 1.517 | 0 | -1 | -1 | 0 | 0.050 |
| Zfp646 | 0.3378 | -0.175 | 0.471 | 0.048 | -0.228 | 1.315 | 0.007 | -0.261 | 2.182 | 0 | -1 | -1 | 0 | 0.050 |
| Zbtb7a | 0.0177 | -0.080 | 1.752 | 0.034 | -0.260 | 1.475 | 0.001 | -0.259 | 2.955 | 0 | -1 | -1 | 0 | 0.050 |
| Inpp5a | 0.1279 | -0.114 | 0.893 | 0.008 | -0.289 | 2.092 | 0.015 | -0.258 | 1.810 | 0 | -1 | -1 | 0 | 0.050 |
| Ece1 | 0.5802 | -0.024 | 0.236 | 0.005 | -0.310 | 2.297 | 0.034 | -0.253 | 1.471 | 0 | -1 | -1 | 0 | 0.050 |
| Sp1 | 0.0933 | -0.192 | 1.030 | 0.034 | -0.238 | 1.467 | 0.008 | -0.250 | 2.101 | 0 | -1 | -1 | 0 | 0.050 |
| Il17rd | 0.1062 | -0.086 | 0.974 | 0.005 | -0.274 | 2.284 | 0.003 | -0.247 | 2.599 | 0 | -1 | -1 | 0 | 0.050 |
| Pip4p2 | 0.3135 | -0.053 | 0.504 | 0.032 | -0.182 | 1.498 | 0.004 | -0.247 | 2.371 | 0 | -1 | -1 | 0 | 0.050 |
| Haghl | 0.1512 | -0.252 | 0.821 | 0.009 | -0.298 | 2.058 | 0.029 | -0.245 | 1.539 | 0 | -1 | -1 | 0 | 0.050 |
| Gps2 | 0.0273 | -0.089 | 1.565 | 0.008 | -0.203 | 2.080 | 0.006 | -0.243 | 2.251 | 0 | -1 | -1 | 0 | 0.050 |
| Gmppb | 0.6290 | -0.020 | 0.201 | 0.047 | -0.170 | 1.325 | 0.017 | -0.243 | 1.766 | 0 | -1 | -1 | 0 | 0.050 |
| Ltbp1 | 0.0463 | -0.074 | 1.335 | 0.002 | -0.303 | 2.676 | 0.037 | -0.243 | 1.434 | 0 | -1 | -1 | 0 | 0.050 |
| Matn2 | 0.8632 | -0.026 | 0.064 | 0.000 | -0.313 | 3.322 | 0.033 | -0.242 | 1.477 | 0 | -1 | -1 | 0 | 0.050 |
| Clstn1 | 0.5770 | 0.038 | 0.239 | 0.001 | -0.457 | 2.954 | 0.017 | -0.242 | 1.776 | 0 | -1 | -1 | 0 | 0.050 |
| Znf319 | 0.2881 | -0.066 | 0.541 | 0.049 | -0.160 | 1.312 | 0.013 | -0.241 | 1.874 | 0 | -1 | -1 | 0 | 0.050 |
| Enpp1 | 0.1309 | -0.061 | 0.883 | 0.016 | -0.302 | 1.787 | 0.001 | -0.237 | 2.965 | 0 | -1 | -1 | 0 | 0.050 |
| Tnfrsf11b | 0.6155 | -0.068 | 0.211 | 0.004 | -0.309 | 2.356 | 0.030 | -0.236 | 1.517 | 0 | -1 | -1 | 0 | 0.050 |
| Dnase1l1 | 0.0039 | -0.121 | 2.410 | 0.004 | -0.330 | 2.409 | 0.006 | -0.235 | 2.244 | 0 | -1 | -1 | 0 | 0.050 |
| Minar1 | 0.3451 | -0.106 | 0.462 | 0.039 | -0.238 | 1.411 | 0.009 | -0.233 | 2.054 | 0 | -1 | -1 | 0 | 0.050 |
| Cdh13 | 0.0760 | -0.109 | 1.119 | 0.014 | -0.327 | 1.842 | 0.017 | -0.233 | 1.765 | 0 | -1 | -1 | 0 | 0.050 |
| Btg3 | 0.1834 | -0.098 | 0.737 | 0.004 | -0.290 | 2.452 | 0.001 | -0.232 | 3.035 | 0 | -1 | -1 | 0 | 0.050 |
| Ankh | 0.0242 | -0.104 | 1.617 | 0.005 | -0.213 | 2.300 | 0.000 | -0.232 | 3.694 | 0 | -1 | -1 | 0 | 0.050 |
| Tcaf1 | 0.1748 | -0.108 | 0.757 | 0.013 | -0.241 | 1.889 | 0.000 | -0.230 | 3.344 | 0 | -1 | -1 | 0 | 0.050 |
| Tenm4 | 0.0648 | -0.129 | 1.188 | 0.002 | -0.248 | 2.723 | 0.049 | -0.230 | 1.309 | 0 | -1 | -1 | 0 | 0.050 |
| Lox | 0.1645 | -0.043 | 0.784 | 0.027 | -0.219 | 1.567 | 0.003 | -0.228 | 2.557 | 0 | -1 | -1 | 0 | 0.050 |
| Coq4 | 0.8523 | -0.023 | 0.069 | 0.006 | -0.143 | 2.228 | 0.044 | -0.228 | 1.358 | 0 | -1 | -1 | 0 | 0.050 |
| Zwint | 0.2835 | -0.100 | 0.547 | 0.010 | -0.200 | 2.014 | 0.001 | -0.226 | 3.181 | 0 | -1 | -1 | 0 | 0.050 |
| Yipf1 | 0.8343 | -0.024 | 0.079 | 0.008 | -0.250 | 2.111 | 0.007 | -0.226 | 2.179 | 0 | -1 | -1 | 0 | 0.050 |
| Orc6 | 0.3845 | -0.052 | 0.415 | 0.014 | -0.240 | 1.839 | 0.000 | -0.224 | 3.426 | 0 | -1 | -1 | 0 | 0.050 |
| Il13ra1 | 0.7437 | -0.016 | 0.129 | 0.017 | -0.297 | 1.762 | 0.010 | -0.223 | 2.011 | 0 | -1 | -1 | 0 | 0.050 |
| Flrt2 | 0.2392 | -0.103 | 0.621 | 0.012 | -0.266 | 1.938 | 0.003 | -0.221 | 2.465 | 0 | -1 | -1 | 0 | 0.050 |
| Tfrc | 0.9505 | -0.004 | 0.022 | 0.023 | -0.338 | 1.641 | 0.030 | -0.221 | 1.525 | 0 | -1 | -1 | 0 | 0.050 |
| Soat1 | 0.0151 | -0.118 | 1.822 | 0.005 | -0.232 | 2.317 | 0.000 | -0.220 | 3.817 | 0 | -1 | -1 | 0 | 0.050 |
| Chek1 | 0.1004 | -0.139 | 0.998 | 0.002 | -0.268 | 2.703 | 0.006 | -0.219 | 2.196 | 0 | -1 | -1 | 0 | 0.050 |
| Lrrc32 | 0.0648 | -0.185 | 1.188 | 0.010 | -0.172 | 1.990 | 0.034 | -0.218 | 1.473 | 0 | -1 | -1 | 0 | 0.050 |

| | | | | | | | | | | | | | | |
|---|---|---|---|---|---|---|---|---|---|---|---|---|---|---|
| Serinc1 | 0.0568 | -0.194 | 1.246 | 0.003 | -0.219 | 2.496 | 0.001 | -0.217 | 2.905 | 0 | -1 | -1 | 0 | 0.050 |
| Cdh10 | 0.3102 | -0.107 | 0.508 | 0.021 | -0.228 | 1.675 | 0.029 | -0.217 | 1.544 | 0 | -1 | -1 | 0 | 0.050 |
| Ly75 | 0.8825 | -0.004 | 0.054 | 0.030 | -0.284 | 1.530 | 0.045 | -0.217 | 1.344 | 0 | -1 | -1 | 0 | 0.050 |
| Mme | 0.0262 | -0.119 | 1.582 | 0.042 | -0.219 | 1.377 | 0.050 | -0.216 | 1.305 | 0 | -1 | -1 | 0 | 0.050 |
| Itfg1 | 0.8323 | -0.025 | 0.080 | 0.006 | -0.275 | 2.228 | 0.029 | -0.215 | 1.531 | 0 | -1 | -1 | 0 | 0.050 |
| Ext2 | 0.0496 | -0.053 | 1.304 | 0.004 | -0.286 | 2.371 | 0.006 | -0.214 | 2.224 | 0 | -1 | -1 | 0 | 0.050 |
| Iglon5 | 0.4365 | -0.063 | 0.360 | 0.002 | -0.218 | 2.670 | 0.017 | -0.214 | 1.779 | 0 | -1 | -1 | 0 | 0.050 |
| Nr1d2 | 0.3233 | -0.049 | 0.490 | 0.027 | -0.265 | 1.567 | 0.023 | -0.213 | 1.639 | 0 | -1 | -1 | 0 | 0.050 |
| Rps12 | 0.1265 | -0.147 | 0.898 | 0.017 | -0.201 | 1.775 | 0.014 | -0.213 | 1.847 | 0 | -1 | -1 | 0 | 0.050 |
| Abca1 | 0.6311 | -0.023 | 0.200 | 0.005 | -0.262 | 2.292 | 0.017 | -0.212 | 1.758 | 0 | -1 | -1 | 0 | 0.050 |
| Apbb2 | 0.7960 | -0.006 | 0.099 | 0.019 | -0.237 | 1.720 | 0.037 | -0.211 | 1.436 | 0 | -1 | -1 | 0 | 0.050 |
| Slfn9 | 0.1579 | -0.091 | 0.802 | 0.016 | -0.144 | 1.799 | 0.010 | -0.211 | 2.019 | 0 | -1 | -1 | 0 | 0.050 |
| Fam234a | 0.0090 | -0.135 | 2.045 | 0.002 | -0.239 | 2.767 | 0.042 | -0.209 | 1.372 | 0 | -1 | -1 | 0 | 0.050 |
| Tyro3 | 0.0604 | -0.115 | 1.219 | 0.009 | -0.224 | 2.045 | 0.014 | -0.204 | 1.868 | 0 | -1 | -1 | 0 | 0.050 |
| Eno1 | 0.1091 | -0.099 | 0.962 | 0.005 | -0.191 | 2.269 | 0.019 | -0.204 | 1.711 | 0 | -1 | -1 | 0 | 0.050 |
| Tspyl1 | 0.1678 | -0.093 | 0.775 | 0.016 | -0.281 | 1.804 | 0.007 | -0.204 | 2.138 | 0 | -1 | -1 | 0 | 0.050 |
| Zfyve19 | 0.0166 | -0.115 | 1.780 | 0.003 | -0.223 | 2.536 | 0.002 | -0.202 | 2.791 | 0 | -1 | -1 | 0 | 0.050 |
| Ror1 | 0.0961 | -0.145 | 1.017 | 0.014 | -0.209 | 1.848 | 0.030 | -0.202 | 1.529 | 0 | -1 | -1 | 0 | 0.050 |
| Sema4f | 0.2533 | -0.102 | 0.596 | 0.016 | -0.239 | 1.792 | 0.011 | -0.202 | 1.951 | 0 | -1 | -1 | 0 | 0.050 |
| Znf385a | 0.4831 | -0.052 | 0.316 | 0.044 | -0.206 | 1.359 | 0.004 | -0.201 | 2.410 | 0 | -1 | -1 | 0 | 0.050 |
| Met | 0.0219 | -0.127 | 1.660 | 0.003 | -0.255 | 2.512 | 0.032 | -0.201 | 1.498 | 0 | -1 | -1 | 0 | 0.050 |
| Col5a1 | 0.9177 | 0.008 | 0.037 | 0.020 | -0.429 | 1.704 | 0.041 | -0.201 | 1.383 | 0 | -1 | -1 | 0 | 0.050 |
| Loxl4 | 0.2709 | -0.098 | 0.567 | 0.002 | -0.178 | 2.605 | 0.003 | -0.199 | 2.567 | 0 | -1 | -1 | 0 | 0.050 |
| Epha5 | 0.0973 | -0.143 | 1.012 | 0.016 | -0.224 | 1.795 | 0.039 | -0.196 | 1.405 | 0 | -1 | -1 | 0 | 0.050 |
| Sgcb | 0.1147 | -0.108 | 0.941 | 0.014 | -0.226 | 1.861 | 0.038 | -0.194 | 1.417 | 0 | -1 | -1 | 0 | 0.050 |
| Lrrc15 | 0.2152 | -0.079 | 0.667 | 0.006 | -0.164 | 2.203 | 0.032 | -0.193 | 1.500 | 0 | -1 | -1 | 0 | 0.050 |
| Ptprg | 0.2915 | -0.078 | 0.535 | 0.003 | -0.209 | 2.503 | 0.047 | -0.191 | 1.329 | 0 | -1 | -1 | 0 | 0.050 |
| B4galnt1 | 0.7627 | -0.033 | 0.118 | 0.045 | -0.160 | 1.350 | 0.033 | -0.191 | 1.475 | 0 | -1 | -1 | 0 | 0.050 |
| Nrcam | 0.3175 | -0.089 | 0.498 | 0.013 | -0.249 | 1.881 | 0.001 | -0.191 | 2.905 | 0 | -1 | -1 | 0 | 0.050 |
| Moxd1 | 0.5481 | -0.056 | 0.261 | 0.011 | -0.254 | 1.945 | 0.035 | -0.188 | 1.455 | 0 | -1 | -1 | 0 | 0.050 |
| Mccc1 | 0.2157 | -0.068 | 0.666 | 0.023 | -0.184 | 1.645 | 0.019 | -0.188 | 1.726 | 0 | -1 | -1 | 0 | 0.050 |
| Znhit6 | 0.0623 | -0.091 | 1.205 | 0.011 | -0.143 | 1.962 | 0.005 | -0.187 | 2.264 | 0 | -1 | -1 | 0 | 0.050 |
| S1pr1 | 0.0166 | -0.083 | 1.781 | 0.009 | -0.215 | 2.063 | 0.040 | -0.186 | 1.397 | 0 | -1 | -1 | 0 | 0.050 |
| Ap1s1 | 0.0541 | -0.050 | 1.267 | 0.015 | -0.181 | 1.831 | 0.021 | -0.183 | 1.668 | 0 | -1 | -1 | 0 | 0.050 |
| Gcnt1 | 0.2979 | -0.215 | 0.526 | 0.023 | -0.382 | 1.644 | 0.046 | -0.182 | 1.334 | 0 | -1 | -1 | 0 | 0.050 |
| Galnt1 | 0.6515 | 0.050 | 0.186 | 0.004 | -0.185 | 2.437 | 0.045 | -0.181 | 1.344 | 0 | -1 | -1 | 0 | 0.050 |
| Supt20 | 0.2100 | -0.153 | 0.678 | 0.032 | -0.212 | 1.497 | 0.039 | -0.180 | 1.411 | 0 | -1 | -1 | 0 | 0.050 |
| Itga6 | 0.0307 | -0.087 | 1.513 | 0.035 | -0.274 | 1.451 | 0.037 | -0.180 | 1.438 | 0 | -1 | -1 | 0 | 0.050 |
| Adgrl3 | 0.0067 | -0.111 | 2.173 | 0.046 | -0.218 | 1.337 | 0.008 | -0.179 | 2.077 | 0 | -1 | -1 | 0 | 0.050 |
| Plxnb3 | 0.2045 | -0.116 | 0.689 | 0.009 | -0.232 | 2.030 | 0.048 | -0.179 | 1.322 | 0 | -1 | -1 | 0 | 0.050 |
| Galnt2 | 0.9236 | 0.004 | 0.035 | 0.001 | -0.186 | 2.889 | 0.008 | -0.179 | 2.084 | 0 | -1 | -1 | 0 | 0.050 |
| Xaf1 | 0.0053 | -0.130 | 2.277 | 0.005 | -0.149 | 2.295 | 0.002 | -0.178 | 2.733 | 0 | -1 | -1 | 0 | 0.050 |
| Itpa | 0.1550 | -0.087 | 0.810 | 0.007 | -0.189 | 2.147 | 0.020 | -0.178 | 1.695 | 0 | -1 | -1 | 0 | 0.050 |
| Rtp4 | 0.3743 | -0.041 | 0.427 | 0.030 | -0.178 | 1.527 | 0.029 | -0.178 | 1.542 | 0 | -1 | -1 | 0 | 0.050 |
| Mad2l1bp | 0.2752 | -0.114 | 0.560 | 0.032 | -0.158 | 1.499 | 0.010 | -0.178 | 1.992 | 0 | -1 | -1 | 0 | 0.050 |
| Wdr6 | 0.0271 | -0.075 | 1.567 | 0.007 | -0.170 | 2.134 | 0.024 | -0.177 | 1.617 | 0 | -1 | -1 | 0 | 0.050 |
| Tmem106b | 0.4396 | 0.045 | 0.357 | 0.026 | -0.298 | 1.587 | 0.027 | -0.173 | 1.573 | 0 | -1 | -1 | 0 | 0.050 |
| Itgb4 | 0.0838 | -0.108 | 1.077 | 0.043 | -0.178 | 1.368 | 0.045 | -0.171 | 1.351 | 0 | -1 | -1 | 0 | 0.050 |
| Lamb3 | 0.0111 | -0.126 | 1.954 | 0.016 | -0.227 | 1.804 | 0.017 | -0.171 | 1.765 | 0 | -1 | -1 | 0 | 0.050 |
| Zbtb1 | 0.0811 | -0.189 | 1.091 | 0.005 | -0.336 | 2.294 | 0.048 | -0.171 | 1.316 | 0 | -1 | -1 | 0 | 0.050 |
| Aldh1l1 | 0.3482 | 0.032 | 0.458 | 0.005 | -0.226 | 2.315 | 0.045 | -0.170 | 1.343 | 0 | -1 | -1 | 0 | 0.050 |
| Itga7 | 0.0633 | -0.139 | 1.199 | 0.019 | -0.207 | 1.723 | 0.040 | -0.169 | 1.401 | 0 | -1 | -1 | 0 | 0.050 |
| Tlr2 | 0.0857 | -0.120 | 1.067 | 0.022 | -0.216 | 1.663 | 0.030 | -0.159 | 1.521 | 0 | -1 | -1 | 0 | 0.050 |
| Frk | 0.0750 | -0.077 | 1.125 | 0.002 | -0.222 | 2.803 | 0.009 | -0.156 | 2.060 | 0 | -1 | -1 | 0 | 0.050 |
| Dpy19l4 | 0.0385 | -0.101 | 1.415 | 0.007 | -0.169 | 2.168 | 0.030 | -0.154 | 1.523 | 0 | -1 | -1 | 0 | 0.050 |
| Nrn1 | 0.1487 | -0.157 | 0.828 | 0.020 | -0.351 | 1.705 | 0.011 | -0.154 | 1.953 | 0 | -1 | -1 | 0 | 0.050 |
| Wls | 0.0757 | -0.076 | 1.121 | 0.014 | -0.179 | 1.846 | 0.027 | -0.153 | 1.571 | 0 | -1 | -1 | 0 | 0.050 |
| Tmem63a | 0.1308 | -0.092 | 0.883 | 0.002 | -0.158 | 2.603 | 0.034 | -0.153 | 1.471 | 0 | -1 | -1 | 0 | 0.050 |
| Carmil2 | 0.8431 | 0.030 | 0.074 | 0.030 | -0.148 | 1.519 | 0.005 | -0.152 | 2.275 | 0 | -1 | -1 | 0 | 0.050 |
| Nr2c1 | 0.8485 | -0.017 | 0.071 | 0.001 | -0.223 | 2.978 | 0.020 | -0.151 | 1.701 | 0 | -1 | -1 | 0 | 0.050 |
| Tmem231 | 0.1336 | -0.116 | 0.874 | 0.007 | -0.163 | 2.133 | 0.038 | -0.151 | 1.416 | 0 | -1 | -1 | 0 | 0.050 |
| Grb10 | 0.0858 | -0.076 | 1.066 | 0.027 | -0.139 | 1.575 | 0.001 | -0.148 | 2.857 | 0 | -1 | -1 | 0 | 0.050 |
| Itgb1 | 0.0230 | -0.090 | 1.638 | 0.008 | -0.239 | 2.107 | 0.026 | -0.147 | 1.584 | 0 | -1 | -1 | 0 | 0.050 |
| Atic | 0.3927 | 0.022 | 0.406 | 0.008 | -0.157 | 2.120 | 0.027 | -0.146 | 1.570 | 0 | -1 | -1 | 0 | 0.050 |
| Gatd3a | 0.2629 | -0.083 | 0.580 | 0.007 | -0.230 | 2.152 | 0.008 | -0.145 | 2.115 | 0 | -1 | -1 | 0 | 0.050 |
| Shroom4 | 0.2402 | -0.071 | 0.619 | 0.046 | -0.143 | 1.340 | 0.013 | -0.144 | 1.899 | 0 | -1 | -1 | 0 | 0.050 |
| Pcca | 0.3784 | -0.039 | 0.422 | 0.036 | -0.275 | 1.443 | 0.010 | -0.144 | 2.010 | 0 | -1 | -1 | 0 | 0.050 |
| Unc93b1 | 0.7463 | -0.056 | 0.127 | 0.002 | -0.180 | 2.806 | 0.002 | -0.143 | 2.811 | 0 | -1 | -1 | 0 | 0.050 |
| Aurkb | 0.0593 | -0.126 | 1.227 | 0.029 | -0.197 | 1.533 | 0.003 | -0.143 | 2.487 | 0 | -1 | -1 | 0 | 0.050 |
| Prc1 | 0.2482 | -0.059 | 0.605 | 0.014 | -0.184 | 1.855 | 0.002 | -0.139 | 2.675 | 0 | -1 | -1 | 0 | 0.050 |
| Sulf1 | 0.5662 | -0.052 | 0.247 | 0.002 | -0.551 | 2.634 | 0.061 | -0.454 | 1.218 | 0 | -1 | 0 | 0 | 0.050 |

| | | | | | | | | | | | | | | |
|---|---|---|---|---|---|---|---|---|---|---|---|---|---|---|
| Pcdhb20 | 0.0434 | -0.093 | 1.363 | 0.007 | -0.532 | 2.184 | 0.157 | -0.439 | 0.804 | 0 | -1 | 0 | 0 | 0.050 |
| Ctnna1 | 0.3585 | -0.062 | 0.445 | 0.010 | -0.650 | 2.017 | 0.077 | -0.424 | 1.115 | 0 | -1 | 0 | 0 | 0.050 |
| Ccdc80 | 0.3602 | -0.044 | 0.444 | 0.004 | -0.531 | 2.349 | 0.102 | -0.401 | 0.992 | 0 | -1 | 0 | 0 | 0.050 |
| Lifr | 0.6928 | -0.020 | 0.159 | 0.005 | -0.446 | 2.274 | 0.106 | -0.394 | 0.973 | 0 | -1 | 0 | 0 | 0.050 |
| Fbn2 | 0.7483 | -0.020 | 0.126 | 0.020 | -0.449 | 1.699 | 0.055 | -0.385 | 1.263 | 0 | -1 | 0 | 0 | 0.050 |
| Col1a2 | 0.9538 | -0.003 | 0.021 | 0.022 | -0.578 | 1.658 | 0.110 | -0.379 | 0.960 | 0 | -1 | 0 | 0 | 0.050 |
| Hdac6 | 0.1101 | -0.038 | 0.958 | 0.032 | -0.404 | 1.501 | 0.100 | -0.374 | 0.998 | 0 | -1 | 0 | 0 | 0.050 |
| Loxl2 | 0.8564 | -0.005 | 0.067 | 0.040 | -0.512 | 1.394 | 0.060 | -0.354 | 1.225 | 0 | -1 | 0 | 0 | 0.050 |
| Col1a1 | 0.4436 | -0.044 | 0.353 | 0.023 | -0.656 | 1.641 | 0.114 | -0.345 | 0.943 | 0 | -1 | 0 | 0 | 0.050 |
| Pcdhb16 | 0.3807 | -0.152 | 0.419 | 0.003 | -0.405 | 2.544 | 0.074 | -0.332 | 1.131 | 0 | -1 | 0 | 0 | 0.050 |
| Gmds | 0.6729 | -0.023 | 0.172 | 0.004 | -0.317 | 2.366 | 0.053 | -0.323 | 1.274 | 0 | -1 | 0 | 0 | 0.050 |
| Cct8l1 | 0.1824 | -0.130 | 0.739 | 0.001 | -0.347 | 3.234 | 0.065 | -0.319 | 1.186 | 0 | -1 | 0 | 0 | 0.050 |
| Ptx3 | 0.7317 | -0.020 | 0.136 | 0.026 | -0.337 | 1.583 | 0.077 | -0.318 | 1.113 | 0 | -1 | 0 | 0 | 0.050 |
| Mgme1 | 0.4891 | -0.021 | 0.311 | 0.005 | -0.242 | 2.311 | 0.100 | -0.314 | 1.002 | 0 | -1 | 0 | 0 | 0.050 |
| Dkk2 | 0.0085 | -0.101 | 2.069 | 0.022 | -0.243 | 1.666 | 0.050 | -0.312 | 1.299 | 0 | -1 | 0 | 0 | 0.050 |
| Znf318 | 0.7695 | -0.037 | 0.114 | 0.030 | -0.457 | 1.527 | 0.070 | -0.301 | 1.154 | 0 | -1 | 0 | 0 | 0.050 |
| Tgfb1 | 0.2632 | -0.120 | 0.580 | 0.018 | -0.289 | 1.740 | 0.080 | -0.299 | 1.095 | 0 | -1 | 0 | 0 | 0.050 |
| Osmr | 0.0159 | -0.119 | 1.798 | 0.000 | -0.381 | 3.672 | 0.059 | -0.296 | 1.227 | 0 | -1 | 0 | 0 | 0.050 |
| Col2a1 | 0.3324 | 0.070 | 0.478 | 0.040 | -0.502 | 1.397 | 0.055 | -0.287 | 1.258 | 0 | -1 | 0 | 0 | 0.050 |
| Tenm2 | 0.1433 | -0.147 | 0.844 | 0.009 | -0.347 | 2.029 | 0.051 | -0.286 | 1.293 | 0 | -1 | 0 | 0 | 0.050 |
| Frmpd1 | 0.9215 | 0.009 | 0.036 | 0.004 | -0.357 | 2.358 | 0.104 | -0.281 | 0.985 | 0 | -1 | 0 | 0 | 0.050 |
| Pcolce2 | 0.4226 | -0.071 | 0.374 | 0.028 | -0.192 | 1.549 | 0.063 | -0.279 | 1.203 | 0 | -1 | 0 | 0 | 0.050 |
| Gm8909 | 0.0345 | -0.133 | 1.462 | 0.004 | -0.364 | 2.437 | 0.053 | -0.265 | 1.280 | 0 | -1 | 0 | 0 | 0.050 |
| Ccdc8 | 0.2144 | -0.145 | 0.669 | 0.049 | -0.302 | 1.312 | 0.114 | -0.259 | 0.944 | 0 | -1 | 0 | 0 | 0.050 |
| Zfand6 | 0.0845 | -0.176 | 1.073 | 0.019 | -0.308 | 1.732 | 0.306 | -0.255 | 0.514 | 0 | -1 | 0 | 0 | 0.050 |
| Zfp267 | 0.4559 | -0.056 | 0.341 | 0.038 | -0.343 | 1.425 | 0.122 | -0.249 | 0.915 | 0 | -1 | 0 | 0 | 0.050 |
| Spint1 | 0.0050 | -0.091 | 2.302 | 0.007 | -0.286 | 2.132 | 0.065 | -0.247 | 1.187 | 0 | -1 | 0 | 0 | 0.050 |
| Gpc4 | 0.0762 | -0.080 | 1.118 | 0.002 | -0.351 | 2.678 | 0.161 | -0.242 | 0.793 | 0 | -1 | 0 | 0 | 0.050 |
| Flrt3 | 0.0224 | -0.114 | 1.649 | 0.001 | -0.301 | 2.889 | 0.074 | -0.242 | 1.133 | 0 | -1 | 0 | 0 | 0.050 |
| 2610005L07R | 0.1935 | -0.206 | 0.713 | 0.006 | -0.171 | 2.243 | 0.076 | -0.241 | 1.122 | 0 | -1 | 0 | 0 | 0.050 |
| Chchd6 | 0.9282 | -0.024 | 0.032 | 0.017 | -0.169 | 1.766 | 0.059 | -0.241 | 1.230 | 0 | -1 | 0 | 0 | 0.050 |
| Fgfr2 | 0.0796 | -0.115 | 1.099 | 0.001 | -0.257 | 3.128 | 0.050 | -0.237 | 1.300 | 0 | -1 | 0 | 0 | 0.050 |
| Flrt1 | 0.0686 | -0.118 | 1.164 | 0.001 | -0.282 | 2.902 | 0.052 | -0.235 | 1.284 | 0 | -1 | 0 | 0 | 0.050 |
| Klhdc2 | 0.0817 | -0.249 | 1.088 | 0.049 | -0.185 | 1.309 | 0.097 | -0.234 | 1.015 | 0 | -1 | 0 | 0 | 0.050 |
| Adamtsl4 | 0.3046 | 0.036 | 0.516 | 0.026 | -0.233 | 1.590 | 0.066 | -0.233 | 1.179 | 0 | -1 | 0 | 0 | 0.050 |
| Igsf3 | 0.0553 | -0.086 | 1.257 | 0.000 | -0.311 | 3.813 | 0.068 | -0.231 | 1.168 | 0 | -1 | 0 | 0 | 0.050 |
| Cracd | 0.7132 | -0.080 | 0.147 | 0.010 | -0.309 | 1.992 | 0.193 | -0.231 | 0.714 | 0 | -1 | 0 | 0 | 0.050 |
| Slit3 | 0.3088 | -0.054 | 0.510 | 0.005 | -0.292 | 2.335 | 0.113 | -0.230 | 0.945 | 0 | -1 | 0 | 0 | 0.050 |
| Fzd7 | 0.0681 | -0.097 | 1.167 | 0.000 | -0.254 | 3.322 | 0.062 | -0.228 | 1.209 | 0 | -1 | 0 | 0 | 0.050 |
| Pja1 | 0.0785 | -0.216 | 1.105 | 0.039 | -0.419 | 1.404 | 0.234 | -0.228 | 0.631 | 0 | -1 | 0 | 0 | 0.050 |
| H2-Q7 | 0.0601 | -0.097 | 1.221 | 0.001 | -0.264 | 2.947 | 0.054 | -0.227 | 1.269 | 0 | -1 | 0 | 0 | 0.050 |
| Gapdhs | 0.8574 | -0.017 | 0.067 | 0.008 | -0.528 | 2.121 | 0.453 | -0.225 | 0.344 | 0 | -1 | 0 | 0 | 0.050 |
| Klhdc3 | 0.2441 | 0.154 | 0.612 | 0.013 | -0.286 | 1.878 | 0.075 | -0.225 | 1.123 | 0 | -1 | 0 | 0 | 0.050 |
| Prg4 | 0.0442 | 0.078 | 1.354 | 0.000 | -0.308 | 3.629 | 0.183 | -0.224 | 0.738 | 0 | -1 | 0 | 0 | 0.050 |
| Fzd1 | 0.0899 | -0.120 | 1.046 | 0.002 | -0.253 | 2.775 | 0.125 | -0.224 | 0.904 | 0 | -1 | 0 | 0 | 0.050 |
| Gtf3a | 0.6790 | -0.029 | 0.168 | 0.007 | -0.246 | 2.186 | 0.140 | -0.223 | 0.853 | 0 | -1 | 0 | 0 | 0.050 |
| L1cam | 0.9508 | 0.002 | 0.022 | 0.011 | -0.343 | 1.978 | 0.083 | -0.223 | 1.081 | 0 | -1 | 0 | 0 | 0.050 |
| Itih5 | 0.9946 | 0.000 | 0.002 | 0.023 | -0.285 | 1.639 | 0.075 | -0.221 | 1.128 | 0 | -1 | 0 | 0 | 0.050 |
| Ptgfrn | 0.0403 | -0.103 | 1.394 | 0.001 | -0.275 | 2.891 | 0.074 | -0.221 | 1.128 | 0 | -1 | 0 | 0 | 0.050 |
| Gpc2 | 0.5539 | -0.076 | 0.257 | 0.004 | -0.205 | 2.399 | 0.143 | -0.220 | 0.846 | 0 | -1 | 0 | 0 | 0.050 |
| Trim8 | 0.0898 | -0.162 | 1.047 | 0.025 | -0.324 | 1.599 | 0.060 | -0.218 | 1.221 | 0 | -1 | 0 | 0 | 0.050 |
| Lamp2 | 0.1260 | 0.173 | 0.899 | 0.000 | -0.311 | 3.672 | 0.082 | -0.217 | 1.086 | 0 | -1 | 0 | 0 | 0.050 |
| Alcam | 0.0079 | -0.091 | 2.100 | 0.000 | -0.280 | 3.349 | 0.239 | -0.216 | 0.622 | 0 | -1 | 0 | 0 | 0.050 |
| Pcdhb18 | 0.1315 | -0.164 | 0.881 | 0.006 | -0.143 | 2.220 | 0.114 | -0.215 | 0.942 | 0 | -1 | 0 | 0 | 0.050 |
| Epha4 | 0.1679 | -0.080 | 0.775 | 0.007 | -0.207 | 2.166 | 0.096 | -0.207 | 1.020 | 0 | -1 | 0 | 0 | 0.050 |
| Mmp14 | 0.1031 | -0.076 | 0.987 | 0.014 | -0.258 | 1.869 | 0.073 | -0.207 | 1.137 | 0 | -1 | 0 | 0 | 0.050 |
| 3000002C10F | 0.1168 | -0.172 | 0.933 | 0.022 | -0.529 | 1.652 | 0.498 | -0.206 | 0.303 | 0 | -1 | 0 | 0 | 0.050 |
| Sirpa | 0.0699 | -0.176 | 1.155 | 0.002 | -0.253 | 2.815 | 0.090 | -0.206 | 1.048 | 0 | -1 | 0 | 0 | 0.050 |
| Igsf10 | 0.6728 | -0.010 | 0.172 | 0.006 | -0.296 | 2.208 | 0.191 | -0.205 | 0.719 | 0 | -1 | 0 | 0 | 0.050 |
| Cemip2 | 0.1707 | -0.072 | 0.768 | 0.005 | -0.261 | 2.294 | 0.053 | -0.205 | 1.280 | 0 | -1 | 0 | 0 | 0.050 |
| Clmp | 0.1642 | -0.073 | 0.785 | 0.004 | -0.236 | 2.377 | 0.080 | -0.205 | 1.097 | 0 | -1 | 0 | 0 | 0.050 |
| Znf787 | 0.4065 | 0.062 | 0.391 | 0.044 | -0.188 | 1.356 | 0.150 | -0.204 | 0.825 | 0 | -1 | 0 | 0 | 0.050 |
| Enpp3 | 0.7055 | -0.034 | 0.152 | 0.009 | -0.206 | 2.064 | 0.101 | -0.204 | 0.997 | 0 | -1 | 0 | 0 | 0.050 |
| Cd248 | 0.0208 | -0.134 | 1.682 | 0.002 | -0.245 | 2.820 | 0.110 | -0.203 | 0.960 | 0 | -1 | 0 | 0 | 0.050 |
| Cd44 | 0.0718 | -0.208 | 1.144 | 0.010 | -0.197 | 1.979 | 0.052 | -0.203 | 1.288 | 0 | -1 | 0 | 0 | 0.050 |
| Edil3 | 0.2346 | -0.029 | 0.630 | 0.009 | -0.232 | 2.042 | 0.051 | -0.202 | 1.295 | 0 | -1 | 0 | 0 | 0.050 |
| Cdh17 | 0.5690 | 0.013 | 0.245 | 0.006 | -0.282 | 2.204 | 0.075 | -0.201 | 1.123 | 0 | -1 | 0 | 0 | 0.050 |
| Plxna4 | 0.0687 | -0.169 | 1.163 | 0.006 | -0.194 | 2.259 | 0.052 | -0.201 | 1.286 | 0 | -1 | 0 | 0 | 0.050 |
| Slc44a1 | 0.1694 | -0.085 | 0.771 | 0.001 | -0.243 | 3.028 | 0.053 | -0.199 | 1.278 | 0 | -1 | 0 | 0 | 0.050 |
| Tenm3 | 0.4081 | -0.052 | 0.389 | 0.006 | -0.248 | 2.254 | 0.163 | -0.198 | 0.787 | 0 | -1 | 0 | 0 | 0.050 |
| Robo2 | 0.0911 | -0.051 | 1.040 | 0.005 | -0.218 | 2.314 | 0.180 | -0.197 | 0.745 | 0 | -1 | 0 | 0 | 0.050 |

| | | | | | | | | | | | | | |
|---|---|---|---|---|---|---|---|---|---|---|---|---|---|
| Il1r1 | 0.0023 | -0.122 | 2.639 | 0.014 | -0.241 | 1.864 | 0.059 | -0.196 | 1.231 | 0 | -1 | 0 | 0 | 0.050 |
| Plxna1 | 0.0696 | -0.142 | 1.158 | 0.011 | -0.225 | 1.955 | 0.070 | -0.195 | 1.156 | 0 | -1 | 0 | 0 | 0.050 |
| Itga5 | 0.0477 | -0.101 | 1.321 | 0.016 | -0.251 | 1.798 | 0.054 | -0.195 | 1.270 | 0 | -1 | 0 | 0 | 0.050 |
| Acyp2 | 0.0018 | -0.125 | 2.743 | 0.011 | -0.154 | 1.963 | 0.061 | -0.194 | 1.216 | 0 | -1 | 0 | 0 | 0.050 |
| Ephb2 | 0.0775 | -0.114 | 1.111 | 0.006 | -0.205 | 2.251 | 0.064 | -0.194 | 1.194 | 0 | -1 | 0 | 0 | 0.050 |
| Itgb5 | 0.6434 | -0.024 | 0.192 | 0.000 | -0.239 | 3.338 | 0.112 | -0.193 | 0.951 | 0 | -1 | 0 | 0 | 0.050 |
| Cd101 | 0.0165 | 0.120 | 1.782 | 0.001 | -0.251 | 3.093 | 0.092 | -0.191 | 1.037 | 0 | -1 | 0 | 0 | 0.050 |
| Itga11 | 0.6665 | -0.021 | 0.176 | 0.007 | -0.346 | 2.169 | 0.356 | -0.189 | 0.448 | 0 | -1 | 0 | 0 | 0.050 |
| Il10rb | 0.0362 | -0.127 | 1.441 | 0.009 | -0.249 | 2.040 | 0.056 | -0.189 | 1.252 | 0 | -1 | 0 | 0 | 0.050 |
| Chmp4b | 0.2586 | -0.062 | 0.587 | 0.010 | -0.282 | 1.997 | 0.050 | -0.189 | 1.297 | 0 | -1 | 0 | 0 | 0.050 |
| Insr | 0.0791 | -0.122 | 1.102 | 0.001 | -0.229 | 3.083 | 0.072 | -0.188 | 1.142 | 0 | -1 | 0 | 0 | 0.050 |
| Ncstn | 0.1392 | -0.148 | 0.856 | 0.033 | -0.175 | 1.480 | 0.066 | -0.187 | 1.178 | 0 | -1 | 0 | 0 | 0.050 |
| Jam3 | 0.1114 | -0.161 | 0.953 | 0.001 | -0.197 | 2.935 | 0.089 | -0.186 | 1.052 | 0 | -1 | 0 | 0 | 0.050 |
| Plxdc2 | 0.0277 | -0.082 | 1.558 | 0.016 | -0.182 | 1.797 | 0.051 | -0.186 | 1.291 | 0 | -1 | 0 | 0 | 0.050 |
| Erbb2 | 0.0380 | -0.112 | 1.420 | 0.015 | -0.226 | 1.829 | 0.069 | -0.186 | 1.160 | 0 | -1 | 0 | 0 | 0.050 |
| Sgcd | 0.1829 | -0.105 | 0.738 | 0.049 | -0.240 | 1.312 | 0.061 | -0.184 | 1.217 | 0 | -1 | 0 | 0 | 0.050 |
| H2-D1 | 0.0850 | -0.112 | 1.070 | 0.004 | -0.211 | 2.449 | 0.109 | -0.184 | 0.964 | 0 | -1 | 0 | 0 | 0.050 |
| Pdgfrb | 0.1453 | -0.076 | 0.838 | 0.005 | -0.221 | 2.331 | 0.138 | -0.183 | 0.862 | 0 | -1 | 0 | 0 | 0.050 |
| Atrnl1 | 0.0050 | -0.097 | 2.299 | 0.009 | -0.219 | 2.040 | 0.054 | -0.183 | 1.268 | 0 | -1 | 0 | 0 | 0.050 |
| Nipsnap3b | 0.4489 | -0.029 | 0.348 | 0.034 | -0.238 | 1.472 | 0.053 | -0.183 | 1.277 | 0 | -1 | 0 | 0 | 0.050 |
| Itga1 | 0.3655 | -0.021 | 0.437 | 0.009 | -0.214 | 2.033 | 0.276 | -0.183 | 0.559 | 0 | -1 | 0 | 0 | 0.050 |
| Kctd12 | 0.9035 | -0.013 | 0.044 | 0.001 | -0.274 | 3.215 | 0.449 | -0.182 | 0.348 | 0 | -1 | 0 | 0 | 0.050 |
| Fat4 | 0.0189 | -0.084 | 1.722 | 0.002 | -0.241 | 2.777 | 0.154 | -0.181 | 0.814 | 0 | -1 | 0 | 0 | 0.050 |
| Ntn1 | 0.0197 | -0.120 | 1.706 | 0.010 | -0.297 | 2.012 | 0.227 | -0.181 | 0.644 | 0 | -1 | 0 | 0 | 0.050 |
| Adgrl2 | 0.0642 | -0.112 | 1.192 | 0.002 | -0.214 | 2.633 | 0.083 | -0.181 | 1.078 | 0 | -1 | 0 | 0 | 0.050 |
| Slc39a6 | 0.2055 | -0.125 | 0.687 | 0.013 | -0.187 | 1.890 | 0.104 | -0.181 | 0.983 | 0 | -1 | 0 | 0 | 0.050 |
| Emb | 0.4763 | -0.052 | 0.322 | 0.048 | -0.206 | 1.319 | 0.079 | -0.180 | 1.102 | 0 | -1 | 0 | 0 | 0.050 |
| Dag1 | 0.3500 | -0.046 | 0.456 | 0.003 | -0.240 | 2.598 | 0.131 | -0.178 | 0.884 | 0 | -1 | 0 | 0 | 0.050 |
| Anpep | 0.2179 | -0.059 | 0.662 | 0.016 | -0.271 | 1.783 | 0.103 | -0.176 | 0.988 | 0 | -1 | 0 | 0 | 0.050 |
| Sgce | 0.1697 | -0.082 | 0.770 | 0.014 | -0.203 | 1.868 | 0.169 | -0.175 | 0.772 | 0 | -1 | 0 | 0 | 0.050 |
| Atrn | 0.0210 | -0.137 | 1.677 | 0.040 | -0.222 | 1.401 | 0.062 | -0.174 | 1.207 | 0 | -1 | 0 | 0 | 0.050 |
| Vta1 | 0.0408 | -0.124 | 1.389 | 0.009 | -0.268 | 2.061 | 0.211 | -0.173 | 0.676 | 0 | -1 | 0 | 0 | 0.050 |
| Adgrg2 | 0.0734 | -0.144 | 1.134 | 0.039 | -0.183 | 1.411 | 0.073 | -0.173 | 1.137 | 0 | -1 | 0 | 0 | 0.050 |
| Plxnb2 | 0.0681 | -0.146 | 1.167 | 0.006 | -0.193 | 2.234 | 0.093 | -0.172 | 1.031 | 0 | -1 | 0 | 0 | 0.050 |
| Ptch1 | 0.2232 | -0.088 | 0.651 | 0.018 | -0.188 | 1.756 | 0.188 | -0.172 | 0.725 | 0 | -1 | 0 | 0 | 0.050 |
| Fzd2 | 0.1575 | -0.088 | 0.803 | 0.005 | -0.270 | 2.329 | 0.170 | -0.172 | 0.770 | 0 | -1 | 0 | 0 | 0.050 |
| Cenpi | 0.3514 | -0.078 | 0.454 | 0.035 | -0.156 | 1.458 | 0.224 | -0.171 | 0.649 | 0 | -1 | 0 | 0 | 0.050 |
| F2r | 0.8057 | 0.073 | 0.094 | 0.041 | -0.285 | 1.389 | 0.177 | -0.171 | 0.752 | 0 | -1 | 0 | 0 | 0.050 |
| Fas | 0.0556 | -0.164 | 1.255 | 0.002 | -0.164 | 2.669 | 0.090 | -0.170 | 1.047 | 0 | -1 | 0 | 0 | 0.050 |
| Ldha | 0.8741 | 0.006 | 0.058 | 0.004 | -0.191 | 2.390 | 0.060 | -0.170 | 1.224 | 0 | -1 | 0 | 0 | 0.050 |
| Cspg4 | 0.0429 | -0.094 | 1.367 | 0.002 | -0.221 | 2.726 | 0.083 | -0.170 | 1.082 | 0 | -1 | 0 | 0 | 0.050 |
| Gm45935 | 0.1972 | -0.149 | 0.705 | 0.032 | -0.150 | 1.498 | 0.189 | -0.169 | 0.724 | 0 | -1 | 0 | 0 | 0.050 |
| Tmem30a | 0.1775 | -0.129 | 0.751 | 0.036 | -0.153 | 1.445 | 0.090 | -0.167 | 1.044 | 0 | -1 | 0 | 0 | 0.050 |
| Nid2 | 0.0677 | -0.127 | 1.169 | 0.044 | -0.236 | 1.361 | 0.149 | -0.167 | 0.827 | 0 | -1 | 0 | 0 | 0.050 |
| Icam1 | 0.0132 | -0.092 | 1.880 | 0.003 | -0.241 | 2.471 | 0.118 | -0.167 | 0.928 | 0 | -1 | 0 | 0 | 0.050 |
| Znf24 | 0.0746 | -0.097 | 1.127 | 0.003 | -0.163 | 2.567 | 0.057 | -0.166 | 1.241 | 0 | -1 | 0 | 0 | 0.050 |
| Igf1r | 0.0673 | -0.119 | 1.172 | 0.001 | -0.218 | 2.895 | 0.091 | -0.165 | 1.043 | 0 | -1 | 0 | 0 | 0.050 |
| Adam17 | 0.1303 | -0.080 | 0.885 | 0.013 | -0.221 | 1.895 | 0.122 | -0.165 | 0.914 | 0 | -1 | 0 | 0 | 0.050 |
| Nlgn4 | 0.2092 | -0.093 | 0.679 | 0.003 | -0.220 | 2.566 | 0.148 | -0.165 | 0.828 | 0 | -1 | 0 | 0 | 0.050 |
| Impa1 | 0.2253 | -0.053 | 0.647 | 0.016 | -0.153 | 1.785 | 0.061 | -0.164 | 1.217 | 0 | -1 | 0 | 0 | 0.050 |
| Smpdl3b | 0.1666 | -0.136 | 0.778 | 0.014 | -0.201 | 1.844 | 0.055 | -0.164 | 1.263 | 0 | -1 | 0 | 0 | 0.050 |
| Slc44a2 | 0.3407 | -0.070 | 0.468 | 0.008 | -0.187 | 2.115 | 0.107 | -0.162 | 0.972 | 0 | -1 | 0 | 0 | 0.050 |
| Npr3 | 0.1494 | -0.052 | 0.826 | 0.028 | -0.143 | 1.559 | 0.052 | -0.162 | 1.288 | 0 | -1 | 0 | 0 | 0.050 |
| Tenm1 | 0.2411 | -0.051 | 0.618 | 0.007 | -0.213 | 2.160 | 0.142 | -0.161 | 0.848 | 0 | -1 | 0 | 0 | 0.050 |
| Sema3a | 0.9591 | 0.003 | 0.018 | 0.024 | -0.298 | 1.616 | 0.111 | -0.161 | 0.953 | 0 | -1 | 0 | 0 | 0.050 |
| Grb2 | 0.0131 | -0.091 | 1.883 | 0.035 | -0.192 | 1.457 | 0.083 | -0.160 | 1.081 | 0 | -1 | 0 | 0 | 0.050 |
| Acat2 | 0.7527 | -0.022 | 0.123 | 0.002 | -0.218 | 2.740 | 0.053 | -0.160 | 1.279 | 0 | -1 | 0 | 0 | 0.050 |
| Itgav | 0.8798 | 0.005 | 0.056 | 0.008 | -0.237 | 2.090 | 0.082 | -0.159 | 1.088 | 0 | -1 | 0 | 0 | 0.050 |
| Cdh11 | 0.0532 | -0.073 | 1.274 | 0.021 | -0.176 | 1.678 | 0.225 | -0.159 | 0.648 | 0 | -1 | 0 | 0 | 0.050 |
| Mcam | 0.0639 | -0.091 | 1.195 | 0.002 | -0.185 | 2.810 | 0.215 | -0.159 | 0.668 | 0 | -1 | 0 | 0 | 0.050 |
| Slc3a2 | 0.1409 | -0.114 | 0.851 | 0.033 | -0.213 | 1.483 | 0.110 | -0.158 | 0.958 | 0 | -1 | 0 | 0 | 0.050 |
| Cr1l | 0.0328 | -0.128 | 1.484 | 0.033 | -0.188 | 1.483 | 0.054 | -0.158 | 1.264 | 0 | -1 | 0 | 0 | 0.050 |
| Ass1 | 0.5913 | -0.033 | 0.228 | 0.009 | -0.182 | 2.040 | 0.056 | -0.156 | 1.252 | 0 | -1 | 0 | 0 | 0.050 |
| Mertk | 0.0757 | -0.053 | 1.121 | 0.017 | -0.198 | 1.768 | 0.057 | -0.156 | 1.247 | 0 | -1 | 0 | 0 | 0.050 |
| Nr2c2 | 0.8454 | -0.017 | 0.073 | 0.040 | -0.187 | 1.403 | 0.075 | -0.154 | 1.128 | 0 | -1 | 0 | 0 | 0.050 |
| Tpbg | 0.0874 | -0.161 | 1.059 | 0.004 | -0.199 | 2.411 | 0.220 | -0.154 | 0.657 | 0 | -1 | 0 | 0 | 0.050 |
| Myorg | 0.0099 | -0.106 | 2.005 | 0.014 | -0.202 | 1.846 | 0.138 | -0.153 | 0.859 | 0 | -1 | 0 | 0 | 0.050 |
| Abca7 | 0.0498 | -0.101 | 1.302 | 0.003 | -0.232 | 2.487 | 0.053 | -0.153 | 1.277 | 0 | -1 | 0 | 0 | 0.050 |
| Kdm2b | 0.6500 | -0.093 | 0.187 | 0.008 | -0.179 | 2.115 | 0.091 | -0.152 | 1.039 | 0 | -1 | 0 | 0 | 0.050 |
| Tmem67 | 0.1311 | -0.117 | 0.883 | 0.020 | -0.154 | 1.704 | 0.080 | -0.152 | 1.097 | 0 | -1 | 0 | 0 | 0.050 |
| Cdh18 | 0.4665 | -0.034 | 0.331 | 0.015 | -0.181 | 1.829 | 0.172 | -0.152 | 0.764 | 0 | -1 | 0 | 0 | 0.050 |

| | | | | | | | | | | | | | |
|---|---|---|---|---|---|---|---|---|---|---|---|---|---|
| Cggbp1 | 0.7130 | -0.025 | 0.147 | 0.015 | -0.150 | 1.822 | 0.062 | -0.152 | 1.209 | 0 | -1 | 0 | 0 | 0.050 |
| Cd47 | 0.1245 | -0.078 | 0.905 | 0.036 | -0.211 | 1.442 | 0.137 | -0.151 | 0.862 | 0 | -1 | 0 | 0 | 0.050 |
| Zfp619 | 0.2742 | -0.129 | 0.562 | 0.048 | -0.259 | 1.319 | 0.349 | -0.150 | 0.458 | 0 | -1 | 0 | 0 | 0.050 |
| Tacstd2 | 0.1010 | -0.132 | 0.996 | 0.011 | -0.140 | 1.941 | 0.101 | -0.149 | 0.996 | 0 | -1 | 0 | 0 | 0.050 |
| Adam10 | 0.0564 | -0.123 | 1.249 | 0.011 | -0.214 | 1.963 | 0.098 | -0.149 | 1.007 | 0 | -1 | 0 | 0 | 0.050 |
| Scube3 | 0.5098 | -0.061 | 0.293 | 0.028 | -0.273 | 1.551 | 0.260 | -0.149 | 0.585 | 0 | -1 | 0 | 0 | 0.050 |
| Nectin1 | 0.0094 | -0.126 | 2.026 | 0.003 | -0.184 | 2.480 | 0.196 | -0.149 | 0.707 | 0 | -1 | 0 | 0 | 0.050 |
| Ncam1 | 0.1139 | -0.113 | 0.944 | 0.004 | -0.214 | 2.425 | 0.140 | -0.148 | 0.854 | 0 | -1 | 0 | 0 | 0.050 |
| Adgra3 | 0.2844 | -0.088 | 0.546 | 0.017 | -0.181 | 1.760 | 0.100 | -0.147 | 0.999 | 0 | -1 | 0 | 0 | 0.050 |
| Slc39a14 | 0.2875 | -0.087 | 0.541 | 0.042 | -0.143 | 1.380 | 0.100 | -0.145 | 0.998 | 0 | -1 | 0 | 0 | 0.050 |
| Anks1b | 0.0714 | -0.080 | 1.146 | 0.010 | -0.143 | 2.022 | 0.097 | -0.145 | 1.015 | 0 | -1 | 0 | 0 | 0.050 |
| Gapdh | 0.5313 | -0.043 | 0.275 | 0.025 | -0.452 | 1.601 | 0.588 | -0.144 | 0.231 | 0 | -1 | 0 | 0 | 0.050 |
| Loxl3 | 0.4991 | 0.014 | 0.302 | 0.004 | -0.154 | 2.388 | 0.201 | -0.143 | 0.697 | 0 | -1 | 0 | 0 | 0.050 |
| Gpc1 | 0.0060 | -0.120 | 2.225 | 0.009 | -0.199 | 2.061 | 0.055 | -0.143 | 1.257 | 0 | -1 | 0 | 0 | 0.050 |
| Entpd3 | 0.3155 | -0.116 | 0.501 | 0.015 | -0.275 | 1.824 | 0.144 | -0.143 | 0.843 | 0 | -1 | 0 | 0 | 0.050 |
| SIRP-beta | 0.1226 | -0.119 | 0.912 | 0.023 | -0.222 | 1.633 | 0.134 | -0.142 | 0.873 | 0 | -1 | 0 | 0 | 0.050 |
| Cdh2 | 0.0415 | -0.127 | 1.382 | 0.043 | -0.157 | 1.369 | 0.097 | -0.141 | 1.014 | 0 | -1 | 0 | 0 | 0.050 |
| Ralgapa2 | 0.0608 | -0.087 | 1.216 | 0.023 | -0.305 | 1.629 | 0.156 | -0.141 | 0.808 | 0 | -1 | 0 | 0 | 0.050 |
| Epha1 | 0.1093 | -0.122 | 0.962 | 0.002 | -0.203 | 2.721 | 0.194 | -0.140 | 0.713 | 0 | -1 | 0 | 0 | 0.050 |
| Plxna3 | 0.1785 | -0.101 | 0.748 | 0.012 | -0.168 | 1.920 | 0.144 | -0.140 | 0.842 | 0 | -1 | 0 | 0 | 0.050 |
| Tmem59 | 0.1684 | -0.093 | 0.774 | 0.007 | -0.197 | 2.141 | 0.382 | -0.139 | 0.418 | 0 | -1 | 0 | 0 | 0.050 |
| Plxnd1 | 0.0373 | -0.106 | 1.428 | 0.021 | -0.180 | 1.672 | 0.169 | -0.138 | 0.773 | 0 | -1 | 0 | 0 | 0.050 |
| Itgb8 | 0.2252 | -0.070 | 0.647 | 0.012 | -0.339 | 1.926 | 0.052 | -0.137 | 1.284 | 0 | -1 | 0 | 0 | 0.050 |
| F630007L15R | 0.4407 | -0.082 | 0.356 | 0.006 | -0.162 | 2.258 | 0.153 | -0.136 | 0.817 | 0 | -1 | 0 | 0 | 0.050 |
| Brip1os | 0.3969 | -0.063 | 0.401 | 0.010 | -0.164 | 1.992 | 0.178 | -0.135 | 0.749 | 0 | -1 | 0 | 0 | 0.050 |
| Mpp7 | 0.0148 | -0.069 | 1.830 | 0.018 | -0.138 | 1.736 | 0.056 | -0.135 | 1.252 | 0 | -1 | 0 | 0 | 0.050 |
| Gpx8 | 0.0200 | -0.131 | 1.699 | 0.022 | -0.160 | 1.651 | 0.003 | -0.134 | 2.539 | 0 | -1 | 0 | 0 | 0.050 |
| Fmnl1 | 0.3642 | -0.046 | 0.439 | 0.007 | -0.189 | 2.129 | 0.059 | -0.133 | 1.231 | 0 | -1 | 0 | 0 | 0.050 |
| Prxl2c | 0.3412 | -0.068 | 0.467 | 0.047 | -0.173 | 1.330 | 0.052 | -0.133 | 1.283 | 0 | -1 | 0 | 0 | 0.050 |
| Itgb3 | 0.1594 | -0.051 | 0.797 | 0.005 | -0.212 | 2.272 | 0.098 | -0.133 | 1.009 | 0 | -1 | 0 | 0 | 0.050 |
| Enc1 | 0.4714 | -0.062 | 0.327 | 0.007 | -0.241 | 2.154 | 0.047 | -0.133 | 1.329 | 0 | -1 | 0 | 0 | 0.050 |
| Ppic | 0.9183 | -0.006 | 0.037 | 0.001 | -0.197 | 2.958 | 0.143 | -0.131 | 0.844 | 0 | -1 | 0 | 0 | 0.050 |
| Mapkapk2 | 0.3608 | 0.036 | 0.443 | 0.009 | -0.140 | 2.048 | 0.043 | -0.131 | 1.369 | 0 | -1 | 0 | 0 | 0.050 |
| Tapbpl | 0.6161 | 0.055 | 0.210 | 0.037 | -0.160 | 1.429 | 0.093 | -0.130 | 1.029 | 0 | -1 | 0 | 0 | 0.050 |
| Itga3 | 0.2075 | -0.090 | 0.683 | 0.017 | -0.195 | 1.758 | 0.015 | -0.130 | 1.827 | 0 | -1 | 0 | 0 | 0.050 |
| Rlim | 0.0201 | -0.081 | 1.696 | 0.029 | -0.157 | 1.539 | 0.168 | -0.130 | 0.775 | 0 | -1 | 0 | 0 | 0.050 |
| Acp2 | 0.5746 | 0.035 | 0.241 | 0.001 | -0.270 | 3.051 | 0.183 | -0.128 | 0.737 | 0 | -1 | 0 | 0 | 0.050 |
| Cadm4 | 0.4659 | -0.024 | 0.332 | 0.002 | -0.205 | 2.740 | 0.095 | -0.128 | 1.023 | 0 | -1 | 0 | 0 | 0.050 |
| Fam102b | 0.3180 | -0.088 | 0.498 | 0.042 | -0.179 | 1.378 | 0.140 | -0.127 | 0.852 | 0 | -1 | 0 | 0 | 0.050 |
| Pfdn1 | 0.3731 | -0.053 | 0.428 | 0.048 | -0.218 | 1.322 | 0.602 | -0.127 | 0.221 | 0 | -1 | 0 | 0 | 0.050 |
| Cacna2d1 | 0.3653 | -0.040 | 0.437 | 0.039 | -0.225 | 1.410 | 0.236 | -0.125 | 0.627 | 0 | -1 | 0 | 0 | 0.050 |
| Aldh1a1 | 0.2899 | -0.086 | 0.538 | 0.010 | -0.178 | 2.002 | 0.081 | -0.124 | 1.090 | 0 | -1 | 0 | 0 | 0.050 |
| Zfp263 | 0.8217 | 0.023 | 0.085 | 0.006 | -0.261 | 2.194 | 0.333 | -0.124 | 0.477 | 0 | -1 | 0 | 0 | 0.050 |
| Mettl17 | 0.3194 | -0.067 | 0.496 | 0.001 | -0.204 | 3.095 | 0.120 | -0.124 | 0.922 | 0 | -1 | 0 | 0 | 0.050 |
| Slc16a1 | 0.1003 | -0.215 | 0.999 | 0.022 | -0.161 | 1.663 | 0.066 | -0.123 | 1.183 | 0 | -1 | 0 | 0 | 0.050 |
| Plxnb1 | 0.3124 | -0.043 | 0.505 | 0.022 | -0.152 | 1.657 | 0.106 | -0.123 | 0.975 | 0 | -1 | 0 | 0 | 0.050 |
| Fam171a2 | 0.1067 | -0.127 | 0.972 | 0.042 | -0.140 | 1.375 | 0.087 | -0.122 | 1.058 | 0 | -1 | 0 | 0 | 0.050 |
| Psat1 | 0.1854 | -0.047 | 0.732 | 0.014 | -0.140 | 1.850 | 0.019 | -0.118 | 1.721 | 0 | -1 | 0 | 0 | 0.050 |
| Cchcr1 | 0.4007 | -0.101 | 0.397 | 0.044 | -0.185 | 1.355 | 0.343 | -0.118 | 0.464 | 0 | -1 | 0 | 0 | 0.050 |
| Znf513 | 0.4215 | 0.034 | 0.375 | 0.032 | -0.184 | 1.501 | 0.569 | -0.117 | 0.245 | 0 | -1 | 0 | 0 | 0.050 |
| Gmcl1 | 0.1230 | -0.313 | 0.910 | 0.009 | -0.227 | 2.061 | 0.369 | -0.116 | 0.433 | 0 | -1 | 0 | 0 | 0.050 |
| Sdf4 | 0.2580 | -0.108 | 0.588 | 0.041 | -0.185 | 1.390 | 0.126 | -0.115 | 0.901 | 0 | -1 | 0 | 0 | 0.050 |
| Col3a1 | 0.0537 | -0.131 | 1.270 | 0.030 | -0.189 | 1.522 | 0.142 | -0.115 | 0.848 | 0 | -1 | 0 | 0 | 0.050 |
| Cpd | 0.8896 | -0.008 | 0.051 | 0.008 | -0.156 | 2.085 | 0.136 | -0.115 | 0.867 | 0 | -1 | 0 | 0 | 0.050 |
| Romo1 | 0.0570 | 0.155 | 1.244 | 0.005 | -0.216 | 2.320 | 0.276 | -0.114 | 0.558 | 0 | -1 | 0 | 0 | 0.050 |
| Aldh1l2 | 0.9404 | 0.003 | 0.027 | 0.011 | -0.161 | 1.972 | 0.034 | -0.113 | 1.470 | 0 | -1 | 0 | 0 | 0.050 |
| Sall2 | 0.0559 | -0.174 | 1.253 | 0.002 | -0.184 | 2.792 | 0.148 | -0.112 | 0.829 | 0 | -1 | 0 | 0 | 0.050 |
| Spin1 | 0.7181 | -0.020 | 0.144 | 0.014 | -0.174 | 1.869 | 0.163 | -0.111 | 0.787 | 0 | -1 | 0 | 0 | 0.050 |
| Dcaf11 | 0.0381 | -0.061 | 1.419 | 0.012 | -0.147 | 1.927 | 0.016 | -0.110 | 1.794 | 0 | -1 | 0 | 0 | 0.050 |
| Pdpn | 0.9487 | 0.014 | 0.023 | 0.041 | -0.275 | 1.385 | 0.020 | -0.110 | 1.689 | 0 | -1 | 0 | 0 | 0.050 |
| Bst2 | 0.1324 | -0.211 | 0.878 | 0.000 | -0.295 | 3.702 | 0.195 | -0.109 | 0.710 | 0 | -1 | 0 | 0 | 0.050 |
| Fbxw8 | 0.0866 | -0.090 | 1.062 | 0.008 | -0.140 | 2.074 | 0.019 | -0.108 | 1.728 | 0 | -1 | 0 | 0 | 0.050 |
| Actr10 | 0.0506 | -0.108 | 1.296 | 0.014 | -0.189 | 1.856 | 0.050 | -0.108 | 1.300 | 0 | -1 | 0 | 0 | 0.050 |
| Bace1 | 0.7780 | -0.032 | 0.109 | 0.007 | -0.180 | 2.165 | 0.158 | -0.107 | 0.800 | 0 | -1 | 0 | 0 | 0.050 |
| Sco2 | 0.2715 | -0.054 | 0.566 | 0.031 | -0.196 | 1.514 | 0.140 | -0.106 | 0.852 | 0 | -1 | 0 | 0 | 0.050 |
| Reck | 0.6658 | 0.046 | 0.177 | 0.001 | -0.237 | 2.890 | 0.404 | -0.106 | 0.394 | 0 | -1 | 0 | 0 | 0.050 |
| Bmp8b | 0.0689 | -0.118 | 1.162 | 0.008 | -0.138 | 2.084 | 0.077 | -0.104 | 1.115 | 0 | -1 | 0 | 0 | 0.050 |
| Wnt5a | 0.1969 | -0.082 | 0.706 | 0.032 | -0.139 | 1.499 | 0.161 | -0.103 | 0.793 | 0 | -1 | 0 | 0 | 0.050 |
| Cpm | 0.5262 | 0.036 | 0.279 | 0.002 | -0.207 | 2.614 | 0.254 | -0.102 | 0.595 | 0 | -1 | 0 | 0 | 0.050 |
| Iba57 | 0.9916 | 0.001 | 0.004 | 0.016 | -0.160 | 1.793 | 0.060 | -0.102 | 1.221 | 0 | -1 | 0 | 0 | 0.050 |
| Rxra | 0.1763 | -0.107 | 0.754 | 0.022 | -0.246 | 1.664 | 0.108 | -0.101 | 0.967 | 0 | -1 | 0 | 0 | 0.050 |

| | | | | | | | | | | | | | |
|---|---|---|---|---|---|---|---|---|---|---|---|---|---|
| Gle1 | 0.4694 | -0.046 | 0.328 | 0.004 | -0.211 | 2.391 | 0.282 | -0.101 | 0.549 | 0 | -1 | 0 | 0 | 0.050 |
| P4ha2 | 0.3047 | 0.067 | 0.516 | 0.046 | -0.197 | 1.334 | 0.236 | -0.100 | 0.628 | 0 | -1 | 0 | 0 | 0.050 |
| Glt8d1 | 0.9917 | -0.001 | 0.004 | 0.022 | -0.152 | 1.660 | 0.082 | -0.100 | 1.084 | 0 | -1 | 0 | 0 | 0.050 |
| Kmt2c | 0.0599 | -0.194 | 1.223 | 0.014 | -0.238 | 1.863 | 0.363 | -0.100 | 0.441 | 0 | -1 | 0 | 0 | 0.050 |
| Zadh2 | 0.5990 | -0.034 | 0.223 | 0.021 | -0.146 | 1.674 | 0.241 | -0.099 | 0.617 | 0 | -1 | 0 | 0 | 0.050 |
| Zfp513 | 0.5916 | 0.104 | 0.228 | 0.000 | -0.326 | 3.674 | 0.035 | -0.098 | 1.454 | 0 | -1 | 0 | 0 | 0.050 |
| Lancl1 | 0.3456 | -0.063 | 0.461 | 0.003 | -0.195 | 2.535 | 0.043 | -0.098 | 1.362 | 0 | -1 | 0 | 0 | 0.050 |
| Adamtsl3 | 0.3374 | 0.048 | 0.472 | 0.018 | -0.198 | 1.743 | 0.383 | -0.095 | 0.417 | 0 | -1 | 0 | 0 | 0.050 |
| Pdap1 | 0.2794 | -0.062 | 0.554 | 0.027 | -0.143 | 1.570 | 0.351 | -0.094 | 0.455 | 0 | -1 | 0 | 0 | 0.050 |
| Mccc2 | 0.8964 | 0.007 | 0.047 | 0.043 | -0.145 | 1.368 | 0.411 | -0.093 | 0.386 | 0 | -1 | 0 | 0 | 0.050 |
| Lipt1 | 0.7214 | -0.032 | 0.142 | 0.041 | -0.248 | 1.390 | 0.011 | -0.090 | 1.976 | 0 | -1 | 0 | 0 | 0.050 |
| Adgrl1 | 0.0620 | -0.064 | 1.208 | 0.013 | -0.165 | 1.879 | 0.178 | -0.087 | 0.751 | 0 | -1 | 0 | 0 | 0.050 |
| Cdk5rap3 | 0.0488 | -0.059 | 1.312 | 0.003 | -0.191 | 2.597 | 0.129 | -0.087 | 0.890 | 0 | -1 | 0 | 0 | 0.050 |
| Nt5e | 0.3494 | -0.045 | 0.457 | 0.014 | -0.245 | 1.840 | 0.142 | -0.085 | 0.846 | 0 | -1 | 0 | 0 | 0.050 |
| Bak1 | 0.1923 | -0.126 | 0.716 | 0.046 | -0.184 | 1.333 | 0.327 | -0.085 | 0.485 | 0 | -1 | 0 | 0 | 0.050 |
| Nlgn2 | 0.3530 | -0.023 | 0.452 | 0.005 | -0.152 | 2.268 | 0.392 | -0.084 | 0.407 | 0 | -1 | 0 | 0 | 0.050 |
| Usp37 | 0.2100 | -0.105 | 0.678 | 0.013 | -0.140 | 1.880 | 0.332 | -0.083 | 0.479 | 0 | -1 | 0 | 0 | 0.050 |
| Chrne | 0.0502 | -0.132 | 1.300 | 0.048 | -0.236 | 1.316 | 0.237 | -0.083 | 0.624 | 0 | -1 | 0 | 0 | 0.050 |
| Ech1 | 0.2411 | 0.152 | 0.618 | 0.028 | -0.211 | 1.549 | 0.207 | -0.081 | 0.684 | 0 | -1 | 0 | 0 | 0.050 |
| Tmem106a | 0.9313 | 0.010 | 0.031 | 0.016 | -0.141 | 1.801 | 0.422 | -0.079 | 0.374 | 0 | -1 | 0 | 0 | 0.050 |
| Heca | 0.7490 | -0.039 | 0.126 | 0.003 | -0.207 | 2.528 | 0.186 | -0.074 | 0.730 | 0 | -1 | 0 | 0 | 0.050 |
| Glg1 | 0.0043 | -0.132 | 2.362 | 0.003 | -0.152 | 2.484 | 0.217 | -0.073 | 0.663 | 0 | -1 | 0 | 0 | 0.050 |
| Dlc1 | 0.8955 | -0.010 | 0.048 | 0.012 | -0.182 | 1.925 | 0.166 | -0.071 | 0.779 | 0 | -1 | 0 | 0 | 0.050 |
| Ppm1d | 0.6294 | -0.018 | 0.201 | 0.048 | -0.156 | 1.323 | 0.474 | -0.070 | 0.325 | 0 | -1 | 0 | 0 | 0.050 |
| Psmb1 | 0.1812 | 0.136 | 0.742 | 0.009 | -0.147 | 2.051 | 0.137 | -0.069 | 0.865 | 0 | -1 | 0 | 0 | 0.050 |
| Cx3cl1 | 0.0703 | -0.193 | 1.153 | 0.017 | -0.181 | 1.760 | 0.301 | -0.068 | 0.522 | 0 | -1 | 0 | 0 | 0.050 |
| Sdhd | 0.7480 | 0.071 | 0.126 | 0.032 | -0.261 | 1.492 | 0.727 | -0.065 | 0.138 | 0 | -1 | 0 | 0 | 0.050 |
| Tbl1xr1 | 0.5313 | 0.052 | 0.275 | 0.006 | -0.192 | 2.201 | 0.487 | -0.063 | 0.312 | 0 | -1 | 0 | 0 | 0.050 |
| Tmem168 | 0.1138 | -0.134 | 0.944 | 0.004 | -0.163 | 2.385 | 0.210 | -0.063 | 0.677 | 0 | -1 | 0 | 0 | 0.050 |
| Npr1 | 0.0120 | -0.108 | 1.920 | 0.013 | -0.146 | 1.901 | 0.623 | -0.061 | 0.206 | 0 | -1 | 0 | 0 | 0.050 |
| Mfap3l | 0.1683 | -0.075 | 0.774 | 0.002 | -0.158 | 2.603 | 0.461 | -0.060 | 0.336 | 0 | -1 | 0 | 0 | 0.050 |
| Raet1a | 0.3515 | -0.173 | 0.454 | 0.004 | -0.188 | 2.411 | 0.545 | -0.060 | 0.264 | 0 | -1 | 0 | 0 | 0.050 |
| Gpaa1 | 0.0508 | -0.066 | 1.294 | 0.012 | -0.145 | 1.939 | 0.206 | -0.059 | 0.687 | 0 | -1 | 0 | 0 | 0.050 |
| Dync2i1 | 0.5425 | 0.023 | 0.266 | 0.031 | -0.152 | 1.506 | 0.350 | -0.059 | 0.456 | 0 | -1 | 0 | 0 | 0.050 |
| P4ha1 | 0.4396 | 0.087 | 0.357 | 0.009 | -0.204 | 2.023 | 0.188 | -0.059 | 0.725 | 0 | -1 | 0 | 0 | 0.050 |
| Cd9 | 0.0981 | -0.130 | 1.008 | 0.022 | -0.191 | 1.657 | 0.636 | -0.058 | 0.196 | 0 | -1 | 0 | 0 | 0.050 |
| Fbxl13 | 0.3991 | -0.191 | 0.399 | 0.003 | -0.149 | 2.593 | 0.503 | -0.058 | 0.298 | 0 | -1 | 0 | 0 | 0.050 |
| Idi1 | 0.3004 | -0.090 | 0.522 | 0.041 | -0.142 | 1.391 | 0.633 | -0.058 | 0.199 | 0 | -1 | 0 | 0 | 0.050 |
| Psma3 | 0.1159 | 0.062 | 0.936 | 0.019 | -0.146 | 1.724 | 0.334 | -0.058 | 0.476 | 0 | -1 | 0 | 0 | 0.050 |
| Jup | 0.0908 | -0.092 | 1.042 | 0.003 | -0.229 | 2.522 | 0.646 | -0.056 | 0.190 | 0 | -1 | 0 | 0 | 0.050 |
| Dynlt1 | 0.9285 | 0.006 | 0.032 | 0.019 | -0.139 | 1.722 | 0.140 | -0.054 | 0.853 | 0 | -1 | 0 | 0 | 0.050 |
| Nceh1 | 0.0631 | -0.189 | 1.200 | 0.004 | -0.237 | 2.416 | 0.555 | -0.054 | 0.255 | 0 | -1 | 0 | 0 | 0.050 |
| Pdhb | 0.1724 | 0.133 | 0.764 | 0.005 | -0.166 | 2.343 | 0.539 | -0.052 | 0.268 | 0 | -1 | 0 | 0 | 0.050 |
| Tbc1d13 | 0.9406 | -0.004 | 0.027 | 0.035 | -0.184 | 1.461 | 0.419 | -0.050 | 0.378 | 0 | -1 | 0 | 0 | 0.050 |
| Atp1b2 | 0.7465 | -0.029 | 0.127 | 0.047 | -0.177 | 1.325 | 0.611 | -0.050 | 0.214 | 0 | -1 | 0 | 0 | 0.050 |
| Pxdn | 0.6155 | 0.027 | 0.211 | 0.009 | -0.140 | 2.058 | 0.261 | -0.050 | 0.583 | 0 | -1 | 0 | 0 | 0.050 |
| Mfap3 | 0.5362 | -0.030 | 0.271 | 0.021 | -0.144 | 1.683 | 0.654 | -0.049 | 0.184 | 0 | -1 | 0 | 0 | 0.050 |
| Myoc | 0.0735 | -0.072 | 1.134 | 0.020 | -0.213 | 1.705 | 0.365 | -0.048 | 0.438 | 0 | -1 | 0 | 0 | 0.050 |
| Hmgb3 | 0.0957 | -0.102 | 1.019 | 0.034 | -0.217 | 1.474 | 0.666 | -0.047 | 0.177 | 0 | -1 | 0 | 0 | 0.050 |
| Slc16a3 | 0.8227 | -0.019 | 0.085 | 0.040 | -0.158 | 1.398 | 0.702 | -0.044 | 0.153 | 0 | -1 | 0 | 0 | 0.050 |
| Pde8a | 0.4073 | -0.041 | 0.390 | 0.010 | -0.193 | 1.983 | 0.321 | -0.040 | 0.494 | 0 | -1 | 0 | 0 | 0.050 |
| Acadm | 0.7856 | 0.017 | 0.105 | 0.007 | -0.162 | 2.140 | 0.444 | -0.038 | 0.353 | 0 | -1 | 0 | 0 | 0.050 |
| Samd8 | 0.0665 | -0.079 | 1.177 | 0.014 | -0.146 | 1.859 | 0.498 | -0.037 | 0.303 | 0 | -1 | 0 | 0 | 0.050 |
| Foxp1 | 0.3321 | 0.097 | 0.479 | 0.050 | -0.177 | 1.305 | 0.855 | -0.035 | 0.068 | 0 | -1 | 0 | 0 | 0.050 |
| Sephs1 | 0.6228 | -0.039 | 0.206 | 0.009 | -0.180 | 2.050 | 0.279 | -0.033 | 0.554 | 0 | -1 | 0 | 0 | 0.050 |
| Dstyk | 0.1602 | -0.054 | 0.795 | 0.008 | -0.141 | 2.121 | 0.499 | -0.032 | 0.302 | 0 | -1 | 0 | 0 | 0.050 |
| Psmb2 | 0.0953 | -0.085 | 1.021 | 0.005 | -0.175 | 2.261 | 0.725 | -0.027 | 0.140 | 0 | -1 | 0 | 0 | 0.050 |
| Acads | 0.1128 | -0.130 | 0.948 | 0.005 | -0.157 | 2.283 | 0.453 | -0.021 | 0.344 | 0 | -1 | 0 | 0 | 0.050 |
| Dus2 | 0.1287 | -0.552 | 0.890 | 0.007 | -0.197 | 2.154 | 0.752 | -0.019 | 0.124 | 0 | -1 | 0 | 0 | 0.050 |
| Cdk19 | 0.2603 | 0.076 | 0.585 | 0.026 | -0.150 | 1.593 | 0.868 | -0.011 | 0.062 | 0 | -1 | 0 | 0 | 0.050 |
| Tkt | 0.1003 | 0.047 | 0.999 | 0.006 | -0.147 | 2.218 | 0.850 | -0.009 | 0.071 | 0 | -1 | 0 | 0 | 0.050 |
| Rfx1 | 0.0982 | 0.077 | 1.008 | 0.016 | -0.157 | 1.792 | 0.946 | -0.004 | 0.024 | 0 | -1 | 0 | 0 | 0.050 |
| Ppif | 0.0433 | -0.064 | 1.364 | 0.016 | -0.189 | 1.790 | 0.962 | 0.004 | 0.017 | 0 | -1 | 0 | 0 | 0.050 |
| Ccdc88b | 0.8457 | -0.045 | 0.073 | 0.047 | -0.142 | 1.329 | 0.946 | 0.007 | 0.024 | 0 | -1 | 0 | 0 | 0.050 |
| Kiaa0100 | 0.6909 | -0.041 | 0.161 | 0.039 | -0.199 | 1.414 | 0.910 | 0.016 | 0.041 | 0 | -1 | 0 | 0 | 0.050 |
| Cmtm3 | 0.6100 | -0.091 | 0.215 | 0.012 | -0.241 | 1.926 | 0.657 | 0.017 | 0.182 | 0 | -1 | 0 | 0 | 0.050 |
| Elf4 | 0.7025 | -0.042 | 0.153 | 0.024 | -0.151 | 1.620 | 0.888 | 0.019 | 0.051 | 0 | -1 | 0 | 0 | 0.050 |
| Dalrd3 | 0.4325 | 0.119 | 0.364 | 0.014 | -0.219 | 1.868 | 0.702 | 0.028 | 0.154 | 0 | -1 | 0 | 0 | 0.050 |
| Prpf38a | 0.0439 | 0.088 | 1.358 | 0.015 | -0.144 | 1.838 | 0.788 | 0.029 | 0.103 | 0 | -1 | 0 | 0 | 0.050 |
| Dapk2 | 0.0339 | -0.073 | 1.470 | 0.032 | -0.153 | 1.501 | 0.390 | 0.030 | 0.409 | 0 | -1 | 0 | 0 | 0.050 |
| B9d1 | 0.5724 | -0.030 | 0.242 | 0.017 | -0.167 | 1.775 | 0.504 | 0.035 | 0.297 | 0 | -1 | 0 | 0 | 0.050 |

| | | | | | | | | | | | | | |
|---|---|---|---|---|---|---|---|---|---|---|---|---|---|
| Tssc4 | 0.0889 | -0.095 | 1.051 | 0.042 | -0.138 | 1.372 | 0.271 | 0.038 | 0.567 | 0 | -1 | 0 | 0 | 0.050 |
| Dnajc5 | 0.3038 | 0.069 | 0.517 | 0.021 | -0.138 | 1.668 | 0.166 | 0.088 | 0.779 | 0 | -1 | 0 | 0 | 0.050 |
| Fv4 | 0.9744 | -0.003 | 0.011 | 0.008 | -0.358 | 2.119 | 0.586 | 0.132 | 0.232 | 0 | -1 | 0 | 0 | 0.050 |
| Cnot9 | 0.1334 | -0.087 | 0.875 | 0.028 | -0.157 | 1.549 | 0.218 | 0.146 | 0.661 | 0 | -1 | 0 | 0 | 0.050 |
| Fundc2 | 0.8216 | 0.033 | 0.085 | 0.007 | -0.167 | 2.128 | 0.004 | 0.186 | 2.400 | 0 | -1 | 1 | 0 | 0.050 |
| Fosl1 | 0.1569 | -0.266 | 0.804 | 0.539 | -0.116 | 0.269 | 0.007 | -0.473 | 2.153 | 0 | 0 | -1 | 0 | 0.050 |
| Txnl1 | 0.0110 | -0.087 | 1.959 | 0.285 | -0.148 | 0.545 | 0.003 | -0.456 | 2.551 | 0 | 0 | -1 | 0 | 0.050 |
| Fzd5 | 0.0684 | -0.263 | 1.165 | 0.073 | -0.316 | 1.135 | 0.009 | -0.382 | 2.037 | 0 | 0 | -1 | 0 | 0.050 |
| Guf1 | 0.1460 | -0.279 | 0.836 | 0.383 | -0.183 | 0.417 | 0.043 | -0.374 | 1.371 | 0 | 0 | -1 | 0 | 0.050 |
| Nlrc3 | 0.0691 | -0.577 | 1.160 | 0.097 | -0.770 | 1.013 | 0.001 | -0.368 | 2.981 | 0 | 0 | -1 | 0 | 0.050 |
| Avpi1 | 0.2096 | -0.184 | 0.679 | 0.097 | -0.232 | 1.013 | 0.004 | -0.347 | 2.373 | 0 | 0 | -1 | 0 | 0.050 |
| Fkbp2 | 0.0161 | -0.119 | 1.793 | 0.243 | -0.135 | 0.614 | 0.000 | -0.347 | 3.777 | 0 | 0 | -1 | 0 | 0.050 |
| Zfp451 | 0.7471 | -0.062 | 0.127 | 0.644 | -0.099 | 0.191 | 0.002 | -0.332 | 2.728 | 0 | 0 | -1 | 0 | 0.050 |
| Trim35 | 0.1704 | -0.182 | 0.769 | 0.108 | -0.220 | 0.967 | 0.026 | -0.331 | 1.586 | 0 | 0 | -1 | 0 | 0.050 |
| Ece2 | 0.9915 | 0.001 | 0.004 | 0.078 | -0.170 | 1.110 | 0.001 | -0.331 | 3.017 | 0 | 0 | -1 | 0 | 0.050 |
| Ptov1 | 0.0070 | -0.101 | 2.155 | 0.073 | -0.310 | 1.135 | 0.004 | -0.328 | 2.379 | 0 | 0 | -1 | 0 | 0.050 |
| Arrb1 | 0.8383 | 0.025 | 0.077 | 0.169 | -0.204 | 0.771 | 0.016 | -0.304 | 1.792 | 0 | 0 | -1 | 0 | 0.050 |
| Cxxc5 | 0.8008 | 0.028 | 0.096 | 0.069 | -0.196 | 1.163 | 0.009 | -0.298 | 2.050 | 0 | 0 | -1 | 0 | 0.050 |
| Rpl34 | 0.3869 | -0.051 | 0.412 | 0.480 | -0.126 | 0.319 | 0.014 | -0.296 | 1.852 | 0 | 0 | -1 | 0 | 0.050 |
| Thbs3 | 0.0867 | 0.074 | 1.062 | 0.085 | -0.154 | 1.070 | 0.027 | -0.286 | 1.563 | 0 | 0 | -1 | 0 | 0.050 |
| Nkiras1 | 0.1566 | -0.159 | 0.805 | 0.130 | -0.221 | 0.886 | 0.006 | -0.286 | 2.190 | 0 | 0 | -1 | 0 | 0.050 |
| Trhde | 0.7414 | -0.037 | 0.130 | 0.081 | -0.169 | 1.093 | 0.004 | -0.277 | 2.380 | 0 | 0 | -1 | 0 | 0.050 |
| Klf5 | 0.0144 | -0.124 | 1.842 | 0.083 | -0.205 | 1.083 | 0.001 | -0.275 | 3.270 | 0 | 0 | -1 | 0 | 0.050 |
| Wrap73 | 0.1212 | -0.121 | 0.916 | 0.078 | -0.232 | 1.107 | 0.001 | -0.273 | 2.968 | 0 | 0 | -1 | 0 | 0.050 |
| Irf7 | 0.3852 | -0.127 | 0.414 | 0.316 | -0.109 | 0.500 | 0.026 | -0.270 | 1.589 | 0 | 0 | -1 | 0 | 0.050 |
| Ca13 | 0.1513 | -0.049 | 0.820 | 0.089 | -0.155 | 1.052 | 0.002 | -0.266 | 2.610 | 0 | 0 | -1 | 0 | 0.050 |
| Actl11 | 0.2198 | -0.128 | 0.658 | 0.094 | -0.294 | 1.027 | 0.021 | -0.262 | 1.670 | 0 | 0 | -1 | 0 | 0.050 |
| Txndc9 | 0.0808 | -0.044 | 1.093 | 0.177 | -0.161 | 0.753 | 0.007 | -0.260 | 2.152 | 0 | 0 | -1 | 0 | 0.050 |
| Tex44 | 0.4871 | 0.116 | 0.312 | 0.703 | 0.031 | 0.153 | 0.000 | -0.257 | 4.081 | 0 | 0 | -1 | 0 | 0.050 |
| Lrrc41 | 0.0217 | -0.135 | 1.663 | 0.099 | -0.230 | 1.005 | 0.003 | -0.256 | 2.540 | 0 | 0 | -1 | 0 | 0.050 |
| Mthfr | 0.3683 | -0.039 | 0.434 | 0.596 | -0.060 | 0.224 | 0.000 | -0.256 | 3.582 | 0 | 0 | -1 | 0 | 0.050 |
| Znf668 | 0.0314 | -0.137 | 1.502 | 0.264 | -0.089 | 0.578 | 0.001 | -0.248 | 3.184 | 0 | 0 | -1 | 0 | 0.050 |
| Phf21a | 0.7826 | -0.041 | 0.106 | 0.248 | 0.077 | 0.606 | 0.036 | -0.247 | 1.445 | 0 | 0 | -1 | 0 | 0.050 |
| Znf48 | 0.1212 | -0.059 | 0.917 | 0.130 | -0.129 | 0.885 | 0.019 | -0.246 | 1.729 | 0 | 0 | -1 | 0 | 0.050 |
| Rps16 | 0.0639 | -0.149 | 1.194 | 0.070 | -0.126 | 1.154 | 0.000 | -0.245 | 3.879 | 0 | 0 | -1 | 0 | 0.050 |
| Timp3 | 0.0733 | -0.089 | 1.135 | 0.141 | -0.185 | 0.850 | 0.000 | -0.240 | 3.309 | 0 | 0 | -1 | 0 | 0.050 |
| Lztr1 | 0.1066 | -0.124 | 0.972 | 0.325 | -0.098 | 0.488 | 0.006 | -0.238 | 2.250 | 0 | 0 | -1 | 0 | 0.050 |
| Fam171b | 0.0775 | -0.155 | 1.111 | 0.099 | -0.173 | 1.003 | 0.047 | -0.236 | 1.326 | 0 | 0 | -1 | 0 | 0.050 |
| Rnf6 | 0.0918 | -0.194 | 1.037 | 0.784 | -0.053 | 0.106 | 0.034 | -0.235 | 1.469 | 0 | 0 | -1 | 0 | 0.050 |
| Dkk1 | 0.2732 | -0.110 | 0.563 | 0.282 | -0.145 | 0.550 | 0.045 | -0.230 | 1.344 | 0 | 0 | -1 | 0 | 0.050 |
| Lasp1 | 0.9578 | 0.005 | 0.019 | 0.659 | -0.016 | 0.181 | 0.001 | -0.230 | 2.929 | 0 | 0 | -1 | 0 | 0.050 |
| Xpnpep2 | 0.1470 | -0.110 | 0.833 | 0.311 | -0.121 | 0.507 | 0.000 | -0.229 | 3.722 | 0 | 0 | -1 | 0 | 0.050 |
| A630089N07F | 0.0594 | -0.134 | 1.227 | 0.091 | -0.157 | 1.042 | 0.000 | -0.227 | 3.872 | 0 | 0 | -1 | 0 | 0.050 |
| Vhl | 0.2794 | -0.059 | 0.554 | 0.112 | -0.219 | 0.952 | 0.030 | -0.226 | 1.525 | 0 | 0 | -1 | 0 | 0.050 |
| PcgF3 | 0.0722 | -0.179 | 1.142 | 0.127 | -0.209 | 0.895 | 0.005 | -0.224 | 2.301 | 0 | 0 | -1 | 0 | 0.050 |
| Ube2s | 0.2852 | -0.069 | 0.545 | 0.063 | -0.087 | 1.203 | 0.000 | -0.223 | 3.441 | 0 | 0 | -1 | 0 | 0.050 |
| Sesn1 | 0.0738 | -0.056 | 1.132 | 0.260 | -0.141 | 0.585 | 0.029 | -0.223 | 1.537 | 0 | 0 | -1 | 0 | 0.050 |
| Nmnat1 | 0.7918 | 0.040 | 0.101 | 0.070 | -0.128 | 1.158 | 0.010 | -0.222 | 1.979 | 0 | 0 | -1 | 0 | 0.050 |
| Plpp3 | 0.1396 | -0.132 | 0.855 | 0.060 | -0.130 | 1.221 | 0.012 | -0.220 | 1.906 | 0 | 0 | -1 | 0 | 0.050 |
| Phf5a | 0.0275 | -0.131 | 1.561 | 0.154 | -0.096 | 0.811 | 0.018 | -0.218 | 1.740 | 0 | 0 | -1 | 0 | 0.050 |
| Scarb1 | 0.0358 | -0.110 | 1.446 | 0.050 | -0.211 | 1.300 | 0.031 | -0.218 | 1.503 | 0 | 0 | -1 | 0 | 0.050 |
| Zfp518b | 0.1460 | -0.172 | 0.836 | 0.457 | -0.037 | 0.340 | 0.005 | -0.216 | 2.278 | 0 | 0 | -1 | 0 | 0.050 |
| BC024063 | 0.2853 | -0.196 | 0.545 | 0.669 | -0.063 | 0.174 | 0.013 | -0.214 | 1.891 | 0 | 0 | -1 | 0 | 0.050 |
| Nrp1 | 0.0507 | -0.073 | 1.295 | 0.076 | -0.224 | 1.121 | 0.038 | -0.214 | 1.418 | 0 | 0 | -1 | 0 | 0.050 |
| Cgrrf1 | 0.1350 | -0.124 | 0.870 | 0.228 | -0.163 | 0.642 | 0.005 | -0.213 | 2.309 | 0 | 0 | -1 | 0 | 0.050 |
| Zfp143 | 0.2323 | -0.113 | 0.634 | 0.065 | -0.195 | 1.185 | 0.007 | -0.210 | 2.145 | 0 | 0 | -1 | 0 | 0.050 |
| Epha2 | 0.0714 | -0.172 | 1.146 | 0.064 | -0.225 | 1.195 | 0.007 | -0.210 | 2.131 | 0 | 0 | -1 | 0 | 0.050 |
| Gvin1 | 0.5870 | -0.059 | 0.231 | 0.760 | 0.058 | 0.119 | 0.002 | -0.209 | 2.773 | 0 | 0 | -1 | 0 | 0.050 |
| Pdlim5 | 0.0516 | -0.106 | 1.287 | 0.071 | -0.116 | 1.146 | 0.022 | -0.208 | 1.660 | 0 | 0 | -1 | 0 | 0.050 |
| Cryz | 0.0465 | -0.123 | 1.332 | 0.278 | -0.063 | 0.556 | 0.016 | -0.206 | 1.783 | 0 | 0 | -1 | 0 | 0.050 |
| Cox18 | 0.0768 | -0.092 | 1.114 | 0.396 | 0.032 | 0.402 | 0.005 | -0.202 | 2.287 | 0 | 0 | -1 | 0 | 0.050 |
| Mrpl32 | 0.0036 | -0.115 | 2.442 | 0.446 | 0.054 | 0.350 | 0.002 | -0.201 | 2.778 | 0 | 0 | -1 | 0 | 0.050 |
| Klhdc4 | 0.0460 | -0.133 | 1.337 | 0.052 | -0.351 | 1.288 | 0.019 | -0.200 | 1.729 | 0 | 0 | -1 | 0 | 0.050 |
| Hat1 | 0.4761 | 0.019 | 0.322 | 0.262 | -0.096 | 0.582 | 0.006 | -0.199 | 2.220 | 0 | 0 | -1 | 0 | 0.050 |
| Adamts4 | 0.3298 | -0.102 | 0.482 | 0.219 | -0.206 | 0.660 | 0.003 | -0.197 | 2.470 | 0 | 0 | -1 | 0 | 0.050 |
| Ajuba | 0.0186 | -0.091 | 1.730 | 0.064 | -0.165 | 1.196 | 0.037 | -0.196 | 1.434 | 0 | 0 | -1 | 0 | 0.050 |
| Klf4 | 0.5845 | 0.181 | 0.233 | 0.653 | -0.050 | 0.185 | 0.048 | -0.196 | 1.318 | 0 | 0 | -1 | 0 | 0.050 |
| Steap1 | 0.0802 | -0.188 | 1.096 | 0.059 | -0.168 | 1.230 | 0.022 | -0.195 | 1.659 | 0 | 0 | -1 | 0 | 0.050 |
| Plpp2 | 0.2286 | -0.136 | 0.641 | 0.266 | -0.088 | 0.575 | 0.016 | -0.192 | 1.788 | 0 | 0 | -1 | 0 | 0.050 |
| Tcf7l2 | 0.1154 | -0.063 | 0.938 | 0.476 | -0.025 | 0.322 | 0.011 | -0.190 | 1.956 | 0 | 0 | -1 | 0 | 0.050 |
| Nipal1 | 0.1692 | -0.104 | 0.772 | 0.512 | -0.047 | 0.290 | 0.001 | -0.189 | 3.086 | 0 | 0 | -1 | 0 | 0.050 |

| | | | | | | | | | | | | | | |
|---|---|---|---|---|---|---|---|---|---|---|---|---|---|---|
| Errfi1 | 0.2935 | -0.179 | 0.532 | 0.059 | -0.185 | 1.226 | 0.016 | -0.188 | 1.786 | 0 | 0 | -1 | 0 | 0.050 |
| Csrp2 | 0.0551 | -0.136 | 1.259 | 0.066 | -0.113 | 1.179 | 0.002 | -0.187 | 2.655 | 0 | 0 | -1 | 0 | 0.050 |
| Ogg1 | 0.4201 | -0.185 | 0.377 | 0.270 | 0.066 | 0.569 | 0.023 | -0.186 | 1.637 | 0 | 0 | -1 | 0 | 0.050 |
| Lrrc9 | 0.0113 | -0.116 | 1.949 | 0.203 | -0.224 | 0.692 | 0.020 | -0.185 | 1.695 | 0 | 0 | -1 | 0 | 0.050 |
| Tinagl1 | 0.3543 | -0.101 | 0.451 | 0.161 | -0.263 | 0.793 | 0.033 | -0.184 | 1.482 | 0 | 0 | -1 | 0 | 0.050 |
| Lman2l | 0.2159 | -0.107 | 0.666 | 0.295 | -0.100 | 0.531 | 0.000 | -0.184 | 3.450 | 0 | 0 | -1 | 0 | 0.050 |
| Sorcs2 | 0.2430 | -0.054 | 0.614 | 0.081 | -0.232 | 1.093 | 0.047 | -0.184 | 1.329 | 0 | 0 | -1 | 0 | 0.050 |
| Wnt10b | 0.6931 | -0.037 | 0.159 | 0.064 | -0.115 | 1.196 | 0.039 | -0.184 | 1.407 | 0 | 0 | -1 | 0 | 0.050 |
| Zc3h11a | 0.0790 | -0.078 | 1.102 | 0.223 | -0.077 | 0.651 | 0.004 | -0.183 | 2.416 | 0 | 0 | -1 | 0 | 0.050 |
| Gm4832 | 0.1011 | -0.139 | 0.995 | 0.623 | -0.041 | 0.206 | 0.001 | -0.183 | 3.068 | 0 | 0 | -1 | 0 | 0.050 |
| Hspg2 | 0.2914 | -0.107 | 0.535 | 0.161 | -0.184 | 0.794 | 0.009 | -0.183 | 2.040 | 0 | 0 | -1 | 0 | 0.050 |
| Frg1 | 0.0038 | -0.116 | 2.417 | 0.014 | -0.101 | 1.842 | 0.003 | -0.182 | 2.497 | 0 | 0 | -1 | 0 | 0.050 |
| Bpnt1 | 0.1869 | -0.108 | 0.728 | 0.028 | -0.105 | 1.555 | 0.000 | -0.182 | 3.476 | 0 | 0 | -1 | 0 | 0.050 |
| Atp1b3 | 0.1188 | -0.176 | 0.925 | 0.083 | -0.194 | 1.083 | 0.038 | -0.182 | 1.424 | 0 | 0 | -1 | 0 | 0.050 |
| Taf6l | 0.1948 | -0.130 | 0.710 | 0.071 | -0.166 | 1.151 | 0.010 | -0.182 | 2.014 | 0 | 0 | -1 | 0 | 0.050 |
| Sorl1 | 0.9488 | -0.014 | 0.023 | 0.290 | -0.118 | 0.537 | 0.017 | -0.180 | 1.771 | 0 | 0 | -1 | 0 | 0.050 |
| Tmem177 | 0.5514 | 0.050 | 0.259 | 0.566 | -0.044 | 0.247 | 0.027 | -0.179 | 1.563 | 0 | 0 | -1 | 0 | 0.050 |
| Dclk1 | 0.0204 | -0.107 | 1.690 | 0.063 | -0.140 | 1.203 | 0.011 | -0.179 | 1.978 | 0 | 0 | -1 | 0 | 0.050 |
| Sytl5 | 0.3140 | -0.038 | 0.503 | 0.065 | -0.120 | 1.186 | 0.012 | -0.177 | 1.925 | 0 | 0 | -1 | 0 | 0.050 |
| Insrr | 0.5427 | -0.037 | 0.265 | 0.009 | -0.131 | 2.034 | 0.013 | -0.177 | 1.872 | 0 | 0 | -1 | 0 | 0.050 |
| Iqub | 0.4649 | -0.080 | 0.333 | 0.284 | -0.054 | 0.547 | 0.030 | -0.177 | 1.530 | 0 | 0 | -1 | 0 | 0.050 |
| Scaf1 | 0.7032 | 0.096 | 0.153 | 0.005 | -0.123 | 2.276 | 0.042 | -0.176 | 1.378 | 0 | 0 | -1 | 0 | 0.050 |
| Zfp36 | 0.0321 | -0.056 | 1.494 | 0.072 | -0.151 | 1.146 | 0.003 | -0.175 | 2.456 | 0 | 0 | -1 | 0 | 0.050 |
| mKIAA1466 | 0.1731 | 0.050 | 0.762 | 0.073 | -0.169 | 1.135 | 0.016 | -0.174 | 1.806 | 0 | 0 | -1 | 0 | 0.050 |
| Tp53rk | 0.2940 | -0.071 | 0.532 | 0.252 | -0.160 | 0.598 | 0.003 | -0.174 | 2.538 | 0 | 0 | -1 | 0 | 0.050 |
| Mrpl16 | 0.7964 | -0.020 | 0.099 | 0.119 | -0.060 | 0.926 | 0.000 | -0.174 | 3.313 | 0 | 0 | -1 | 0 | 0.050 |
| Wdr73 | 0.1394 | -0.186 | 0.856 | 0.295 | -0.102 | 0.531 | 0.047 | -0.173 | 1.324 | 0 | 0 | -1 | 0 | 0.050 |
| Igf2r | 0.3075 | 0.037 | 0.512 | 0.096 | -0.207 | 1.017 | 0.013 | -0.173 | 1.879 | 0 | 0 | -1 | 0 | 0.050 |
| Tle1 | 0.0073 | 0.118 | 2.134 | 0.628 | -0.049 | 0.202 | 0.001 | -0.172 | 3.182 | 0 | 0 | -1 | 0 | 0.050 |
| Hdac2 | 0.0601 | -0.063 | 1.221 | 0.084 | -0.118 | 1.074 | 0.006 | -0.172 | 2.257 | 0 | 0 | -1 | 0 | 0.050 |
| Unkl | 0.0593 | -0.197 | 1.227 | 0.158 | -0.199 | 0.801 | 0.020 | -0.172 | 1.692 | 0 | 0 | -1 | 0 | 0.050 |
| Myrf | 0.0327 | -0.134 | 1.486 | 0.152 | -0.110 | 0.819 | 0.020 | -0.171 | 1.699 | 0 | 0 | -1 | 0 | 0.050 |
| Blvra | 0.2482 | -0.054 | 0.605 | 0.036 | -0.104 | 1.444 | 0.012 | -0.170 | 1.917 | 0 | 0 | -1 | 0 | 0.050 |
| Lxn | 0.2581 | -0.054 | 0.588 | 0.109 | -0.110 | 0.964 | 0.000 | -0.169 | 3.379 | 0 | 0 | -1 | 0 | 0.050 |
| Mtarc2 | 0.3077 | -0.046 | 0.512 | 0.095 | -0.128 | 1.022 | 0.024 | -0.169 | 1.626 | 0 | 0 | -1 | 0 | 0.050 |
| Fam111a | 0.1345 | 0.031 | 0.871 | 0.800 | -0.014 | 0.097 | 0.005 | -0.168 | 2.269 | 0 | 0 | -1 | 0 | 0.050 |
| Bnip2 | 0.7932 | -0.033 | 0.101 | 0.153 | -0.130 | 0.816 | 0.001 | -0.167 | 3.095 | 0 | 0 | -1 | 0 | 0.050 |
| Dsg2 | 0.0834 | -0.085 | 1.079 | 0.215 | -0.162 | 0.667 | 0.038 | -0.166 | 1.422 | 0 | 0 | -1 | 0 | 0.050 |
| Ppp2r2d | 0.8470 | 0.014 | 0.072 | 0.216 | -0.158 | 0.666 | 0.038 | -0.166 | 1.421 | 0 | 0 | -1 | 0 | 0.050 |
| Gata3 | 0.9903 | -0.001 | 0.004 | 0.571 | -0.034 | 0.243 | 0.000 | -0.166 | 3.396 | 0 | 0 | -1 | 0 | 0.050 |
| Nol12 | 0.0279 | 0.071 | 1.554 | 0.144 | -0.135 | 0.843 | 0.010 | -0.165 | 2.021 | 0 | 0 | -1 | 0 | 0.050 |
| Ptgr1 | 0.3488 | -0.034 | 0.457 | 0.021 | -0.130 | 1.683 | 0.017 | -0.164 | 1.760 | 0 | 0 | -1 | 0 | 0.050 |
| Slc29a2 | 0.0711 | -0.258 | 1.148 | 0.125 | -0.208 | 0.902 | 0.022 | -0.164 | 1.649 | 0 | 0 | -1 | 0 | 0.050 |
| Knop1 | 0.8279 | 0.020 | 0.082 | 0.695 | -0.021 | 0.158 | 0.017 | -0.163 | 1.766 | 0 | 0 | -1 | 0 | 0.050 |
| Ptprn2 | 0.8177 | 0.032 | 0.087 | 0.068 | -0.263 | 1.167 | 0.041 | -0.163 | 1.389 | 0 | 0 | -1 | 0 | 0.050 |
| Med25 | 0.4475 | -0.196 | 0.349 | 0.703 | -0.018 | 0.153 | 0.047 | -0.163 | 1.329 | 0 | 0 | -1 | 0 | 0.050 |
| Pip4p1 | 0.8249 | 0.021 | 0.084 | 0.046 | -0.134 | 1.337 | 0.049 | -0.163 | 1.309 | 0 | 0 | -1 | 0 | 0.050 |
| Xylb | 0.5260 | 0.024 | 0.279 | 0.119 | -0.054 | 0.924 | 0.001 | -0.162 | 2.977 | 0 | 0 | -1 | 0 | 0.050 |
| Rps20 | 0.0013 | -0.137 | 2.878 | 0.195 | -0.116 | 0.710 | 0.013 | -0.162 | 1.885 | 0 | 0 | -1 | 0 | 0.050 |
| Znf706 | 0.0772 | -0.090 | 1.112 | 0.035 | -0.114 | 1.456 | 0.025 | -0.161 | 1.606 | 0 | 0 | -1 | 0 | 0.050 |
| Itga2 | 0.3980 | -0.020 | 0.400 | 0.072 | -0.185 | 1.141 | 0.002 | -0.160 | 2.659 | 0 | 0 | -1 | 0 | 0.050 |
| H6pd | 0.5043 | -0.024 | 0.297 | 0.028 | -0.091 | 1.546 | 0.001 | -0.160 | 2.911 | 0 | 0 | -1 | 0 | 0.050 |
| Lrrc45 | 0.2970 | -0.070 | 0.527 | 0.509 | -0.078 | 0.293 | 0.035 | -0.160 | 1.455 | 0 | 0 | -1 | 0 | 0.050 |
| Rxrb | 0.0463 | -0.105 | 1.334 | 0.088 | -0.170 | 1.058 | 0.004 | -0.159 | 2.360 | 0 | 0 | -1 | 0 | 0.050 |
| Wnt3 | 0.3340 | -0.047 | 0.476 | 0.084 | -0.154 | 1.076 | 0.012 | -0.159 | 1.923 | 0 | 0 | -1 | 0 | 0.050 |
| Zfp948 | 0.7639 | -0.016 | 0.117 | 0.737 | -0.022 | 0.132 | 0.010 | -0.158 | 2.010 | 0 | 0 | -1 | 0 | 0.050 |
| Cycs | 0.0576 | 0.113 | 1.239 | 0.449 | -0.080 | 0.348 | 0.006 | -0.158 | 2.230 | 0 | 0 | -1 | 0 | 0.050 |
| Shq1 | 0.4406 | -0.040 | 0.356 | 0.581 | -0.086 | 0.236 | 0.002 | -0.157 | 2.726 | 0 | 0 | -1 | 0 | 0.050 |
| Dxo | 0.1076 | -0.120 | 0.968 | 0.677 | -0.028 | 0.170 | 0.000 | -0.157 | 3.387 | 0 | 0 | -1 | 0 | 0.050 |
| Cdc123 | 0.0156 | -0.110 | 1.808 | 0.279 | -0.092 | 0.555 | 0.001 | -0.157 | 2.942 | 0 | 0 | -1 | 0 | 0.050 |
| Tor1b | 0.6334 | 0.023 | 0.198 | 0.506 | -0.064 | 0.296 | 0.003 | -0.156 | 2.461 | 0 | 0 | -1 | 0 | 0.050 |
| Pttg1ip | 0.6257 | -0.045 | 0.204 | 0.800 | -0.038 | 0.097 | 0.001 | -0.155 | 3.241 | 0 | 0 | -1 | 0 | 0.050 |
| Ing5 | 0.6854 | 0.031 | 0.164 | 0.555 | -0.069 | 0.256 | 0.019 | -0.155 | 1.715 | 0 | 0 | -1 | 0 | 0.050 |
| Leng1 | 0.4685 | -0.057 | 0.329 | 0.535 | -0.049 | 0.271 | 0.003 | -0.154 | 2.526 | 0 | 0 | -1 | 0 | 0.050 |
| Ift80 | 0.0479 | -0.132 | 1.320 | 0.022 | -0.094 | 1.649 | 0.027 | -0.154 | 1.572 | 0 | 0 | -1 | 0 | 0.050 |
| Lef1 | 0.4260 | -0.091 | 0.371 | 0.319 | -0.096 | 0.496 | 0.001 | -0.154 | 3.016 | 0 | 0 | -1 | 0 | 0.050 |
| Plau | 0.1832 | -0.066 | 0.737 | 0.552 | -0.079 | 0.258 | 0.009 | -0.154 | 2.057 | 0 | 0 | -1 | 0 | 0.050 |
| Gsto1 | 0.0172 | -0.110 | 1.765 | 0.598 | -0.028 | 0.223 | 0.005 | -0.154 | 2.338 | 0 | 0 | -1 | 0 | 0.050 |
| Rrad | 0.3011 | -0.080 | 0.521 | 0.054 | -0.177 | 1.268 | 0.001 | -0.153 | 3.245 | 0 | 0 | -1 | 0 | 0.050 |
| Otud6a | 0.5969 | -0.061 | 0.224 | 0.642 | -0.050 | 0.192 | 0.048 | -0.153 | 1.316 | 0 | 0 | -1 | 0 | 0.050 |
| Rfwd3 | 0.3988 | -0.079 | 0.399 | 0.037 | -0.113 | 1.431 | 0.023 | -0.153 | 1.643 | 0 | 0 | -1 | 0 | 0.050 |

| | | | | | | | | | | | | | |
|---|---|---|---|---|---|---|---|---|---|---|---|---|---|
| Thsd1 | 0.0696 | -0.137 | 1.158 | 0.204 | -0.091 | 0.690 | 0.037 | -0.152 | 1.432 | 0 | 0 | -1 | 0 | 0.050 |
| Cpox | 0.6510 | 0.028 | 0.186 | 0.303 | -0.103 | 0.519 | 0.031 | -0.151 | 1.507 | 0 | 0 | -1 | 0 | 0.050 |
| Rnf25 | 0.0124 | -0.119 | 1.907 | 0.098 | -0.081 | 1.010 | 0.001 | -0.151 | 2.955 | 0 | 0 | -1 | 0 | 0.050 |
| Furin | 0.1850 | 0.108 | 0.733 | 0.118 | -0.238 | 0.929 | 0.001 | -0.150 | 3.001 | 0 | 0 | -1 | 0 | 0.050 |
| Sgo1 | 0.7715 | -0.043 | 0.113 | 0.646 | -0.065 | 0.190 | 0.031 | -0.150 | 1.515 | 0 | 0 | -1 | 0 | 0.050 |
| Zc3hc1 | 0.9762 | 0.002 | 0.010 | 0.900 | -0.012 | 0.046 | 0.001 | -0.149 | 3.271 | 0 | 0 | -1 | 0 | 0.050 |
| Zfp91 | 0.1911 | 0.046 | 0.719 | 0.643 | 0.027 | 0.192 | 0.001 | -0.149 | 2.878 | 0 | 0 | -1 | 0 | 0.050 |
| Dennd3 | 0.5194 | -0.064 | 0.285 | 0.902 | 0.012 | 0.045 | 0.008 | -0.147 | 2.119 | 0 | 0 | -1 | 0 | 0.050 |
| Rtf2 | 0.5892 | -0.030 | 0.230 | 0.482 | -0.070 | 0.317 | 0.003 | -0.147 | 2.456 | 0 | 0 | -1 | 0 | 0.050 |
| Mat2b | 0.1306 | 0.065 | 0.884 | 0.600 | -0.030 | 0.222 | 0.047 | -0.146 | 1.331 | 0 | 0 | -1 | 0 | 0.050 |
| Shkbp1 | 0.5114 | -0.092 | 0.291 | 0.058 | -0.084 | 1.235 | 0.012 | -0.146 | 1.936 | 0 | 0 | -1 | 0 | 0.050 |
| Crtap | 0.0954 | -0.047 | 1.020 | 0.124 | -0.098 | 0.905 | 0.033 | -0.145 | 1.479 | 0 | 0 | -1 | 0 | 0.050 |
| Ext1 | 0.5222 | 0.065 | 0.282 | 0.061 | -0.108 | 1.213 | 0.024 | -0.145 | 1.616 | 0 | 0 | -1 | 0 | 0.050 |
| Nufip1 | 0.0152 | -0.076 | 1.817 | 0.011 | -0.110 | 1.974 | 0.036 | -0.145 | 1.441 | 0 | 0 | -1 | 0 | 0.050 |
| Pdik1l | 0.1608 | -0.200 | 0.794 | 0.978 | 0.005 | 0.009 | 0.048 | -0.142 | 1.322 | 0 | 0 | -1 | 0 | 0.050 |
| Gpr176 | 0.1291 | -0.223 | 0.889 | 0.218 | -0.077 | 0.661 | 0.017 | -0.141 | 1.773 | 0 | 0 | -1 | 0 | 0.050 |
| Dpp3 | 0.3407 | -0.078 | 0.468 | 0.061 | -0.059 | 1.213 | 0.021 | -0.140 | 1.685 | 0 | 0 | -1 | 0 | 0.050 |
| Cep70 | 0.2666 | -0.236 | 0.574 | 0.209 | -0.078 | 0.681 | 0.033 | -0.139 | 1.482 | 0 | 0 | -1 | 0 | 0.050 |
| Uqcr10 | 0.2703 | -0.167 | 0.568 | 0.013 | -0.122 | 1.895 | 0.031 | -0.139 | 1.507 | 0 | 0 | -1 | 0 | 0.050 |
| Mtr | 0.2978 | -0.047 | 0.526 | 0.148 | -0.048 | 0.829 | 0.023 | -0.139 | 1.631 | 0 | 0 | -1 | 0 | 0.050 |
| Ttc39c | 0.9973 | 0.000 | 0.001 | 0.307 | -0.066 | 0.513 | 0.036 | -0.139 | 1.444 | 0 | 0 | -1 | 0 | 0.050 |
| Cpt1c | 0.4400 | -0.082 | 0.357 | 0.041 | -0.116 | 1.391 | 0.022 | -0.139 | 1.649 | 0 | 0 | -1 | 0 | 0.050 |
| Nme7 | 0.1674 | -0.058 | 0.776 | 0.036 | -0.136 | 1.449 | 0.004 | -0.138 | 2.354 | 0 | 0 | -1 | 0 | 0.050 |
| Ctnna3 | 0.2999 | -0.227 | 0.523 | 0.061 | -0.610 | 1.213 | 0.059 | -0.656 | 1.232 | 0 | 0 | 0 | 0 | 0.050 |
| C3 | 0.1041 | -0.098 | 0.983 | 0.489 | -0.216 | 0.311 | 0.090 | -0.459 | 1.046 | 0 | 0 | 0 | 0 | 0.050 |
| Ralgapa1 | 0.7342 | 0.033 | 0.134 | 0.683 | -0.144 | 0.165 | 0.120 | -0.350 | 0.923 | 0 | 0 | 0 | 0 | 0.050 |
| Rilp | 0.8875 | -0.016 | 0.052 | 0.098 | -0.211 | 1.011 | 0.080 | -0.342 | 1.099 | 0 | 0 | 0 | 0 | 0.050 |
| Epdr1 | 0.7656 | -0.049 | 0.116 | 0.608 | -0.075 | 0.216 | 0.093 | -0.335 | 1.031 | 0 | 0 | 0 | 0 | 0.050 |
| Col11a1 | 0.7525 | -0.035 | 0.124 | 0.113 | -0.548 | 0.945 | 0.262 | -0.331 | 0.581 | 0 | 0 | 0 | 0 | 0.050 |
| Mettl25 | 0.1059 | -0.093 | 0.975 | 0.447 | -0.231 | 0.350 | 0.229 | -0.296 | 0.639 | 0 | 0 | 0 | 0 | 0.050 |
| Cavin3 | 0.8234 | -0.037 | 0.084 | 0.057 | -0.337 | 1.241 | 0.145 | -0.294 | 0.838 | 0 | 0 | 0 | 0 | 0.050 |
| Fbn1 | 0.2585 | -0.024 | 0.588 | 0.065 | -0.242 | 1.186 | 0.059 | -0.289 | 1.229 | 0 | 0 | 0 | 0 | 0.050 |
| Suv39h2 | 0.0224 | 0.075 | 1.649 | 0.420 | 0.050 | 0.377 | 0.109 | -0.280 | 0.961 | 0 | 0 | 0 | 0 | 0.050 |
| Decr1 | 0.2697 | -0.082 | 0.569 | 0.114 | -0.409 | 0.944 | 0.242 | -0.280 | 0.617 | 0 | 0 | 0 | 0 | 0.050 |
| Ppp1r15b | 0.5063 | -0.226 | 0.296 | 0.401 | -0.082 | 0.397 | 0.086 | -0.280 | 1.067 | 0 | 0 | 0 | 0 | 0.050 |
| Midn | 0.1740 | -0.248 | 0.759 | 0.067 | -0.264 | 1.177 | 0.055 | -0.277 | 1.258 | 0 | 0 | 0 | 0 | 0.050 |
| Apoa1 | 0.0842 | -0.436 | 1.075 | 0.324 | -0.146 | 0.490 | 0.359 | -0.276 | 0.445 | 0 | 0 | 0 | 0 | 0.050 |
| Mpst | 0.0596 | -0.076 | 1.225 | 0.127 | -0.315 | 0.895 | 0.136 | -0.268 | 0.867 | 0 | 0 | 0 | 0 | 0.050 |
| Ppil3 | 0.7161 | -0.054 | 0.145 | 0.186 | -0.058 | 0.731 | 0.106 | -0.266 | 0.973 | 0 | 0 | 0 | 0 | 0.050 |
| Postn | 0.8615 | -0.035 | 0.065 | 0.315 | -0.160 | 0.501 | 0.266 | -0.263 | 0.575 | 0 | 0 | 0 | 0 | 0.050 |
| Smoc2 | 0.6374 | 0.019 | 0.196 | 0.085 | -0.211 | 1.069 | 0.062 | -0.262 | 1.211 | 0 | 0 | 0 | 0 | 0.050 |
| Ccdc124 | 0.7679 | -0.028 | 0.115 | 0.273 | -0.078 | 0.564 | 0.053 | -0.255 | 1.277 | 0 | 0 | 0 | 0 | 0.050 |
| Sp2 | 0.1058 | -0.224 | 0.975 | 0.893 | -0.006 | 0.049 | 0.116 | -0.249 | 0.934 | 0 | 0 | 0 | 0 | 0.050 |
| Otud6b | 0.0734 | -0.077 | 1.134 | 0.960 | 0.005 | 0.018 | 0.064 | -0.247 | 1.193 | 0 | 0 | 0 | 0 | 0.050 |
| Cmtm4 | 0.0809 | -0.259 | 1.092 | 0.084 | 0.096 | 1.078 | 0.201 | -0.245 | 0.697 | 0 | 0 | 0 | 0 | 0.050 |
| Aebp1 | 0.0369 | 0.117 | 1.433 | 0.130 | -0.406 | 0.888 | 0.326 | -0.244 | 0.487 | 0 | 0 | 0 | 0 | 0.050 |
| Zfp36l2 | 0.0441 | -0.094 | 1.356 | 0.188 | -0.161 | 0.726 | 0.086 | -0.243 | 1.064 | 0 | 0 | 0 | 0 | 0.050 |
| Dusp11 | 0.6502 | -0.037 | 0.187 | 0.122 | -0.182 | 0.914 | 0.298 | -0.242 | 0.526 | 0 | 0 | 0 | 0 | 0.050 |
| Ifi44 | 0.0055 | 0.093 | 2.258 | 0.385 | -0.041 | 0.415 | 0.092 | -0.242 | 1.035 | 0 | 0 | 0 | 0 | 0.050 |
| Tmem230 | 0.4798 | -0.044 | 0.319 | 0.922 | -0.005 | 0.035 | 0.085 | -0.234 | 1.070 | 0 | 0 | 0 | 0 | 0.050 |
| Rapgef4 | 0.0985 | -0.313 | 1.007 | 0.120 | -0.346 | 0.921 | 0.111 | -0.227 | 0.955 | 0 | 0 | 0 | 0 | 0.050 |
| Ecscr | 0.5575 | -0.115 | 0.254 | 0.352 | -0.096 | 0.453 | 0.062 | -0.224 | 1.207 | 0 | 0 | 0 | 0 | 0.050 |
| Coil | 0.0487 | 0.084 | 1.313 | 0.215 | -0.161 | 0.667 | 0.160 | -0.223 | 0.797 | 0 | 0 | 0 | 0 | 0.050 |
| Iap | 0.1441 | -0.077 | 0.841 | 0.261 | -0.146 | 0.583 | 0.086 | -0.221 | 1.067 | 0 | 0 | 0 | 0 | 0.050 |
| Hivep2 | 0.1537 | -0.271 | 0.813 | 0.096 | -0.180 | 1.016 | 0.444 | -0.219 | 0.353 | 0 | 0 | 0 | 0 | 0.050 |
| Znhit3 | 0.0361 | -0.130 | 1.443 | 0.079 | -0.308 | 1.103 | 0.161 | -0.216 | 0.794 | 0 | 0 | 0 | 0 | 0.050 |
| Runx1 | 0.4730 | -0.141 | 0.325 | 0.040 | -0.095 | 1.401 | 0.325 | -0.213 | 0.488 | 0 | 0 | 0 | 0 | 0.050 |
| Arhgef26 | 0.6870 | -0.069 | 0.163 | 0.058 | -0.312 | 1.233 | 0.067 | -0.212 | 1.172 | 0 | 0 | 0 | 0 | 0.050 |
| Klf9 | 0.0550 | -0.148 | 1.260 | 0.141 | -0.137 | 0.850 | 0.360 | -0.210 | 0.443 | 0 | 0 | 0 | 0 | 0.050 |
| Zfp512b | 0.4727 | 0.064 | 0.325 | 0.256 | -0.125 | 0.591 | 0.150 | -0.210 | 0.823 | 0 | 0 | 0 | 0 | 0.050 |
| Rybp | 0.2005 | -0.129 | 0.698 | 0.670 | -0.034 | 0.174 | 0.092 | -0.209 | 1.037 | 0 | 0 | 0 | 0 | 0.050 |
| Gdpd5 | 0.2745 | -0.119 | 0.561 | 0.162 | -0.147 | 0.790 | 0.074 | -0.208 | 1.133 | 0 | 0 | 0 | 0 | 0.050 |
| Ppip5k1 | 0.0017 | 0.128 | 2.759 | 0.093 | -0.241 | 1.032 | 0.147 | -0.207 | 0.832 | 0 | 0 | 0 | 0 | 0.050 |
| Mgp | 0.6668 | 0.018 | 0.176 | 0.161 | -0.176 | 0.794 | 0.062 | -0.206 | 1.207 | 0 | 0 | 0 | 0 | 0.050 |
| Tmbim6 | 0.9300 | -0.017 | 0.032 | 0.131 | -0.417 | 0.884 | 0.173 | -0.202 | 0.761 | 0 | 0 | 0 | 0 | 0.050 |
| Hdhd5 | 0.1851 | -0.129 | 0.733 | 0.633 | 0.019 | 0.199 | 0.185 | -0.201 | 0.733 | 0 | 0 | 0 | 0 | 0.050 |
| Hoxd13 | 0.2861 | -0.043 | 0.543 | 0.279 | -0.100 | 0.554 | 0.069 | -0.198 | 1.162 | 0 | 0 | 0 | 0 | 0.050 |
| Ppp2r2b | 0.8681 | -0.012 | 0.061 | 0.186 | -0.162 | 0.730 | 0.072 | -0.198 | 1.144 | 0 | 0 | 0 | 0 | 0.050 |
| Ecm1 | 0.0199 | -0.085 | 1.701 | 0.070 | -0.167 | 1.158 | 0.074 | -0.197 | 1.133 | 0 | 0 | 0 | 0 | 0.050 |
| Abca13 | 0.7651 | 0.052 | 0.116 | 0.080 | -0.149 | 1.097 | 0.114 | -0.197 | 0.943 | 0 | 0 | 0 | 0 | 0.050 |
| Wnt10a | 0.2613 | -0.038 | 0.583 | 0.401 | -0.064 | 0.397 | 0.066 | -0.197 | 1.182 | 0 | 0 | 0 | 0 | 0.050 |

| | | | | | | | | | | | | | | |
|---|---|---|---|---|---|---|---|---|---|---|---|---|---|---|
| Ebf1 | 0.5580 | -0.154 | 0.253 | 0.426 | -0.138 | 0.371 | 0.221 | -0.197 | 0.655 | 0 | 0 | 0 | 0 | 0.050 |
| Fbxo25 | 0.1037 | -0.228 | 0.984 | 0.955 | 0.007 | 0.020 | 0.087 | -0.193 | 1.063 | 0 | 0 | 0 | 0 | 0.050 |
| Tspan6 | 0.0403 | -0.112 | 1.395 | 0.273 | -0.113 | 0.564 | 0.136 | -0.193 | 0.867 | 0 | 0 | 0 | 0 | 0.050 |
| Pptc7 | 0.8182 | 0.026 | 0.087 | 0.231 | -0.068 | 0.636 | 0.066 | -0.192 | 1.182 | 0 | 0 | 0 | 0 | 0.050 |
| Micall1 | 0.2207 | 0.160 | 0.656 | 0.547 | -0.039 | 0.262 | 0.369 | -0.190 | 0.433 | 0 | 0 | 0 | 0 | 0.050 |
| Akr1b10 | 0.1583 | -0.098 | 0.800 | 0.140 | -0.187 | 0.855 | 0.086 | -0.188 | 1.067 | 0 | 0 | 0 | 0 | 0.050 |
| Gm12500 | 0.2469 | -0.200 | 0.608 | 0.143 | 0.351 | 0.844 | 0.221 | -0.188 | 0.656 | 0 | 0 | 0 | 0 | 0.050 |
| Tnfaip8 | 0.8905 | 0.030 | 0.050 | 0.250 | -0.251 | 0.602 | 0.395 | -0.187 | 0.404 | 0 | 0 | 0 | 0 | 0.050 |
| G0s2 | 0.1745 | -0.119 | 0.758 | 0.357 | -0.067 | 0.448 | 0.183 | -0.186 | 0.737 | 0 | 0 | 0 | 0 | 0.050 |
| Lama4 | 0.7063 | -0.045 | 0.151 | 0.143 | -0.190 | 0.844 | 0.059 | -0.185 | 1.233 | 0 | 0 | 0 | 0 | 0.050 |
| Mfsd5 | 0.9542 | 0.003 | 0.020 | 0.908 | 0.011 | 0.042 | 0.077 | -0.184 | 1.114 | 0 | 0 | 0 | 0 | 0.050 |
| Cybrd1 | 0.9771 | -0.003 | 0.010 | 0.262 | -0.178 | 0.581 | 0.369 | -0.183 | 0.433 | 0 | 0 | 0 | 0 | 0.050 |
| Heph | 0.8433 | -0.027 | 0.074 | 0.676 | -0.020 | 0.170 | 0.174 | -0.183 | 0.759 | 0 | 0 | 0 | 0 | 0.050 |
| Phlpp1 | 0.1460 | -0.227 | 0.836 | 0.468 | -0.112 | 0.330 | 0.083 | -0.181 | 1.082 | 0 | 0 | 0 | 0 | 0.050 |
| Fchsd2 | 0.1195 | -0.163 | 0.923 | 0.160 | -0.184 | 0.795 | 0.224 | -0.179 | 0.650 | 0 | 0 | 0 | 0 | 0.050 |
| Rpl23 | 0.0661 | -0.087 | 1.180 | 0.167 | -0.148 | 0.778 | 0.067 | -0.179 | 1.177 | 0 | 0 | 0 | 0 | 0.050 |
| Shank1 | 0.6920 | -0.072 | 0.160 | 0.407 | -0.094 | 0.390 | 0.193 | -0.178 | 0.714 | 0 | 0 | 0 | 0 | 0.050 |
| Nans | 0.9334 | -0.006 | 0.030 | 0.144 | -0.082 | 0.841 | 0.302 | -0.176 | 0.520 | 0 | 0 | 0 | 0 | 0.050 |
| Lrr1 | 0.7193 | -0.047 | 0.143 | 0.090 | -0.133 | 1.047 | 0.101 | -0.175 | 0.995 | 0 | 0 | 0 | 0 | 0.050 |
| Ifi203 | 0.2767 | 0.070 | 0.558 | 0.104 | -0.155 | 0.982 | 0.066 | -0.175 | 1.180 | 0 | 0 | 0 | 0 | 0.050 |
| Znf271 | 0.1559 | -0.112 | 0.807 | 0.293 | -0.060 | 0.533 | 0.067 | -0.175 | 1.176 | 0 | 0 | 0 | 0 | 0.050 |
| Igdcc4 | 0.0720 | -0.131 | 1.142 | 0.608 | -0.069 | 0.216 | 0.246 | -0.174 | 0.608 | 0 | 0 | 0 | 0 | 0.050 |
| Aida | 0.5330 | 0.035 | 0.273 | 0.538 | -0.076 | 0.270 | 0.268 | -0.174 | 0.572 | 0 | 0 | 0 | 0 | 0.050 |
| Zfp106 | 0.1849 | -0.146 | 0.733 | 0.030 | -0.115 | 1.525 | 0.220 | -0.173 | 0.658 | 0 | 0 | 0 | 0 | 0.050 |
| Icam5 | 0.7189 | 0.010 | 0.143 | 0.021 | -0.087 | 1.679 | 0.195 | -0.173 | 0.710 | 0 | 0 | 0 | 0 | 0.050 |
| Atg4b | 0.9823 | -0.005 | 0.008 | 0.405 | -0.158 | 0.392 | 0.070 | -0.172 | 1.155 | 0 | 0 | 0 | 0 | 0.050 |
| Kyat1 | 0.2555 | 0.051 | 0.593 | 0.645 | 0.093 | 0.190 | 0.208 | -0.172 | 0.683 | 0 | 0 | 0 | 0 | 0.050 |
| Tor2a | 0.2286 | 0.091 | 0.641 | 0.050 | -0.128 | 1.304 | 0.053 | -0.171 | 1.277 | 0 | 0 | 0 | 0 | 0.050 |
| Rbm41 | 0.1186 | -0.205 | 0.926 | 0.508 | -0.046 | 0.294 | 0.314 | -0.170 | 0.503 | 0 | 0 | 0 | 0 | 0.050 |
| Lsr | 0.1606 | -0.103 | 0.794 | 0.067 | -0.150 | 1.172 | 0.186 | -0.169 | 0.730 | 0 | 0 | 0 | 0 | 0.050 |
| B4galt1 | 0.7226 | 0.021 | 0.141 | 0.041 | -0.109 | 1.391 | 0.225 | -0.168 | 0.649 | 0 | 0 | 0 | 0 | 0.050 |
| Ppp2r2a | 0.9346 | 0.005 | 0.029 | 0.148 | -0.156 | 0.829 | 0.078 | -0.167 | 1.110 | 0 | 0 | 0 | 0 | 0.050 |
| Prdx6 | 0.6495 | -0.015 | 0.187 | 0.153 | -0.166 | 0.815 | 0.089 | -0.167 | 1.052 | 0 | 0 | 0 | 0 | 0.050 |
| Vcan | 0.4175 | -0.132 | 0.379 | 0.109 | -0.114 | 0.961 | 0.164 | -0.167 | 0.785 | 0 | 0 | 0 | 0 | 0.050 |
| Rgs9 | 0.0583 | 0.472 | 1.234 | 0.330 | -0.251 | 0.482 | 0.374 | -0.167 | 0.428 | 0 | 0 | 0 | 0 | 0.050 |
| Steap3 | 0.0758 | -0.126 | 1.120 | 0.066 | -0.152 | 1.177 | 0.050 | -0.166 | 1.299 | 0 | 0 | 0 | 0 | 0.050 |
| Hyal2 | 0.2635 | -0.245 | 0.579 | 0.092 | -0.156 | 1.038 | 0.151 | -0.166 | 0.820 | 0 | 0 | 0 | 0 | 0.050 |
| Wdr59 | 0.3287 | 0.056 | 0.483 | 0.792 | -0.016 | 0.101 | 0.124 | -0.165 | 0.908 | 0 | 0 | 0 | 0 | 0.050 |
| Casp6 | 0.0476 | -0.084 | 1.322 | 0.068 | -0.106 | 1.168 | 0.058 | -0.165 | 1.235 | 0 | 0 | 0 | 0 | 0.050 |
| Rad9a | 0.0821 | -0.085 | 1.086 | 0.476 | -0.072 | 0.323 | 0.059 | -0.165 | 1.230 | 0 | 0 | 0 | 0 | 0.050 |
| Timp2 | 0.0045 | 0.095 | 2.344 | 0.679 | -0.041 | 0.168 | 0.088 | -0.165 | 1.054 | 0 | 0 | 0 | 0 | 0.050 |
| Fyb2 | 0.2320 | -0.153 | 0.635 | 0.385 | -0.085 | 0.414 | 0.185 | -0.164 | 0.733 | 0 | 0 | 0 | 0 | 0.050 |
| Tpmt | 0.5289 | -0.053 | 0.277 | 0.134 | -0.131 | 0.872 | 0.054 | -0.163 | 1.268 | 0 | 0 | 0 | 0 | 0.050 |
| Acadsb | 0.2772 | -0.187 | 0.557 | 0.666 | -0.022 | 0.176 | 0.170 | -0.163 | 0.771 | 0 | 0 | 0 | 0 | 0.050 |
| Donson | 0.1190 | -0.092 | 0.925 | 0.231 | 0.082 | 0.636 | 0.271 | -0.163 | 0.567 | 0 | 0 | 0 | 0 | 0.050 |
| Marchf7 | 0.3032 | -0.183 | 0.518 | 0.192 | -0.194 | 0.717 | 0.073 | -0.162 | 1.136 | 0 | 0 | 0 | 0 | 0.050 |
| Rnaseh1 | 0.7569 | -0.021 | 0.121 | 0.669 | 0.072 | 0.175 | 0.149 | -0.161 | 0.826 | 0 | 0 | 0 | 0 | 0.050 |
| Ogfod3 | 0.7544 | 0.032 | 0.122 | 0.482 | -0.090 | 0.317 | 0.067 | -0.160 | 1.171 | 0 | 0 | 0 | 0 | 0.050 |
| Ptma | 0.7296 | -0.050 | 0.137 | 0.702 | 0.030 | 0.154 | 0.227 | -0.159 | 0.645 | 0 | 0 | 0 | 0 | 0.050 |
| Tgfb2 | 0.1938 | -0.059 | 0.713 | 0.104 | -0.145 | 0.982 | 0.088 | -0.159 | 1.055 | 0 | 0 | 0 | 0 | 0.050 |
| Zscan21 | 0.7350 | -0.022 | 0.134 | 0.029 | -0.116 | 1.542 | 0.090 | -0.159 | 1.048 | 0 | 0 | 0 | 0 | 0.050 |
| Gtf2h2 | 0.1915 | -0.107 | 0.718 | 0.407 | -0.111 | 0.391 | 0.161 | -0.159 | 0.792 | 0 | 0 | 0 | 0 | 0.050 |
| Slc7a11 | 0.7758 | -0.039 | 0.110 | 0.046 | -0.135 | 1.341 | 0.156 | -0.158 | 0.806 | 0 | 0 | 0 | 0 | 0.050 |
| Map3k10 | 0.4315 | -0.070 | 0.365 | 0.021 | -0.136 | 1.680 | 0.065 | -0.158 | 1.190 | 0 | 0 | 0 | 0 | 0.050 |
| Lipt2 | 0.2963 | -0.181 | 0.528 | 0.288 | -0.031 | 0.541 | 0.114 | -0.157 | 0.943 | 0 | 0 | 0 | 0 | 0.050 |
| Tcf4 | 0.7072 | -0.072 | 0.150 | 0.097 | -0.185 | 1.014 | 0.208 | -0.157 | 0.682 | 0 | 0 | 0 | 0 | 0.050 |
| Dcps | 0.4743 | -0.032 | 0.324 | 0.453 | -0.076 | 0.344 | 0.062 | -0.157 | 1.206 | 0 | 0 | 0 | 0 | 0.050 |
| Plpp1 | 0.4356 | -0.146 | 0.361 | 0.038 | -0.113 | 1.421 | 0.193 | -0.156 | 0.714 | 0 | 0 | 0 | 0 | 0.050 |
| Fn3krp | 0.5054 | -0.060 | 0.296 | 0.069 | -0.142 | 1.161 | 0.204 | -0.156 | 0.690 | 0 | 0 | 0 | 0 | 0.050 |
| Nek1 | 0.1100 | -0.103 | 0.959 | 0.149 | -0.109 | 0.826 | 0.184 | -0.156 | 0.734 | 0 | 0 | 0 | 0 | 0.050 |
| Zdhhc2 | 0.2213 | -0.199 | 0.655 | 0.209 | -0.063 | 0.680 | 0.085 | -0.154 | 1.072 | 0 | 0 | 0 | 0 | 0.050 |
| Zc2hc1a | 0.0767 | -0.040 | 1.115 | 0.046 | -0.065 | 1.338 | 0.076 | -0.154 | 1.118 | 0 | 0 | 0 | 0 | 0.050 |
| Tm4sf1 | 0.0624 | -0.318 | 1.205 | 0.144 | -0.173 | 0.841 | 0.467 | -0.153 | 0.330 | 0 | 0 | 0 | 0 | 0.050 |
| Mindy2 | 0.9073 | 0.025 | 0.042 | 0.512 | -0.118 | 0.291 | 0.172 | -0.153 | 0.765 | 0 | 0 | 0 | 0 | 0.050 |
| Mrc2 | 0.3686 | -0.025 | 0.433 | 0.111 | -0.186 | 0.954 | 0.147 | -0.153 | 0.833 | 0 | 0 | 0 | 0 | 0.050 |
| Eef1akmt4 | 0.2185 | -0.089 | 0.661 | 0.951 | 0.004 | 0.022 | 0.131 | -0.153 | 0.882 | 0 | 0 | 0 | 0 | 0.050 |
| Pnkd | 0.3555 | -0.053 | 0.449 | 0.620 | -0.056 | 0.208 | 0.152 | -0.152 | 0.817 | 0 | 0 | 0 | 0 | 0.050 |
| Slc38a4 | 0.9958 | 0.000 | 0.002 | 0.093 | -0.119 | 1.031 | 0.079 | -0.152 | 1.100 | 0 | 0 | 0 | 0 | 0.050 |
| Vps72 | 0.0967 | -0.138 | 1.014 | 0.757 | 0.021 | 0.121 | 0.125 | -0.151 | 0.903 | 0 | 0 | 0 | 0 | 0.050 |
| Rad1 | 0.0714 | -0.081 | 1.146 | 0.132 | -0.103 | 0.879 | 0.068 | -0.151 | 1.170 | 0 | 0 | 0 | 0 | 0.050 |
| Psrc1 | 0.3841 | -0.265 | 0.416 | 0.098 | -0.142 | 1.007 | 0.405 | -0.151 | 0.392 | 0 | 0 | 0 | 0 | 0.050 |

**Tab b_Down-regulated:** Contains pathway enrichment analysis (Enrichr) of 244 cell surface proteins significantly down-regulated by glucose starvation in both WT and CT MEFs.

| GO_Biological_Process (down) | Overlap | P-value | Adjusted P-value | Genes |
|---|---|---|---|---|
| Axon Guidance (GO:0007411) | 18/152 | 4.60E-13 | 6.71E-10 | EPHB6;EPHA7;EPHA6;NOTCH1;SEMA3C;UNC5B;RYK;SEMA4B;SEMA4C;PTPRM;UNC5C;EFNA5;EFNB2;EFNB1;SLIT2;EPHB1;EPHA3;EPHB3 |
| Neuron Projection Guidance (GO:0097485) | 16/132 | 6.59E-12 | 4.81E-09 | EPHB6;EPHA7;EPHA6;NOTCH1;SEMA3C;RYK;SEMA4B;SEMA4C;UNC5C;EFNA5;EFNB2;EFNB1;SLIT2;EPHB1;EPHA3;EPHB3 |
| Ephrin Receptor Signaling Pathway (GO:0048013) | 10/43 | 8.18E-11 | 3.85E-08 | EPHB6;EFNB2;EFNB1;EPHA7;EPHA6;EFNA5;EPHB1;EPHB4;EPHA3;EPHB3 |
| Cell Surface Receptor Protein Tyrosine Kinase Signaling Pathway (GO:0007169) | 21/296 | 1.05E-10 | 3.85E-08 | EPHB6;DDR1;PDGFRA;EPHA7;EPHA6;RYK;FLT4;NRG1;EFNA5;EFNB2;GHR;EFNB1;EPHB1;FGFR3;EPHB4;EPHA3;ANGPTL1;MPZL1;EPHB3;DDR2;FGFR1 |
| Cell-Cell Adhesion via Plasma-Membrane Adhesion Molecules (GO:0098742) | 16/171 | 3.40E-10 | 9.93E-08 | KIRREL3;CXADR;VCAM1;PTPRS;PCDHGB4;PCDHGC3;PTPRM;PCDHB14;PVR;FGFRL1;PTPRF;ROBO1;PTPRD;PTK7;NECTIN3;NECTIN2 |
| Axonogenesis (GO:0007409) | 17/200 | 4.24E-10 | 1.03E-07 | EPHB6;EPHA7;EPHA6;NOTCH1;SEMA3C;RYK;APLP2;SEMA4B;SEMA4C;UNC5C;EFNA5;EFNB2;EFNB1;SLIT2;EPHB1;EPHA3;EPHB3 |
| Aortic Valve Development (GO:0003176) | 9/41 | 1.29E-09 | 2.68E-07 | BMPR2;NOTCH1;JAG1;SNAI1;EMILIN1;SLIT2;TNFRSF1B;ROBO1;TNFRSF1A |
| Pulmonary Valve Development (GO:0003177) | 7/21 | 3.71E-09 | 6.77E-07 | BMPR2;JAG1;NOTCH1;SLIT2;TNFRSF1B;ROBO1;TNFRSF1A |
| Negative Chemotaxis (GO:0050919) | 7/25 | 1.47E-08 | 2.38E-06 | EPHA7;SEMA3C;SEMA4B;SEMA4C;UNC5C;SLIT2;ROBO1 |
| Enzyme-Linked Receptor Protein Signaling Pathway (GO:0007167) | 12/121 | 3.00E-08 | 4.38E-06 | PTPRD;DDR1;PDGFRA;BMPR2;RYK;FLT4;NRG1;IL6ST;PTPRF;ANGPTL1;MPZL1;DDR2 |
| Negative Regulation of Extracellular Matrix Organization (GO:1903054) | 5/11 | 1.13E-07 | 1.50E-05 | TGFBR3;NOTCH1;EMILIN1;TNFRSF1B;TNFRSF1A |
| Nervous System Development (GO:0007399) | 21/480 | 4.92E-07 | 5.16E-05 | KIRREL3;IGSF8;PRNP;JAG1;RYK;APLP2;CRIM1;PCDHB14;NRG1;EFNA5;PTPRF;ROBO1;LRP6;PTPRD;ZEB2;ADGRA2;ADGRG6;PLXNA2;EPHB1;ITM2B;CDON |
| Peptidyl-Tyrosine Phosphorylation (GO:0018108) | 8/58 | 4.96E-07 | 5.16E-05 | DDR1;PDGFRA;FLT4;BTK;EPHA3;EGFR;DDR2;FGFR1 |
| Mesenchymal Cell Differentiation (GO:0048762) | 8/58 | 4.96E-07 | 5.16E-05 | TGFBR3;NOTCH1;SEMA3C;SNAI1;SLC39A10;CTNNB1;LRP6;FGFR1 |
| Peptidyl-Tyrosine Modification (GO:0018212) | 7/46 | 1.32E-06 | 1.28E-04 | PDGFRA;FLT4;BTK;EGFR;EPHA3;DDR2;FGFR1 |
| Regulation of Cell Migration (GO:0030334) | 20/479 | 1.91E-06 | 1.68E-04 | IGSF8;PDGFRA;JAG1;NOTCH1;SEMA3C;FLT4;SEMA4B;SEMA4C;PTPRK;EGFR;ROBO1;TGFBR3;SNAI1;PLXNA2;EMILIN2;AMOTL2;EMILIN1;SLIT2;TMSB10;HBEGF |
| Positive Regulation of Homotypic Cell-Cell Adhesion (GO:0034112) | 5/18 | 1.95E-06 | 1.68E-04 | CTNNB1;EMILIN2;EMILIN1;F11R;IL6ST |
| Cell Surface Receptor Signaling Pathway via STAT (GO:0097696) | 7/55 | 4.55E-06 | 3.69E-04 | GHR;IFNAR2;CLCF1;IL6ST;FGFR3;TNFRSF1A;IFNAR1 |
| + Reg of Phosphatidylinositol 3-Kinase/Prot Kinase B Signal Transduction (GO:0051897) | 11/162 | 5.03E-06 | 3.87E-04 | DDR1;PDGFRA;UNC5B;RYK;EFNA5;FGFR3;EGFR;PIK3R5;FGFR1;DDR2;HBEGF |
| Aortic Valve Morphogenesis (GO:0003180) | 6/38 | 6.17E-06 | 4.50E-04 | NOTCH1;JAG1;SNAI1;EMILIN1;SLIT2;ROBO1 |
| Positive Regulation of Platelet Aggregation (GO:1901731) | 4/11 | 6.67E-06 | 4.63E-04 | EMILIN2;EMILIN1;F11R;IL6ST |
| Homophilic Cell Adhesion via Plasma Membrane Adhesion Molecules (GO:0007156) | 7/59 | 7.34E-06 | 4.87E-04 | KIRREL3;PTK7;PTPRM;PVR;NECTIN3;NECTIN2;ROBO1 |
| Positive Regulation of Cell Population Proliferation (GO:0008284) | 19/484 | 8.39E-06 | 5.29E-04 | PDGFRA;NOTCH1;FLT4;NRG1;LAMC1;EGFR;EFNB2;GHR;TGFBR3;CDK4;CLCF1;CTNNB1;S1PR2;IL6ST;FGFR3;HBEGF;DDR2;CRLF2;FGFR1 |
| Protein Autophosphorylation (GO:0046777) | 10/140 | 8.71E-06 | 5.29E-04 | DDR1;PDGFRA;MEX3B;FLT4;EPHB1;EPHB4;EGFR;EPHB3;FGFR1;DDR2 |
| Regulation of Extracellular Matrix Organization (GO:1903053) | 5/24 | 9.12E-06 | 5.32E-04 | DDR1;NOTCH1;LAMC1;NID1;DDR2 |
| Ventricular Septum Morphogenesis (GO:0060412) | 5/28 | 2.03E-05 | 0.001137389 | TGFBR3;BMPR2;NOTCH1;SLIT2;ROBO1 |
| Cell Surface Receptor Signaling Pathway via JAK-STAT (GO:0007259) | 6/47 | 2.19E-05 | 0.001183577 | GHR;IFNAR2;CLCF1;FGFR3;TNFRSF1A;IFNAR1 |
| Positive Regulation of Cellular Process (GO:0048522) | 21/622 | 2.71E-05 | 0.001413836 | PDGFRA;NOTCH1;FLT4;NRG1;LAMC1;NID1;TNFRSF1B;EGFR;LRP6;EFNB2;GHR;CDK4;CLCF1;CTNNB1;AMOTL2;S1PR2;IL6ST;FGFR3;HBEGF;CRLF2;FGFR1 |
| Retinal Ganglion Cell Axon Guidance (GO:0031290) | 3/6 | 3.49E-05 | 0.001715369 | PTPRM;SLIT2;EPHB1 |
| Epithelial to Mesenchymal Transition (GO:0001837) | 6/51 | 3.53E-05 | 0.001715369 | TGFBR3;NOTCH1;SNAI1;SLC39A10;CTNNB1;FGFR1 |
| Positive Regulation of Muscle Cell Differentiation (GO:0051149) | 4/18 | 5.78E-05 | 0.002720888 | NOTCH1;NRG1;LAMC1;NID1 |
| Regulation of Extracellular Matrix Constituent Secretion (GO:0003330) | 3/7 | 6.05E-05 | 0.002760452 | NOTCH1;TNFRSF1B;TNFRSF1A |
| Synapse Organization (GO:0050808) | 9/144 | 7.04E-05 | 0.00299203 | KIRREL3;PTPRD;PTPRS;PTK7;RYK;PLXNA2;PCDHB14;AGRN;PTPRF |
| Response to Interferon-Alpha (GO:0035455) | 4/19 | 7.25E-05 | 0.00299203 | IFITM3;IFNAR2;IFITM2;IFNAR1 |
| Pulmonary Valve Morphogenesis (GO:0003184) | 4/19 | 7.25E-05 | 0.00299203 | JAG1;NOTCH1;SLIT2;ROBO1 |
| Sprouting Angiogenesis (GO:0002040) | 6/58 | 7.38E-05 | 0.00299203 | EFNB2;ADGRA2;FLT4;SLIT2;EPHB4;ROBO1 |
| Regulation of Phosphatidylinositol 3-Kinase/Protein Kinase B Signal Transduction (GO:0051896) | 11/219 | 8.28E-05 | 0.003266034 | DDR1;PDGFRA;UNC5B;RYK;EFNA5;FGFR3;EGFR;PIK3R5;DDR2;HBEGF;FGFR1 |
| Ventricular Septum Development (GO:0003281) | 5/38 | 9.37E-05 | 0.003598501 | TGFBR3;BMPR2;NOTCH1;SLIT2;ROBO1 |
| Canonical Wnt Signaling Pathway (GO:0060070) | 6/63 | 1.18E-04 | 0.004406849 | RYK;FZD6;SNAI1;FZD8;CTNNB1;LRP6 |
| Cell Migration Involved in Sprouting Angiogenesis (GO:0002042) | 4/22 | 1.33E-04 | 0.004851142 | EFNB2;SLIT2;EPHB4;ROBO1 |
| Regulation of Neuron Apoptotic Process (GO:0043523) | 8/129 | 1.88E-04 | 0.006674188 | PRNP;EPHA7;UNC5B;CLCF1;TMBIM1;CTNNB1;IL6ST;MCL1 |

| | | | | |
|---|---|---|---|---|
| Regulation of Cell Population Proliferation (GO:0042127) | 22/769 | 1.97E-04 | 0.006842189 | KLF10;PDGFRA;JAG1;NOTCH1;BMPR2;FLT4;NRG1;PTPRK;EGFR;EFNB2;GHR;CDK4;CLCF1;CTNNB1;EMILIN2;EMILIN1;S1PR2;IL6ST;FGFR3;HBEGF;CRLF2;FGFR1 |
| Regulation of Extracellular Matrix Assembly (GO:1901201) | 3/10 | 2.02E-04 | 0.006854616 | TGFBR3;NOTCH1;EMILIN1 |
| Regulation of Platelet Aggregation (GO:0090330) | 4/26 | 2.62E-04 | 0.008481676 | EMILIN2;EMILIN1;F11R;IL6ST |
| Blood Vessel Endothelial Cell Migration (GO:0043534) | 4/26 | 2.62E-04 | 0.008481676 | EFNB2;SLIT2;EPHB4;ROBO1 |
| **GO_Cellular_Component (down)** | **Overlap** | **P-value** | **Adjusted P-va** | **Genes** |
| Cell-Cell Junction (GO:0005911) | 18/316 | 7.17E-08 | 9.39E-06 | KIRREL3;KIRREL1;CXADR;JAG1;NOTCH1;BMPR2;HEG1;PTPRM;PTPRK;F11R;PVR;FGFRL1;PDLIM2;PTK7;FAT1;CTNNB1;NECTIN3;NECTIN2 |
| Adherens Junction (GO:0005912) | 10/151 | 1.69E-05 | 0.001108256 | CXADR;NOTCH1;JAG1;BMPR2;PDLIM2;PTPRM;CTNNB1;PVR;NECTIN3;NECTIN2 |
| Focal Adhesion (GO:0005925) | 14/387 | 3.05E-04 | 0.012274365 | LAYN;PVR;EGFR;EFNB2;IL1RL1;ADGRE5;PTK7;TSPAN4;FAT1;CTNNB1;FGFR3;MPZL1;NECTIN2;DDR2 |
| Cell-Substrate Junction (GO:0030055) | 14/395 | 3.75E-04 | 0.012274365 | LAYN;PVR;EGFR;EFNB2;IL1RL1;ADGRE5;PTK7;TSPAN4;FAT1;CTNNB1;FGFR3;MPZL1;NECTIN2;DDR2 |
| Cul3-RING Ubiquitin Ligase Complex (GO:0031463) | 5/58 | 7.02E-04 | 0.018389854 | KLHL9;KCTD10;KLHL11;TNFAIP1;KLHL13 |
| Collagen-Containing Extracellular Matrix (GO:0062023) | 13/383 | 8.99E-04 | 0.019630165 | LAMA5;BGN;LAMC1;NID1;BCAM;LGALS1;CLCF1;ANGPTL2;EMILIN2;EMILIN1;AGRN;ANGPTL1;CDON |
| Multivesicular Body, Internal Vesicle (GO:0097487) | 2/5 | 0.00144679 | 0.027075643 | CD63;EGFR |
| Basement Membrane (GO:0005604) | 4/50 | 0.003206373 | 0.052504352 | LAMA5;LAMC1;NID1;AGRN |
| Membrane Raft (GO:0045121) | 7/171 | 0.005102291 | 0.066554015 | PRNP;CXADR;BTK;IL6ST;TNFRSF1B;EGFR;TNFRSF1A |
| Filopodium (GO:0030175) | 4/57 | 0.005146994 | 0.066554015 | CXADR;VCAM1;UNC5C;ANTXR1 |
| cullin-RING Ubiquitin Ligase Complex (GO:0031461) | 8/219 | 0.005588505 | 0.066554015 | KLHL9;KCTD10;KLHL11;FEM1C;KLHL13;TNFAIP1;DCAF7;ANAPC10 |
| Apical Junction Complex (GO:0043296) | 5/107 | 0.010074411 | 0.109978983 | CXADR;AMOTL2;F11R;NECTIN3;NECTIN2 |
| Lysosomal Lumen (GO:0043202) | 4/86 | 0.021072687 | 0.185083261 | SDC4;BGN;AGRN;GPC6 |
| Multivesicular Body Membrane (GO:0032585) | 2/19 | 0.02211166 | 0.185083261 | CD63;CHMP1A |
| Dendrite (GO:0030425) | 8/281 | 0.022407911 | 0.185083261 | EPHB6;KIRREL3;PRNP;EPHA7;EPHA6;EPHB1;EPHA3;EPHB3 |
| Endosome Membrane (GO:0010008) | 10/392 | 0.022605589 | 0.185083261 | IFITM3;CD63;IFITM2;NOTCH1;TMBIM1;TAB3;ANTXR1;EPHB1;EGFR;LRP6 |
| Podosome (GO:0002102) | 2/20 | 0.024373661 | 0.187820565 | VCAM1;AMOTL2 |
| Lysosome (GO:0005764) | 12/532 | 0.030724483 | 0.20647864 | IFITM3;CD63;IFITM2;AP5M1;SDC4;CHMP1A;SPPL2A;TMBIM1;BGN;AGRN;GPC6;VASN |
| Golgi-associated Vesicle Membrane (GO:0030660) | 2/23 | 0.031690919 | 0.20647864 | SPPL2A;ITM2B |
| Cytoplasmic Vesicle Membrane (GO:0030659) | 10/415 | 0.031716337 | 0.20647864 | CD63;NOTCH1;ADGRE5;SPPL2A;TMBIM1;TAB3;ANTXR1;ITM2B;EGFR;HBEGF |
| Golgi Lumen (GO:0005796) | 4/99 | 0.033099629 | 0.20647864 | SDC4;BGN;AGRN;GPC6 |
| Glutamatergic Synapse (GO:0098978) | 4/102 | 0.036341235 | 0.216395535 | EFNB2;PTPRD;EFNB1;NRG1 |
| Early Endosome (GO:0005769) | 8/320 | 0.0432122 | 0.246121663 | IFITM3;VCAM1;LIPG;CHMP1A;EPHB1;EPHA3;EGFR;LRP6 |
| **GO_Molecular_Function (down)** | **Overlap** | **P-value** | **Adjusted P-va** | **Genes** |
| Transmembrane Receptor Protein Tyrosine Kinase Activity (GO:0004714) | 14/50 | 7.06E-16 | 1.63E-13 | DDR1;EPHB6;PDGFRA;RYK;FLT4;CRIM1;FGFRL1;EGFR;FGFR3;EPHB4;EPHA3;EPHB3;DDR2;FGFR1 |
| Ephrin Receptor Activity (GO:0005003) | 7/15 | 2.19E-10 | 2.52E-08 | EPHB6;EPHA7;EPHA6;EPHB1;EPHA3;EPHB4;EPHB3 |
| Transmembrane-Ephrin Receptor Activity (GO:0005005) | 6/14 | 8.58E-09 | 6.61E-07 | EPHB6;EPHA7;EPHA6;EPHB1;EPHA3;EPHB3 |
| Protein Tyrosine Kinase Activity (GO:0004713) | 11/102 | 4.77E-08 | 2.76E-06 | DDR1;PDGFRA;EPHA7;RYK;FLT4;BTK;FGFR3;EPHB4;EGFR;FGFR1;DDR2 |
| Transmembrane Receptor Protein Kinase Activity (GO:0019199) | 7/37 | 2.78E-07 | 1.28E-05 | DDR1;PDGFRA;RYK;FLT4;EGFR;EPHB4;DDR2 |
| Chemorepellent Activity (GO:0045499) | 5/24 | 9.12E-06 | 3.51E-04 | EPHA7;SEMA3C;SEMA4B;SEMA4C;NRG1 |
| Wnt Receptor Activity (GO:0042813) | 4/14 | 1.97E-05 | 6.49E-04 | RYK;FZD6;FZD8;LRP6 |
| Transmembrane Receptor Protein Phosphatase Activity (GO:0019198) | 4/15 | 2.65E-05 | 6.66E-04 | PTPRD;PTPRM;PTPRK;PTPRF |
| Transmembrane Receptor Protein Tyrosine Phosphatase Activity (GO:0005001) | 4/15 | 2.65E-05 | 6.66E-04 | PTPRD;PTPRM;PTPRK;PTPRF |
| Protein Tyrosine Kinase Activator Activity (GO:0030296) | 5/30 | 2.88E-05 | 6.66E-04 | NRG1;EFNA5;IL6ST;EGFR;HBEGF |
| Cytokine Receptor Activity (GO:0004896) | 7/83 | 6.97E-05 | 0.00146305 | GHR;IL1RL1;IL4R;IL1RAP;IL6ST;IL17RA;CRLF2 |
| PDZ Domain Binding (GO:0030165) | 5/58 | 7.02E-04 | 0.013010216 | TGFBR3;CXADR;CRIM1;FZD8;F11R |
| Transmembrane Receptor Protein Tyrosine Kinase Activator Activity (GO:0030297) | 3/15 | 7.32E-04 | 0.013010216 | NRG1;EFNA5;HBEGF |
| Neuropilin Binding (GO:0038191) | 3/17 | 0.001074694 | 0.017732449 | SEMA3C;SEMA4B;SEMA4C |
| Cell Adhesion Mediator Activity (GO:0098631) | 4/39 | 0.001271294 | 0.019577923 | LAMA5;BCAM;VCAM1;NECTIN3 |
| Vascular Endothelial Growth Factor Receptor Activity (GO:0005021) | 2/5 | 0.00144679 | 0.020888032 | PDGFRA;FLT4 |
| Fibroblast Growth Factor Binding (GO:0017134) | 3/20 | 0.001753698 | 0.023829663 | FGFR3;FGFRL1;FGFR1 |
| Tumor Necrosis Factor Binding (GO:0043120) | 2/6 | 0.002152734 | 0.025595149 | TNFRSF1B;TNFRSF1A |
| Axon Guidance Receptor Activity (GO:0008046) | 2/6 | 0.002152734 | 0.025595149 | PTK7;ROBO1 |
| Ephrin Receptor Binding (GO:0046875) | 3/22 | 0.002326832 | 0.025595149 | EFNB2;EFNB1;EFNA5 |
| Semaphorin Receptor Binding (GO:0030215) | 3/22 | 0.002326832 | 0.025595149 | SEMA3C;SEMA4B;SEMA4C |
| Receptor Ligand Activity (GO:0048018) | 12/380 | 0.002558318 | 0.026862336 | LAMA5;EPHA7;LGALS1;JAG1;SEMA3C;CLCF1;SEMA4B;SEMA4C;NRG1;PVR;NECTIN2;HBEGF |
| Protein Tyrosine Phosphatase Activity (GO:0004725) | 5/82 | 0.003300657 | 0.032559796 | PTPRD;PTPRS;PTPRM;PTPRK;PTPRF |

| | | | | |
|---|---|---|---|---|
| Alcohol Dehydrogenase (NADP+) Activity (GO:0008106) | 3/25 | 0.003382836 | 0.032559796 | AKR7A2;RDH11;AKR1B1 |
| Tumor Necrosis Factor Receptor Activity (GO:0005031) | 2/8 | 0.003954158 | 0.036536416 | TNFRSF1B;TNFRSF1A |
| Volume-Sensitive Anion Channel Activity (GO:0005225) | 2/9 | 0.005043151 | 0.044806454 | TTYH3;TTYH2 |
| Aldose Reductase (NADPH) Activity (GO:0004032) | 2/10 | 0.006253438 | 0.053501639 | AKR7A2;AKR1B1 |
| Transforming Growth Factor Beta Receptor Activity (GO:0005024) | 2/11 | 0.007581924 | 0.060393943 | TGFBR3;BMPR2 |
| Death Receptor Activity (GO:0005035) | 2/11 | 0.007581924 | 0.060393943 | TNFRSF1B;TNFRSF1A |
| Protein Homodimerization Activity (GO:0042803) | 16/680 | 0.009542758 | 0.073479237 | PDGFRA;PRNP;HMGCS1;FLT4;SPPL2A;AHR;F11R;LRP6;GHR;MKLN1;IMPA2;CHMP1A;SLIT2;NECTIN3;NECTIN2;FGFR1 |
| Phospholipase Activator Activity (GO:0016004) | 2/15 | 0.014017875 | 0.10445578 | PDGFRA;BTK |
| Transmembrane Receptor Protein Serine/Threonine Kinase Activity (GO:0004675) | 2/17 | 0.017868616 | 0.12898907 | TGFBR3;BMPR2 |
| Ligand-Gated Monoatomic Anion Channel Activity (GO:0099095) | 3/47 | 0.019606089 | 0.135491279 | TTYH3;TTYH2;PACC1 |
| BMP Binding (GO:0036122) | 2/18 | 0.019942439 | 0.135491279 | TGFBR3;BMPR2 |
| Intracellularly Calcium-Gated Channel Activity (GO:0141147) | 2/19 | 0.02211166 | 0.141883151 | TTYH3;TTYH2 |
| Intracellularly Calcium-Gated Chloride Channel Activity (GO:0005229) | 2/19 | 0.02211166 | 0.141883151 | TTYH3;TTYH2 |
| Growth Factor Activity (GO:0008083) | 4/92 | 0.026219505 | 0.163694749 | JAG1;CLCF1;NRG1;HBEGF |
| Transforming Growth Factor Beta Binding (GO:0050431) | 2/22 | 0.029165786 | 0.177297276 | TGFBR3;VASN |
| Notch Binding (GO:0005112) | 2/23 | 0.031690919 | 0.187707753 | JAG1;KCTD10 |
| Growth Factor Receptor Binding (GO:0070851) | 4/100 | 0.034160535 | 0.19727709 | PDGFRA;IL1RN;IL6ST;HBEGF |
| Cytokine Receptor Binding (GO:0005126) | 4/109 | 0.04459366 | 0.251247206 | TGFBR3;IL1RN;CLCF1;IL6ST |

**Tab c_Up-regulated:** Contains pathway enrichment analysis (Enrichr) of 95 cell surface proteins significantly up-regulated by glucose starvation in both WT and CT MEFs.

| GO_Molecular_Process (up) | Overlap | P-value | Adjusted P-value | Genes |
|---|---|---|---|---|
| Nicotinamide Nucleotide Metabolic Process (GO:0046496) | 6/51 | 1.48E-07 | 4.55E-05 | PKM;PKLR;HKDC1;PGK1;PGK2;GPD1L |
| ADP Catabolic Process (GO:0046032) | 5/28 | 1.96E-07 | 4.55E-05 | PKM;PKLR;HKDC1;PGK1;PGK2 |
| Glycolytic Process (GO:0006096) | 5/28 | 1.96E-07 | 4.55E-05 | PKM;PKLR;HKDC1;PGK1;PGK2 |
| Pyridine Nucleotide Catabolic Process (GO:0019364) | 5/32 | 3.96E-07 | 6.77E-05 | PKM;PKLR;HKDC1;PGK1;PGK2 |
| ATP Metabolic Process (GO:0046034) | 6/62 | 4.86E-07 | 6.77E-05 | PKM;PKLR;HKDC1;PGK1;ATP5PO;PGK2 |
| Carbohydrate Catabolic Process (GO:0016052) | 5/40 | 1.25E-06 | 1.46E-04 | PKM;PKLR;HKDC1;PGK1;PGK2 |
| Pyruvate Metabolic Process (GO:0006090) | 5/48 | 3.17E-06 | 3.15E-04 | PKM;PKLR;HKDC1;PGK1;PGK2 |
| Purine Ribonucleoside Triphosphate Biosynthetic Process (GO:0009206) | 3/11 | 1.67E-05 | 1.45E-03 | IMPDH1;IMPDH2;ATP5PO |
| Primary miRNA Processing (GO:0031053) | 3/12 | 2.21E-05 | 1.71E-03 | SRRT;DGCR8;SRSF3 |
| Polarized Epithelial Cell Differentiation (GO:0030859) | 3/14 | 3.64E-05 | 2.54E-03 | RAB10;SCRIB;RHOA |
| mRNA Splicing, via Spliceosome (GO:0000398) | 7/211 | 6.63E-05 | 4.20E-03 | SYNCRIP;PRMT5;HNRNPUL2;SRSF3;RBM42;PPIH;WDR77 |
| Regulation of mRNA Splicing, via Spliceosome (GO:0048024) | 5/97 | 9.92E-05 | 5.76E-03 | PRMT5;C1QBP;SRSF3;NSRP1;WDR77 |
| Neurotransmitter Receptor Transport, Endosome to Plasma Membrane (GO:0099639) | 2/5 | 2.21E-04 | 1.19E-02 | SCRIB;RAB11A |
| NADH Metabolic Process (GO:0006734) | 3/26 | 2.49E-04 | 1.24E-02 | PGK1;PGK2;GPD1L |
| Neurotransmitter Receptor Transport to Postsynaptic Membrane (GO:0098969) | 2/6 | 3.31E-04 | 1.54E-02 | SCRIB;RAB11A |
| Golgi Organization (GO:0007030) | 5/128 | 3.63E-04 | 1.58E-02 | RAB2A;RAB2B;PRMT5;SPTBN5;GORASP2 |
| miRNA Processing (GO:0035196) | 3/32 | 4.66E-04 | 1.91E-02 | SRRT;DGCR8;SRSF3 |
| Autophagosome-Lysosome Fusion (GO:0061909) | 2/8 | 6.14E-04 | 2.38E-02 | PIP4K2A;PIP4K2B |
| Astral Microtubule Organization (GO:0030953) | 2/9 | 7.86E-04 | 2.89E-02 | PDE4DIP;RAB11A |
| Establishment of Protein Localization to Plasma Membrane (GO:0061951) | 3/41 | 9.71E-04 | 3.25E-02 | RAB10;GORASP2;RAB11A |
| Establishment of Endothelial Intestinal Barrier (GO:0090557) | 2/10 | 9.80E-04 | 3.25E-02 | RAP2B;TJP2 |
| Macroautophagy (GO:0016236) | 5/162 | 1.06E-03 | 3.35E-02 | RAB2A;PIP4K2A;PIP4K2B;ARHGAP26;CHMP7 |
| Protein Localization to Endosome (GO:0036010) | 2/11 | 1.19E-03 | 3.62E-02 | TOLLIP;RAB35 |
| Purine Ribonucleotide Biosynthetic Process (GO:0009152) | 3/45 | 1.27E-03 | 3.70E-02 | IMPDH1;IMPDH2;ATP5PO |
| Morphogenesis of a Polarized Epithelium (GO:0001738) | 2/12 | 1.43E-03 | 3.98E-02 | RAB10;SCRIB |
| Establishment of Apical/Basal Cell Polarity (GO:0035089) | 2/14 | 1.96E-03 | 4.62E-02 | SCRIB;RHOA |
| GTP Metabolic Process (GO:0046039) | 2/14 | 1.96E-03 | 4.62E-02 | IMPDH1;IMPDH2 |
| Negative Regulation of Transcription Elongation by RNA Polymerase II (GO:0034244) | 2/14 | 1.96E-03 | 4.62E-02 | TCERG1;NELFA |
| Small GTPase-mediated Signal Transduction (GO:0007264) | 5/188 | 2.04E-03 | 4.62E-02 | RAP2B;KRIT1;PLCE1;RHOA;REPS1 |
| Cytoplasmic Microtubule Organization (GO:0031122) | 3/53 | 2.05E-03 | 4.62E-02 | PDE4DIP;RHOA;RAB11A |
| Cellular Component Assembly (GO:0022607) | 6/277 | 2.08E-03 | 4.62E-02 | NIFK;GORASP2;CALD1;PDE4DIP;PPIH;RHOA |
| Glucose Catabolic Process to Pyruvate (GO:0061718) | 2/15 | 2.25E-03 | 4.62E-02 | PGK1;PGK2 |
| Canonical Glycolysis (GO:0061621) | 2/15 | 2.25E-03 | 4.62E-02 | PGK1;PGK2 |
| Positive Regulation of Cytoplasmic Translation (GO:2000767) | 2/15 | 2.25E-03 | 4.62E-02 | SYNCRIP;PKM |
| Glucose Metabolic Process (GO:0006006) | 3/57 | 2.52E-03 | 4.83E-02 | HKDC1;PGK1;PGK2 |
| Glycolytic Process Through Glucose-6-Phosphate (GO:0061620) | 2/16 | 2.57E-03 | 4.83E-02 | PGK1;PGK2 |
| Protein Localization to Cell-Cell Junction (GO:0150105) | 2/16 | 2.57E-03 | 4.83E-02 | SCRIB;TJP2 |
| **GO_Cellular_Component (up)** | **Overlap** | **P-value** | **Adjusted P-value** | **Genes** |
| Ficolin-1-Rich Granule (GO:0101002) | 6/184 | 2.47E-04 | 1.84E-02 | CSTB;PKM;IMPDH1;IMPDH2;PLEKHO2;RHOA |
| Ficolin-1-Rich Granule Lumen (GO:1904813) | 5/123 | 3.02E-04 | 1.84E-02 | CSTB;PKM;IMPDH1;IMPDH2;PLEKHO2 |
| Cell-Cell Junction (GO:0005911) | 7/316 | 7.75E-04 | 3.15E-02 | RAB10;RAP2B;MAGI3;KRIT1;SCRIB;RHOA;TJP2 |
| Cytoplasmic Vesicle Lumen (GO:0060205) | 4/117 | 2.35E-03 | 7.14E-02 | CSTB;PKM;IMPDH1;IMPDH2 |
| Kinetochore Microtubule (GO:0005828) | 2/18 | 3.25E-03 | 7.14E-02 | CHMP7;RAB11A |
| Pericentric Heterochromatin (GO:0005721) | 2/21 | 4.42E-03 | 7.14E-02 | CBX5;BAZ1B |
| Recycling Endosome Membrane (GO:0055038) | 3/70 | 4.52E-03 | 7.14E-02 | RAP2B;RAB35;RAB11A |
| Nucleolus (GO:0005730) | 10/805 | 4.88E-03 | 7.14E-02 | CSTB;RPS25;CBX5;NIFK;PRKRIP1;CTCF;ANP32B;BAZ1B;RPF2;GNL3 |
| Nuclear Lumen (GO:0031981) | 10/814 | 5.27E-03 | 7.14E-02 | CSTB;RPS25;CBX5;NIFK;PRKRIP1;CTCF;ANP32B;BAZ1B;RPF2;GNL3 |
| Recycling Endosome (GO:0055037) | 4/155 | 6.40E-03 | 7.52E-02 | RAB10;RAP2B;RAB35;RAB11A |
| Autophagosome (GO:0005776) | 3/81 | 6.78E-03 | 7.52E-02 | PIP4K2A;PIP4K2B;CHMP7 |
| Chromosome (GO:0005694) | 4/179 | 1.05E-02 | 8.61E-02 | NIFK;CTCF;RPF2;GNL3 |
| Endosome Membrane (GO:0010008) | 6/392 | 1.10E-02 | 8.61E-02 | RAB10;RAP2B;RAB15;RAB35;ARHGAP26;RAB11A |
| Intracellular Organelle Lumen (GO:0070013) | 10/912 | 1.13E-02 | 8.61E-02 | CSTB;PKM;GLDC;IMPDH1;C1QBP;IMPDH2;PGK1;PLEKHO2;LIAS;PDIA5 |
| U4/U6 X U5 tri-snRNP Complex (GO:0046540) | 2/35 | 1.20E-02 | 8.61E-02 | RBM42;PPIH |
| Spliceosomal tri-snRNP Complex (GO:0097526) | 2/35 | 1.20E-02 | 8.61E-02 | RBM42;PPIH |

| | | | | |
|---|---|---|---|---|
| Phagocytic Vesicle (GO:0045335) | 3/100 | 1.20E-02 | 8.61E-02 | NCF2;SRGAP2;RAB11A |
| Glutamatergic Synapse (GO:0098978) | 3/102 | 1.27E-02 | 8.61E-02 | TANC1;RHOA;RAB11A |
| Cytoplasmic Vesicle Membrane (GO:0030659) | 6/415 | 1.42E-02 | 8.81E-02 | RAB10;RAB15;RAB35;ARHGAP26;RHOA;RAB11A |
| Apical Junction Complex (GO:0043296) | 3/107 | 1.44E-02 | 8.81E-02 | RAP2B;RHOA;TJP2 |
| Secretory Granule Lumen (GO:0034774) | 5/316 | 1.74E-02 | 1.01E-01 | CSTB;PKM;IMPDH1;IMPDH2;TOLLIP |
| Ribosome (GO:0005840) | 2/46 | 2.02E-02 | 1.07E-01 | RPS25;MRPS34 |
| Intracellular Membraneless Organelle (GO:0043232) | 12/1310 | 2.11E-02 | 1.07E-01 | CSTB;RPS25;CBX5;NIFK;CALD1;PRKRIP1;MRPS34;CTCF;ANP32B;BAZ1B;RPF2;GNL3 |
| Cell-Cell Contact Zone (GO:0044291) | 2/48 | 2.19E-02 | 1.07E-01 | RAP2B;TJP2 |
| Multivesicular Body (GO:0005771) | 2/49 | 2.27E-02 | 1.07E-01 | CHMP7;RAB11A |
| Phagolysosome (GO:0032010) | 1/5 | 2.35E-02 | 1.07E-01 | NCF2 |
| Bounding Membrane of Organelle (GO:0098588) | 9/881 | 2.41E-02 | 1.07E-01 | RAB2A;RAB10;RAB2B;GORASP2;RAB15;RAB35;ARHGAP26;RHOA;RAB11A |
| Asymmetric Synapse (GO:0032279) | 3/131 | 2.46E-02 | 1.07E-01 | RPS25;SCRIB;SRGAP2 |
| Insulin-Responsive Compartment (GO:0032593) | 1/6 | 2.82E-02 | 1.18E-01 | RAB10 |
| Condensed Chromosome (GO:0000793) | 2/62 | 3.51E-02 | 1.37E-01 | NIFK;CTCF |
| Transferase Complex, Transferring Phosphorus-Containing Groups (GO:0061695) | 1/8 | 3.74E-02 | 1.37E-01 | PKLR |
| Intracellular Membrane-Bounded Organelle (GO:0043231) | 35/5597 | 3.74E-02 | 1.37E-01 | TCERG1;CSTB;WIZ;CTCF;CLINT1;SYNCRIP;ALKBH5;C1QBP;DGCR8;PIP4K2B;NELFA;PGK2;REPS1;RAB2A;PRMT5;CBX5;TSC22D1;PDE4DIP;BAZ1B;WDR77;GNL3;RPS25;PKM;PER3;HNRNPUL2;GORASP2;IMPDH1;IMPDH2;ATP5PO;RBM42;POLR3G;PPIH;ANP32B;NSRP1;TJP2 |
| Specific Granule (GO:0042581) | 3/159 | 4.01E-02 | 1.37E-01 | PGRMC1;RAP2B;TOLLIP |
| Amphisome Membrane (GO:1904930) | 1/9 | 4.20E-02 | 1.37E-01 | CHMP7 |
| Tertiary Granule (GO:0070820) | 3/163 | 4.27E-02 | 1.37E-01 | CSTB;RAP2B;RHOA |
| Nucleus (GO:0005634) | 31/4879 | 4.28E-02 | 1.37E-01 | TCERG1;CSTB;WIZ;CTCF;SYNCRIP;ALKBH5;C1QBP;DGCR8;PIP4K2B;NELFA;PGK2;RAB2A;PRMT5;CBX5;TSC22D1;PDE4DIP;BAZ1B;WDR77;GNL3;RPS25;PKM;PER3;HNRNPUL2;IMPDH1;IMPDH2;ATP5PO;RBM42;POLR3G;ANP32B;NSRP1;TJP2 |
| Postsynaptic Density (GO:0014069) | 3/164 | 4.33E-02 | 1.37E-01 | RPS25;SCRIB;SRGAP2 |
| Secretory Granule Membrane (GO:0030667) | 4/279 | 4.40E-02 | 1.37E-01 | RAB10;PGRMC1;RAP2B;RHOA |
| Mitochondrial Matrix (GO:0005759) | 5/407 | 4.46E-02 | 1.37E-01 | GLDC;C1QBP;MRPS34;PGK1;LIAS |
| Spindle Microtubule (GO:0005876) | 2/71 | 4.49E-02 | 1.37E-01 | CHMP7;RAB11A |
| **Go_Molecular_Function (up)** | **Overlap** | **P-value** | **Adjusted P-value** | **Genes** |
| GTP Binding (GO:0005525) | 9/206 | 5.99E-07 | 7.37E-05 | RAB2A;RAB10;RAB2B;RAP2B;RAB35;PIP4K2B;RHOA;RAB11A;GNL3 |
| Guanyl Ribonucleotide Binding (GO:0032561) | 9/232 | 1.61E-06 | 9.90E-05 | RAB2A;RAB10;RAB2B;RAP2B;RAB35;PIP4K2B;RHOA;RAB11A;GNL3 |
| mRNA Binding (GO:0003729) | 8/310 | 1.15E-04 | 4.72E-03 | SYNCRIP;PKM;STAU2;C1QBP;SRSF3;RBM42;NSRP1;GNL3 |
| Myosin Binding (GO:0017022) | 4/57 | 1.56E-04 | 4.78E-03 | RAB10;CALD1;RHOA;RAB11A |
| GDP Binding (GO:0019003) | 4/66 | 2.75E-04 | 6.76E-03 | RAB2A;RAB10;RAP2B;RAB35 |
| Primary miRNA Binding (GO:0070878) | 2/6 | 3.31E-04 | 6.78E-03 | DGCR8;SRSF3 |
| 1-Phosphatidylinositol-4-Phosphate 5-Kinase Activity (GO:0016308) | 2/7 | 4.62E-04 | 8.11E-03 | PIP4K2A;PIP4K2B |
| Guanylate Kinase Activity (GO:0004385) | 2/8 | 6.14E-04 | 9.00E-03 | MAGI3;TJP2 |
| Anion Binding (GO:0043168) | 8/402 | 6.58E-04 | 9.00E-03 | RAB2A;RAB10;RAP2B;GLDC;RAB35;PGK1;KRIT1;PGK2 |
| Cadherin Binding (GO:0045296) | 7/317 | 7.89E-04 | 9.71E-03 | RAB10;PLCB3;PKM;CALD1;SCRIB;CLINT1;TJP2 |
| GTPase Activity (GO:0003924) | 6/266 | 1.69E-03 | 1.86E-02 | RAB2A;RAB2B;RAP2B;RAB35;RHOA;RAB11A |
| Myosin V Binding (GO:0031489) | 2/14 | 1.96E-03 | 1.86E-02 | RAB10;RAB11A |
| Ribonucleoside Triphosphate Phosphatase Activity (GO:0017111) | 6/274 | 1.97E-03 | 1.86E-02 | RAB2A;RAB2B;RAP2B;RAB35;RHOA;RAB11A |
| Nucleoside Monophosphate Kinase Activity (GO:0050145) | 2/18 | 3.25E-03 | 2.86E-02 | MAGI3;TJP2 |
| Oxidoreductase Activity, Acting on the CH-OH Group of Donors, NAD or NADP as Acceptor (GO:0016616) | 3/70 | 4.52E-03 | 3.43E-02 | IMPDH1;IMPDH2;GPD1L |
| Double-Stranded RNA Binding (GO:0003725) | 3/71 | 4.70E-03 | 3.43E-02 | STAU2;PRKRIP1;DGCR8 |
| mRNA 5'-UTR Binding (GO:0048027) | 2/22 | 4.85E-03 | 3.43E-02 | SYNCRIP;GNL3 |
| Protein Homodimerization Activity (GO:0042803) | 9/680 | 5.02E-03 | 3.43E-02 | PGRMC1;GALE;PKM;GLDC;GLMN;PIP4K2A;DGCR8;PIP4K2B;SRGAP2 |
| Phosphatidylinositol Kinase Activity (GO:0052742) | 2/24 | 5.76E-03 | 3.54E-02 | PIP4K2A;PIP4K2B |
| Phosphatidylinositol Phospholipase C Activity (GO:0004435) | 2/24 | 5.76E-03 | 3.54E-02 | PLCB3;PLCE1 |
| Phospholipase C Activity (GO:0004629) | 2/26 | 6.74E-03 | 3.95E-02 | PLCB3;PLCE1 |
| ADP Binding (GO:0043531) | 2/28 | 7.79E-03 | 4.35E-02 | PGK1;PGK2 |
| methyl-CpG Binding (GO:0008327) | 2/29 | 8.34E-03 | 4.46E-02 | PRMT5;WDR77 |
| Ubiquitin-Like Protein Conjugating Enzyme Binding (GO:0044390) | 2/34 | 1.13E-02 | 5.81E-02 | TCERG1;TOLLIP |
| Transcription Coregulator Binding (GO:0001221) | 3/103 | 1.30E-02 | 6.42E-02 | PER3;WIZ;CTCF |
| Phosphoric Diester Hydrolase Activity (GO:0008081) | 2/44 | 1.86E-02 | 8.78E-02 | PLCB3;PLCE1 |
| Transcription Corepressor Binding (GO:0001222) | 2/46 | 2.02E-02 | 9.19E-02 | PER3;WIZ |
| Sulfurtransferase Activity (GO:0016783) | 1/7 | 3.28E-02 | 1.44E-01 | LIAS |
| 5S rRNA Binding (GO:0008097) | 1/8 | 3.74E-02 | 1.48E-01 | RPF2 |

| | | | | |
|---|---|---|---|---|
| Superoxide-Generating NADPH Oxidase Activator Activity (GO:0016176) | 1/8 | 3.74E-02 | 1.48E-01 | NCF2 |
| Arginine N-methyltransferase Activity (GO:0016273) | 1/8 | 3.74E-02 | 1.48E-01 | PRMT5 |
| Filamin Binding (GO:0031005) | 1/10 | 4.65E-02 | 1.51E-01 | CRMP1 |
| Histone H3 Kinase Activity (GO:0140996) | 1/10 | 4.65E-02 | 1.51E-01 | PKM |
| Racemase and Epimerase Activity, Acting on Carbohydrates and Derivatives (GO:0016857) | 1/10 | 4.65E-02 | 1.51E-01 | GALE |
| Dipeptidyl-Peptidase Activity (GO:0008239) | 1/10 | 4.65E-02 | 1.51E-01 | GLMN |
| Myosin Heavy Chain Binding (GO:0032036) | 1/10 | 4.65E-02 | 1.51E-01 | SPTBN5 |

**Tab d_Sec24c-sependent down:** Contains pathway enrichment analysis (DAVID, v2023q4) of cell surface proteins whose down-regulation in response to glucose starvation is dependent on Sec24c.

| GO_Biological_Process (down) | Overlap | P-value | Adjusted P-value | Genes |
|---|---|---|---|---|
| Cell-Cell Adhesion Via Plasma-Membrane Adhesion Molecules (GO:0098742) | 9/172 | 3.31E-09 | 1.95E-06 | CDH3;CXADR;CDH1;PTK7;PTPRM;NECTIN3;PTPRF;NECTIN2;ROBO1 |
| Homophilic Cell Adhesion Via Plasma Membrane Adhesion Molecules (GO:0007156) | 6/60 | 3.36E-08 | 9.86E-06 | CDH1;PTK7;PTPRM;NECTIN3;NECTIN2;ROBO1 |
| Ephrin Receptor Signaling Pathway (GO:0048013) | 4/43 | 1.01E-05 | 1.97E-03 | EFNB2;EFNB1;EPHB4;EPHB3 |
| Cell Migration Involved In Sprouting Angiogenesis (GO:0002042) | 3/17 | 1.96E-05 | 2.31E-03 | EFNB2;EPHB4;ROBO1 |
| Response To Interferon-Alpha (GO:0035455) | 3/17 | 1.96E-05 | 2.31E-03 | IFITM3;IFNAR2;IFITM2 |
| Blood Vessel Endothelial Cell Migration (GO:0043534) | 3/22 | 4.40E-05 | 4.31E-03 | EFNB2;EPHB4;ROBO1 |
| Response To Interferon-Beta (GO:0035456) | 3/29 | 1.03E-04 | 8.40E-03 | IFITM3;IFNAR2;IFITM2 |
| Positive Regulation Of Notch Signaling Pathway (GO:0045747) | 3/33 | 1.52E-04 | 8.40E-03 | TSPAN14;JAG1;ROBO1 |
| Positive Regulation Of Neuron Death (GO:1901216) | 3/34 | 1.66E-04 | 8.40E-03 | EFNB2;PRNP;CTNNB1 |
| Adherens Junction Organization (GO:0034332) | 3/35 | 1.81E-04 | 8.40E-03 | CDH3;CDH1;CTNNB1 |
| Positive Regulation Of Signal Transduction (GO:0009967) | 6/266 | 1.88E-04 | 8.40E-03 | RIMS1;TSPAN14;CDH3;JAG1;LAMC1;ROBO1 |
| Substrate Adhesion-Dependent Cell Spreading (GO:0034446) | 3/36 | 1.97E-04 | 8.40E-03 | LAMA5;LAMC1;EPHB3 |
| Positive Regulation Of Melanin Biosynthetic Process (GO:0048023) | 2/7 | 2.03E-04 | 8.40E-03 | ZEB2;CDH3 |
| Positive Regulation Of Secondary Metabolite Biosynthetic Process (GO:1900378) | 2/7 | 2.03E-04 | 8.40E-03 | ZEB2;CDH3 |
| Type I Interferon-Mediated Signaling Pathway (GO:0060337) | 3/37 | 2.14E-04 | 8.40E-03 | IFITM3;IFNAR2;IFITM2 |
| Cellular Response To Type I Interferon (GO:0071357) | 3/38 | 2.32E-04 | 8.54E-03 | IFITM3;IFNAR2;IFITM2 |
| Regulation Of Cell-Cell Adhesion (GO:0022407) | 3/43 | 3.36E-04 | 1.16E-02 | JAG1;CDH1;EPHB3 |
| Regulation Of Viral Entry Into Host Cell (GO:0046596) | 3/44 | 3.60E-04 | 1.18E-02 | IFITM3;IFITM2;NECTIN2 |
| Regulation Of Melanin Biosynthetic Process (GO:0048021) | 2/10 | 4.32E-04 | 1.34E-02 | ZEB2;CDH3 |
| Interferon-Mediated Signaling Pathway (GO:0140888) | 3/49 | 4.95E-04 | 1.45E-02 | IFITM3;IFNAR2;IFITM2 |
| Sprouting Angiogenesis (GO:0002040) | 3/52 | 5.90E-04 | 1.65E-02 | EFNB2;EPHB4;ROBO1 |
| Synapse Organization (GO:0050808) | 4/131 | 7.77E-04 | 2.08E-02 | CDH1;PTK7;AGRN;PTPRF |
| Aorta Development (GO:0035904) | 2/14 | 8.67E-04 | 2.12E-02 | JAG1;ROBO1 |
| Cardiac Cell Development (GO:0055006) | 2/14 | 8.67E-04 | 2.12E-02 | CXADR;JAG1 |
| Protein Localization To Plasma Membrane (GO:0072659) | 4/137 | 9.18E-04 | 2.16E-02 | TSPAN14;CDH1;F11R;NECTIN3 |
| Axon Guidance (GO:0007411) | 4/149 | 1.25E-03 | 2.81E-02 | EFNB2;EFNB1;PTPRM;EPHB3 |
| Cell-Cell Junction Organization (GO:0045216) | 3/68 | 1.29E-03 | 2.81E-02 | CDH3;CXADR;CDH1 |
| Negative Regulation Of Viral Entry Into Host Cell (GO:0046597) | 2/18 | 1.45E-03 | 2.86E-02 | IFITM3;IFITM2 |
| Pulmonary Valve Morphogenesis (GO:0003184) | 2/18 | 1.45E-03 | 2.86E-02 | JAG1;ROBO1 |
| **GO_Cellular_Component (down)** | **Overlap** | **P-value** | **Adjusted P-value** | **Genes** |
| Cell-Cell Junction (GO:0005911) | 11/299 | 2.14E-09 | 1.59E-07 | CDH3;CXADR;JAG1;CDH1;PTK7;FAT1;PTPRM;CTNNB1;F11R;NECTIN3;NECTIN2 |
| Adherens Junction (GO:0005912) | 8/150 | 2.27E-08 | 8.39E-07 | CDH3;CXADR;JAG1;CDH1;PTPRM;CTNNB1;NECTIN3;NECTIN2 |
| Apical Junction Complex (GO:0043296) | 6/106 | 1.02E-06 | 2.52E-05 | CXADR;CDH1;AMOTL2;F11R;NECTIN3;NECTIN2 |
| Catenin Complex (GO:0016342) | 3/28 | 9.22E-05 | 1.71E-03 | CDH3;CDH1;CTNNB1 |
| Basement Membrane (GO:0005604) | 3/46 | 4.11E-04 | 6.08E-03 | LAMA5;LAMC1;AGRN |
| Lysosome (GO:0005764) | 7/503 | 1.00E-03 | 1.23E-02 | IFITM3;IFITM2;AP5M1;TMBIM1;AGRN;GPC6;VASN |
| Focal Adhesion (GO:0005925) | 6/387 | 1.35E-03 | 1.39E-02 | EFNB2;ADGRE5;PTK7;FAT1;CTNNB1;NECTIN2 |
| Cell-Substrate Junction (GO:0030055) | 6/395 | 1.50E-03 | 1.39E-02 | EFNB2;ADGRE5;PTK7;FAT1;CTNNB1;NECTIN2 |
| Bicellular Tight Junction (GO:0005923) | 3/81 | 2.13E-03 | 1.75E-02 | CXADR;AMOTL2;F11R |
| Tight Junction (GO:0070160) | 3/93 | 3.16E-03 | 2.34E-02 | CXADR;AMOTL2;F11R |
| Lytic Vacuole Membrane (GO:0098852) | 4/287 | 1.28E-02 | 8.58E-02 | IFITM3;IFITM2;TMBIM1;VASN |
| Zonula Adherens (GO:0005915) | 1/5 | 1.57E-02 | 8.91E-02 | NECTIN2 |
| Apicolateral Plasma Membrane (GO:0016327) | 1/5 | 1.57E-02 | 8.91E-02 | CXADR |
| Cortical Cytoskeleton (GO:0030863) | 2/63 | 1.69E-02 | 8.91E-02 | RIMS1;CDH1 |
| Flotillin Complex (GO:0016600) | 1/6 | 1.88E-02 | 9.25E-02 | CDH1 |

| | | | | |
|---|---|---|---|---|
| Cytoskeleton Of Presynaptic Active Zone (GO:0048788) | 1/7 | 2.18E-02 | 1.01E-01 | RIMS1 |
| Phosphatidylinositol 3-Kinase Complex, Class I (GO:0097651) | 1/8 | 2.49E-02 | 1.01E-01 | PIK3R5 |
| Phosphatidylinositol 3-Kinase Complex, Class IA (GO:0005943) | 1/8 | 2.49E-02 | 1.01E-01 | PIK3R5 |
| Lysosomal Membrane (GO:0005765) | 4/356 | 2.58E-02 | 1.01E-01 | IFITM3;IFITM2;TMBIM1;VASN |
| Collagen-Containing Extracellular Matrix (GO:0062023) | 4/373 | 3.00E-02 | 1.04E-01 | LAMA5;LAMC1;AGRN;ANGPTL1 |
| Lysosomal Lumen (GO:0043202) | 2/86 | 3.01E-02 | 1.04E-01 | AGRN;GPC6 |
| Neuron Projection (GO:0043005) | 5/557 | 3.08E-02 | 1.04E-01 | PRNP;CXADR;PTK7;ROBO1;EPHB3 |
| Specific Granule Membrane (GO:0035579) | 2/90 | 3.28E-02 | 1.05E-01 | TSPAN14;TMBIM1 |
| beta-catenin-TCF Complex (GO:1990907) | 1/12 | 3.72E-02 | 1.15E-01 | CTNNB1 |
| Golgi Lumen (GO:0005796) | 2/100 | 3.97E-02 | 1.17E-01 | AGRN;GPC6 |
| Proton-Transporting ATP Synthase Complex (GO:0045259) | 1/18 | 5.52E-02 | 1.51E-01 | ATP5MC1 |
| Mitochondrial Proton-Transporting ATP Synthase Complex (GO:0005753) | 1/18 | 5.52E-02 | 1.51E-01 | ATP5MC1 |
| Secretory Granule Membrane (GO:0030667) | 3/279 | 5.79E-02 | 1.53E-01 | TSPAN14;ADGRE5;TMBIM1 |
| Anaphase-Promoting Complex (GO:0005680) | 1/20 | 6.12E-02 | 1.56E-01 | ANAPC10 |
| **GO_Molecular_Function (down)** | **Overlap** | **P-value** | **Adjusted P-value** | **Genes** |
| Coreceptor Activity Involved In Wnt Signaling Pathway, Planar Cell Polarity Pathway (GO:1904929) | 2/5 | 9.71E-05 | 8.64E-03 | PTK7;GPC6 |
| Coreceptor Activity Involved In Wnt Signaling Pathway (GO:0071936) | 2/7 | 2.03E-04 | 9.03E-03 | PTK7;GPC6 |
| Cadherin Binding (GO:0045296) | 6/319 | 4.97E-04 | 1.45E-02 | CDH3;CDH1;PTPRM;CTNNB1;F11R;VASN |
| Ephrin Receptor Activity (GO:0005003) | 2/15 | 9.99E-04 | 1.45E-02 | EPHB4;EPHB3 |
| Transmembrane Receptor Protein Phosphatase Activity (GO:0019198) | 2/16 | 1.14E-03 | 1.45E-02 | PTPRM;PTPRF |
| Transmembrane Receptor Protein Tyrosine Phosphatase Activity (GO:0005001) | 2/16 | 1.14E-03 | 1.45E-02 | PTPRM;PTPRF |
| Ubiquitin-Protein Transferase Activator Activity (GO:0097027) | 2/16 | 1.14E-03 | 1.45E-02 | EFNB1;UBE2J1 |
| Ephrin Receptor Binding (GO:0046875) | 2/21 | 1.97E-03 | 2.20E-02 | EFNB2;EFNB1 |
| Transmembrane Receptor Protein Tyrosine Kinase Activity (GO:0004714) | 2/50 | 1.09E-02 | 1.07E-01 | EPHB4;EPHB3 |
| Axon Guidance Receptor Activity (GO:0008046) | 1/5 | 1.57E-02 | 1.31E-01 | PTK7 |
| PDZ Domain Binding (GO:0030165) | 2/62 | 1.64E-02 | 1.31E-01 | CXADR;F11R |
| Aspartic-Type Endopeptidase Inhibitor Activity (GO:0019828) | 1/6 | 1.88E-02 | 1.31E-01 | PRNP |
| G Protein-Coupled Glutamate Receptor Binding (GO:0035256) | 1/7 | 2.18E-02 | 1.31E-01 | PRNP |
| Calcium Ion Binding (GO:0005509) | 4/346 | 2.36E-02 | 1.31E-01 | MEX3B;CDH3;CDH1;AGRN |
| Protein Tyrosine Phosphatase Activity (GO:0004725) | 2/76 | 2.40E-02 | 1.31E-01 | PTPRM;PTPRF |
| Cuprous Ion Binding (GO:1903136) | 1/8 | 2.49E-02 | 1.31E-01 | PRNP |
| RNA Polymerase I Core Promoter Sequence-Specific DNA Binding (GO:0001164) | 1/8 | 2.49E-02 | 1.31E-01 | RRN3 |
| Protein Binding Involved In Heterotypic Cell-Cell Adhesion (GO:0086080) | 1/9 | 2.80E-02 | 1.31E-01 | CXADR |
| RNA Polymerase I Transcription Regulatory Region Sequence-Specific DNA Binding (GO:0001163) | 1/9 | 2.80E-02 | 1.31E-01 | RRN3 |
| alditol:NADP+ 1-Oxidoreductase Activity (GO:0004032) | 1/10 | 3.11E-02 | 1.38E-01 | AKR7A2 |
| Histone Methyltransferase Binding (GO:1990226) | 1/11 | 3.41E-02 | 1.42E-01 | CTNNB1 |
| LRR Domain Binding (GO:0030275) | 1/12 | 3.72E-02 | 1.42E-01 | ROBO1 |
| Solute:Monoatomic Cation Symporter Activity (GO:0015294) | 1/12 | 3.72E-02 | 1.42E-01 | SLC39A10 |
| I-SMAD Binding (GO:0070411) | 1/13 | 4.02E-02 | 1.42E-01 | CTNNB1 |
| 1-Phosphatidylinositol-3-Kinase Regulator Activity (GO:0046935) | 1/14 | 4.32E-02 | 1.42E-01 | PIK3R5 |
| Aldo-Keto Reductase (NADP) Activity (GO:0004033) | 1/14 | 4.32E-02 | 1.42E-01 | AKR7A2 |
| Transmembrane-Ephrin Receptor Activity (GO:0005005) | 1/14 | 4.32E-02 | 1.42E-01 | EPHB3 |
| Death Receptor Binding (GO:0005123) | 1/15 | 4.62E-02 | 1.47E-01 | TMBIM1 |
| Phosphatidylinositol 3-Kinase Regulator Activity (GO:0035014) | 1/17 | 5.22E-02 | 1.60E-01 | PIK3R5 |
| **MSigDB_Hallmark (down)** | **Overlap** | **P-value** | **Adjusted P-value** | **Genes** |
| Myogenesis | 4/200 | 3.65E-03 | 4.92E-02 | PRNP;SMTN;AGRN;EPHB3 |

| | | | | |
|---|---|---|---|---|
| Apical Junction | 4/200 | 3.65E-03 | 4.92E-02 | CDH3;CDH1;NECTIN3;NECTIN2 |
| Wnt-beta Catenin Signaling | 2/42 | 7.75E-03 | 6.98E-02 | JAG1;CTNNB1 |
| TGF-beta Signaling | 2/54 | 1.26E-02 | 8.49E-02 | CDH1;CTNNB1 |
| Cholesterol Homeostasis | 2/74 | 2.28E-02 | 1.14E-01 | JAG1;CTNNB1 |
| Interferon Gamma Response | 3/200 | 2.52E-02 | 1.14E-01 | IFITM3;IFNAR2;IFITM2 |
| Interferon Alpha Response | 2/97 | 3.75E-02 | 1.45E-01 | IFITM3;IFITM2 |
| UV Response Dn | 2/144 | 7.57E-02 | 2.55E-01 | PTPRM;LAMC1 |
| Apoptosis | 2/161 | 9.15E-02 | 2.59E-01 | IFITM3;CTNNB1 |
| Notch Signaling | 1/32 | 9.61E-02 | 2.59E-01 | JAG1 |
| Angiogenesis | 1/36 | 1.07E-01 | 2.64E-01 | JAG1 |
| Apical Surface | 1/44 | 1.30E-01 | 2.70E-01 | EPHB4 |
| IL-2/STAT5 Signaling | 2/199 | 1.30E-01 | 2.70E-01 | IFITM3;PRNP |
| Reactive Oxygen Species Pathway | 1/49 | 1.43E-01 | 2.77E-01 | PRNP |
| IL-6/JAK/STAT3 Signaling | 1/87 | 2.40E-01 | 4.33E-01 | PIK3R5 |
| Androgen Response | 1/100 | 2.71E-01 | 4.58E-01 | UBE2J1 |
| DNA Repair | 1/150 | 3.78E-01 | 4.70E-01 | BCAM |
| TNF-alpha Signaling via NF-kB | 1/200 | 4.70E-01 | 4.70E-01 | JAG1 |
| Estrogen Response Late | 1/200 | 4.70E-01 | 4.70E-01 | CDH1 |
| Complement | 1/200 | 4.70E-01 | 4.70E-01 | PIK3R5 |
| mTORC1 Signaling | 1/200 | 4.70E-01 | 4.70E-01 | ATP5MC1 |
| Epithelial Mesenchymal Transition | 1/200 | 4.70E-01 | 4.70E-01 | LAMC1 |
| Inflammatory Response | 1/200 | 4.70E-01 | 4.70E-01 | PIK3R5 |
| Oxidative Phosphorylation | 1/200 | 4.70E-01 | 4.70E-01 | ATP5MC1 |
| Glycolysis | 1/200 | 4.70E-01 | 4.70E-01 | AGRN |
| heme Metabolism | 1/200 | 4.70E-01 | 4.70E-01 | BCAM |
| Allograft Rejection | 1/200 | 4.70E-01 | 4.70E-01 | IFNAR2 |
