## Supplemental Table S2 for "ER-to-Golgi Trafficking is a Nutrient-Sensitive Checkpoint Linking Glucose Starvation to Cell Surface Remodeling"

### Table S2. ULK1-regulated phosphorylation sites on SEC31: motif scores, conservation, and mass spectrometry results.

This supplemental table summarizes the analysis of phosphorylation sites on SEC31A, with an emphasis on ULK1-dependent regulation. ULK1 motif-matching scores (PSSM) and evolutionary conservation across vertebrate SEC31A were determined using a motif prediction program[84, 85]. Phosphorylation status was evaluated by mass spectrometry in two experimental contexts: (1) overexpressed EYFP-tagged SEC31A isolated from cells co-expressing empty vector (EV), wild-type (WT) ULK1, or kinase-dead (KD) ULK1; and (2) endogenous FLAG-tagged SEC31A from cells cultured in Dulbecco's Modified Eagle Medium with or without glucose for 1 h to model nutrient stress. By integrating motif prediction, conservation, and empirical phosphorylation data, this data highlights candidate ULK1-regulated sites on SEC31A and their sensitivity to cellular nutrient stress. Although several ULK1-dependent phosphorylation sites (identified in the overexpression context) exhibit high PSSM scores, S352 is the only site for which mass spectrometry detected phosphorylation on endogenous SEC31A. Abbreviations: D, detected; ND, not detected.

Column Descriptions:

*All values reflect changes in cell surface protein abundance as measured by TMT-based proteomics following NHS-biotinylation and*

| Column Heading | Description |
|---|---|
| Site | Phosphorylation site on SEC31A, indicated by residue and position (e.g., S352). |
| Peptide | Amino acid sequence of the phosphopeptide detected by mass spectrometry containing the indicated site. |
| PSSM | Motif-matching score (Position-Specific Scoring Matrix) indicating the similarity of the site to the ULK1 consensus phosphorylation motif. |
| Conservation (Cons) | Degree of evolutionary conservation of the phosphorylation site among vertebrate SEC31A |
| Coverage (Cov) | Indicates whether the peptide containing the site was detected by mass spectrometry in the specified experimental condition. If multiple "Coverage" columns are present, each corresponds to a specific condition (e.g., EV, WT ULK1, KD ULK1, +Glucose, –Glucose). |
| Phosphorylation (Phosph) | Indicates whether phosphorylation at the specific site was detected by mass spectrometry in the corresponding experimental condition. Multiple columns correspond to the respective conditions. |
| Flag | Categorical annotation for each phosphorylation site: 1 = phosphorylation is ULK1-dependent + endogenous phosphorylation is detected; 2 = phosphorylation is ULK1-dependent; endogenous phosphorylation is not detected; 3 = phosphorylation is not ULK1-dependent + endogenous phosphorylation is detected; 4 = no specific pattern of phosphorylation; 5 = inconsistent coverage; 6 = not covered by mass spectrometry. |

| EYFP SEC31A+WT ULK1 | | | | | | | | | | Endogenous SEC31A | | | | Flag |
|---|---|---|---|---|---|---|---|---|---|---|---|---|---|---|
| | | | | EV | | WT ULK1 | | KD ULK1 | | Control | | Starved | | |
| Site | Peptide | PSSM | Cons | Cov | Phosph | Cov | Phosph | Cov | Phosph | Cov | Phosph | Cov | Phosph | |
| S352 | DKLSSSFGNL | 2.62 | 100% | + | ND | + | D | + | ND | + | D | + | D | 1 |
| S337 | SIMGGSTDGL | 5.63 | 100% | + | ND | + | D | + | ND | + | ND | + | ND | 2 |
| S332 | RISVYSIMGG | 4.15 | 100% | + | ND | + | D | + | ND | + | ND | + | ND | 2 |
| T62 | MKSCATFSSS | 3.12 | 100% | + | ND | + | D | + | ND | + | ND | + | ND | 2 |
| S208 | PIIKVSDHSN | 1.70 | 100% | + | ND | + | D | + | ND | + | ND | + | ND | 2 |
| S323 | VLSAASFDGR | 1.29 | 100% | + | ND | + | D | + | ND | + | ND | + | ND | 2 |
| S351 | VDKLSSSFGN | 1.03 | 100% | + | ND | + | D | + | ND | + | ND | + | ND | 2 |
| S430 | HHVFISQVVT | 1.02 | 100% | + | ND | + | D | + | ND | + | ND | + | ND | 2 |
| S712 | KAQDGSHPLS | 0.15 | 100% | + | ND | + | D | + | ND | + | ND | + | ND | 2 |
| S442 | EFLSRSDQLQ | -0.38 | 100% | + | ND | + | D | + | ND | + | ND | + | ND | 2 |
| S521 | VALKDSDQVA | -0.38 | 100% | + | ND | + | D | + | ND | + | ND | + | ND | 2 |
| S682 | ENEGDSLLQT | -2.99 | 100% | + | ND | + | D | + | ND | + | ND | + | ND | 2 |
| S329 | FDGRISVYSI | 4.78 | 86% | + | ND | + | D | + | ND | + | ND | + | ND | 2 |
| T741 | AMDTSTVGVL | -3.48 | 86% | + | ND | + | D | + | ND | + | ND | + | ND | 2 |
| S1168 | SPTITSGLHN | -2.99 | 57% | + | ND | + | D | + | ND | + | ND | + | ND | 2 |
| S740 | QAMDTSTVGV | 7.31 | 43% | + | ND | + | D | + | ND | + | ND | + | ND | 2 |
| T1161 | KLREQTLSPT | 0.22 | 100% | + | D | + | D | + | D | + | D | + | D | 3 |
| S1163 | REQTLSPTIT | -0.92 | 100% | + | D | + | D | + | D | + | D | + | D | 3 |
| S532 | SDGEESPAAE | -3.72 | 100% | + | D | + | D | + | D | + | D | + | D | 3 |
| S799 | VAGHESPKIP | -3.45 | 86% | + | D | + | D | + | D | + | D | + | D | 3 |
| S527 | DQVAQSDGEE | -4.11 | 71% | + | D | + | D | + | D | + | D | + | D | 3 |
| S1194 | HTHIVSTSNF | 4.58 | 100% | + | ND | + | ND | + | D | + | ND | + | ND | 4 |
| S397 | RPVGASFSFG | 3.97 | 100% | + | ND | + | D | + | D | + | ND | + | ND | 4 |
| T1165 | QTLSPTITSG | 3.23 | 100% | + | ND | + | ND | + | D | + | ND | + | ND | 4 |
| T1190 | GLTMHTHIVS | 2.16 | 100% | + | ND | + | D | + | D | + | ND | + | ND | 4 |
| S186 | QHILASASPS | 1.48 | 100% | + | ND | + | ND | + | D | + | ND | + | ND | 4 |
| S1196 | HIVSTSNFSE | 0.20 | 100% | + | ND | + | ND | + | D | + | ND | + | ND | 4 |
| S399 | VGASFSFGGK | -0.88 | 100% | + | ND | + | ND | + | D | + | ND | + | ND | 4 |
| T165 | TPGAKTQPPE | -3.57 | 100% | + | ND | + | ND | + | D | + | ND | + | ND | 4 |
| T1187 | YSEGLTMHTH | 3.75 | 86% | + | D | + | ND | + | D | + | ND | + | ND | 4 |
| T1195 | THIVSTSNFS | 0.05 | 86% | + | ND | + | ND | + | D | + | ND | + | ND | 4 |
| S414 | NVRMPSHQGA | 1.31 | 57% | + | ND | + | ND | + | D | + | ND | + | ND | 4 |
| S751 | LAAKMSQYAN | 3.01 | 100% | | | + | ND | | | | | | | 5 |
| S465 | KKIDASQTEF | 1.63 | 100% | | | + | D | | | | | | | 5 |
| S440 | EKEFLSRSDQ | 1.04 | 71% | | | + | D | | | | | | | 5 |
| S564 | GTFNISVSGD | 4.97 | 100% | | | | | | | | | | | 6 |
| T36 | QQLDATFSTN | 4.89 | 100% | | | | | | | | | | | 6 |
| S65 | CATFSSSHRY | 4.58 | 100% | | | | | | | | | | | 6 |
| S918 | ISSASSYTGQ | 4.57 | 100% | | | | | | | | | | | 6 |
| S621 | KYFAKSQSKI | 4.29 | 100% | | | | | | | | | | | 6 |
| S66 | ATFSSSHRYH | 3.76 | 100% | | | | | | | | | | | 6 |
| T194 | PSGRATVWDL | 3.50 | 100% | | | | | | | | | | | 6 |
| S986 | SELPASQRTG | 3.38 | 100% | | | | | | | | | | | 6 |
| S475 | EKNVWSFLKV | 2.48 | 100% | | | | | | | | | | | 6 |
| S269 | LAIAWSMADP | 2.42 | 100% | | | | | | | | | | | 6 |
| S172 | PPEDISCIAW | 2.36 | 100% | | | | | | | | | | | 6 |
| S217 | NRMHCSGLAW | 2.32 | 100% | | | | | | | | | | | 6 |
| S278 | PELLLSCGKD | 2.32 | 100% | | | | | | | | | | | 6 |
| T1122 | LILKTTFEDL | 2.11 | 100% | | | | | | | | | | | 6 |
| T936 | QASSPTSSPA | 1.88 | 100% | | | | | | | | | | | 6 |
| T303 | ELPTNTQWCF | 1.85 | 100% | | | | | | | | | | | 6 |
| T1121 | HLILKTTFED | 1.70 | 100% | | | | | | | | | | | 6 |
| S965 | APPSSSAYAL | 1.61 | 100% | | | | | | | | | | | 6 |
| S64 | SCATFSSSHR | 1.52 | 100% | | | | | | | | | | | 6 |
| S963 | PGAPPSSSAY | 1.52 | 100% | | | | | | | | | | | 6 |
| S104 | ILYDPSKIIA | 1.43 | 100% | | | | | | | | | | | 6 |
| S1176 | HNIARSIETR | 1.37 | 100% | | | | | | | | | | | 6 |
| T560 | PSSGGTFNIS | 1.34 | 100% | | | | | | | | | | | 6 |
| S584 | TGNFESAVDL | 1.23 | 100% | | | | | | | | | | | 6 |
